# Discovering cellular programs of intrinsic and extrinsic drivers of metabolic traits using LipocyteProfiler

**DOI:** 10.1101/2021.07.17.452050

**Authors:** Samantha Laber, Sophie Strobel, Josep-Maria Mercader, Hesam Dashti, Alina Ainbinder, Julius Honecker, Garrett Garborcauskas, David R. Stirling, Aaron Leong, Katherine Figueroa, Nasa Sinnott-Armstrong, Maria Kost-Alimova, Giacomo Deodato, Alycen Harney, Gregory P. Way, Alham Saadat, Sierra Harken, Saskia Reibe-Pal, Hannah Ebert, Yixin Zhang, Virtu Calabuig-Navarro, Elizabeth McGonagle, Adam Stefek, Josée Dupuis, Beth A. Cimini, Hans Hauner, Miriam S. Udler, Anne E. Carpenter, Jose C. Florez, Cecilia M. Lindgren, Suzanne B. R. Jacobs, Melina Claussnitzer

**Author notes:** shared lead authors (Samantha Laber, Sophie Strobel).

## Abstract

A primary obstacle in translating genetics and genomics data into therapeutic strategies is elucidating the cellular programs affected by genetic variants and genes associated with human diseases. Broadly applicable high-throughput, unbiased assays offer a path to rapidly characterize gene and variant function and thus illuminate disease mechanisms. Here, we report LipocyteProfiler, an unbiased high-throughput, high-content microscopy assay that is amenable to large-scale morphological and cellular profiling of lipid-accumulating cell types. We apply LipocyteProfiler to adipocytes and hepatocytes and demonstrate its ability to survey diverse cellular mechanisms by generating rich context-, and process-specific morphological and cellular profiles. We then use LipocyteProfiler to identify known and novel cellular programs altered by polygenic risk of metabolic disease, including insulin resistance, waist-to-hip ratio and the polygenic contribution to lipodystrophy. LipocyteProfiler paves the way for large-scale forward and reverse phenotypic profiling in lipid-storing cells, and provides a framework for the unbiased identification of causal relationships between genetic variants and cellular programs relevant to human disease.

## Introduction

With the rise of human genome sequencing data, the number of genetic loci and variants known to be associated with human diseases has increased substantially; however, elucidating the pathogenic mechanisms through which genetic variants impact disease largely remains a non-systematic, labor- and cost-intensive endeavor that is biased towards hypotheses drawn from *a priori* knowledge. Phenotypic profiling is a powerful tool to systematically discover external and internal regulators of biological processes in cellular systems in an unbiased manner <u>(Neumann et al. 2010; Green et al. 2011; Feng et al. 2009; Hughes et al. 2021).</u> In particular, high-content imaging is an established multi-parametric approach that captures and quantifies numerous distinct biological processes from microscopy images, yielding a rich set of morphological and cellular profiles (Bray et al. 2016). To date, imaging-based profiling has been used in small molecule screens to identify compound fingerprints, ascertain compound toxicity, predict compounds’ assay activity <u>(Kepiro et al. 2018; Scheeder et al. 2018; Simm et al. 2018; Wawer et al. 2014)</u>, and in gene expression screens to annotate gene function <u>(Rohban et al. 2017)</u>. In all cases, the basic strategy is to match the profile of a given sample based on similarity to morphological profiles of already-annotated samples. Thus far the phenotypic data ascertained from scalable morphological profiling assays has been limited to features detectable by generic cellular organelle stains <u>(Bray et al. 2016</u>) -- such as structural information about nuclei, endoplasmic reticulum (ER), cytoskeleton and mitochondria -- or generic processes, such as cell growth or proliferation. It is currently unknown how gene and compound effects translate to changes in specific cellular pathways and processes. As such, there is a pressing need to develop foundational technologies that allow systematically linking genetic variation to disease-relevant cellular programs at scale.

In metabolism, lipid droplets represent a highly-relevant feature that is amenable to image-based profiling. Lipid droplets are storage organelles central to both whole body metabolism and energy homeostasis. These droplets are highly dynamic, and found in all cell types <u>(Olzmann and Carvalho 2019) either as part of cellular homeostasis in lipocytes (lipid-accumulating cells), such as adipocytes, hepatocytes, macrophages/foam cells and glial cells (Liu et al. 2015; Olzmann and Carvalho 2019; Wang et al. 2013; Grandl and Schmitz 2010; Robichaud et al. 2021), or as part of pathophysiological processes in cells such as vascular smooth muscle cells, skeletal muscle cells, renal podocytes, and cancer cells (Hershey et al. 2019; Cruz et al. 2020; Wang et al. 2005; Weinert et al. 2013; Prats et al. 2006). Changes in lipid droplet dynamics -- such as the number and size of lipid droplets, and overall lipid content -- are associated with the progression of numerous metabolic diseases, including type 2 diabetes (T2D), obesity, and non-alcoholic fatty liver disease (NAFLD).</u>

Here, we introduce LipocyteProfiler, a metabolic disease-oriented phenotypic profiling system for lipid-accumulating cells that bridges the gap between high-throughput general assays and low-throughput, highly customized, disease-focused readouts. LipocyteProfiler is an adaption of Cell Painting (<u>Gustafsdottir et al. 2013</u>; <u>Bray et al. 2016</u>) that incorporates BODIPY to measure dynamic features of lipid droplets, and thus captures lipocyte-relevant phenotypes in addition to generic morphological profiles. We prototyped LipocyteProfiler in adipocytes and hepatocytes, which are highly specialized cells that store excess energy in the form of lipid droplets and have key roles in metabolic disease. First, we demonstrate that LipocyteProfiler can identify meaningful changes in feature profiles (a) during adipocyte differentiation, (b) across white and brown adipocyte lineages, and (c) following genetic and drug perturbations. Next, we correlated LipocyteProfiles with transcriptomic data from RNAseq to link gene sets with morphological and cellular features, capturing a broad range of cellular activity in differentiating adipocytes. We then applied LipocyteProfiler to connect polygenic risk scores for type 2 diabetes (T2D)-related traits to cellular phenotypes, and discover novel trait-specific cellular mechanisms underlying polygenic risk. Finally, we used our method to uncover cellular traits under the genetic control of an individual genetic risk locus, as shown for the *2p23.3* metabolic risk locus at *DNMT3A* <u>(Tachmazidou et al. 2017)</u>. We demonstrated that our customized morphometric approach is capable of identifying diverse cellular mechanisms by generating context-, process-, and allele-specific morphological and cellular profiles.

## Results

### LipocyteProfiler creates meaningful morphological and cellular profiles in differentiating adipocytes

To quantitatively map dynamic and context-dependent morphological and cellular signatures in lipocytes as well as to discover intrinsic and extrinsic drivers of cellular programs, we developed a high-content, image-based profiling approach called LipocyteProfiler (Figure 1a). LipocyteProfiler expands on the Cell Painting protocol <u>(Bray et al. 2016)</u> by replacing the ER stain with BODIPY, which stains intracellular lipid droplets composed of neutral lipids. The result is an unbiased high-throughput profiling assay that generates rich generic and lipocyte-specific cellular profiles from six multiplexed fluorescent dyes imaged in four channels (Figure 1b) in conjunction with an automated image analysis pipeline (see Methods for more details). LipocyteProfiler extracts 3,005 morphological and cellular features that map to three cellular compartments (Cell, Cytoplasm, and Nucleus) across four channels differentiating the organelles, namely *DNA* (Hoechst), *Mito* (MitoTracker Red which stains mitochondria), *AGP* (actin, Golgi, plasma membrane; stained with Phalloidin (F-actin cytoskeleton) and Wheat Germ Agglutinin (Golgi and plasma membranes)), and *Lipid* (BODIPY, which stains neutral lipids, multiplexed with SYTO14, which stains nucleoli and cytoplasmic RNA) (Figure 1c).

**Figure 1.**
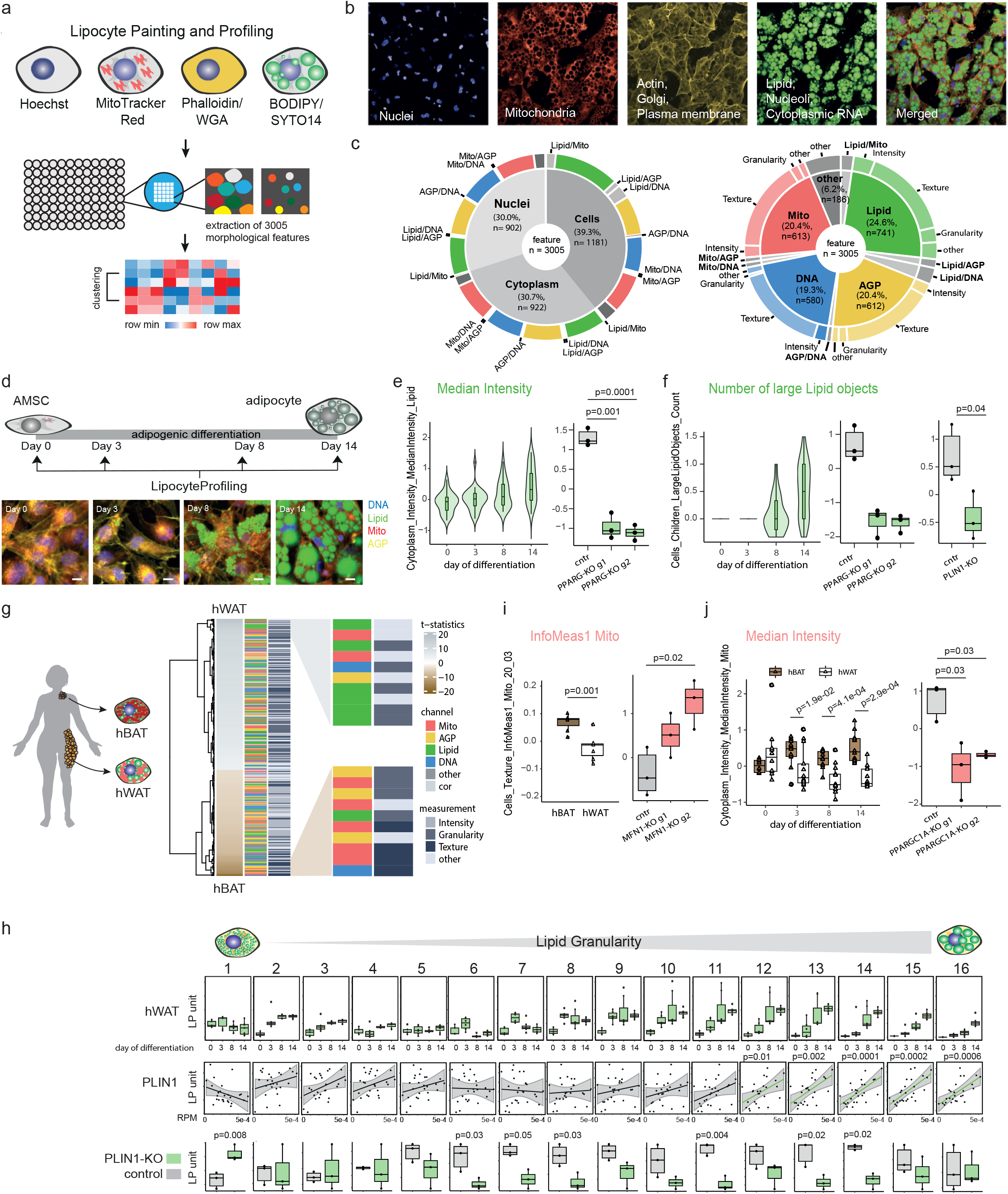
LipocyteProfiler creates rich morphological and cellular profiles in adipocytes that are informative for known cellular functions. (a) Schematic of LipocyteProfiler, which is a high-content imaging assay that multiplexes six fluorescent stains imaged in four channels in conjunction with an automated image analysis pipeline to generate rich morphological and cellular profiles in lipid-storing cell types, such as adipocytes during differentiation. (b) Representative microscope image of fully differentiated adipocytes for four individual channels and a merged representation across channels. (c) LipocyteProfiler extracts 3,005 morphological and cellular features that map equally to three cellular compartments and across four channels. (d) Schematic of LipocyteProfiling in differentiating hWAT at four time points of adipocyte differentiation (day 0, 3, 8, 14). Representative images of AMSCs stained using LipocytePainting at 4 time points of differentiation (day 0, 3, 8, 14). Scale bar = 10um. (e) *Lipid MedianIntensity*, a measurement of lipid content within a cell, significantly increases with adipogenic differentiation and decreases following CRISPR/Cas9-mediated knockdown of *PPARG* in differentiated white adipocytes. Data are shown for two guides used (g1 and g2) and Y-axis shows LP units (normalized LipocyteProfiling (LP) values across three batches, see Methods). (f) *Number of large Lipid objects* informative for large lipid droplets are absent in the progenitor state (day 0) and in early differentiation (day 3) and progressively increase in later stages of differentiation (day 8 and 14). *Number of large Lipid objects* are reduced following CRISPR/Cas9-mediated KO of *PPARG* (data are shown for two guides used (g1 and g2)) and *PLIN1,* at day 14 of differentiation. Y-axis shows LP units (normalized LP values across three batches, see Methods). (g) Morphological profiles of white (hWAT) and brown (hBAT) adipocytes at day 14 of differentiation differ significantly across all feature classes (FDR<0.1%). Features are clustered based on effect size. Features with the highest effect size in hWAT and hBAT adipocytes are lipid- and mitochondria-related, respectively. Graph shows zoom-in for top 10 features with largest effect sizes in hWAT (top panel) and hBAT (bottom panel). (h) *Lipid Granularity* measures, as a spectra of 16 lipid droplet size measures, show size-specific changes in hWAT and hBAT during differentiation. See also Figure S1e Granularity features informative for larger lipid droplets (*Lipid Granularity 10*-*16*) correlate positively with *PLIN1* gene expression and are reduced in *PLIN1*-KO adipocytes. See also Figures S1f-g (PLIN2, FASN-KO). Y-axis shows autoscaled LP units (normalized LP values across three batches, see Methods) (i) Brown adipocytes (hBAT) show higher *Mito*_*Texture_InfoMeas1*, a measure of spatial relationship between specific intensity values, compared to white adipocytes (hWAT). CRISPR/Cas9-mediated knockout of *MFN1*, a mitochondrial fusion gene, changes *Mito*_*Texture_InfoMeas1* (data is shown for two guides used (g1 and g2). Y-axis shows LP units (normalized LP values across three batches (hBAT/hWAT) or normalized across CRISPR-KO data, see Methods). (j) *Mito MedianIntensity* is higher in brown (hBAT) compared to white (hWAT) adipocytes throughout differentiation and decreased after CRISPR/Cas9-mediated knockout of *PPARGC1A* in hWAT. Y-axis shows LP units (normalized LP values across three batches, see Methods).

We tested the ability of LipocyteProfiler to detect i) changes associated with adipocyte differentiation, ii) differences between white and brown adipocytes, and iii) phenotypic effects of directed gene perturbation using CRISPR/Cas9 to knockout key regulators of adipocyte function.

First, we applied LipocyteProfiler to a model of adipocyte differentiation using an established white adipocyte line (hWAT) <u>(Xue et al. 2015)</u>, which undergoes phenotypic changes from fibroblast-shaped to spherical lipid-filled cells during differentiation (days 0, 3, 8, 14; Figure 1d). We mapped the phenotypic signature of progressive lipid accumulation and cytoskeletal remodelling during adipocyte differentiation in hWAT using tractable *Lipid* and *AGP* features. We show that cytoplasmic intensity of *Lipid*, a proxy of overall lipid content within a cell, increased with adipogenic differentiation (Figure 1e). In addition, *large Lipid objects* (large lipid droplets) were absent in the progenitor state (day 0) and in early differentiation (day 3), and increase in later stages of differentiation (Figure 1f). We confirmed that CRISPR/Cas9-directed perturbation of *PPARG*, the master regulator of adipogenesis, decreases the overall *Lipid* intensity in differentiated white adipocytes (guide 1 *p=*1.0x10^-3^, guide 2 *p=*1.0x10^-4^; Figure 1e). Further, *large Lipid objects* present at day 14 of differentiation were reduced when we perturb regulators of lipid accumulation, *PPARG* (guide 1 *p*=4.0x10^-3^, guide 2 *p=*6.0x10^-3^) and *PLIN1* (*p*=4.0x10^-2^), a key regulator of lipid droplet homeostasis (Figure 1f). These data demonstrate that LipocyteProfiler detects expected changes in lipid dynamics associated with adipocyte differentiation. Another cellular change that occurs during adipocyte differentiation is a drastic reorganization of the actin cytoskeleton, which transitions from well-defined stress fibers in pre-adipocytes, to relatively thick cortical actin lining composed of patches of punctate F-actin at the inner surface of the plasma membrane in fully differentiated adipocytes <u>(Kanzaki 2006)</u> (Figure S1a). This cytoskeletal remodelling is stimulated by insulin and essential for GLUT4 translocation into the membrane to facilitate insulin-responsive glucose uptake in the cell <u>(Kanzaki 2006)</u>. Concordantly, we found that CRISPR/Cas9-mediated disruption of the insulin receptor (*INSR*) and insulin receptor substrate 1 (*IRS1*) in pre-adipocytes altered *AGP Texture* features (describing the smoothness of a given stain) in mature adipocytes at day 14 of differentiation (Figure S1b). Specifically, *INSR*- and *IRS*-knock-out reduced variation of cytoplasmic *AGP* stain intensities most significantly near the plasma membrane (*Cytoplasm_RadialDisribution_RadialCV_AGP_4_of_4, IRS1* guide 1 *p=*0.002, guide 2 *p=*0.03*; INSR* guide 1 *p=*0.02, guide 2 *p=*0.005), indicative of less punctuated *AGP*, which is in line with less cortical actin in *INSR*- and *IRS-*knock-out cells.

Next, we use brown and white adipocyte model systems to elucidate mitochondrial and lipid-related informational content. Intrinsic differences distinguishing white and brown adipocytes are known to be predominantly driven by changes in mitochondrial number and activity that translate into differential lipid accumulation <u>(Cedikova et al. 2016)</u>. Using an established brown adipocyte line derived from human neck fat (hBAT) from the same individual as for the hWAT line, we showed that morphological profiles from differentiated hWAT and hBAT differ significantly in every channel and feature category (Figure 1g). *Lipid Granularity* measures, a class of metrics that capture the typical sizes of bright spots for a stain, predominated among those increased in hWAT. During adipocyte differentiation, lipid droplets typically increase first in number and then enlarge and fuse to form larger lipid droplets over the course of maturation <u>(Fei et al. 2011)</u>. We observed that the number of small and medium sized lipid droplets (*Lipid Granularity* measures 1-9) present in early differentiating hWAT saturate in early stages of differentiation (Figure 1h). Larger lipid droplets (*Lipid Granularity* measures 10-16) increase in terminal differentiation, indicating that lipid droplets form in early differentiation and grow in size thereafter, a process that is reflected in *Lipid Granularity* measures and *Lipid objects* count (Figure 1h, Figure S1c). Consistent with the notion that adipocytes from brown adipose tissue have smaller lipid droplets, we found that during differentiation, hBAT generally accumulate fewer medium and large lipid droplets as seen by lower values across the spectra of granularity (Figure 1h, Figure S1e). Intuitively, LipocyteProfiler-derived size estimates showed that white hWAT are larger than brown hBAT adipocytes after 14 days of adipogenic differentiation as cells become lipid-laden (*Cells_AreaShape_Area; p*=5.1x10^-5^; Figure S1d). To test if lipid droplet-associated perilipins can be linked to lipid droplet sizes, we correlated *Lipid Granularity* measures with mRNA expression levels of *PLIN1*, which is specifically expressed in adipocytes where it directs the formation of large lipid droplets <u>(Shijun et al. 2020; Gandotra et al. 2011)</u> and *PLIN2,* the only constitutively and ubiquitously expressed lipid droplet protein which is associated with a range of lipid droplets in diverse cell types <u>(Brasaemle et al. 1997; Tsai et al. 2017)</u>. We observed that mRNA expression levels of *PLIN1* positively correlated with the *Lipid Granularity* features informative for larger spot sizes (*Lipid Granularity measures 12-16*) (Figure 1h). *PLIN2* correlated best with *Lipid Granularity* measures of smaller and larger size spectra (Figure S1f). Accordingly, when we knocked out *PLIN1 and FASN*, genes involved in lipid droplet dynamics and lipid metabolism, we observed a size-specific reduction of *Lipid Granularity* (Figure 1h, Figure S1g), suggesting that *Lipid Granularity* features are a suitable output measure of lipid droplet size spectra and an indicator of adipocyte differentiation.

Consistent with the relevance of mitochondria for brown adipocyte function, mitochondrial measures were among the features that increased the most in hBAT (Figure 1g), particularly the *Texture* feature *Cells*_*Texture_InfoMeas1_Mito* (describing the overall information content based on the smoothness of a given stain). Perturbation of *MFN1*, a mitochondrial fusion gene, increased *Cells*_*Texture_InfoMeas1_Mito* in hWAT adipocytes (guide 2 *p*=0.02; Figure 1i), suggesting that the higher values of this measurement in differentiated hBAT could be indicative of higher mitochondrial fission in hBAT compared to hWAT. This finding is consistent with brown adipocytes elevating mitochondrial thermogenesis by increasing mitochondrial fission <u>(Gao and Houtkooper 2014).</u> hBAT adipocytes are further characterized by increased *Mito Intensity* compared to hWAT adipocytes throughout differentiation, with the most substantial increase in the fully differentiated state (Median, day3 *p*=0.02, day8 *p*=0.0004, day14 *p*=0.0003; Figure 1j), demonstrating that LipocyteProfiler can identify known cellular programs that distinguish different adipocyte lineages. Indeed, when we perturbed *PPARGC1A*, the master regulator of mitochondrial biogenesis and thermogenesis in adipocytes, using CRISPR-Cas9 mediated knockout in hWAT, mitochondrial intensity decreased (guide 1 *p*=0.03, guide 2 *p=*0.03; Figure 1j). Taken together our data demonstrate that LipocyteProfiler is capable of generating rich sets of morphological and cellular features that correlate with cellular function.

### LipocyteProfiler identifies distinct depot-specific morphological and cellular signatures associated with differentiation trajectories in both visceral and subcutaneous adipocytes

We next used LipocyteProfiler to distinguish phenotypes of primary human adipose-derived mesenchymal stem cells (AMSCs) derived from the two main adipose tissue depots in the body, namely subcutaneous and visceral, across the course of differentiation (Figure 2a). We differentiated subcutaneous and visceral AMSCs and generated morphological profiles at days 0, 3, 8, and 14 using LipocyteProfiler and validated successful differentiation in both depots by an increase of adipogenesis marker genes (*LIPE*, *PPARG, PLIN1, GLUT4*) (Figure S2a). Concomitantly, we used RNA-sequencing to profile the transcriptome on the same differentiation timepoints. We observed that both the morphological and transcriptomic profiles show time course-specific signatures revealing a differentiation trajectory; however only morphological profiles generated by LipocyteProfiler also resolved adipose depot-specific signatures throughout differentiation (Figure 2b). At day 14 of differentiation, morphological differences between subcutaneous and visceral adipocytes were spread across a large number of features in all feature classes (Figure 2c, Figure S2b). To discover patterns associated with progression through adipocyte differentiation in each depot we performed a sample progression discovery analysis (SPD) <u>(Qiu et al. 2011)</u>. SPD clusters samples to reveal their underlying progression, and simultaneously identifies subsets of features that show the same progression pattern and are illustrative of differentiation. We discovered that subsets of features distinguish the differentiation patterns of subcutaneous and visceral adipocytes, and that most dominant feature classes were dynamically changing over the time course of differentiation (Figure 2d). In visceral adipocytes, the early phase of differentiation was predominantly associated with mitochondrial features whereas terminal phases of differentiation were primarily associated with changes in lipid-related features (Figure 2d). In subcutaneous adipocytes, we observed that the feature classes (actin-cytoskeleton, lipid, mitochondrial, and nucleic-acid) were more evenly involved through adipogenesis and that the contribution of *Lipid* features started in early phases of differentiation, consistent with an earlier initiation of lipid accumulation in subcutaneous compared to visceral adipocytes (Figure 2d).

**Figure 2.**
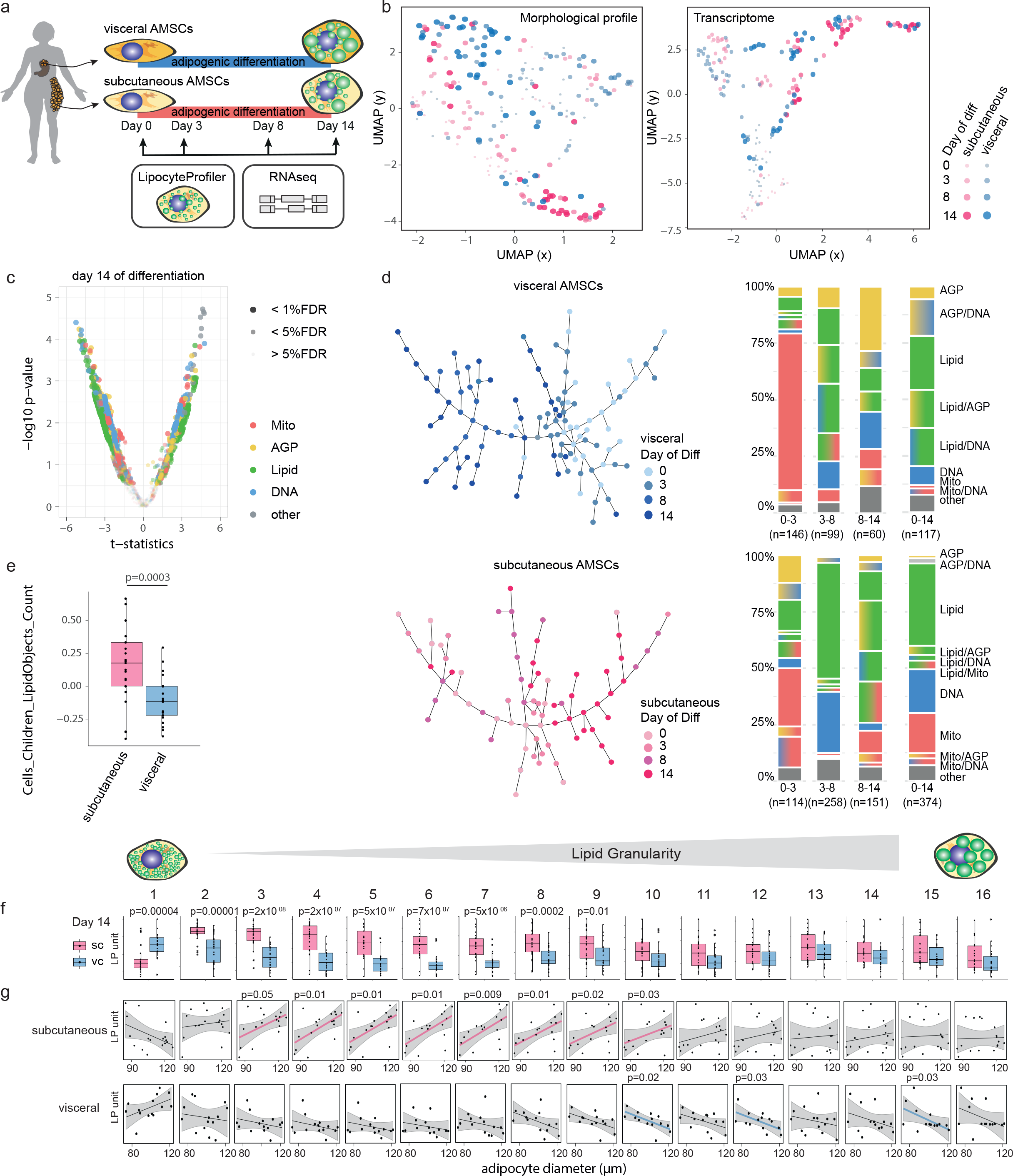
LipocyteProfiler identifies distinct depot-specific morphological and cellular signatures associated with differentiation trajectories in both visceral and subcutaneous AMSCs. (a) Human AMSCs isolated from subcutaneous and visceral adipose depots were differentiated for 14 days, and LipocyteProfiler and RNAseq-based profiling was performed throughout adipocyte differentiation (day 0, 3, 8, and 14). (b) LipocyteProfiler and transcriptome profiles show time course-specific signatures revealing a differentiation trajectory, but only LipocyteProfiler additionally resolves adipose depot-specific signatures. (c) Subcutaneous and visceral AMSCs at terminal differentiation (day 14) have distinct morphological and cellular profiles with differences that are spread across all channels. See also Figure S2b (day 0, 3, and 8). (d) Sample progression discovery analysis (SPD). Proportions of subgroups of features characterizing differentiation differ between subcutaneous and visceral adipocytes and dynamically change over the course of differentiation. In both depots, *Mito* features drive differentiation predominantly in the early phase of differentiation (day 0 to 3) whereas *Lipid* features predominate in the terminal phases (day 8 to 14). (e) The number of lipid droplets is higher in subcutaneous AMSCs compared to visceral AMSCs at terminal differentiation. Y-axis shows LP units (normalized LP values across eight batches, see methods). (f) Mature subcutaneous AMSCs have larger intracellular lipid droplets compared to visceral AMSCs at day 14 of differentiation (*Lipid Granularity*). Y-axis shows autoscaled LP units (normalized LP values across eight batches, see Methods). (g) *Lipid Granularity* from subcutaneous AMSCs at day 14 of differentiation correlates positively with floating mature adipocyte diameter, but shows an inverse relationship for visceral adipose tissue, suggesting distinct cellular mechanisms that lead to adipose tissue hypertrophy in these two depots. Y-axis shows autoscaled LP units (normalized LP values across eight batches; x-axis, histology adipocytes diameter (µm), see Methods).

We next compared lipid-related signatures in mature AMSCs and observed that subcutaneous AMSCs had more lipid droplets compared to visceral AMSCs (*Cells_LipidObject_count,* Figure 2e, *p*=2.9x10^-4^). More specifically, mature subcutaneous AMSCs showed significantly higher *Lipid Granularity* of small-to medium-sized lipid objects, whereas visceral adipocytes showed higher *Lipid Granularity* of very small lipid objects, suggesting that mature subcutaneous AMSCs have larger intracellular lipid droplets compared to visceral AMSCs which present higher abundance of very small lipid droplets (Figure 2f). These apparent intrinsic differences in differentiation capacities and lipid accumulation between subcutaneous and visceral AMSCs are in line with previously described distinct properties of AMSCs from those depots across differentiation <u>(Baglioni et al. 2012)</u>. Our data suggests that LipocyteProfiler can facilitate identifying those distinct lineage differences and programs of cellular differentiation.

Lastly, to assess the *in vivo* relevance of morphological features of *in vitro* differentiated adipocytes, we correlated *Lipid Granularity* features of adipocytes at day 14 of differentiation with diameter estimates of tissue-derived mature adipocytes from the same individual (see Methods). We showed that changes of *Lipid Granularity* of *in vitro* differentiated female subcutaneous adipocytes correlated significantly with the mean diameter of mature adipocytes (Figure 2g). More specifically, medium size granularity measures increased with larger *in vivo* size estimates, suggesting that *in vivo* adipocyte size is reflected by medium-sized lipid droplets in subcutaneous adipocytes that have been differentiated *in vitro*. Strikingly, we found the opposite effect between correlation of visceral *Lipid Granularity* and diameter estimates from mature adipocytes, suggesting that subcutaneous and visceral adipose tissues differ in cellular programs that govern depot-specific adipose tissue expansion, which may account for different depot-specific susceptibility to metabolic diseases. Indeed, white adipose depots have been reported to differ in their respective mechanisms of fat mass expansion under metabolic challenges, with subcutaneous adipose tissue being more capable of hyperplasia whereas visceral adipose tissue expands mainly via hypertrophy <u>(Wang et al. 2013)</u>.

### LipocyteProfiles reflect transcriptional states in adipocytes

To identify relevant processes that manifest in morphological and cellular features, and to identify pathways of a given set of features, we next used a linear mixed model to link the expression of 52,170 genes derived from RNAseq with each of the 2,760 image-based LipocyteProfiler features in subcutaneous adipocytes at day 14 of differentiation across 26 individuals (see Methods). We found 44,736 non-redundant significant feature-gene connections (FDR<0.1) that were composed of 10,931 genes and 869 features (FDR<0.01, 20,296 non-redundant feature-gene connections, 7,012 genes and 669 features (Figure S2c)), and mapped across all channels (Figure 3a). Although features from every channel had significant gene correlations, *Lipid* features showed the highest number of gene connections compared to any other channel. This suggests that lipid droplet structure, localization, and dynamics in adipocytes most closely represent the transcriptional state of the differentiated cell (Figure 3b). Pathway enrichment analyses of feature-gene connections added support that genes that correlated with a particular feature were biologically meaningful. For example *Mito Granularity* associated with a list of genes that were enriched more than expected by chance for biological pathways of tricarboxylic acid cycle (TCA) which oxidizes acetyl-CoA in mitochondria <u>(Kusminski and Scherer 2012)</u>, and *Lipid Intensity* in the cell associated with a gene list significantly enriched in biological pathways of oxidative phosphorylation (OXPHOS) and beta-oxidation (WikiPathway 368 and WikiPathway 143). Similarly, *Lipid Granularity* in the cytoplasm was enriched for genes involved in adipogenesis, apoptosis, and differentiation of white and brown adipocytes. Finally, correlation features (measures capturing the overlap between lipid droplets, mitochondria, and AGP) were enriched for cytoplasmic ribosomal protein and genes in the beta-oxidation pathway (Figure 3b; Table S1).

**Figure 3.**
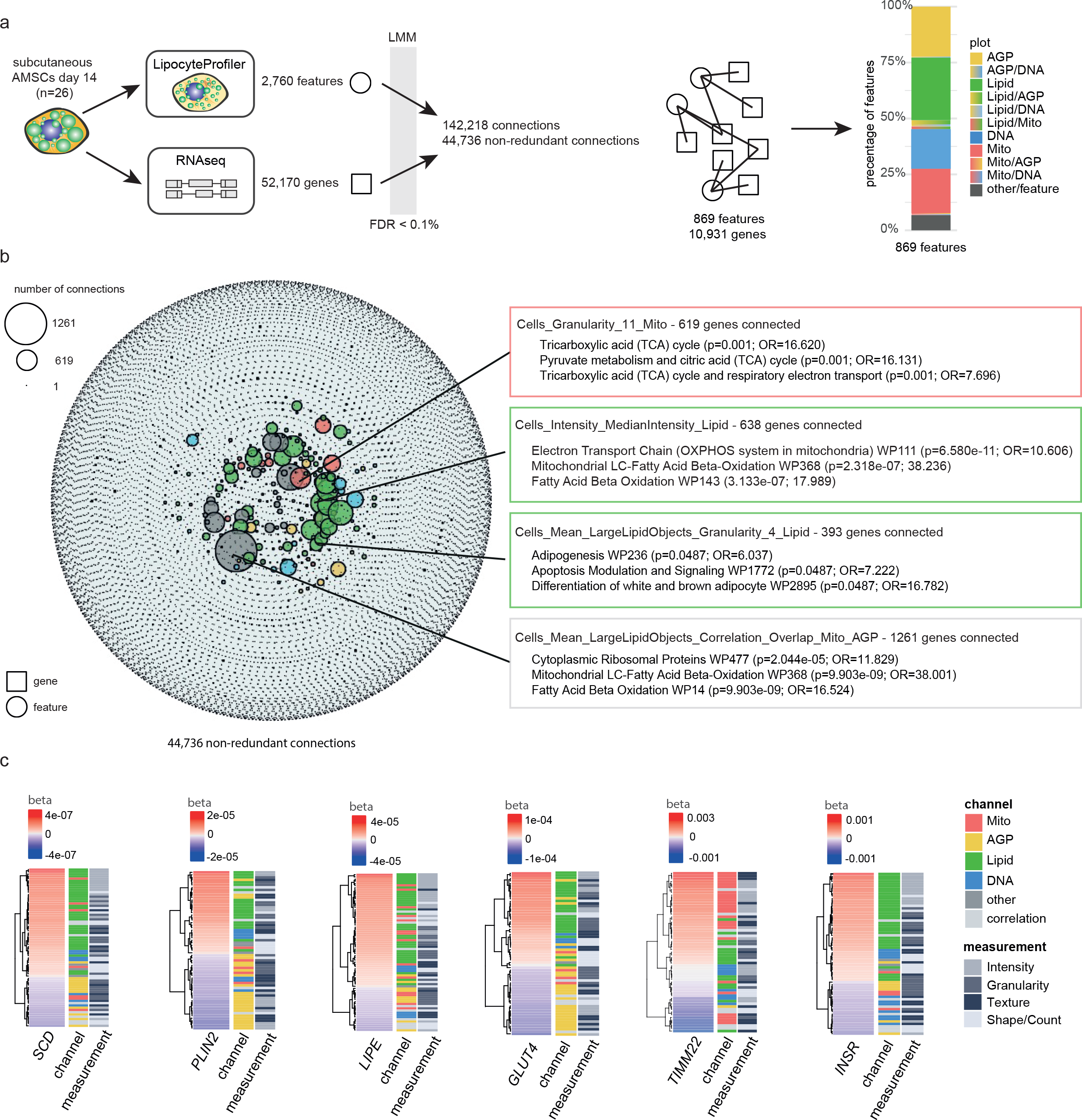
Correlations between morphological and transcriptional profiles. (a) Linear mixed model (LMM) was applied to correlate 2,760 morphological features derived from LipocyteProfiler with 52,170 transcripts derived from RNAseq in matched samples of subcutaneous AMSCs at terminal differentiation (day 14). With FDR < 0.1%, we discover 44,736 non-redundant connections that map to 869 morphological features and 10,931 genes. (b) Network of transcript-LipocyteProfiler feature correlations. Genes correlated to individual LipocyteProfiler features are enriched for relevant pathways. Node size is determined by number of connections Figure S2c (FDR<0.01%) (c) LipocyteProfiler signatures of adipocyte marker genes *SCD, PLIN2, LIPE, GLUT4, TIMM22, and INSR* recapitulate their known cellular function. Features are clustered based on beta of linear regression.

In addition to examining genes connected to particular feature groups, we also explored morphological features connected to specific genes. We found that morphological signatures of *SCD, PLIN2, LIPE, GLUT4*, *TIMM22,* and *INSR* revealed their known cellular functions (Figure 3c; Table S2). For example, the expression of *TIMM22,* a mitochondrial membrane gene, was most strongly correlated with *Mito Texture.* Expression of the insulin receptor (*INSR*) most strongly correlated with *Lipid Intensity* features indicative of lipid accumulation. *PLIN2* and *GLUT4* showed the highest positive and negative correlations with *Lipid* and *AGP* features, respectively. Together these data show that the mechanistic information gained from LipocyteProfiles is not limited to generic cellular organelles but reflects the transcriptional state of the cell and can be deployed to gain relevant mechanistic insights.

### LipocyteProfiler identifies cellular processes affected by drug perturbations in adipocytes and hepatocytes

To investigate whether LipocyteProfiler is capable of identifying effects of drug perturbations on morphological and cellular profiles, we first compared subcutaneous and visceral adipocytes that had been stimulated with the ß-adrenergic agonist isoproterenol (Figure 4a). Isoproterenol is known to induce lipolysis and increase mitochondrial energy dissipation <u>(Miller et al. 2015).</u> We observed that visceral adipocytes (n=4 individuals) responded to 24-hour isoproterenol treatment by changes in *Lipid* and *Mito* features (Figure 4b, Table S3). More specifically, we observed that isoproterenol-treated visceral adipocytes had increased *Mito Intensity* (*p*=0.04) and differences in mitochondrial *Texture Entropy* (*p*=0.009), indicative of a less-smooth appearance of mitochondrial staining compared to DMSO-treated controls (Figure 4c) and suggesting that isoproterenol treatment results in more hyperpolarized and fragmented mitochondria, which is a reported mechanism of norepinephrine-stimulated browning in adipocytes <u>(Gao and Houtkooper 2014)</u>. Isoproterenol-treated visceral adipocytes are further characterized by decreased *Lipid Intensity* (*p*=0.04) and *Lipid Texture Entropy* (*p*=0.03) (Figure 4d) as well as decreased *Lipid Granularity* across the full granularity size spectra, particularly at the smallest lipid droplet sizes (Figure 4e). This pattern suggests less overall lipid content in isoproterenol-treated visceral adipocytes, which may be due to increased lipolysis. Finally, the phenotypic response following isoproterenol treatment was predominant in visceral adipocytes, as we did not observe a significant effect in subcutaneous adipocytes (Figure S3). Indeed, adrenergic induced lipolysis is observed to be higher in visceral than subcutaneous in overweight and obese individuals <u>(Hoffstedt et al. 1997; Morigny et al. 2021)</u>.

**Figure 4.**
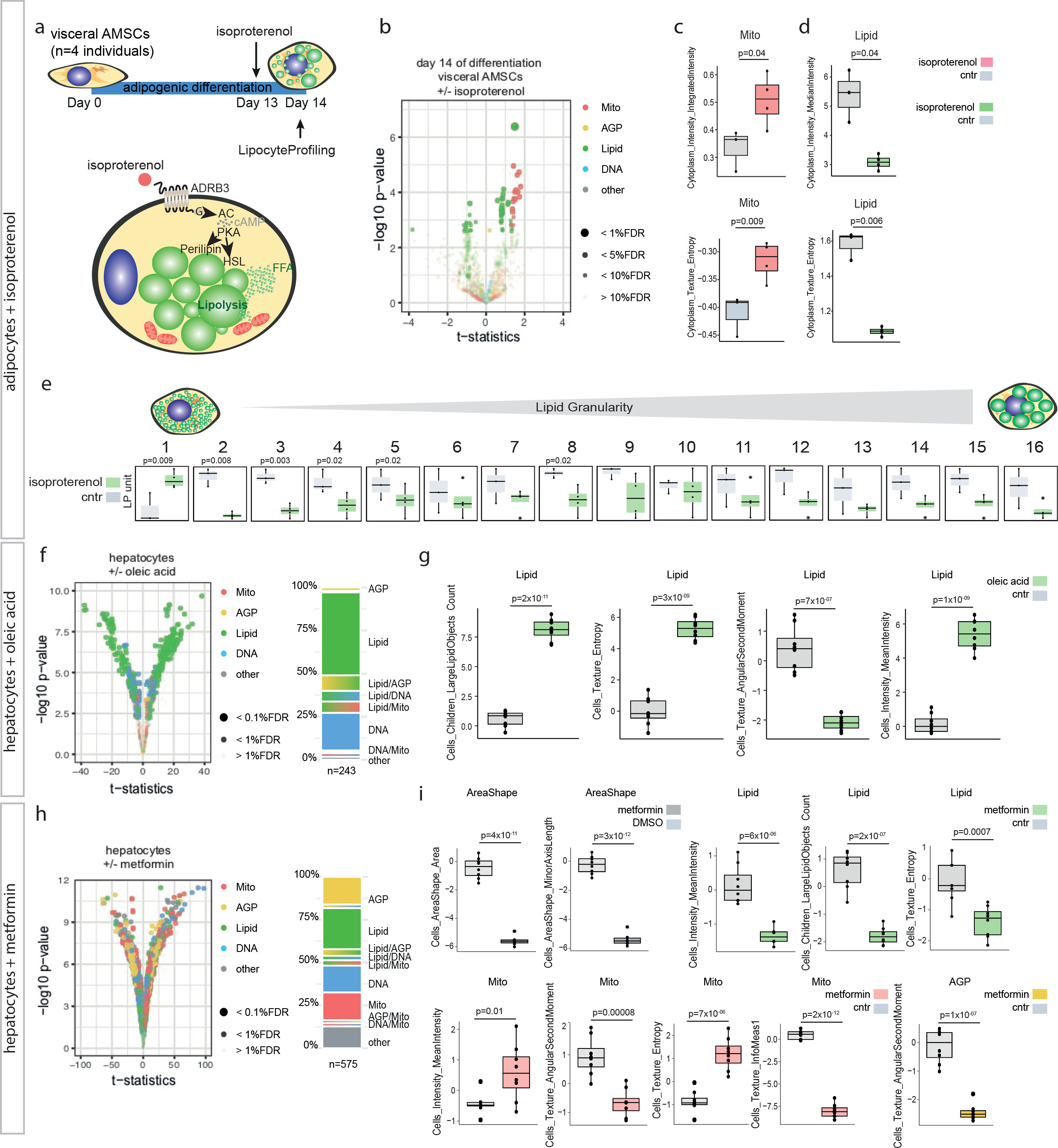
LipocyteProfiler identifies molecular mechanisms of drug stimulations in adipocytes and hepatocytes. (a) LipocyteProfiler was performed in visceral AMSCs (n=4) treated with the beta-adrenergic receptor agonist isoproterenol for 24 hours. (b) Isoproterenol treatment results in changes of lipid-related and mitochondrial traits in visceral AMSCs at day 14 of differentiation. See also Figure S3 (subcutaneous). (c-d) Isoproterenol treatment of visceral AMSC increases *Mito Intensity* and *Mito Texture Entropy*, while decreasing the respective *Lipid* features. Y-axis shows LP units (normalized LP values across eight batches, see Methods). (e) Isoproterenol treatment reduces lipid droplet sizes measured via *Lipid Granularity.* Y-axis shows autoscaled LP units (normalized LP values across eight batches, see Methods). (f) Oleic acid treatment in hepatocytes results in changes of lipid-related features. (g) Oleic acid treatment in hepatocytes affects lipid-related morphological features suggestive of increased lipid droplet size and number. Y-axis shows LP units (normalized LP values across PHH data, see Methods). (h) Metformin treatment in hepatocytes results in global changes affecting features across all channels. (i) Metformin effect in hepatocytes is suggestive of increased mitochondrial activity, while lipid droplet size and number are reduced. Metformin-treated hepatocytes are also smaller and show reduced cytoskeletal randomness. Y-axis shows LP units (normalized LP values across PHH data, see Methods).

To test LipocyteProfiler application to cell types beyond adipocytes we assayed the effects of oleic acid and metformin in primary human hepatocytes (PHH). Consistent with the finding that free fatty acid treatment induces lipid droplet accumulation in PHH <u>(Liu et al. 2014)</u>, our results showed that 24h treatment of PHH with oleic acid (OA) yielded predominantly *Lipid* feature changes in the cell (Figure 4f, Table S3), with a morphological profile indicative of increased lipid droplet number (*LargeLipidObjects_Count*, *p*=2.3x10^-11^) and overall lipid content (*Cells_MeanIntensity_Lipid*, *p*=1.3x10^-09^) as well as differences in *Texture* (*Cells_Texture_Entropy_Lipid*, *p*=2.6x10^-09^; *Cells_Texture_AngularSecondMoment_Lipid*, *p*=7.2x10^-07^; Figure 4g). By contrast, treatment of PHH with 5mM metformin for 24h caused morphological and cellular changes that were spread across all channels (Figure 4h, Table S3), with a profile suggestive of smaller cells (*Cells*_*AreaShape_Area*, *p*=3.9x10^-11^; *Cells*_*AreaShape_MinorAxisLength*, *p*=3.1x10^-12^) with increased mitochondrial membrane potential (*Cells_MeanIntensity_Mito*, *p*=0.01), and mitochondrial heterogeneity (*Cells_Texture_AngularSecondMoment_Mito*, *p*=7.6x10^-05^; *Cells_Texture_Entropy_Mito*, *p*=6.7x10^-06^; *Cells_Texture_InfoMeas1_Mito*, *p*=2.2x10^-12^). Additionally, we observed reduced lipid content (*Cells_MeanIntensity_Lipid*, *p*=6.1x10^-06^), reduced lipid droplet number (*LargeLipidObjects_Count*, *p*=2.3x10^-07^) and differences in *Texture* (*Cells_Texture_Entropy_Lipid*, *p*=7.0x10^-04^) (Figure 4i). This concerted effect of metformin on mitochondrial structure and function as well as lipid-related features is consistent with a less uniform appearance of the cytoskeleton, Golgi, and plasma membrane in metformin-treated hepatocytes compared to control (*Cells_Texture_AngularSecondMoment_AGP*, *p*=10x10^-08^). Indeed, prolonged treatment with high doses of metformin leads to mitochondrial uncoupling, resulting in mitochondrial hyperpolarization and diminished lipid accumulation in PHH <u>(Liu et al. 2014; Forkink et al. 2014; Demine et al. 2019).</u> Together, these data demonstrate that morphological and cellular profiles of drug perturbation in lipocytes yield cellular signatures reflecting known biology and drug action in a single concerted snapshot of cell behavior.

### Polygenic risk effects for insulin resistance affects lipid degradation in differentiated visceral adipocytes

Next, we used LipocyteProfiler to discover cellular programs of metabolic polygenic risk in adipocytes. For systematic profiling of AMSCs in the context of natural genetic variation (Table S4), we first assessed the effect of both technical and biological variance on LipocyteProfiler features. To obtain a measure of batch-to-batch variance associated with our experimental set-up, we differentiated hWAT, hBAT, and SGBS preadipocytes <u>(Fischer-Posovszky et al. 2008)</u> in three independent experiments and found no significant batch effect (BEscore 0.0047, 0.0001, 0.0003, Figure S4a). We also showed that the accuracy of predicting cell type is substantially higher than predicting batch (Figure S4a), indicating that our LipocyteProfiler framework can detect intrinsic versus extrinsic variance in our dataset with low batch effect and high accuracy. Secondly, we performed a variance component analysis across all data collected on adipocyte morphological and cellular traits across 65 donor-derived differentiating AMSCs to assess the contribution of intrinsic genetic variation compared to the contribution of other possible confounding factors such as batch, T2D status, age, sex, BMI, cell density, and passage number. In total, we found that across all samples and batches, the largest contributor to feature variance was donor ID, accounting for 17.03% (interquartile range 11.45%–21.95%) of variance (Figure S4b). Other factors appeared to contribute only marginally to overall variance of the data, including batch effect (6.02%, 3.94%–8.84%) and plating density (3.75%, 1.55%–5.61%). These data suggest that LipocyteProfiler is able to detect and distinguish inter-individual genetic feature variation to a similar degree as reported for human iPSCs, where quantitative assays of cell morphology demonstrated a donor contribution to inter-individual variation in the range of 8-23% <u>(Kilpinen et al. 2017)</u>. To account for the variable feature-specific contributions of batch, sex, age, and BMI to overall feature variance, we corrected for those covariables in our analyses. Together, these data suggest that AMSCs-derived LipocyteProfiles can be used to study the effect of genetic contributions to morphological and cellular programs.

To ascertain the effect of polygenic risk for metabolic disease on cellular programs, we used the latest genome-wide association studies (GWAS) summary statistics for T2D. We constructed individual genome-wide polygenic risk scores (PRSs) for three T2D-related traits that have been linked to adipose tissue: T2D <u>(Mahajan et al. 2018)</u>, insulin resistance by homeostasis measure assessment (HOMA-IR; <u>(Matthews et al. 1985) (Dupuis et al. 2010)), and waist-to-hip ratio adjusted for BMI (WHRadjBMI) (Pulit et al. 2019). We selected donors from the bottom and top 25^th^ percentiles of these genome-wide PRSs (referred to as low and high polygenic risk, respectively) and compared LipocyteProfiles across the time course of visceral and subcutaneous adipocyte differentiation in high and low polygenic risk groups for each of the traits (</u>Figure 5a).

**Figure 5.**
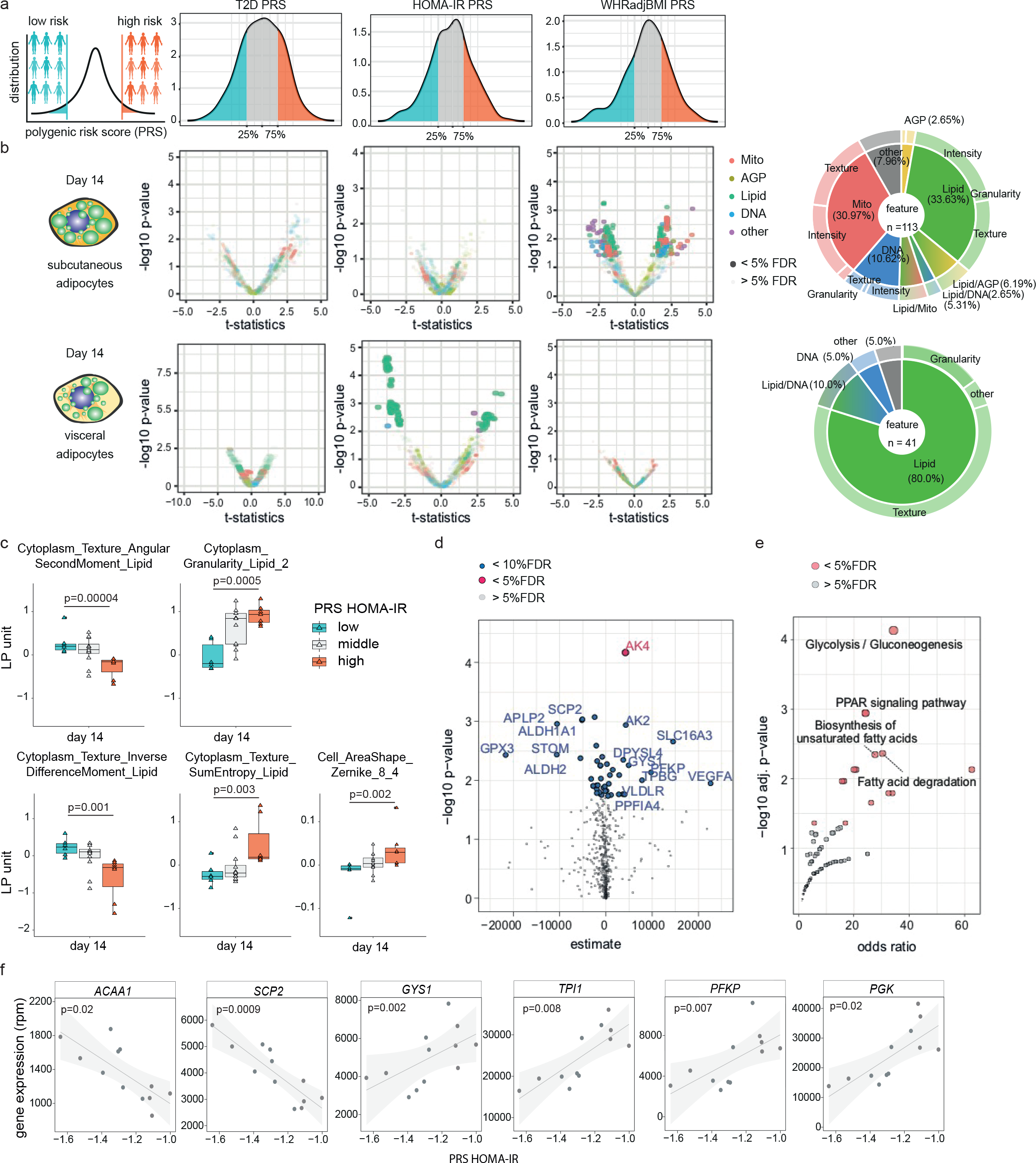
Polygenic risk effects for insulin resistance affects lipid degradation in differentiated visceral adipocytes. (a) Donors from the bottom and top 25 percentiles of genome-wide polygenic risk scores for three T2D-related traits (HOMA-IR, T2D, WHRadjBMI) were selected to compare LipocyteProfiles across the time course of visceral and subcutaneous adipocyte differentiation. (b) LipocyteProfiler applied to visceral and subcutaneous differentiating adipocytes reveals trait-specific polygenic effects on image-based cellular and morphological signatures for HOMA-IR in terminally differentiated visceral AMSCs (day 14; largely *Lipid* features) and WHRadjBMI in subcutaneous AMSCs at day 14 of differentiation (largely *Mito* and *Lipid* features), but no effect for T2D. See also Figure S5a, e (day 0, 3, and 8). (c) HOMA-IR polygenic risk in visceral AMSCs manifested in altered *Lipid Texture*, *Lipid Granularity* and *Cell shape* features, resembling an inhibition of lipolysis. Y-axis shows LP units (normalized LP values across eight batches, see Methods). See also Figure 4d (isoproterenol stimulation) (d-f) Correlation of gene expression of 512 genes known to be involved in adipocyte function with HOMA-IR PRS showed that genes that correlated with HOMA-IR PRSs were enriched for biological processes related to glucose metabolism, fatty acid transport, degradation and lipolysis (KEGG Pathways 2019).

We found significant polygenic effects on image-based cellular signatures for HOMA-IR and WHRadjBMI, but no effect for T2D (Figure 5b, Figure S5a,e; Table S5). More specifically, we observed an effect of HOMA-IR polygenic risk on morphological and cellular profiles at day 14 in visceral adipocytes (41 features, FDR<5%, Figure 5b), indicating a spatiotemporal and depot-specific effect of polygenic risk for insulin resistance. The features that differed between the high and low HOMA-IR PRS carriers were mostly *Lipid* features (Figure 5b). Visceral adipocytes from high polygenic risk individuals showed, in the cytoplasm, increased *Lipid Granularity* (*p*=0.0005, Figure 5c), increased *Cytoplasm_Texture_SumEntropy_Lipid* (*p*=0.003, Figure 5c), increased *Cells_AreaShape_Zernike_8_4* (*p*=0.009, Figure 5c), decreased *Cytoplasm_Texture_InverseDifferenceMoment_Lipid* (*p*=0.002, Figure 5c), and reduced *Cytoplasm_Texture_AngularSecondMoment_Lipid* (*p*=0.00005, Figure 5c) compared to low polygenic risk individuals. These data indicate that visceral adipocytes from individuals with high compared to low polygenic risk for insulin resistance are characterized by a lipid-associated morphological profile, driven by key features informative for increased number of small- to medium-sized lipid droplets, less homogenous lipid droplet distribution, and larger adipocytes, indicating excessive lipid accumulation in visceral adipocytes from high polygenic risk individuals. Notably, the pattern that contrasts between high and low polygenic risk individuals recapitulates signatures that resemble an inhibition of lipolysis and lipid degradation, as demonstrated by the inverse direction of effect in isoproterenol-stimulated visceral AMSCs as shown in Figure 4. More specifically, we observed that HOMA-IR increases the number of small lipid droplets in visceral adipocytes, which are precisely the features affected in response to isoproterenol (Figure S5b and Figure 4e). Together, these image-derived rich representations of morphological and cellular signatures capture a cellular program that is characterized by a metabolic switch towards lipid accumulation rather than lipolysis in visceral adipocytes derived from individuals at high polygenic risk for insulin resistance.

To further resolve the cellular program underlying HOMA-IR PRSs in visceral adipocytes, we integrate image-based information from LipocyteProfiler with an additional phenotypic modality, RNA-Seq data from the same donor-derived samples to ascertain the effects of polygenic risk for HOMA-IR on gene expression. Looking at a subset of mRNA levels for 512 genes known to be involved in adipocyte differentiation and function (GSEA hallmark gene sets for adipogenesis, fatty acid metabolism and glycolysis <u>(Subramanian et al. 2005; Mootha et al. 2003)</u>), we identified 51 genes under the polygenic control of HOMA-IR (FDR <10%) in fully differentiated visceral adipocytes (Figure 5d, Table S6). Genes correlating with the HOMA-IR PRS were enriched for biological processes related to glucose metabolism, fatty acid transport, degradation, and lipolysis (Figure 5e, Table S7). Negatively correlated genes include *ACAA1* (*p*=0.02) and *SCP2* (*p*=0.0009) (Figure 5f), consistent with an inhibition of lipolysis and lipid degradation in visceral adipocytes from individuals at high polygenic risk for HOMA-IR. Positively correlated genes include *GYS1,* which is a regulator of glycogen biosynthesis shown to causally link glycogen metabolism to lipid droplet formation in brown adipocytes <u>(Mayeuf-Louchart et al. 2019)</u> (*p*=0.005, Figure 5f). Additionally, multiple critical enzymes of the glycolysis pathway (*TPI1* (*p*=0.008), *PFKP* (*p*=0.007), *PGK* (*p*=0.02), Figure 5f), and marker genes of energy metabolism (*AK2* and *AK4* (Figure S5d)) were positively correlated suggesting a metabolic switch from lipolytic degradation of triglycerides to glycolytic activity. Although a causal link between visceral adipose mass and insulin resistance has been widely observed <u>(Lebovitz and Banerji 2005)</u>, the mechanism behind this observation is not understood. Together, orthogonal evidence from both high-content image-based and RNAseq-based profiling experiments in subcutaneous and visceral AMSCs suggests that individuals with high polygenic risk for HOMA-IR are characterized by a block of lipid degradation in visceral adipocytes.

### Polygenic risk for lipodystrophy-like phenotype manifests in cellular programs that indicate reduced lipid accumulation capacity in subcutaneous adipocytes

To resolve polygenic effects on adipocyte cellular programs beyond heterogeneous T2D and insulin resistance traits, we used the clinically informed and process-specific partitioned PRS of lipodystrophy, which are enriched for enhancer activity in adipocytes <u>(Udler et al. 2018)</u>, and correlated this score with morphological features throughout adipocyte differentiation. The lipodystrophy PRS was constructed based on 20 T2D-associated loci with a lipodystrophy-like phenotype, signifying insulin resistance with a lower BMI (Figure 6a). We found patients with high vs. low polygenic risk of lipodystrophy had distinctive morphological features in the *Mito*, *AGP*, and *Lipid* categories in their subcutaneous AMSCs at day 14 of differentiation, whereas mostly *Lipid* features were associated with increased lipodystrophy PRS in mature visceral AMSCs (Figure 6b-d, Figure S6a, Table S8), again highlighting a depot-specific and spatiotemporal-dependent effect of polygenic risk on morphological profiles captured with LipocyteProfiler. The patterns that contrast between high and low polygenic risk carriers include key features informative for increased mitochondrial membrane potential (e.g. *Intensity_MeanIntensity_Mito p*=0.0075, *Intensity_MaxIntensityEdge_Mito p*=0.005; Table S8), changes to the actin cytoskeleton indicating decreased cortical actin at the plasma membrane (e.g. *Cells_RadialDistribution_FracAtD_AGP ring 2 of 4 p=*0.012 *and 4 of 4 p=*0.0091, Figure S6b), and decreased lipid accumulation in subcutaneous adipocytes (e.g. *Intensity_IntegratedIntensityEdge_Lipid p*=0.0121, *Intensity_LowerQuartileIntensity_Lipid p*=0.0048; Table S8). Strikingly, representative images of subcutaneous adipocytes derived from individuals at the tail ends of lipodystrophy PRS (top compared to bottom 25^th^ percentiles) confirm that adipocytes from high PRS carriers have increased mitochondrial stain intensity - indicating of higher mitochondrial membrane potential <u>(Xiao et al. 2016)</u> - accompanied by smaller lipid droplets on average compared to adipocytes from individuals with low PRS (Figure 6c). We also note that CRISPR/Cas9 mediated knock-out of the monogenic familial partial lipodystrophy gene *PLIN1*, maps to features informative for decreased number of medium- and large-sized lipid droplets (Figure 1h), matching the polygenic risk effect. To assess whether the identified morphological and cellular changes underlying lipodystrophy polygenic risk resemble cellular drivers of monogenic forms of lipodystrophy more broadly, we next correlated expression of marker genes of monogenic familial partial lipodystrophy syndromes (*PPARG*, *LIPE*, *PLIN1*, *AKT2*, *CIDEC*, *LMNA,* and *ZMPSTE24)* with LipocyteProfiler features across subcutaneous adipocytes from 26 individuals. We found similar morphological signatures between profiles from monogenic lipodystrophy-associated genes and the polygenic lipodystrophy profile, with high effect sizes of *Mito* and *AGP* features (Figure S6c). These results suggest that polygenic and monogenic forms of lipodystrophy converge on similar cellular mechanisms involving increased mitochondrial activity, and decreased lipid accumulation in subcutaneous adipocytes from high PRS donors. This finding is consistent with the fact that different monogenic forms of lipodystrophy (independent of the mutation) showed similar consequences on mitochondrial OXPHOS in patient samples <u>(Sleigh et al. 2012)</u>.

**Figure 6.**
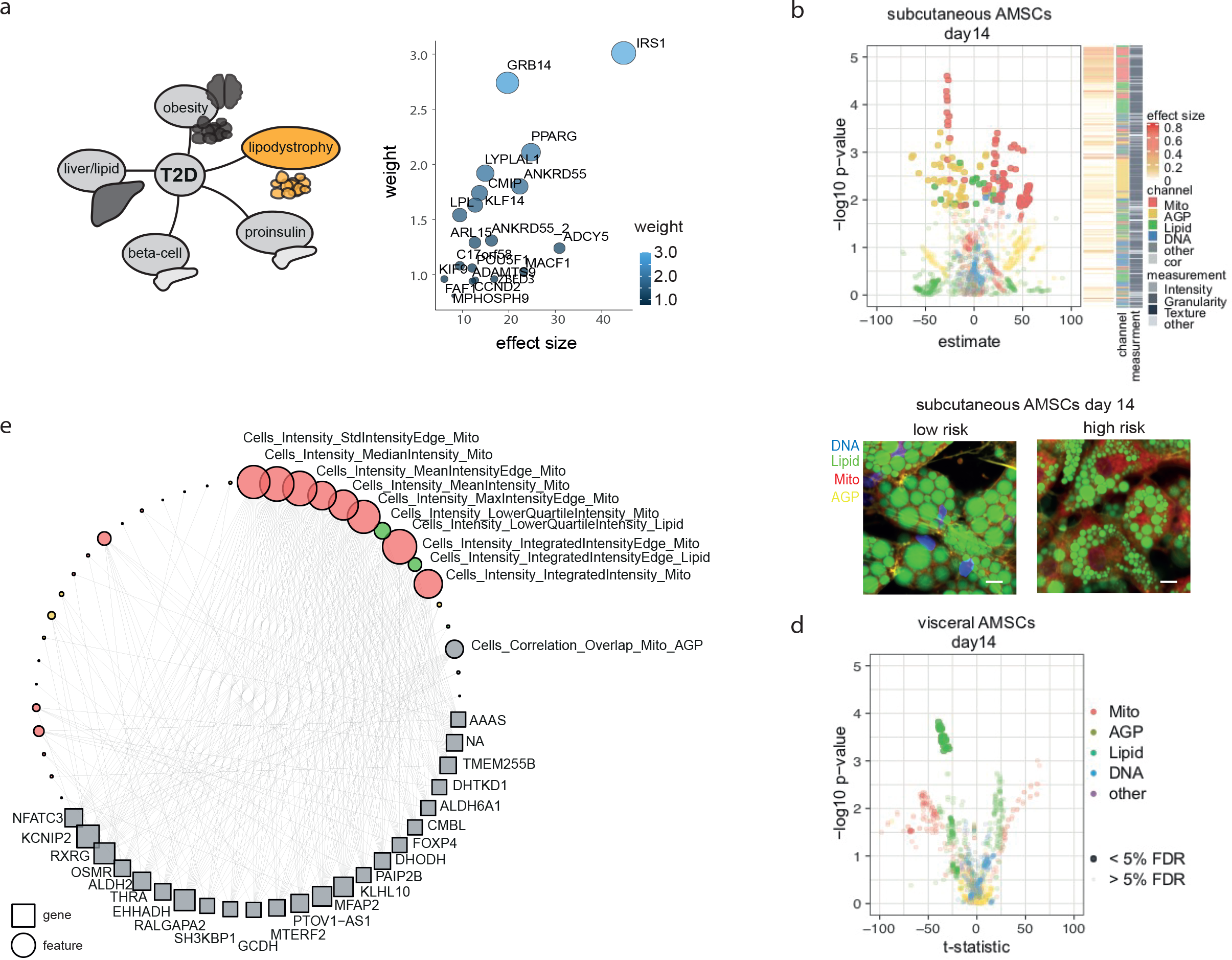
Polygenic risk for lipodystrophy-like phenotype manifests in cellular programs that indicate increased mitochondrial activity, reduced actin cytoskeleton remodeling and reduced lipid accumulation capacity in subcutaneous adipocytes. (a) Schematic of T2D process-specific PRS. Lipodystrophy-specific PRS consists of 20 T2D associated loci contributing to polygenic risk for a lipodystrophy-like phenotype <u>(Udler et al. 2018).</u> (b-d) Depot-specific effects on LipocyteProfiles in AMSCs at day 14 are under the polygenetic control of the lipodystrophy cluster with a mitochondrial and AGP driven profile in subcutaneous AMSCs (b), whereas in visceral AMSCs mostly *Lipid* features were associated with increased polygenic risk (d). See also Figure S6a (day 0, 3, and 8). Representative images of computed averaged subcutaneous AMSCs from low and high risk allele carriers for lipodystrophy PRS show higher mitochondrial intensity, reduced cortical actin and reduced lipid droplet size in high risk carriers (c). (e) Gene - feature connections for lipodystrophy PRS-mediated differential features are enriched for *Mitochondrial Intensity* features informative for mitochondrial membrane potential in subcutaneous AMSCs at day 14 (FDR<0.1%). See also Figure S6d

To further resolve the cellular pathways of lipodystrophy polygenic risk that could underlie the morphological signature in subcutaneous adipocytes, we created a network of genes linked to features identified to be under the control of lipodystrophy polygenic risk. This analysis identified 23 genes that had 10 or more connections to features derived from the lipodystrophy PRS LipocyteProfile (FDR 0.1%, Figure 6e). Eighteen of those genes were significantly correlated with the lipodystrophy PRS (Figure S6d). For example, we found *EHHADH* (a marker gene of peroxisomal β-oxidation) and *NFATC3* (a gene involved in mitochondrial fragmentation and previously linked to a lipodystrophic phenotype in mice <u>(Hu et al. 2018)</u>) to be positively correlated with an increased polygenic risk (*p*=0.04 and *p*=0.004, respectively; Figure S6d), suggesting that gene networks identified through morphological signatures recapitulate mechanisms of polygenic risk, and that LipocyteProfiler can be used to identify molecular mechanisms of disease risk.

Together, these data map aggregated polygenic risk for a lipodystrophy-like phenotype onto cellular programs characterized by increased mitochondrial activity, and decreased lipid accumulation in subcutaneous adipocytes, which is consistent with the notion that limited peripheral storage capacity of adipose tissue underlies polygenic lipodystrophy <u>(Lotta et al. 2017)</u>.

### Allele-specific effect of the *2p23.3* lipodystrophy-like locus on mitochondrial fragmentation and lipid accumulation in visceral adipocytes

To confirm that LipocyteProfiler can link an individual genetic risk locus to meaningful cellular profiles in adipocytes, we investigated a locus on chromosome 2, spanning the *DNMT3A* gene at location *2p23.3*. The metabolic risk haplotype (minor allele frequency of 0.35 in 1000 Genome Phase 3 combined populations) associates with a higher risk for T2D and WHRadjBMI (Figure 7a). To map the *2p23.3* metabolic risk locus to cellular functions, we compared LipocyteProfiles of subcutaneous and visceral AMSCs of risk and non-risk haplotype carriers at 3 time points during adipocyte differentiation (before (day 0), early (day3), and terminal differentiation (day 14)) (Figure 7b). In visceral AMSCs, we identified 83 and 78 features that are significantly different between haplotypes at day 3 and day 14 of differentiation, respectively (Figure 7c, Table S9). At day 3, 70% of significant different image-based features are mitochondrial, and on day 14, 80% of differential features are lipid-related. These findings suggest that the *2p23.3* locus is associated with a mitochondrial function phenotype during early differentiation, which then progresses to altered lipid droplet formation in mature visceral adipocytes. At day 3 of differentiation representative microscopic images show higher mitochondrial stain intensities in risk allele carriers (Figure 7d). Top-scoring, most differential mitochondrial features (*Cells_MaxIntensity_Mito p*=0.004,*Cells_Texture*_*Entropy_Mito p*=0.003 and *Cytoplasm_Granularity_Mito 7 p*=0.02 and *8 p*=0.02; Figure 7e, Figure S7a) were increased in metabolic risk carriers, suggestive of less tubular mitochondria with increased mitochondrial membrane potential and altered function. At day 14 of differentiation, AMSCs from metabolic risk haplotype carriers showed smaller lipid droplets in representative microscopic images (Figure 7f). More specifically, we observed that risk haplotype carriers have decreased *Lipid Intensity* (*p*=0.008; Figure 7g) in the cell and a smaller area of *large Lipid objects* (*LargeLipidobjects_AreaShape p*=0.01; Figure 7g), suggesting a lipid phenotype characterized by reduced lipid droplet stabilization and/or formation. This profile is associated with increased mature adipocyte diameter estimates as shown in Figure 2 and suggests that risk haplotype carriers have a morphological profile that is consistent with visceral WAT hypertrophy. We further note that our findings in human adipocytes are corroborated by organismal perturbation of the candidate effector transcript *DNMT3A* in mice, where deletion of *Dnmt3a* results in changes of whole body fat mass (Figure S7b) <u>(Dickinson et al. 2016)</u> and protects from high-fat-diet induced insulin resistance, mainly attributed to actions in visceral adipose tissue <u>(You et al. 2017)</u>. Together, the data demonstrate that LipocyteProfiler captures complex cellular phenotypes associated with the genetic risk for metabolic diseases and traits, and allows the effective resolution of spatio-temporal context of action. With LipocyteProfiler, we generated a resource that enables unbiased mechanistic interrogation of the hundreds of metabolic disease loci with unknown functions. We have provided all data and software open-access and open-source for the community.

**Figure 7.**
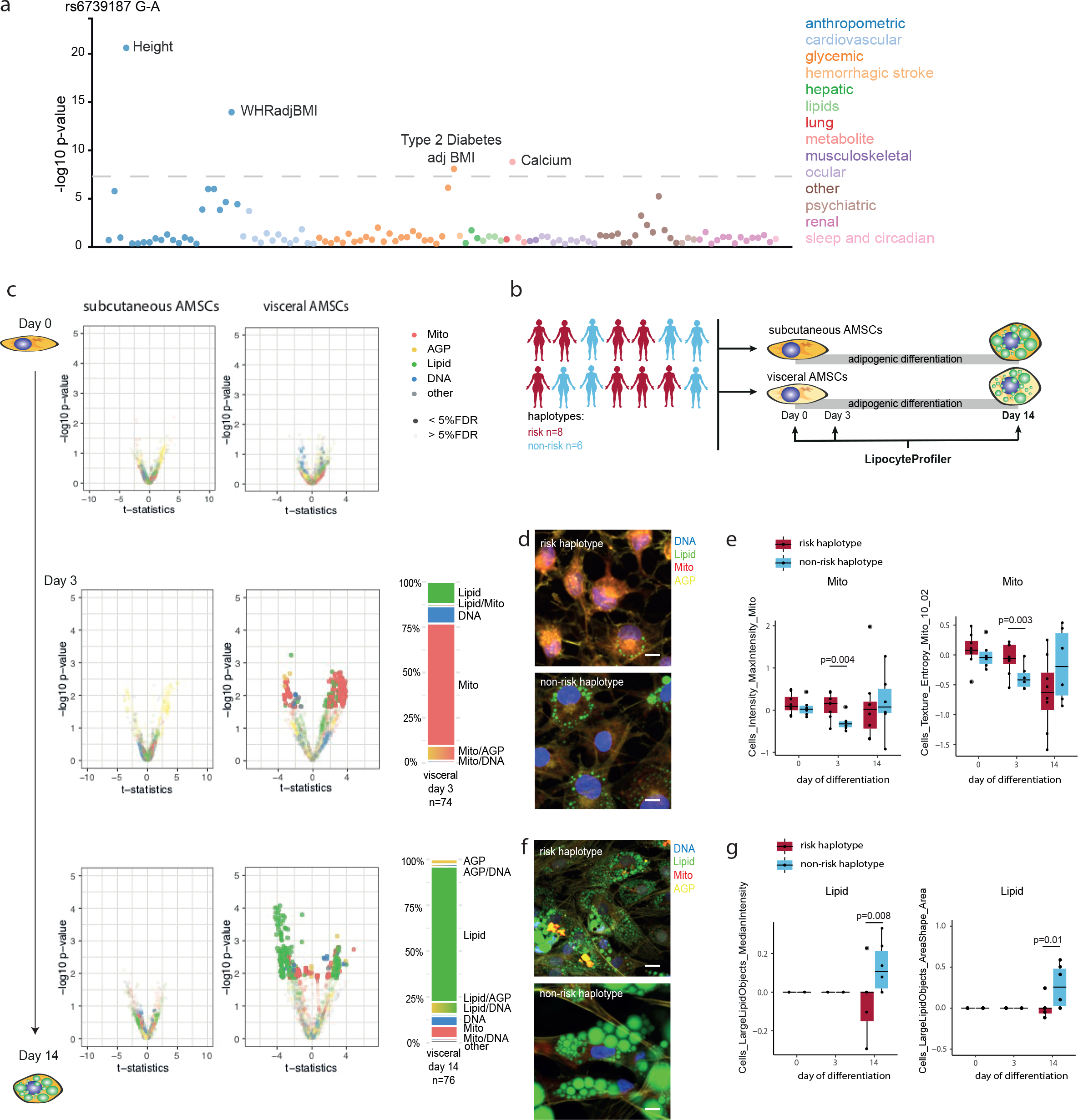
Allele-specific effect of the common *2p23.3* lipodystrophy-like locus on mitochondrial fragmentation and lipid accumulation in visceral adipocytes. (a) PheWAS <u>(Taliun et al. 2020)</u> at the *2q23.3* risk locus shows associations with height, WHRadjBMI, T2D and Calcium. (b) LipocyteProfiler was performed in subcutaneous and visceral AMSCs of eight risk and six non-risk haplotype carriers at three time points during adipocyte differentiation (day 0, 3 and 14). (c) In visceral AMSCs, 74 and 76 features were different between haplotypes at day 3 and day 14 of differentiation, respectively, in visceral AMSCs, with 70% of differential features at day 3 being mitochondrial, and 80% lipid-related at day 14. (d) Representative images of visceral AMSCs from risk (top) and non-risk (bottom) haplotype at day 3 of differentiation stained using LipocytePainting. Scale bar = 10um. (e) *Mito MaxIntensity* and *Mito Texture Entropy* were higher at day 3 of differentiation in visceral AMSCs from 6 risk haplotype carriers, suggesting more fragmented and higher mitochondrial membrane potential. Y-axis shows LP units (normalized LP values across eight batches, see methods). (f) Representative images of visceral AMSCs from risk (top) and non-risk (bottom) haplotype at day 14 of differentiation stained using LipocytePainting. Scale bar = 10um. (g) *LargeLipidObject MedianIntensity* was lower and *Lipid Texture AngularSecondMoment* was higher at day 14 of differentiation in visceral AMSCs from 6 risk haplotype carriers, suggesting a perturbed lipid phenotype characterized by reduced lipid droplet stabilization and/or formation. Y-axis shows LP units (normalized LP values across eight batches, see Methods)

## Discussion***/***Conclusion

In this study, we present a novel high-content image-based profiling framework, LipocyteProfiler, for identification of causal relationships between natural genetic variation, effect of drugs and physiologically relevant stimulations, and effector genes with cellular programs in the context of metabolic disease. We provide proof-of-principle results showcasing that LipoctyeProfiler links genetic variants to distinct morphological and cellular profiles, which demonstrates its usefulness to help unravel disease-relevant complex cellular programs, beyond the information that is typically gained by hypothesis-driven cell-based read-outs. We show that the information gained from LipocyteProfiler can report on both the physiological and pathological states of the cell, and identify morphological and cellular traits underlying cell state transitions, providing a controlled set-up to interrogate dynamic rather than static programs. Using LipocyteProfiler in defined cell states, we are able to robustly detect subtle phenotypic differences driven by drug treatment, genetic perturbation, and natural genetic variation associated with metabolic traits, even with relatively small sample sizes. Our ability to detect these subtle changes might be a consequence of cell morphology capturing the downstream manifestations of genomic, transcriptional, and proteomic effects. We show that polygenic risk for metabolic traits converges into discrete pathways and mechanisms, and demonstrate that LipocyteProfiler elucidates morphological and cellular signatures underlying differential polygenic metabolic risk specific to adipocyte depot, metabolic trait, and cell developmental time point. For example, by selecting AMSCs from donors from the top and the bottom of the PRS distribution for metabolic traits we identify aggregated polygenic effects on lipid degradation in visceral adipocytes for insulin resistance, and on mitochondrial activity and cytoskeleton remodeling in subcutaneous adipocytes for lipodystrophy-specific polygenic contributors to T2D risk. By linking image-based profiles to transcriptional states we further provide a rich resource of gene - cellular trait connections, and provide a foundational image-based assay that enables scalable, unbiased mechanistic interrogation of the hundreds of metabolic disease loci whose functions still remain unknown. We expect that by increasing sample sizes, our approach will help to pave the way to map cellular QTLs in future population-scale image-based profiling endeavours (GWAS in a dish) to link common genetic risk variation to cellular phenotypes in lipocytes, and accelerating therapeutic pathway discoveries.

Limitations:

One limitation in the current study is the relatively low sample size to link genetic variants to cellular and morphological processes. Findings presented here in our proof-of principle study need to be replicated in larger population-scale experiments in the future.

## Acknowledgments

Work in the Claussnitzer lab is supported by the FNIH AMP-T2D RFB8b, NIDDK UM1 DK126185, NIDDK DK102173. M.C. and S.B.R.J. are supported by a Next Generation Award at the Broad Institute of MIT and Harvard. S.L. was supported by a Novo Nordisk Postdoctoral Fellowship run in partnership with the University of Oxford. J.M.Mercader is supported by American Diabetes Association Innovative and Clinical Translational Award 1-19-ICTS-068 H.H. and M.C. received funding within the Clinical Cooperation Group “Nutrigenomics and Type 2 Diabetes” from the Helmholtz Center Munich and the German Center for Diabetes Research. H.H. was funded by the Else Kröner-Fresenius-Foundation, Bad Homburg, Germany. C.M.L is supported by the Li Ka Shing Foundation, NIHR Oxford Biomedical Research Centre, Oxford, NIH (1P50HD104224-01), Gates Foundation (INV-024200), and a Wellcome Trust Investigator Award (221782/Z/20/Z). AL is supported by Grant 2020096 from the Doris Duke Charitable Foundation. The authors gratefully acknowledge the use of the Opera Phenix High-Content/High-Throughput imaging system at the Broad Institute, funded by the S10 Grant NIH OD-026839-01. J.F. is supported by the NIDDK K24 DK110550, UM1 DK126185. M.U. is supported by K23DK114551. S.B.R.J. was supported by NIDDK UM1 DK126185, and NIDDK U01 DK105554. We are grateful for valuable discussions with Xinchen Wang on LipocyteProfiler. The hWAT and hBAT cell lines were kindly shared by Yu-Hua Tseng at the Joslin Diabetes Center Harvard Medical School, and the SGBS line by Martin Wabitsch, University of Ulm, Germany.

## Author Contributions

Conceptualization, S.L., S.S., J-M.M., A.A., J.F., S.J., M.C.;

Methodology, S.L., S.S., M.A., D.S., G.W., B.C., A.C., M.C.,

Formal Analysis, S.L., S.S., J-M.M., K.F., G.D., D.S., G.W., B.C.,M.S.U;

Investigation, S.L., S.S., A.A., J.H., G.G., E.MG., A.S., A.H., A.S., S.H., V.C-N;

Resources, H.H, M.C., J. F..

Data Curation, H.D., S.R-P.,

Visualization, S.L., S.S.;

Supervision, B.C., H.H., M.U., J.F., A.C., C.M.L., S.J., M.C.;

Funding Acquisition, H.H., J.F., A.C., C.M.L., S.J., M.C.;

Writing – Original Draft, S.L., S.S., M.C.;

Writing – Review & Editing, J-M.M., H.D., A.A., A.C., S.J., J.F., M.S.U, C.M.L., H.H.

## Supplemental Information

### Supplementary Figure legends

**Figure S1.**
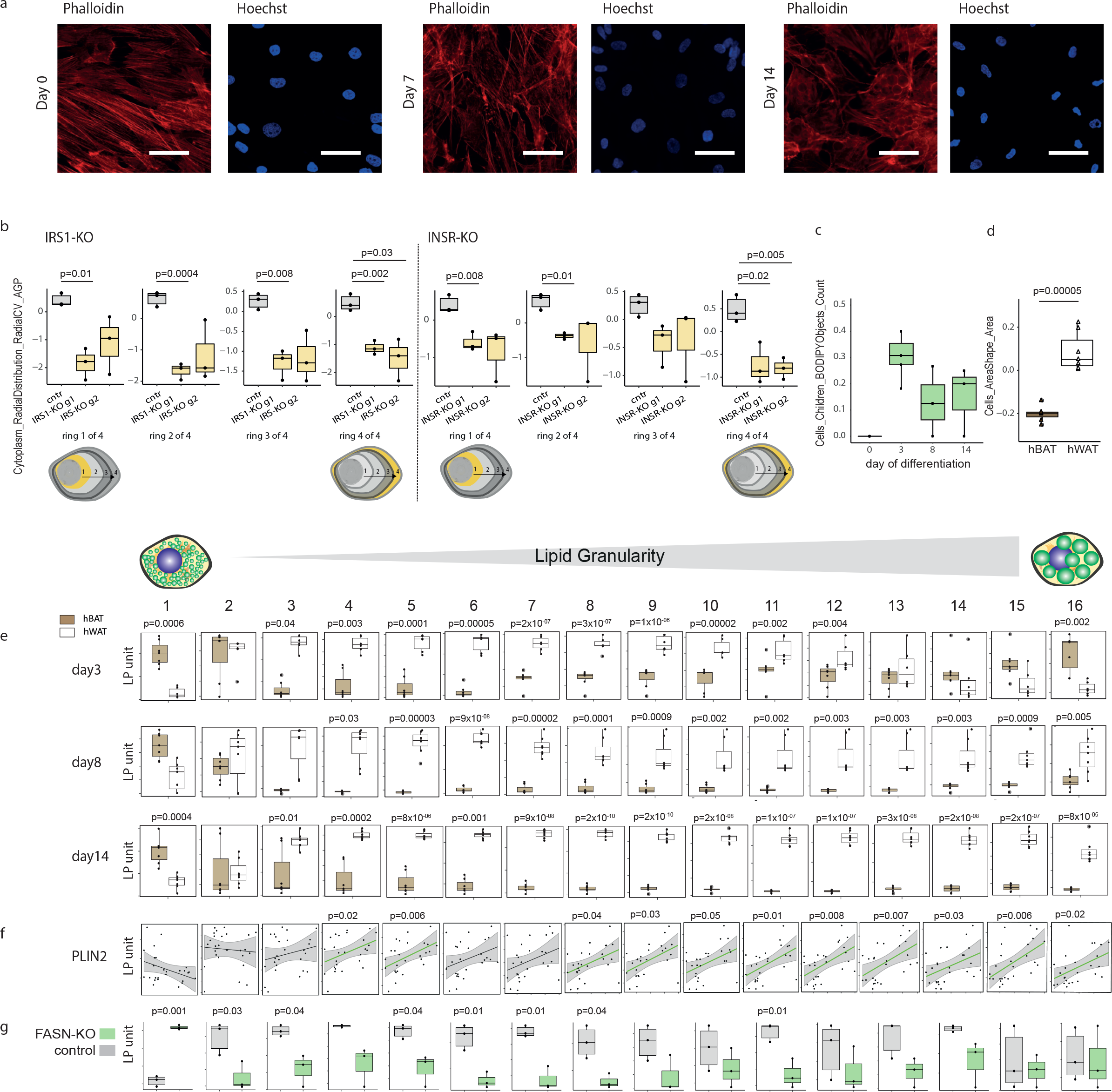
LipocyteProfiler characterise lipid droplet formation and expansion. (a) Representative microscope images of actin-cytoskeleton remodeling during adipocyte differentiation. (left to right day 0, 7, 14; red = Phalloidin, actin-cytoskeleton, blue = Hoechst, DNA). scale bar = 52.8µm (b) *Cytoplasm_RadialDistribution_RadialCV_AGP* measures (ring 1 to 4) are reduced in CRISPR/Cas9-mediated KO of *INSR* and *IRS-1.* Data is shown for two guides targeting *INSR or IRS-1*. (g1 and g2). (c) Lipid droplets form in early differentiation and saturate thereafter. Y-axis shows LP units (normalized LP values across eight batches, see Methods). (d) White (hWAT) are larger than brown (hBAT) adipocytes after 14 days of adipogenic differentiation as cells become lipid-laden, measured by *Cells_AreaShape_Area*. Y-axis shows LP units (normalized LP values across three batches, see Methods). (e) *Lipid Granularity* measurements, captured by spectra of 16 lipid droplet size measures, show size-specific changes in hWAT and hBAT during differentiation, suggesting hBAT generally accumulate less medium-size and large lipid droplets as seen by lower values across the spectra of granularity. Y-axis shows LP units (normalized LP values across three batches, see Methods). (f) Granularity features informative for larger lipid droplets (*Lipid Granularity 10*-*16*) correlate positively with *PLIN2* gene expression. Y-axis shows LP units (normalized LP values across eight batches, see Methods). X-axis *PLIN2* expression (rpm). (g) *Lipid Granularity* measures are reduced in CRISPR/Cas9-mediated KO of *FASN* in hWAT at day 14 of differentiation. Y-axis shows LP units (normalized LP values across CRISPR-KO data, see Methods). Data is shown for one guide targeting *FASN*.

**Figure S2.**
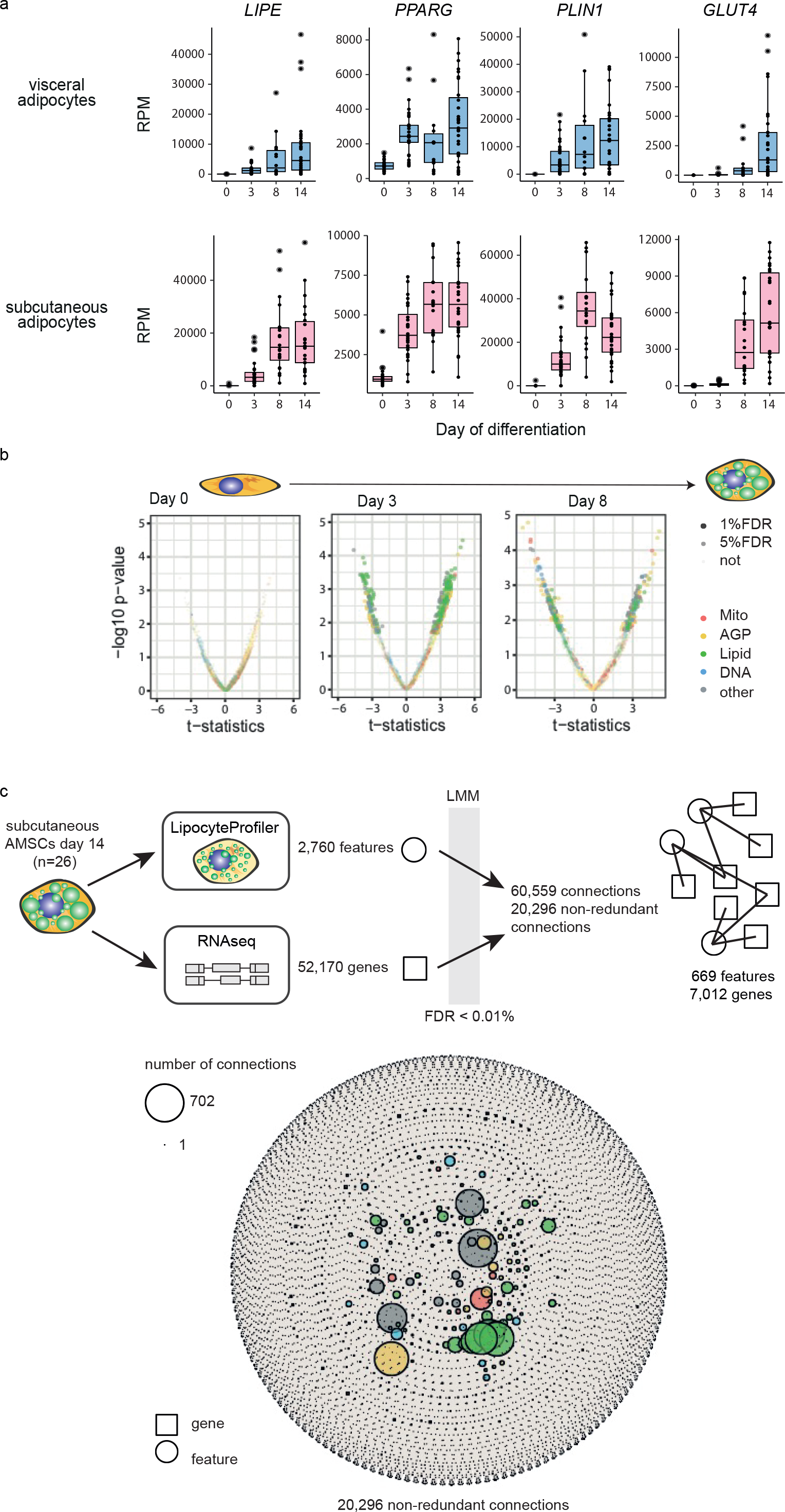
LipocyteProfiler identifies depot-specific cellular signatures in adipocytes. (a) Gene expression from RNAseq of adipogenesis marker genes *LIPE, PPARG, PLIN1* and *GLUT4* in visceral (top) and subcutaneous (bottom) AMSCs throughout differentiation. (b) Subcutaneous and visceral AMSCs have distinct morphological and cellular profiles with differences that are spread across all channels that become apparent at day 3 of differentiation and are maintained at day 8. See also Figure 2c (day 14). (c) A Linear mixed model (LMM) was applied to correlate 2,760 morphological features derived from LipocyteProfiler with 52,170 transcripts derived from RNAseq in matched samples of subcutaneous AMSCs at terminal differentiation (day 14). With FDR < 0.01%, we discover 20,296 non-redundant connections that map to 669 morphological features and 7,012 genes. See also Figure 3a, b (FDR< 0.1%))

**Figure S3:**
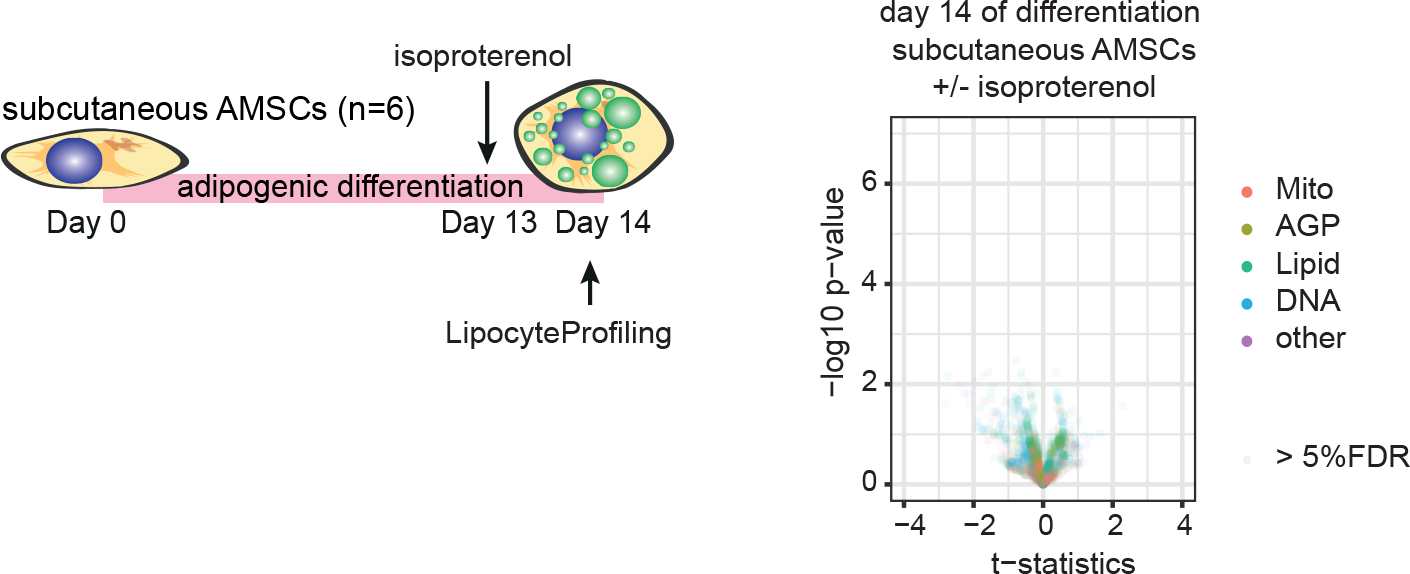
LipocyteProfiler reveals depot-specific isoproterenol-induced cellular signatures in subcutaneous adipocytes compared to visceral adipocytes. Isoproterenol treatment results in no effect on morphological profile in subcutaneous AMSCs at day 14 of differentiation. See also Figure 4b (visceral).

**Figure S4.**
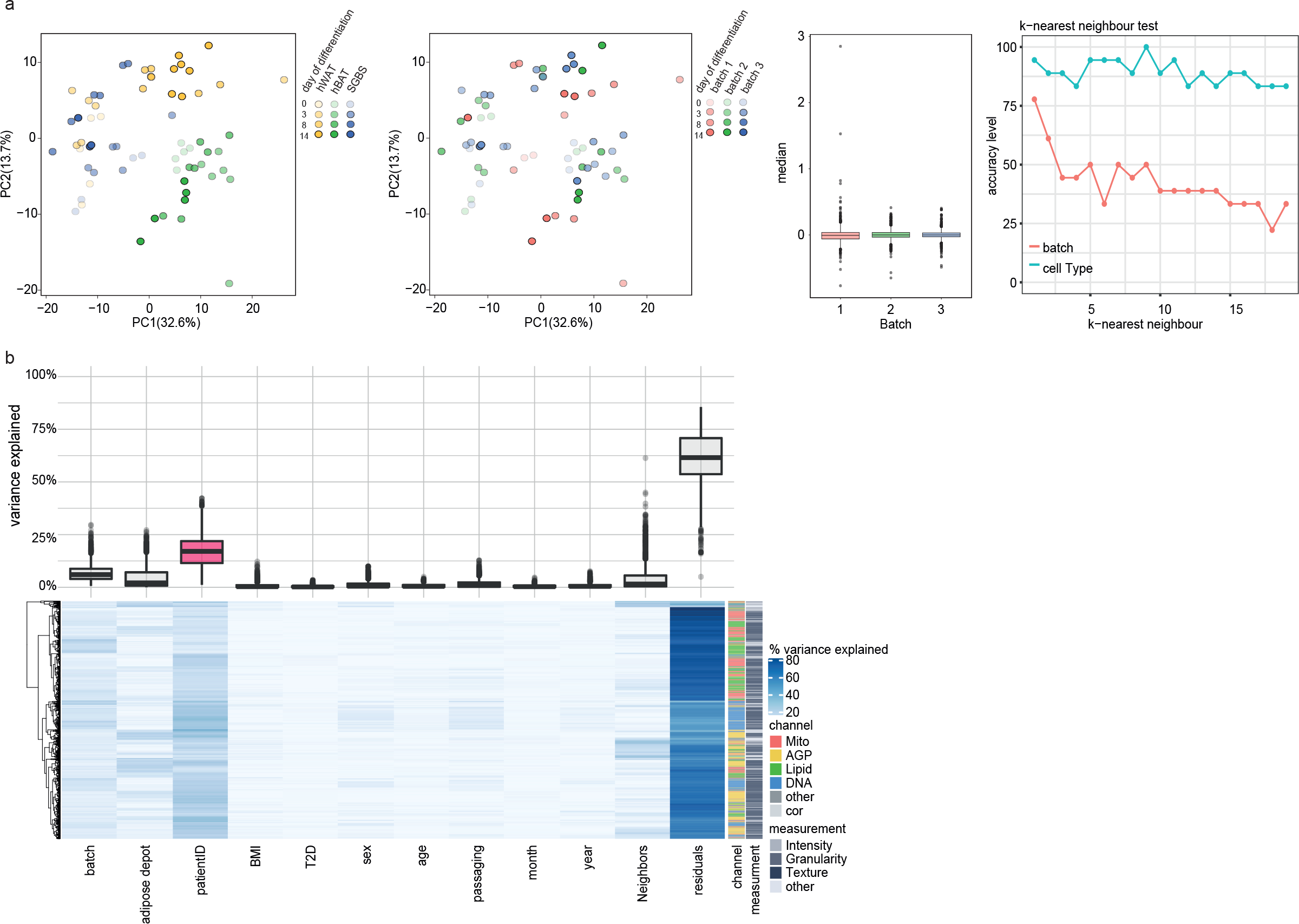
LipocyteProfiler detects and distinguishes inter-individual genetic variation from extrinsic confounding factors. (a) Morphological profiles of hBAT, hWAT and SGBS across differentiation cluster according to cell type and show maturation trajectory in PC1 and PC2, though do not cluster by batch (two plots on left). BEclear analysis shows no significant batch effect and accuracy of predicting cell type is higher than predicting batch using a k-nearest neighbor supervised machine learning algorithm (two plots on right). (b) Variance component analysis across all data to assess contribution of intrinsic genetic variation on adipocyte morphology and cellular traits across 65 donor-derived differentiating AMSCs. This analysis showed that patientID explains the majority of feature variance compared to contribution of other possible confounding factors such as batch, adipose depot, T2D status, age, sex, BMI, cell density, month/year of sampling and passage numbers.

**Figure S5.**
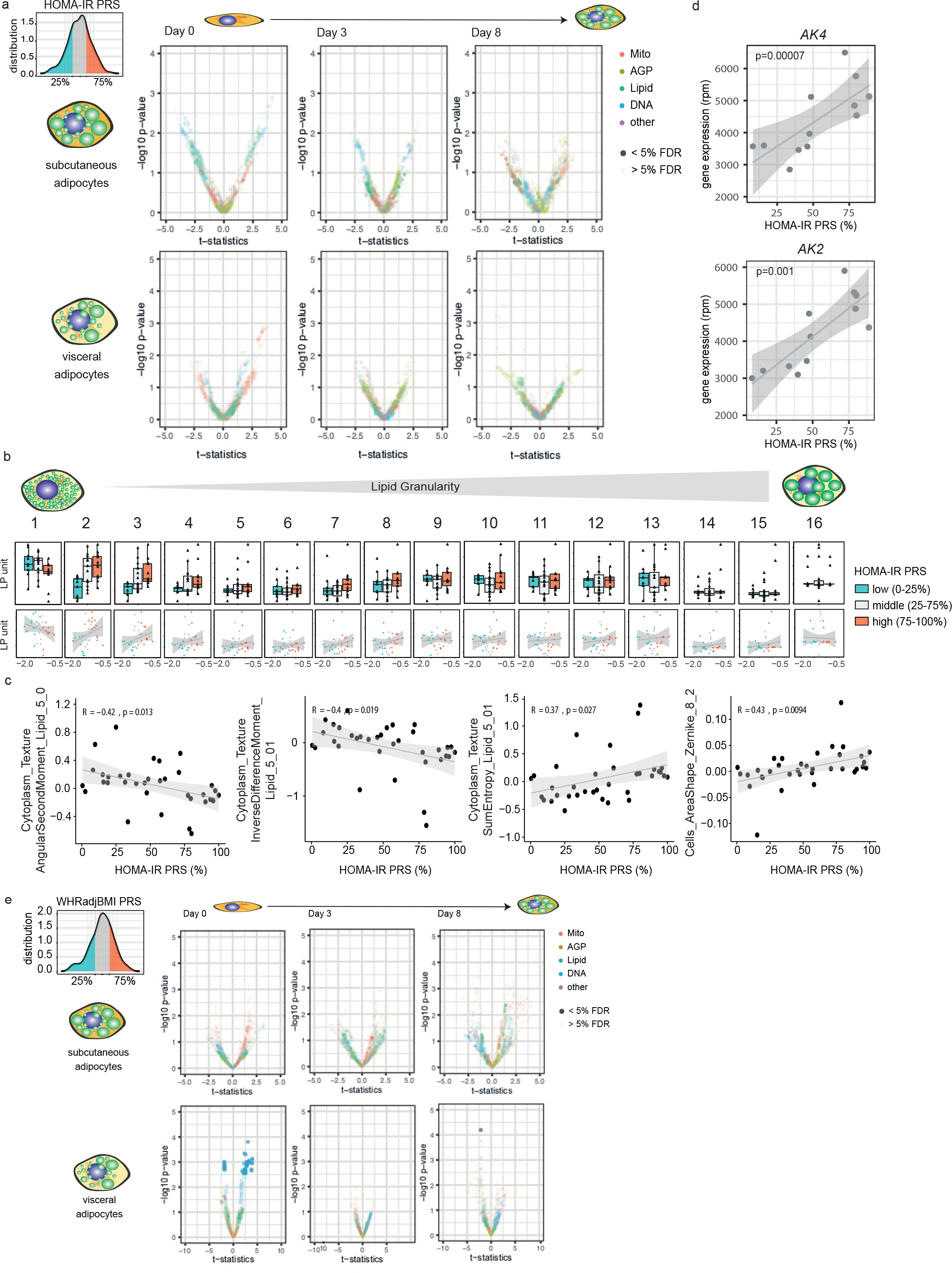
LipocyteProfiler discovers significant polygenic effects on image-based cellular signatures for HOMA-IR and WHRadjBMI. (a) LipocyteProfiler differences between top and bottom 25% of HOMA-IR risk in subcutaneous and visceral AMSCs at day 0, 3, and 8 of adipogenesis. (See also Figure 5b; day 14 visceral) (b) Mature visceral AMSCs (day 14) derived from individuals with high polygenic risk for insulin resistance show an increased number of small- to medium-sized lipid droplets compared to AMSCs from donors with low polygenic risk. *Cytoplasm_Granularity_BODIPY* measures 1-16. Y-axis shows autoscaled LP units (normalized LP values across eight batches, see methods). Bottom panel; x-axis shows PRS for HOMA-IR. (c) Representative significant features of HOMA-IR morphological profile correlate with PRS percentile (See also Figure 5c) Y-axis shows LP units (normalized LP values across eight batches, see methods). (d) *AK4* and *AK2* correlate significantly with HOMA-IR PRS in visceral AMSCs at day 14 of differentiation. (e) LipocyteProfiler differences between top and bottom 25% of WHRadjBMI risk in subcutaneous and visceral AMSCs at day 0, 3, and 8 of adipogenesis. See also Figure 5b; day 14 subcutaneous).

**Figure S6.**
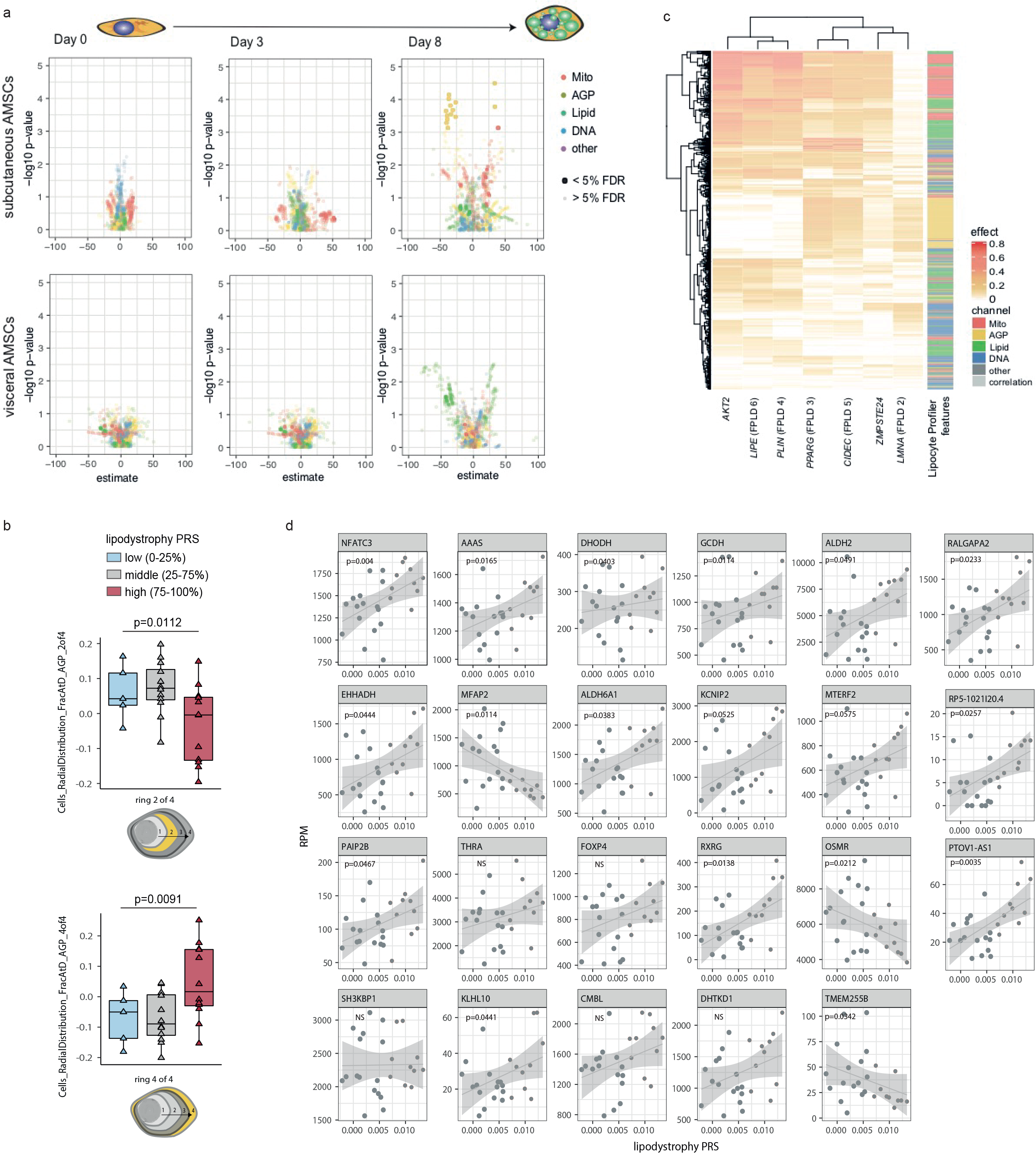
Morphological profile of polygenic risk for lipodystrophy shows similarity to cellular signatures of marker genes of monogenic familial partial lipodystrophy syndromes. (a) LipocyteProfiler differences driven by polygenic risk in subcutaneous and visceral AMSCs at day 0, 3, and 8 of adipogenesis. See also Figure 6b, d; day14). (b) Polygenic risk for lipodystrophy affects the structure of the actin cytoskeleton; golgi; plasma membrane of differentiated subcutaneous AMSCs (*Cells_RadialDistribution_FraAtD_2of4* and *4of4*). Y-axis shows LP units (normalized LP values across eight batches, see Methods). (c) LipocyteProfiler features correlated with genes underlying monogenic familial partial lipodystrophy syndromes including *PPARG*, *LIPE*, *PLIN1*, *AKT2*, *CIDEC*, *LMNA* and *ZMPSTE24* match LipocyteProfiler morphological and cellular signatures characterizing the lipodystrophy-specific PRS effects, and highlight *Mito* and *AGP* features (See also Figure 6b; day14). (d) Genes significantly correlating with the lipodystrophy PRS-mediated differential features (LMM, FDR<0.1%) are under polygenic control for lipodystrophy risk.

**Figure S7.**
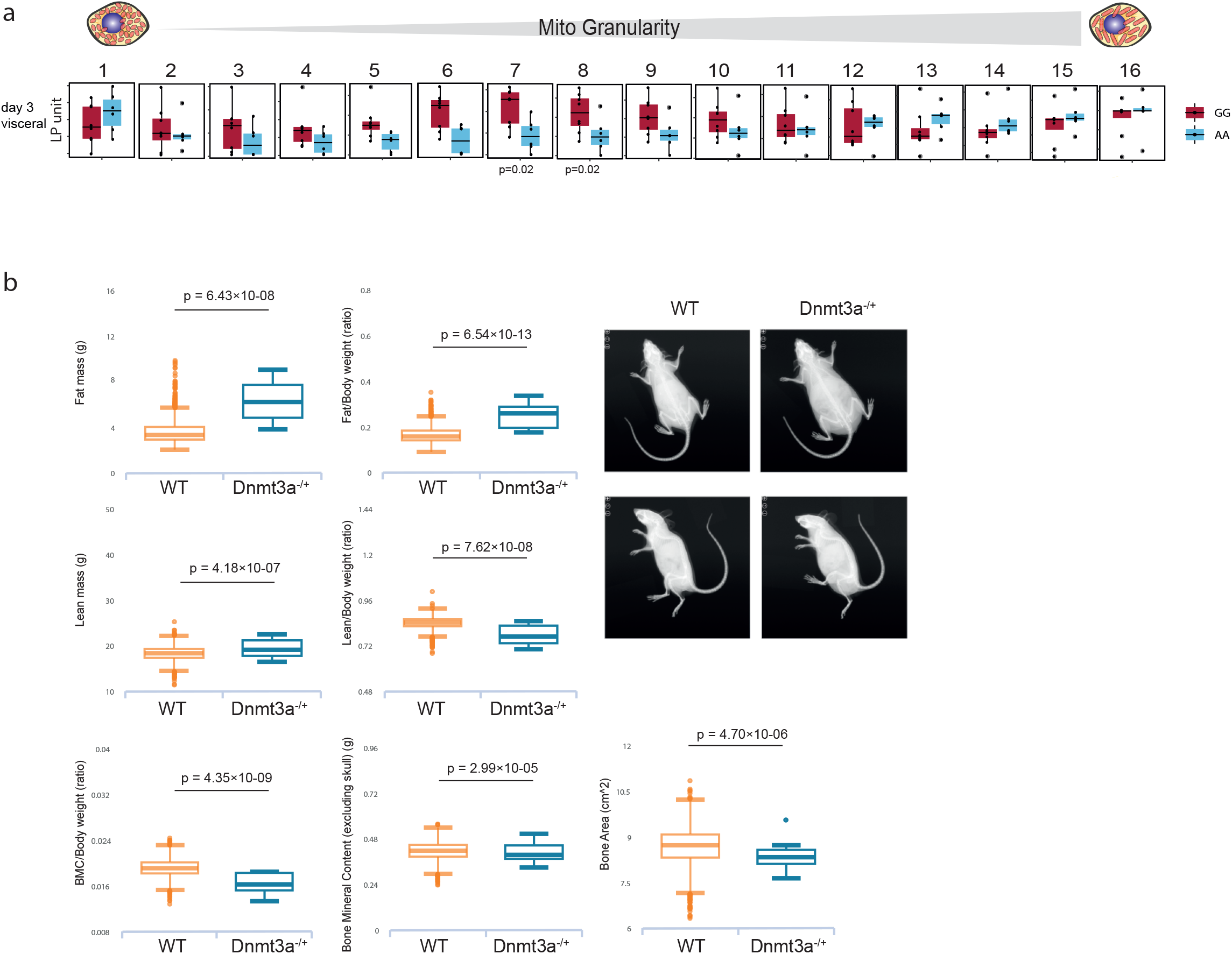
Heterozygous knockout mice for DNMT3A have adipose and bone phenotypes. (a) *Mito Granularity* (7-8 size measures) at day 3 of differentiation in visceral AMSCs was increased in risk allele carriers, suggestive of less tubular mitochondria. Y-axis shows autoscaled LP units (normalized LP values across eight batches, see Methods) (b) Heterozygous knockout mice for DNMT3A show increased body weight due to increased overall fat mass and have reduced bone mineral density. Data retrieved from www.mousephenotype.org <u>(Dickinson et al. 2016)</u>.

## Methods

### *Human primary AMSCs isolation/*Abdominal laparoscopy cohort - Munich Obesity BioBank / MOBB

We obtained AMSCs from subcutaneous and visceral adipose tissue from patients undergoing a range of abdominal laparoscopic surgeries (sleeve gastrectomy, fundoplication or appendectomy). The visceral adipose tissue is derived from the proximity of the angle of His and subcutaneous adipose tissue obtained from beneath the skin at the site of surgical incision. Additionally, human liposuction material was obtained. Each participant gave written informed consent before inclusion and the study protocol was approved by the ethics committee of the Technical University of Munich (Study № 5716/13). Isolation of AMSCs was performed as previously described <u>(Skurk and Hauner 2012).</u> For a subset of donors, purity of AMCSs was assessed as previously described <u>(Raajendiran et al. 2019).</u> Briefly, cells were stained with 0.05ug CD34, 0.125ug CD29, 0.375ug CD31, 0.125ug CD45 per 250K cells and analyzed on CytoFlex together with negative control samples of corresponding AMSCs.

### Differentiation of human AMSCs

For imaging, cells were seeded at 10K cells/well in 96-well plates (Cell Carrier, Perkin Elmer #6005550) and induced 4 days after seeding. For RNAseq, cells were seeded at 40K cells/well in 12-well dishes (Corning). Before Induction cells were cultured in proliferation medium (Basic medium consisting of DMEM-F12 1% Penicillin - Streptomycin, 33µM Biotin and 17µM Pantothenate supplemented with 0.13µM Insulin, 0.01ug/ml EGF, 0.001ug/ml FGF, 2.5%FCS). Adipogenic differentiation was induced by changing culture medium to induction medium. (Basic medium supplemented with 0.861μM Insulin, 1nM T3, 0.1μM Cortisol, 0.01mg/ml Transferrin, 1μM Rosiglitazone, 25nM Dexamethasone, 2.5nM IBMX). On day 3 of adipogenic differentiation culture medium was changed to differentiation medium (Basic medium supplemented with 0.861μM Insulin, 1nM T3, 0.1μM Cortisol, 0.01mg/ml Transferrin). Medium was changed every 3 days. Visceral-derived AMSCs were differentiated by further adding 2% FBS as well as 0.1mM oleic and linoleic acid to the induction and differentiation media. For isoproterenol stimulation experiments, 1uM isoproterenol was added to the differentiation media and cells treated overnight.

### Isolation and adipocyte diameter determination of floating mature adipocytes

Mature adipocyte isolation was carried out as described earlier (https://pubmed.ncbi.nlm.nih.gov/26081284/). Immediately after isolation, approximately 50 μl of the adipocyte suspension was pipetted onto a glass slide and the diameter of 100 cells was manually determined under a light microscope.

### Primary human hepatocytes culture

Primary human hepatocytes (PHH) were purchased from BioIVT. Donor lot YNZ was used in this study. PHH were thawed and immediately resuspended in CP media (BioIVT) supplemented with torpedo antibiotic (BioIVT). Cell count and viability were assessed by trypan blue exclusion test prior to plating. Hepatocytes were plated onto collagen-coated Cellcarrier-96 Ultra Microplates (Perkin Elmer) at a density of 50,000 cells per well in CP media supplemented. Four hours after plating, media was replaced with fresh CP media. After 24 h, media was replaced with fresh CP media or CP media containing oleic acid (0.3mM) or metformin (5mM). Hepatocytes were incubated for an additional 24 h prior to processing.

### LipocytePainting

Human primary AMSCs and PHH were plated in 96-well CellCarrier plates (Perkinelmer #6005550). AMSCs were differentiated for 14 days and high content imaging was performed at day 0, day 3, day 8 and day 14 of adipogenic differentiation. Primary human hepatocytes were stained after 48 h in culture, and 24h following treatment with oleic acid or metformin. On the respective day of the assay, cell culture media was removed and replaced by 0.5uM Mitotracker staining solution (1mM MitoTracker Deep Red stock (Invitrogen #M22426) diluted in culture media) to each well followed by 30 minutes incubation at 37°C protected from light. After 30min Mitotracker staining solution was removed and cells were washed twice with Dulbecco’s Phosphate-Buffered Saline (1X), DPBS (Corning® #21-030-CV) and 2.9uM BODIPY staining solution (3.8mM BODIPY 505/515 stock (Thermofisher #D3921) diluted in DPBS) was added followed by 15 minutes incubation at 37°C protected from light. Subsequently, cells were fixed by adding 16% Methanol-free Paraformaldehyde, PFA (Electron Microscopy Sciences #15710-S) directly to the BODIPY staining solution to a final concentration of 3.2% and incubated for 20 minutes at RT protected from light. PFA was removed and cells were washed once with Hank’s Balanced Salt Solution (1x), HBSS (Gibco #14025076). To permeabilize cells 0.1% Triton X-100 (Sigma Aldrich #X100) was added and incubated at RT for 10 minutes protected from light. After Permeabilization multi-stain solution (10 units of Alexa Fluor™ 568 Phalloidin (ThermoFisher #A12380), 0.01mg/ml Hoechst 33342 (Invitrogen #H3570), 0.0015mg/ml Wheat Germ Agglutinin, Alexa Fluor™ 555 Conjugate (ThermoFisher #W32464), 3uM SYTO™ 14 Green Fluorescent Nucleic Acid Stain (Invitrogen #S7576) diluted in HBSS) was added and cells were incubated at RT for 10 minutes protected from light. Finally, staining solution was removed and cells were washed three times with HBSS. Cells were imaged using a Opera Phenix High content screening system using confocal, 20x objective. Per well we imaged 25 fields.

### LipocyteProfiling

Quantitation was performed using CellProfiler 3.1.9. Prior to processing, flat field illumination correction was performed using functions generated from the median intensity across each plate. Nuclei were identified using the DAPI stain and then expanded to identify whole cells using the Phalloidin/WGA and BODIPY stains. Regions of cytoplasm were then determined by removing the Nuclei from the Cell segmentations. Speckles of BODIPY staining were enhanced to assist in detection of small and large individual Lipid objects. For each object set measurements were collected representing size, shape, intensity, granularity, texture, colocalization and distance to neighbouring objects. After LipocyteProfiler (LP) feature extraction data was filtered by applying automated and manual quality control steps. First, fields with a total cell count less than 50 cells were removed. Second, every field was assessed visually and fields that were corrupted by experimental induced technical artifacts were removed. Furthermore blocklisted features (Way, Gregory (2019): Blocklist Features - Cell Profiler. figshare. Dataset. https://doi.org/10.6084/m9.figshare.10255811.v3), LP-features measurement category *Manders, RWC and Costes*, that are known to be noisy and generally unreliable were removed. Additionally, LP-features named *SmallLipidObjetcs,* that measure small objects stained by SYTO14 rather than lipid informative objects, were also removed. After filtering data were normalized per plate using a robust scaling approach <u>(Pedregosa et al. 2011)</u> that subtracts the median from each variable and divides it by the interquartile range. Individual wells were aggregated for downstream analysis by cell depot and day of differentiation.

Subsequent data analyses were performed in R3.6.1 and Matlab using base packages unless noted. To assess batch effects we visualised the data using a Principle component analysis and quantified it using a Kolmogorov-Smirnov test implemented in the “BEclear” R package <u>(Akulenko et al. 2016).</u> Additionally we performed a k-nearest neighbour (knn) supervised machine learning algorithm implemented in the “class” R package <u>(Venables and Ripley 2002)</u> to investigate the accuracy of predicting biological and technical variation. For this analysis the data set, consisting of 3 different cell types (hWAT, hBAT, SGBS) distributed on the 96-well plate, imaged at 4 days of differentiation, was split into equally balanced testing (n=18) and training (n=56) sets. Accuracy of the classification model was predicted based on three different categories cell type, batch and column of the 96-well plate. (github)

For dimensionality reduction visualisation Uniform manifold approximation and projection maps (UMAP) were created using the UMAP R package version 0.2.7.0 <u>(McInnes et al. 2018)</u> (github). To visualise LipocyteProfiles and their effect size ComplexHeatmap Bioconductor package version 2.7.7 <u>(Gu et al. 2016)</u> was used (github)

To identify patterns of adipocyte differentiation underlying the morphological profiles a sample progression discovery analysis (SPD) was performed using the algorithm previously described <u>(Qiu et al. 2011)</u>. Briefly, the two adipose depots were analysed separately and features were clustered into modules based on correlation (correlation coefficient 0.6). Minimal spanning trees (MST) were constructed for each module and MSTs of each module are correlated to each other. Modules that support common MST were selected and an overall MST based on features of all selected modules were reconstructed.

Variance component analysis was performed by fitting multivariable linear regression models - yi ∼ xi + zi + .. - where y denotes an LipocyteProfiler feature of individual i and x, z, etc. independent variables that could confound identification of biological sources of variability of the dataset. Independent variables are experimental batch, adipose depot, passaging before freezing, season and year of of AMSCs isolation, sex, age, BMI, T2D status of individual, LipocyteProfiler feature *Cells_Neighbors_PercentTouching_Adjacent* corresponding to density of cell seeding and identification numbers of induviduals. (github)

To test whether there is a difference of morphological profiles on the tail ends of polygenic risk scores (PRS) for T2D, HOMA-IR and WHRadjBMI a multi-way analysis of variance (ANOVA) was performed. Individuals belonging to top 25% and bottom 25% of PRS score distribution are categorized into a categorical variable with 2 levels, top 25% or 25% bottom, according to their PRS percentile. Differences of morphological profiles are predicted using the categorised PRS variable adjusted for sex, age, BMI and batch. For the process-specific lipodystrophy polygenic risk score a linear regression model was fitted adjusted for sex, age, BMI and batch to predict differences of morphological and cellular profiles. To overcome multiple testing burden p-values were corrected using false positive rate (FDR) described in R package “qvalue” <u>(qvalue)</u>. Features with FDR < 5% were classified to be significantly impacted by the PRS variable. (github)

### Staining and microscopy of actin-cytoskeleton in subcutaneous AMSCs

To stain the actin cytoskeleton, and nuclei, cells were washed twice with ice cold PBS and fixed with paraformaldehyde Roti-Histofix 4 % (Roth, Karlsruhe, Germany) for 15 min. Cells were washed twice with ice cold PBS for 5 min and incubated with ice cold 0.1 % Triton X/PBS (Roth, Karlsruhe, Germany) for 5 min. Cells were washed twice with PBS and stained with 0.46 % Bisbenzimide H 33258 (Sigma-Aldrich, Steinheim, Germany), and 1% Phalloidin-Atto-565 (Sigma-Aldrich, Steinheim, Germany). Cells were incubated for one hour at RT in the dark. Afterwards, cells were washed twice with PBS for 5 min and kept in PBS at 4 °C until imaging. Images were acquired on a Leica DMi8 microscope using the HC PL APO ×63/1.40 oil objective. Images were processed using the Leica LasX software.

### Generating of average cells

For each group of interest, cells were pooled and divided into 100 clusters via K-Means clustering (scikit-learn). Individual cells were then sampled from the cluster closest to a theoretical point representing the mean of all object measurements, as determined by a euclidean distance matrix.

### RNA-seq

RNA-seq data were processed using FastQC <u>(Krueger and Others 2015)</u> and spliced reads were aligned to human genome assembly (hg19) using STAR <u>(Dobin et al. 2013)</u> followed by counting gene levels using Rsubread R package <u>(Liao et al. 2019)</u>. Next, raw read counts were normalized using DESseq2 R package <u>(Love et al. 2014)</u>. For differential expression analysis on the tail ends of polygenic risk scores (PRS) for HOMA-IR a multi-way analysis of variance (ANOVA) was performed on subset of 512 genes (GSEA hallmark gene sets for adipogenesis, fatty acid metabolism and glycolysis). Individuals belonging to top 25% and bottom 25% of PRS score distribution are categorized into a categorical variable with 2 levels, top 25% or 25% bottom, according to their PRS percentile. Differences in transcriptional profiles are predicted using categorised PRS variable adjusted for for sex, age, BMI and batch. To overcome multiple testing burden p-values were corrected using false positive rate (FDR) described in R package “qvalue” <u>(qvalue)</u>. Genes with FDR < 10% were classified to be significantly impacted by PRS and were uploaded to Enrichr to analyse them as a gene list against the WikiPathways. (github)

### Gene expression and LipocyteProfiler feature network

A linear regression model was fitted of 2,760 LP-features and global transcriptome RNA-seq data adjusted for sex, age, BMI and batch in subcutaneous AMSCs at day 14 of differentiation. Gene LP features association were declared to be significant when passing FDR cut-off of 0.1% FDR. LP features belonging to *Cells* category were used for further analysis. Associations between genes and LP features were visualised using “igraph” R package <u>(Csardi et al. 2006)</u> (github). Genes that are connected to top scoring LP features were uploaded to Enrichr to analyse them as a gene list against WikiPathways or BioPlanet. Adipocyte marker genes, *SCD, PLIN2, LIPE, INSR, GLUT4 and TIMM22,* were chosen to demonstrate morphological profiles matching their known pathways, by identifying LP features that associate with those genes with a global significant level of 5% FDR. (github)

### CRISPR-Cas9 mediated knockout of adipocyte marker genes

We generated a hWAT cell-line stably expressing Cas9 as previously described <u>(Shalem et al. 2014).</u> We validated the generated line by assessing Cas9 activity (90%) and adipocyte differentiation capacity using adipocyte marker gene expression and morphological profiling. CRISPR/Cas9 mediated knockdown of *PPARG, PPARGC1AA, MFN1*, *PLIN1, INSR,* and *IRS1* was performed in pre-adipocytes (5 days before differentiation) using three replicates per guide and two guides per gene (guide sequences targeting *PPARG*: ATACACAGGTGCAATCAAAG and CAACTTTGGGATCAGCTCCG; *PPARGC1A* TATTGAACGCACCTTAAGTG and AGTCCTCACTGGTGGACACG; *MFN1*: CACCAGGTCATCTCTCAAGA and TTATATGGCCAATCCCACTA; *PLIN1*: TCACGGCAGATACTTACCAG and TCTGCACGGTGTATCGAGAG; *INSR*: TTATCGGCGATATGGTGATG and AGTGAGTATGAGGATTCGGC; *IRS1* CCCAGGACCCGCATTCAAAG and CCGAAGCACTAGATCGCCGT) as well as five non-targeted controls (control guide sequences: ATCAGGCCTTGTCCGTGATT, TACGTCATTAAGAGTTCAAC, GACAGTGAAATTAGCTCCCA, GATTCATACTAAACACTCTA, CCTAGTTCATAAGCTACGCC) in an 96-well arrayed format. Guide on-target efficiency was assessed using Next-generation sequencing followed by CRISPResso analysis <u>(Pinello et al. 2016).</u> AMSCs were stained using LipocytePainting (see above) on day 14 of differentiation. After feature extraction and QC steps (see also LipocyteProfiling), we removed samples where guide cutting efficiency was <10% or were discrepancy between the two guides was equal or above 10%. For visualisations we used one non-targeted control, that showed lowest standard deviation of replicates and was closest to the median of all five non-targeted controls across all LipocyteProfiler features.

## Quality Control

Genotyping of all samples was performed in two separate batches using the Infinium HTS assay on Global Screening Array bead-chips. Since the two sets of samples were genotyped with different versions of the beadchips and in different batches, we Qced, imputed, and generated the genome-wide polygenic scores separately and combined the results afterwards.

A 3-step quality control protocol was applied using PLINK <u>(Purcell et al. 2007; Chang et al. 2015)</u>, and included 2 stages of SNP removal and an intermediate stage of sample exclusion.

The exclusion criteria for genetic markers consisted of: proportion of missingness ≥ 0.05, HWE *p* ≤ 1 x 10^-20^ for all the cohort, and MAF < 0.001. This protocol for genetic markers was performed twice, before and after sample exclusion.

For the individuals, we considered the following exclusion criteria: gender discordance, subject relatedness (pairs with PI-HAT ≥ 0.125 from which we removed the individual with the highest proportion of missingness), sample call rates ≥ 0.02 and population structure showing more than 4 standard deviations within the distribution of the study population according to the first seven principal components (Supplementary Figure 27). After QC, 35 subjects remained for the analysis for which we had matched LipocyteProfiler imaging data.

Genotypes were phased with SHAPEIT2 <u>(Delaneau et al. 2013)</u>, and then performed genotype imputation with the Michigan Imputation server, using Haplotype Reference Consortium (HRC) <u>(Consortium and the Haplotype Referenc…)</u> as reference panel. We excluded variants with an info imputation r-squared < 0.5 and a MAF < 0.005.

Genome-wide polygenic scores were computed using PRS-CS <u>(Ge et al. 2019)</u> and using the “auto” parameter to specify the phi shrinkage parameter. We computed the PRS-CS polygenic scores for the following traits: T2D <u>(Mahajan et al. 2018)</u>, BMI, waist-to-hip ratio adjusted and unadjusted by BMI, and stratified by sex and combined <u>(Pulit et al. 2019)</u>. Genome-wide PRS for HOMA-IR were computed with LdPred <u>(Vilhjálmsson et al. 2015)</u>^8^ using summary statistics from Dupuis et al (Dupuis et al. 2010)^9^.

Process-specific PRSs were constructed based on five clusters defined in Udler et al. <u>(Udler et al. 2018)</u> by selecting the SNPs that had weight larger than 0.75 for each of a given cluster. We used the effect sizes described in Mahajan et al as weight for the polygenic scores <u>(Mahajan et al. 2018).</u>

All PRSs were tested for association with T2D and with BMI using the 30,240 MGB Biobank samples from European Ancestry defined based on self-reported and principal components.

## MGB Biobank cohort

The MGB Biobank <u>(Karlson et al. 2016)</u> maintains blood and DNA samples from more than 60,000 consented patients seen at Partners HealthCare hospitals, including Massachusetts General Hospital, Brigham and Women’s Hospital, McLean Hospital, and Spaulding Rehabilitation Hospital, all in the USA. Patients are recruited in the context of clinical care appointments at more than 40 sites, clinics and also electronically through the patient portal at Partners HealthCare. Biobank subjects provide consent for the use of their samples and data in broad-based research. The Partners Biobank works closely with the Partners Research Patient Data Registry (RPDR), the Partners’ enterprise scale data repository designed to foster investigator access to a wide variety of phenotypic data on more than 4 million Partners HealthCare patients. Approval for analysis of Biobank data was obtained by Partners IRB, study 2016P001018.

Type 2 diabetes status was defined based on “curated phenotypes” developed by the Biobank Portal team using both structured and unstructured electronic medical record (EMR) data and clinical, computational and statistical methods. Natural Language Processing (NLP) was used to extract data from narrative text. Chart reviews by disease experts helped identify features and variables associated with particular phenotypes and were also used to validate results of the algorithms. The process produced robust phenotype algorithms that were evaluated using metrics such as sensitivity, the proportion of true positives correctly identified as such, and positive predictive value (PPV), the proportion of individuals classified as cases by the algorithm <u>(Yu et al. 2015).</u>

a. Control selection criteria.

1. Individuals determined by the “curated disease” algorithm employed above to have no history of type 2 diabetes with NPV of 99%.
2. Individuals at least age 55.
3. Individuals with HbA1c less than 5.7
b. Case selection criteria.

1. Individuals determined by the “curated disease” algorithm employed above to have type 2 diabetes with PPV of 99%
2. Individuals at least age 30 given the higher rate of false positive diagnoses in younger individuals.

Genomic data for 30,240 participants was generated with the Illumina Multi-Ethnic Genotyping Array, which covers more than 1.7 million markers, including content from over 36,000 individuals, and is enriched for exome content with >400,000 markers missense, nonsense, indels, and synonymous variants.

For the individuals, we considered the following exclusion criteria: gender discordance, subject relatedness (pairs with PI-HAT ≥ 0.125 from which we removed the individual with the highest proportion of missingness), sample call rates ≥ 0.02 and population structure showing more than 4 standard deviations within the distribution of the study population according to the first seven principal components (Supplementary Figure 27).

For the individuals, we considered the following exclusion criteria: gender discordance, subject relatedness (pairs with PI-HAT ≥ 0.125 from which we removed the individual with the highest proportion of missingness), sample call rates ≥ 0.02 and population structure showing more than 4 standard deviations within the distribution of the study population according to the first seven principal components (Supplementary Figure 27).

---

**Supplemental Table 1:**
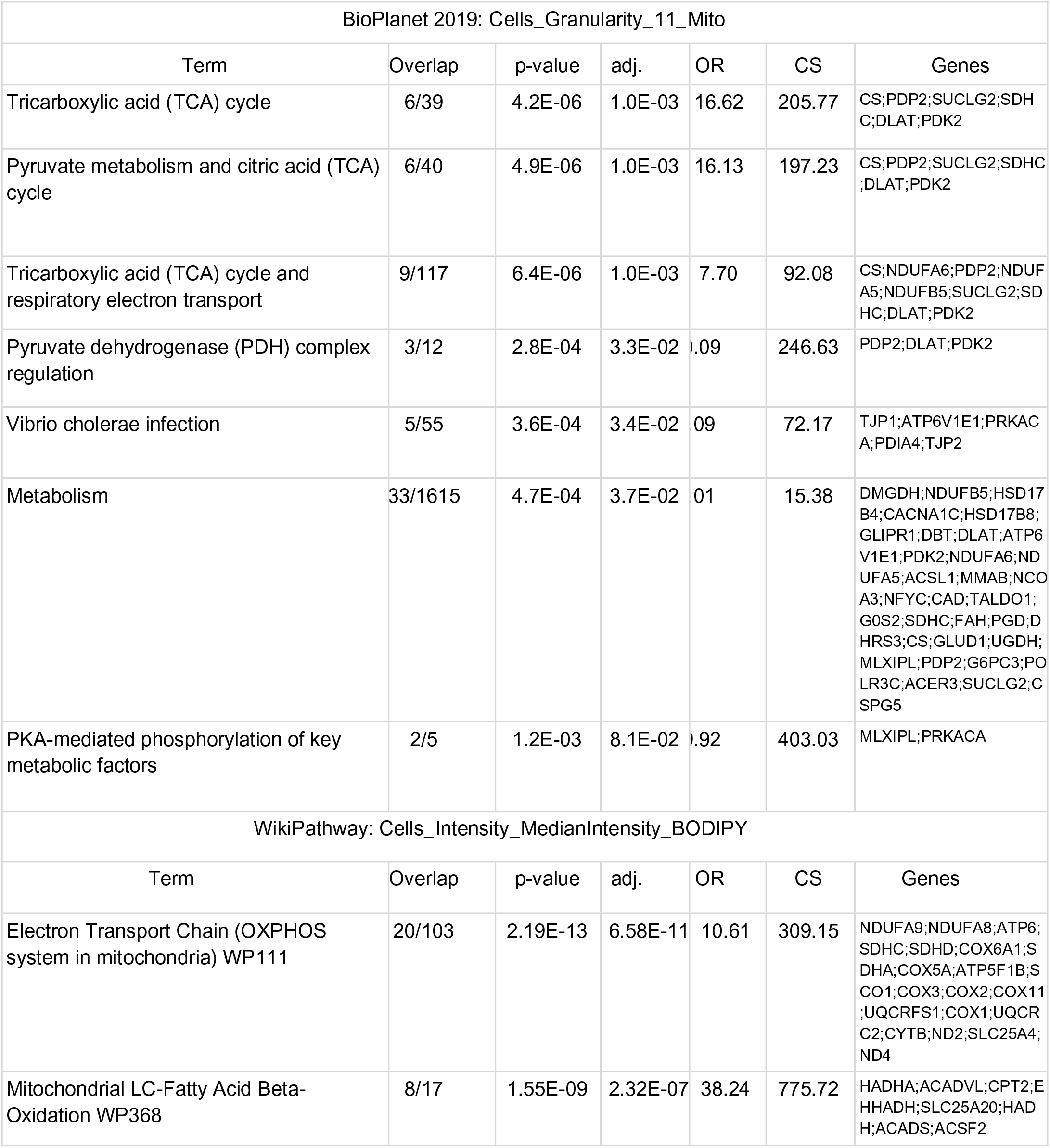
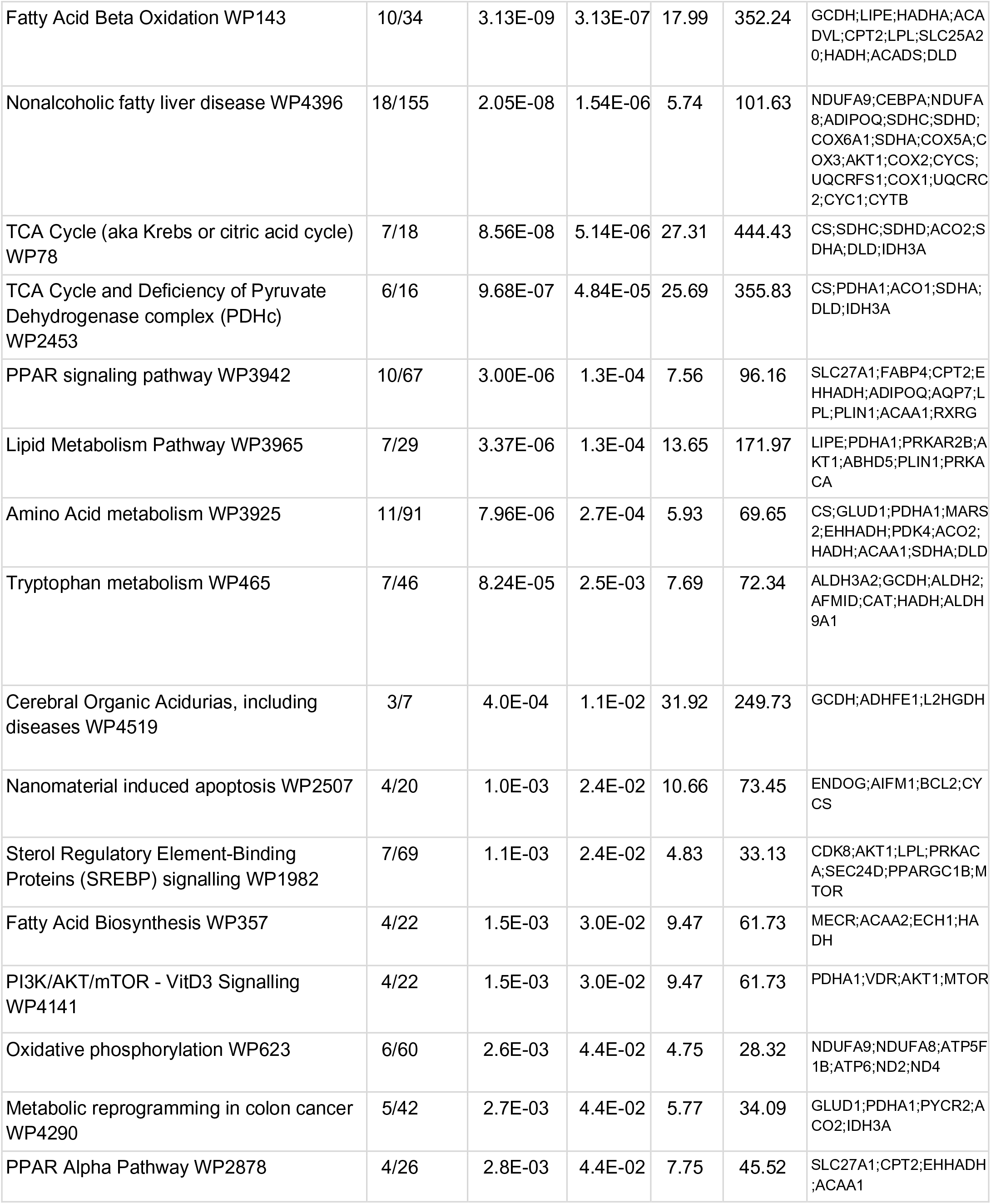
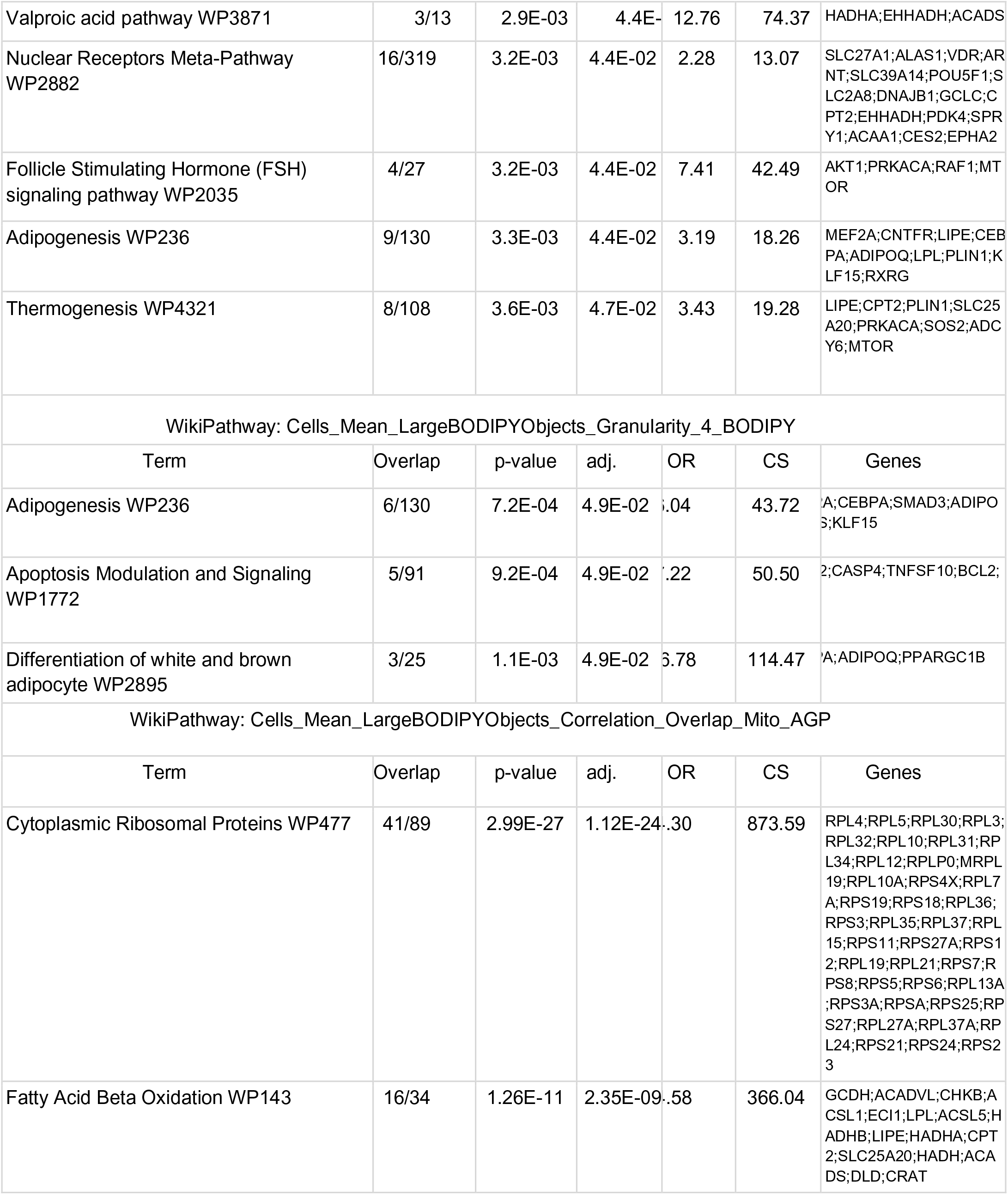
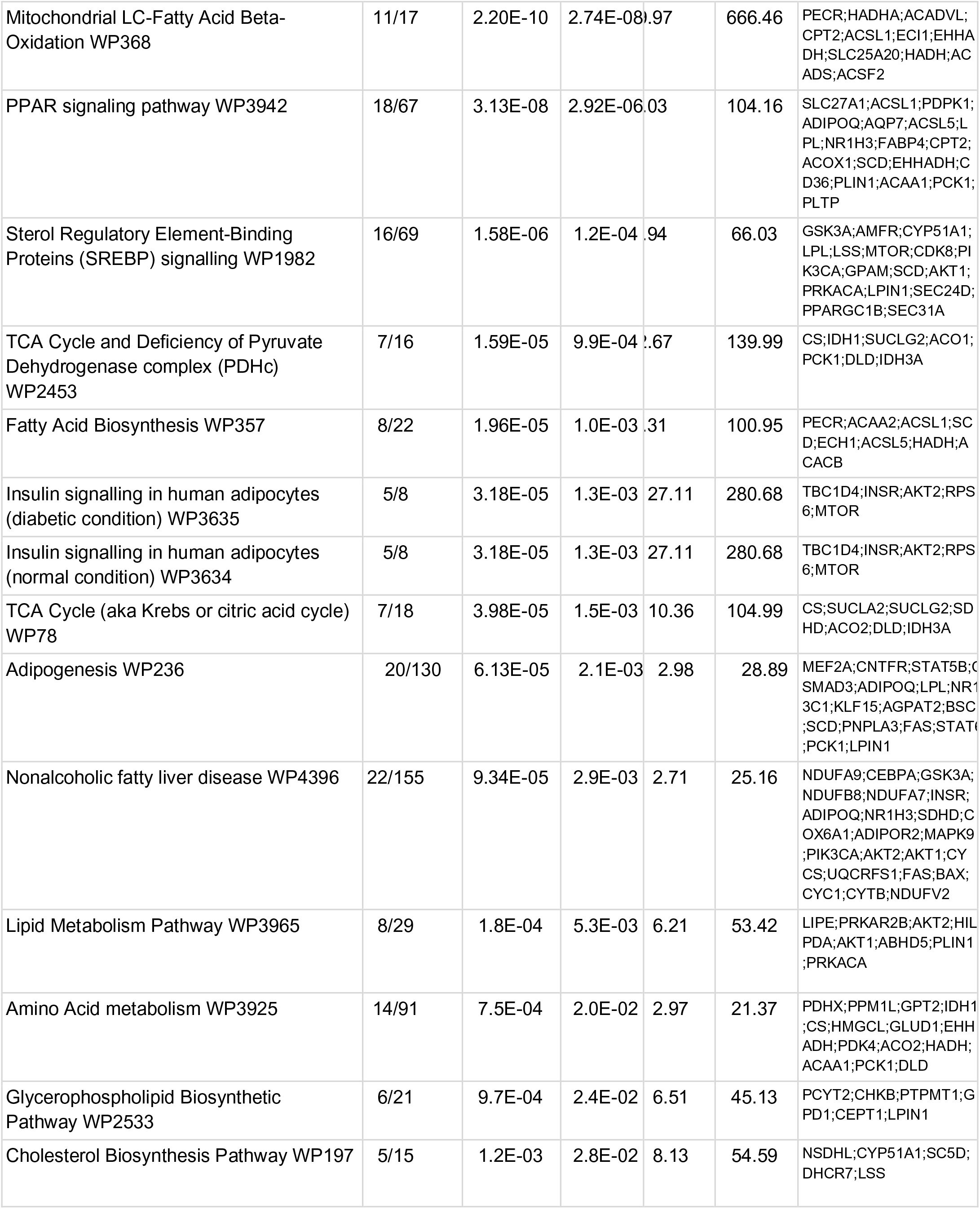
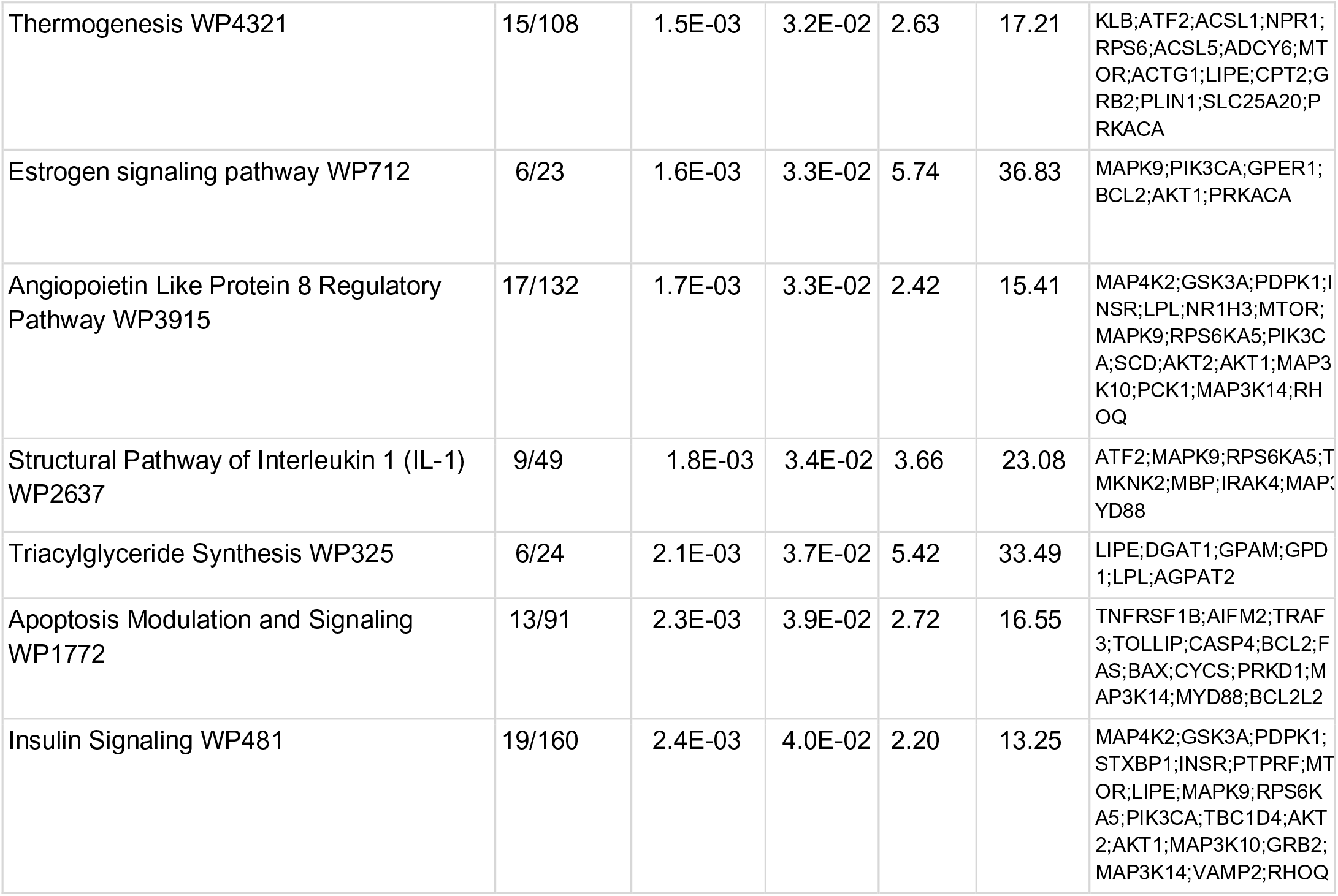
Lists of pathways enriched among representative significant gene LP-features connections. Term, which pathway; Overlap, number of genes that overlap and total genes; p-value, enrichment p-value; adj., adjusted p-value, q-value; OR, odds ratio, enrichment; CS, combined score, approximation of overall association (-log10(P) * log(Odds)), Genes, genes in the pathway which are associated with gene LP-feature connections.

**Supplemental Table 2:**
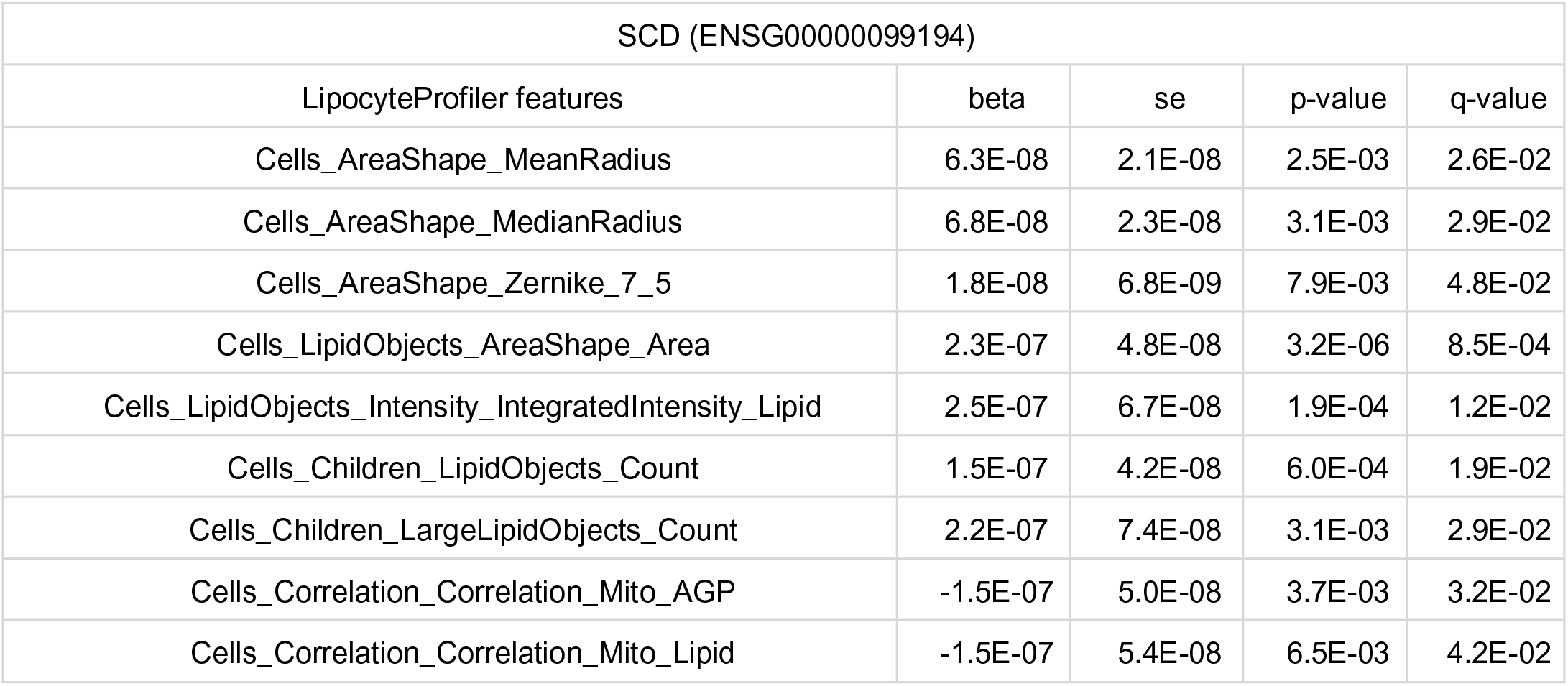
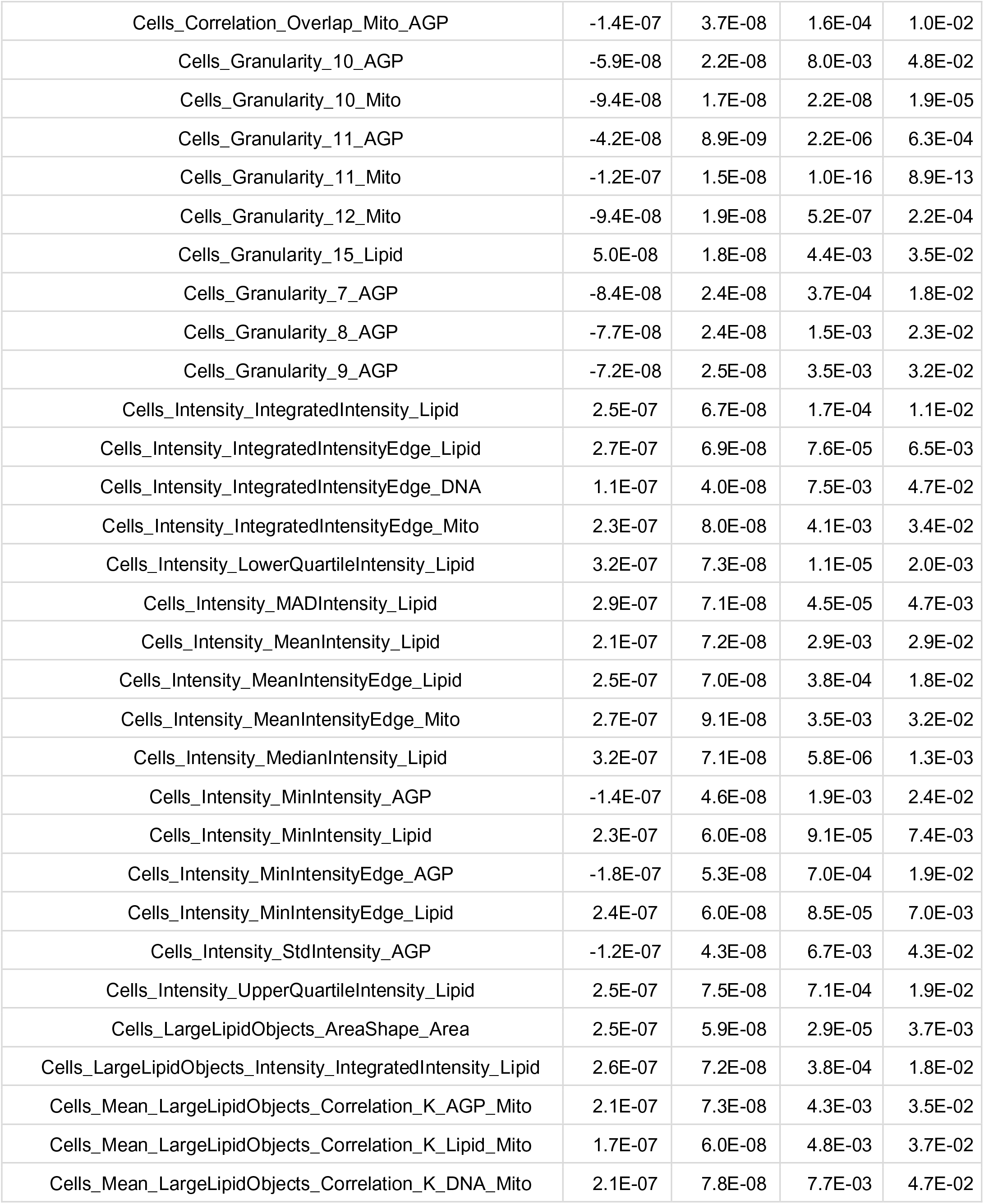
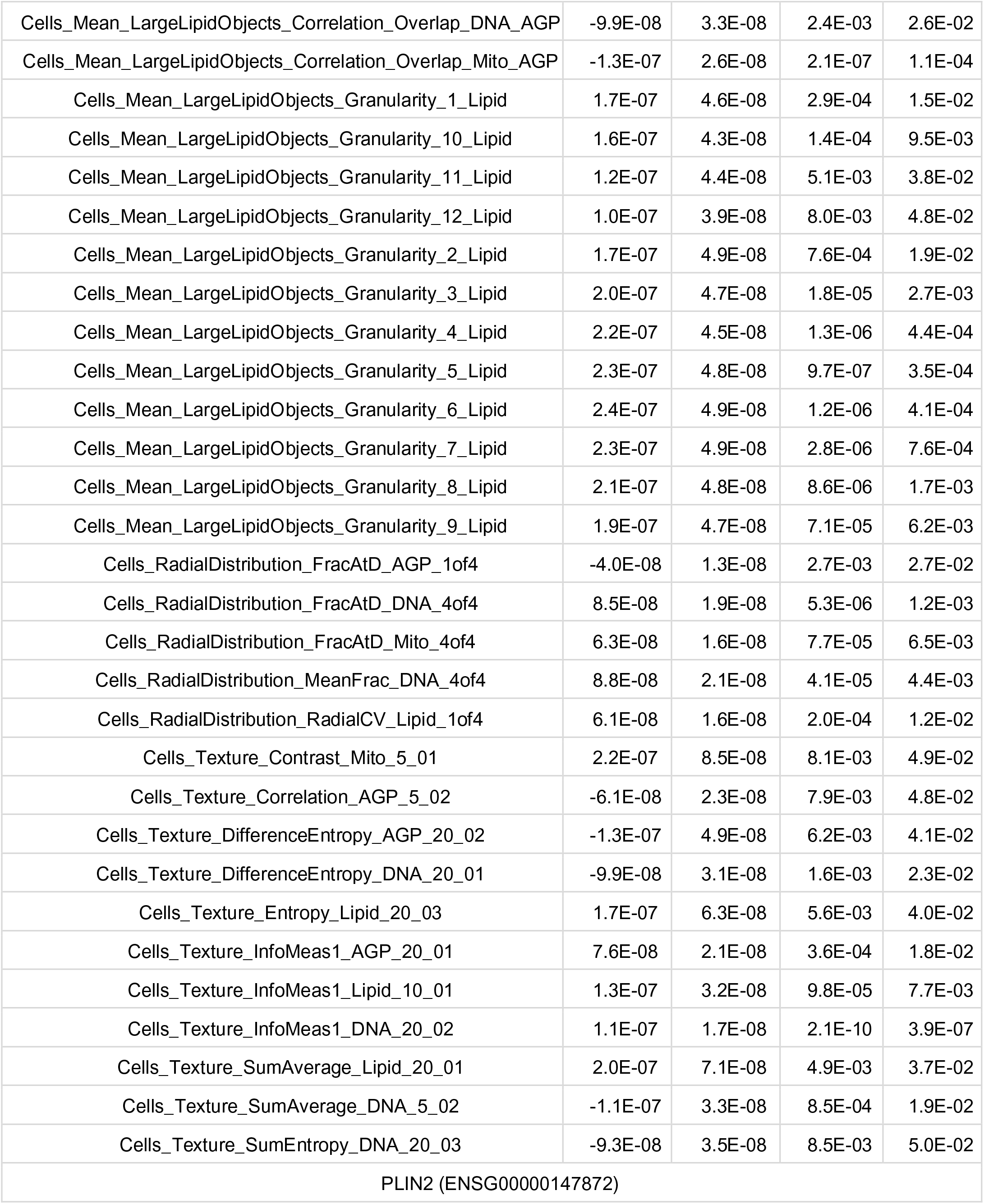
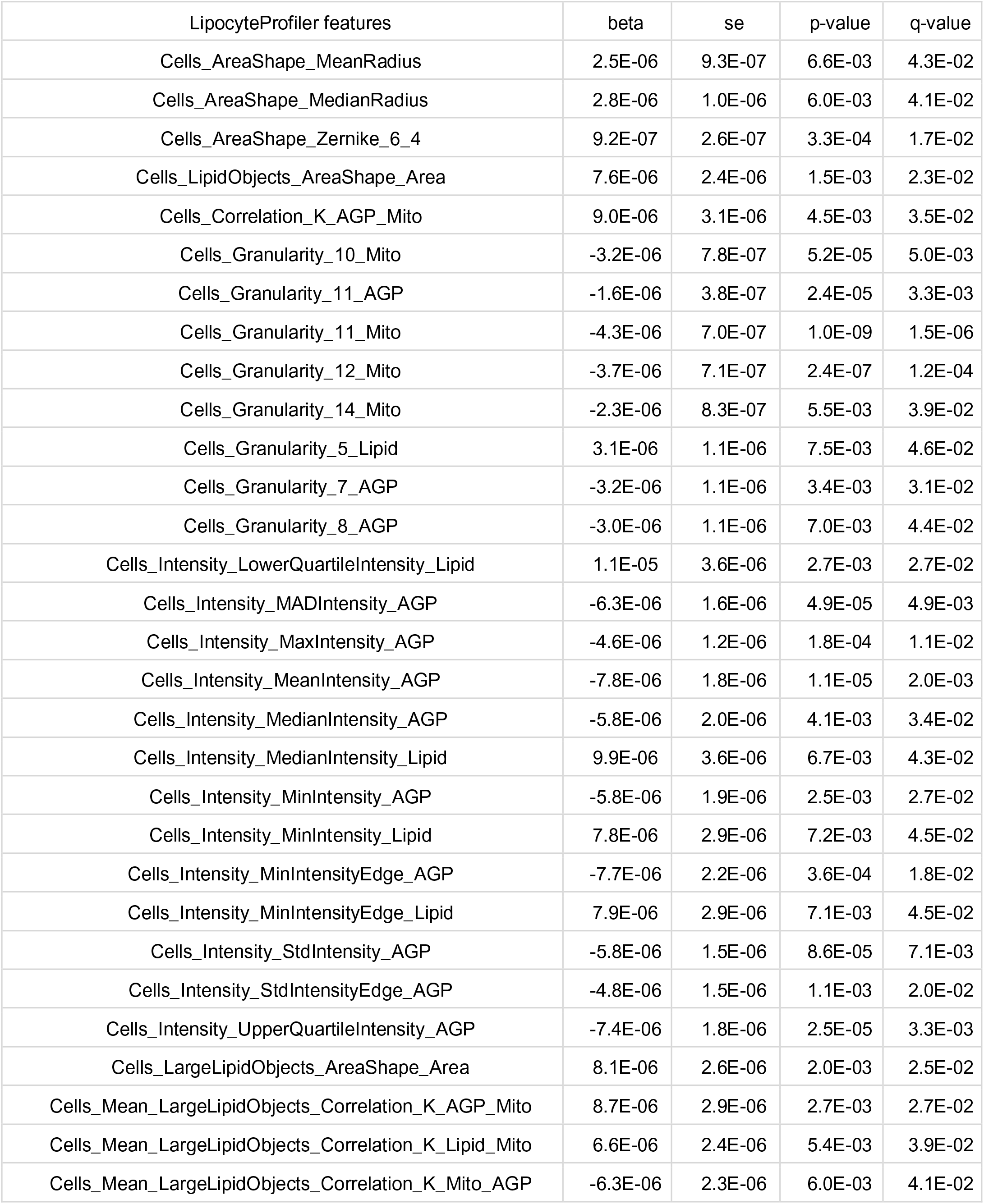
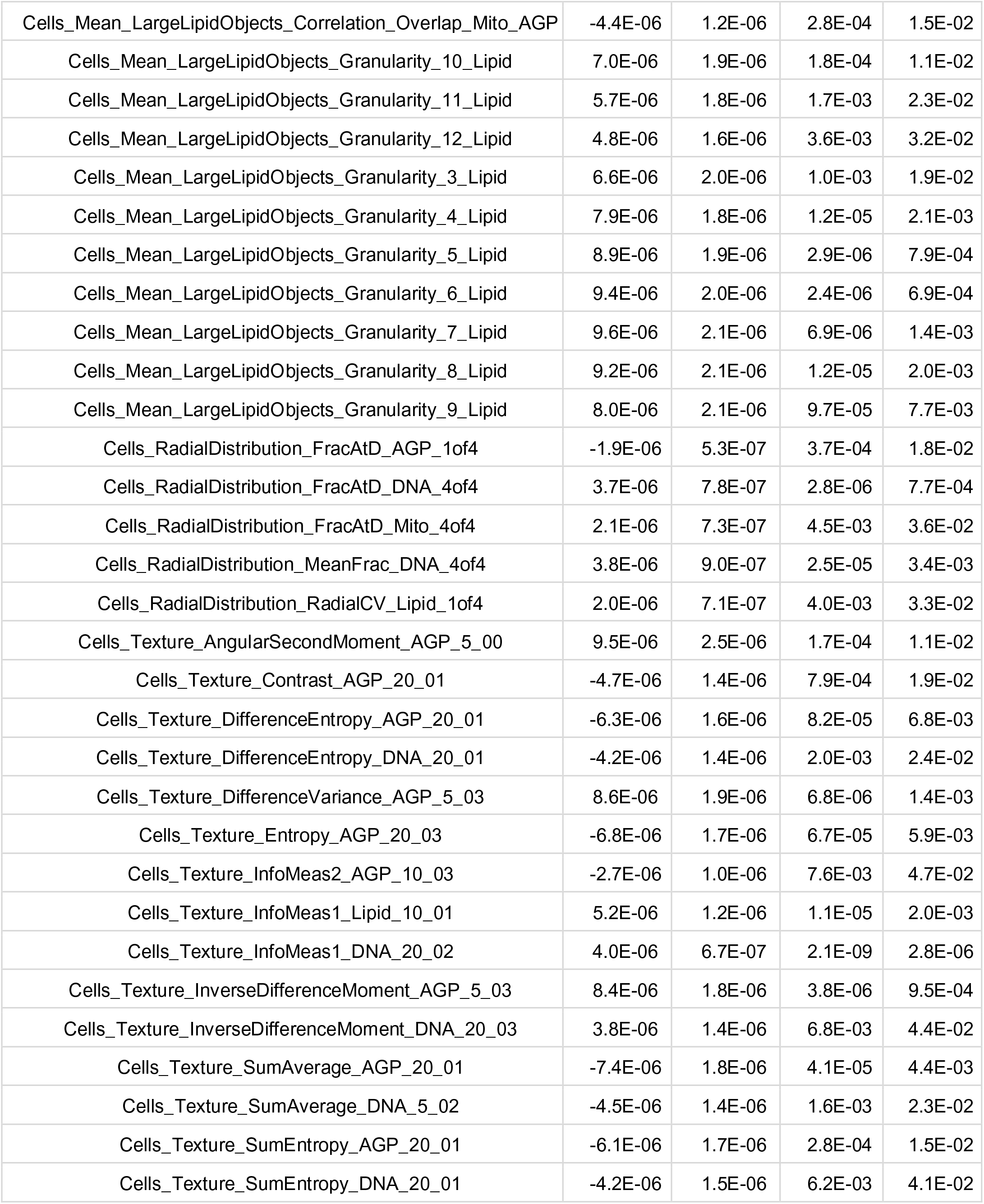
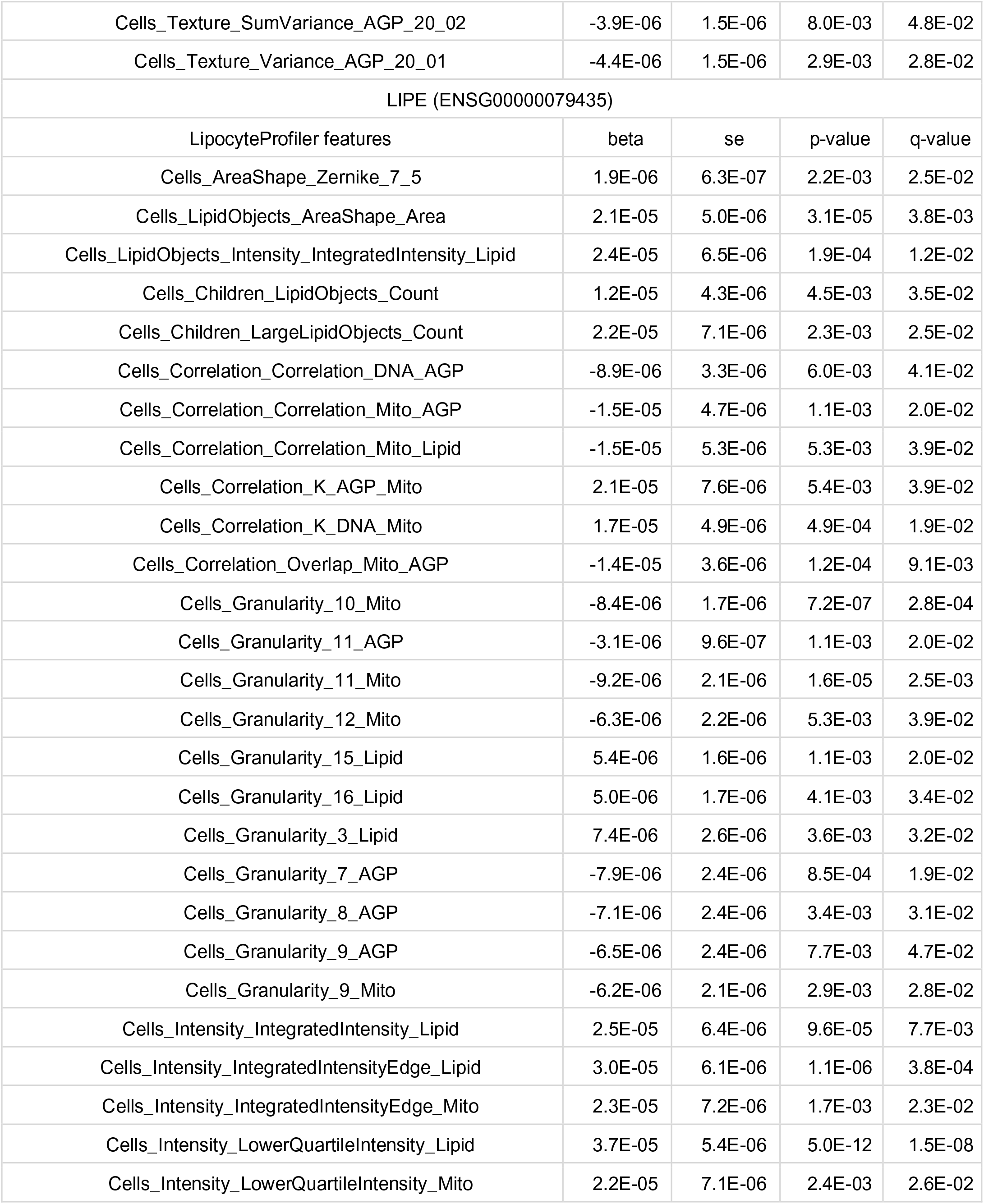
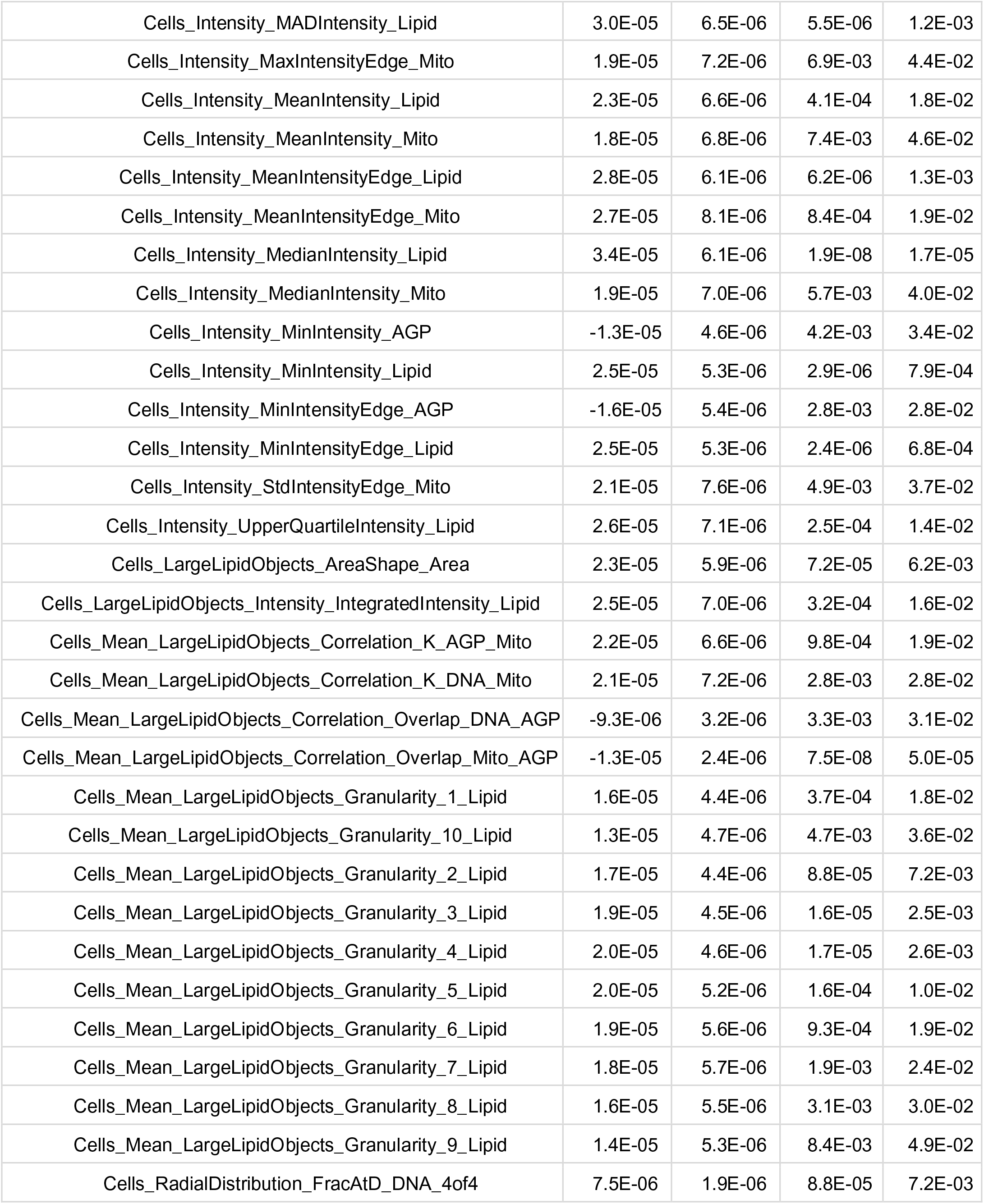
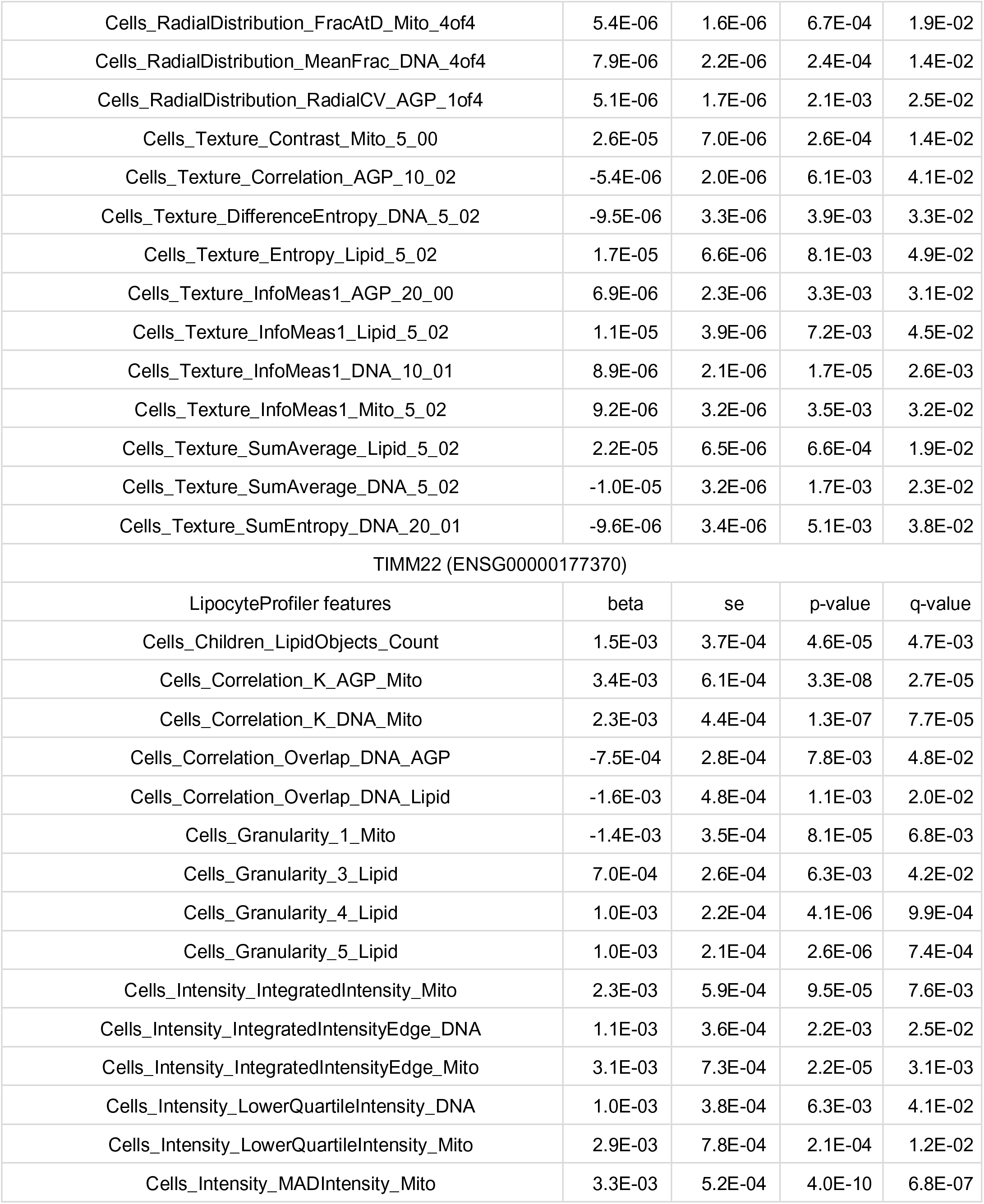
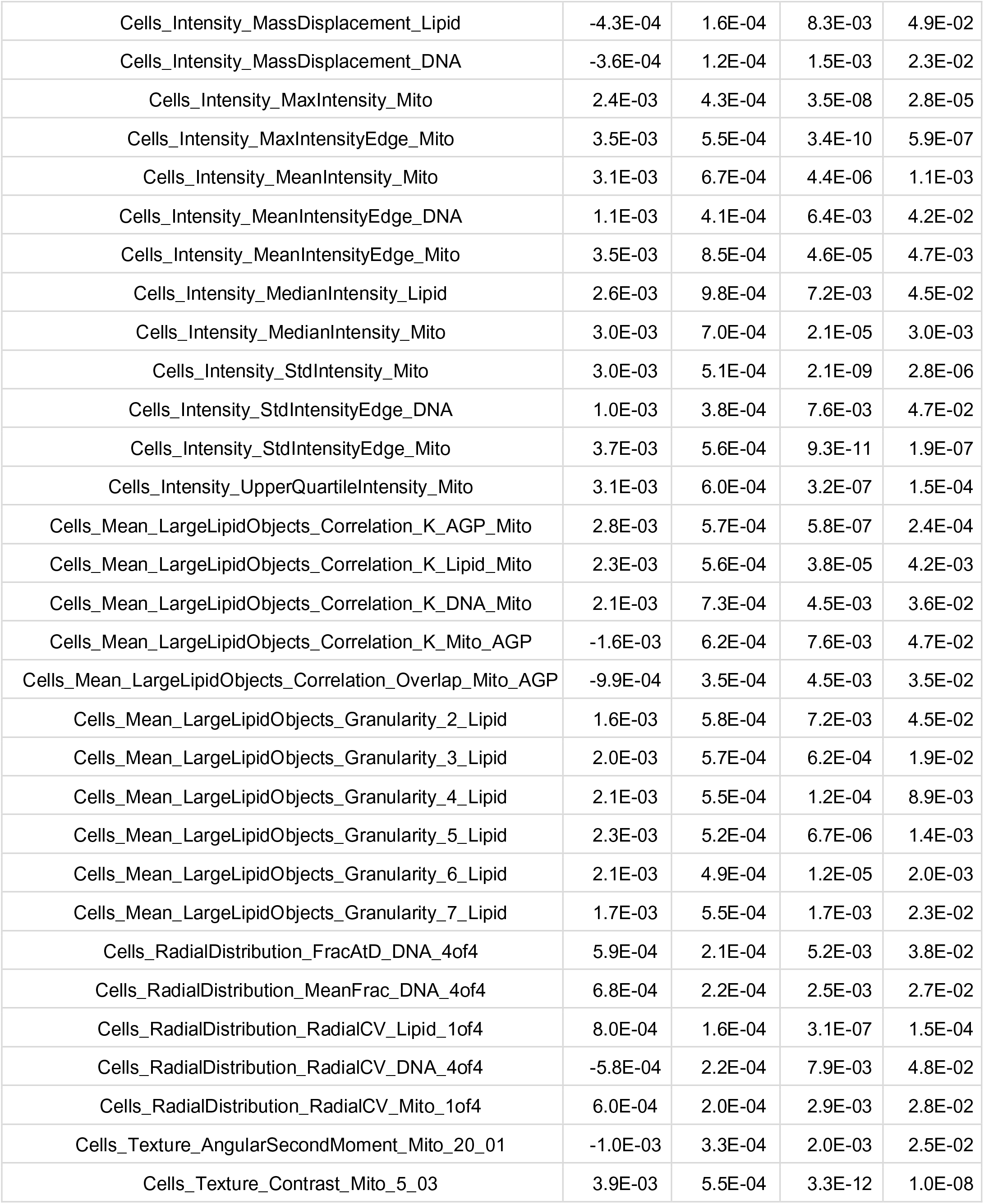
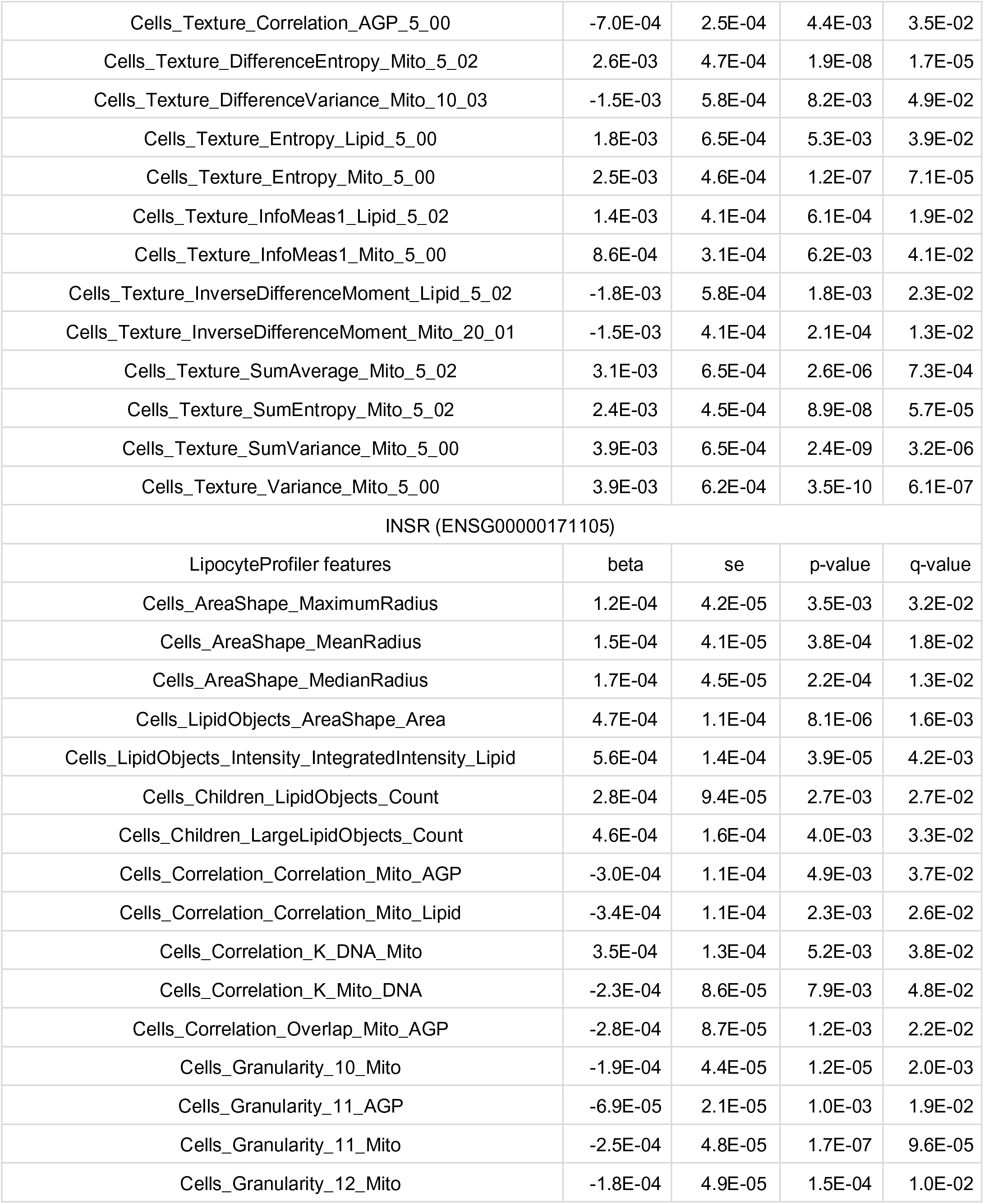
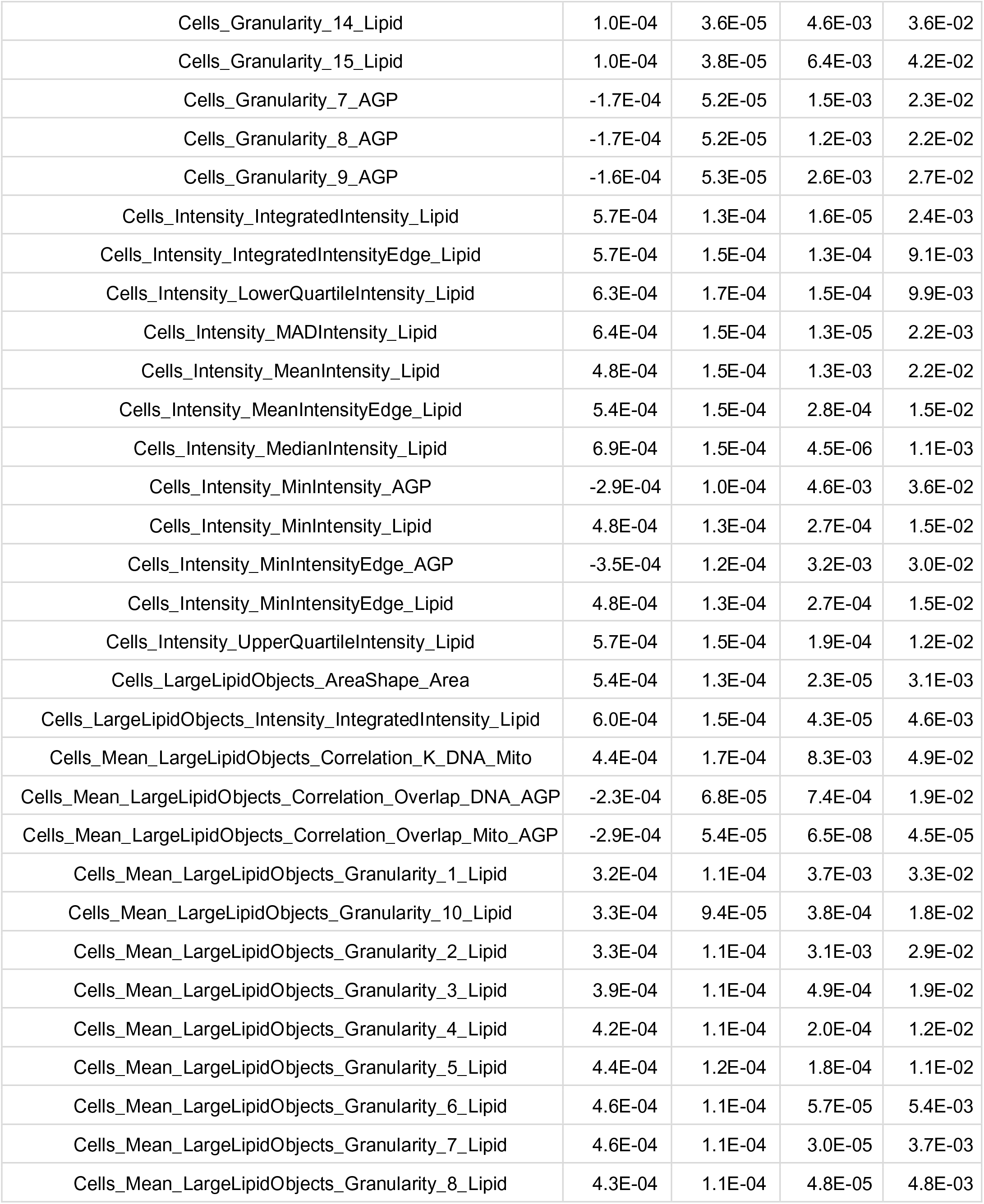
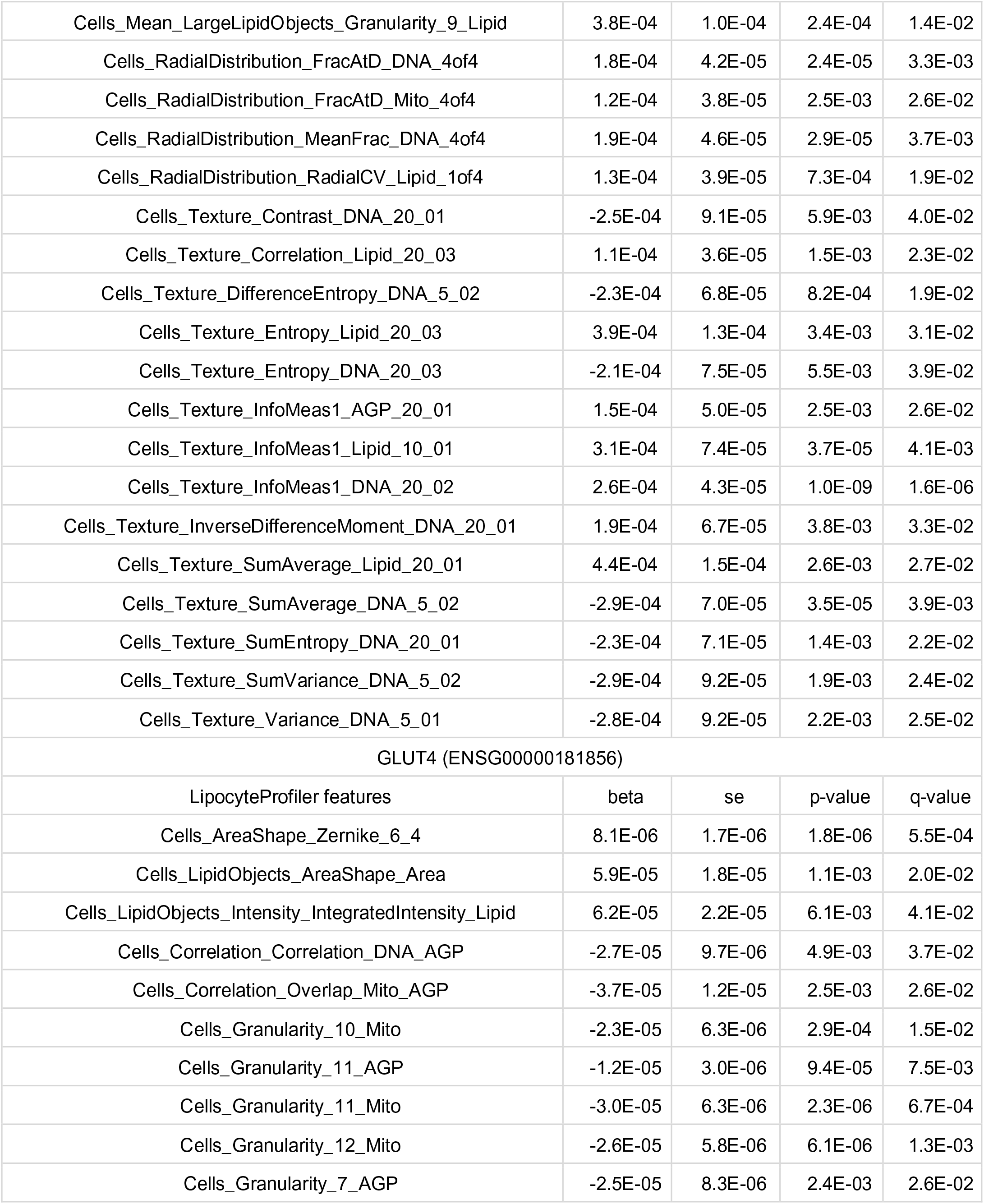
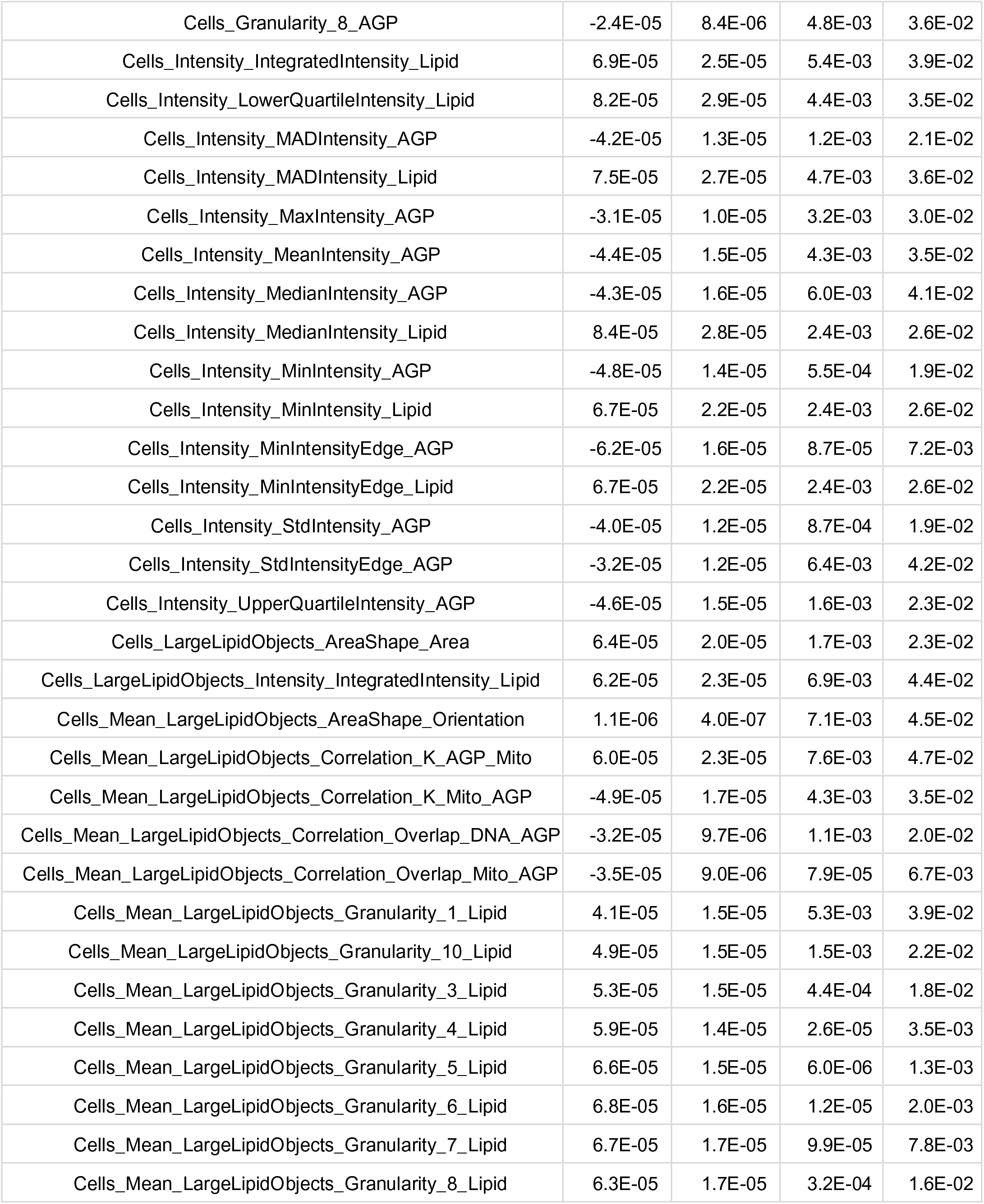
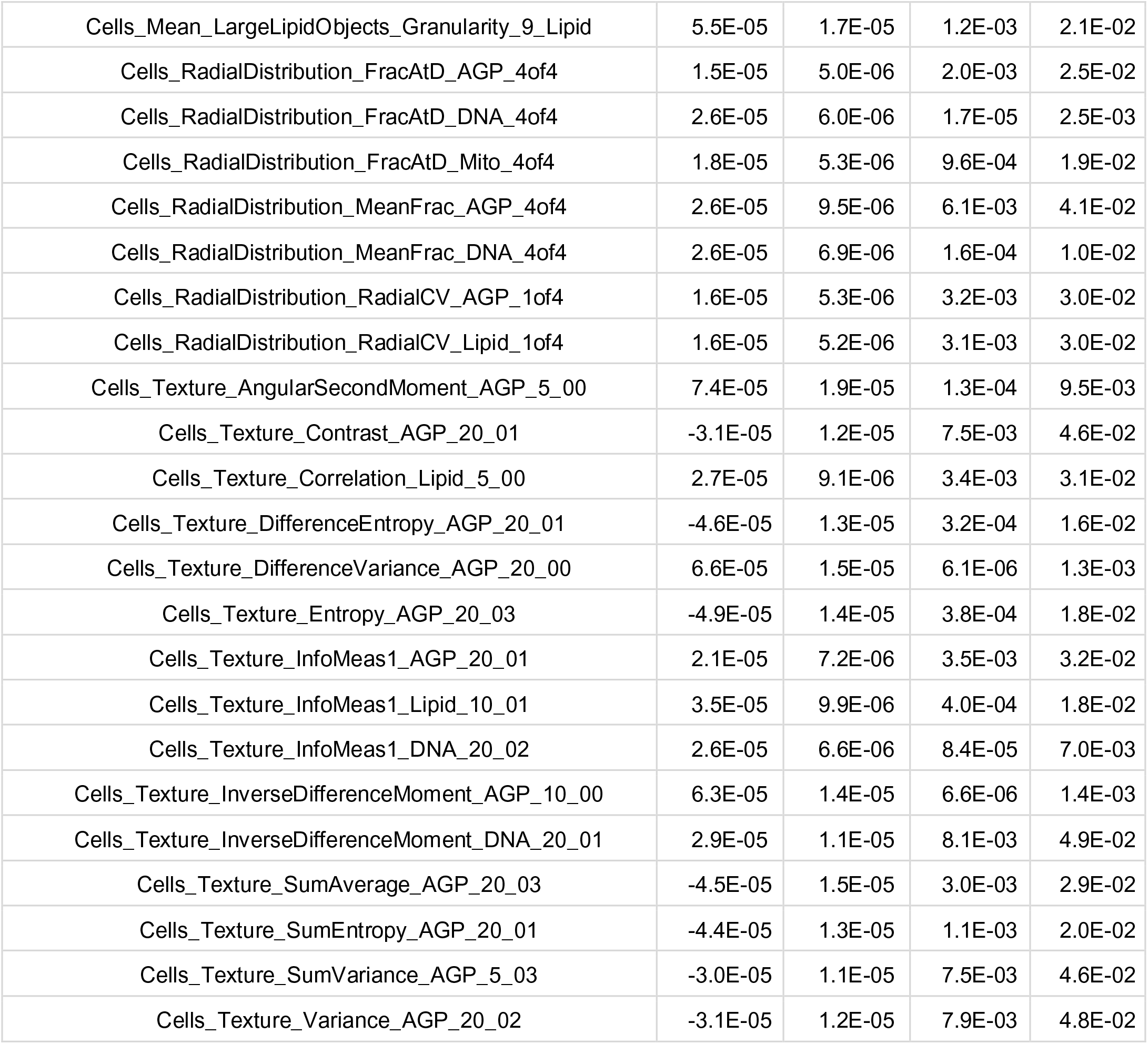
Morphological signatures of adipocyte marker genes SCD, PLIN2, LIPE, TIMM22, INSR and GLUT4 recapitulate their cellular function. LMM output (significance level FDR 5%). Beta, beta of LMM; se, standard error of LMM, p-value of LMM, q-value of LMM

| SCD (ENSG00000099194) |  |  |  |  |
| --- | --- | --- | --- | --- |
| LipocyteProfiler features | beta | se | p-value | q-value |
| Cells_AreaShape_MeanRadius | 6.3E-08 | 2.1E-08 | 2.5E-03 | 2.6E-02 |
| Cells_AreaShape_MedianRadius | 6.8E-08 | 2.3E-08 | 3.1E-03 | 2.9E-02 |
| Cells_AreaShape_Zernike_7_5 | 1.8E-08 | 6.8E-09 | 7.9E-03 | 4.8E-02 |
| Cells_LipidObjects_AreaShape_Area | 2.3E-07 | 4.8E-08 | 3.2E-06 | 8.5E-04 |
| Cells_LipidObjects_Intensity_IntegratedIntensity_Lipid | 2.5E-07 | 6.7E-08 | 1.9E-04 | 1.2E-02 |
| Cells_Children_LipidObjects_Count | 1.5E-07 | 4.2E-08 | 6.0E-04 | 1.9E-02 |
| Cells_Children_LargeLipidObjects_Count | 2.2E-07 | 7.4E-08 | 3.1E-03 | 2.9E-02 |
| Cells_Correlation_Correlation_Mito_AGP | -1.5E-07 | 5.0E-08 | 3.7E-03 | 3.2E-02 |
| Cells_Correlation_Correlation_Mito_Lipid | -1.5E-07 | 5.4E-08 | 6.5E-03 | 4.2E-02 |
| Cells_Correlation_Overlap_Mito_AGP | -1.4E-07 | 3.7E-08 | 1.6E-04 | 1.0E-02 |
| Cells_Granularity_10_AGP | -5.9E-08 | 2.2E-08 | 8.0E-03 | 4.8E-02 |
| Cells_Granularity_10_Mito | -9.4E-08 | 1.7E-08 | 2.2E-08 | 1.9E-05 |
| Cells_Granularity_11_AGP | -4.2E-08 | 8.9E-09 | 2.2E-06 | 6.3E-04 |
| Cells_Granularity_11_Mito | -1.2E-07 | 1.5E-08 | 1.0E-16 | 8.9E-13 |
| Cells_Granularity_12_Mito | -9.4E-08 | 1.9E-08 | 5.2E-07 | 2.2E-04 |
| Cells_Granularity_15_Lipid | 5.0E-08 | 1.8E-08 | 4.4E-03 | 3.5E-02 |
| Cells_Granularity_7_AGP | -8.4E-08 | 2.4E-08 | 3.7E-04 | 1.8E-02 |
| Cells_Granularity_8_AGP | -7.7E-08 | 2.4E-08 | 1.5E-03 | 2.3E-02 |
| Cells_Granularity_9_AGP | -7.2E-08 | 2.5E-08 | 3.5E-03 | 3.2E-02 |
| Cells_Intensity_IntegratedIntensity_Lipid | 2.5E-07 | 6.7E-08 | 1.7E-04 | 1.1E-02 |
| Cells_Intensity_IntegratedIntensityEdge_Lipid | 2.7E-07 | 6.9E-08 | 7.6E-05 | 6.5E-03 |
| Cells_Intensity_IntegratedIntensityEdge_DNA | 1.1E-07 | 4.0E-08 | 7.5E-03 | 4.7E-02 |
| Cells_Intensity_IntegratedIntensityEdge_Mito | 2.3E-07 | 8.0E-08 | 4.1E-03 | 3.4E-02 |
| Cells_Intensity_LowerQuartileIntensity_Lipid | 3.2E-07 | 7.3E-08 | 1.1E-05 | 2.0E-03 |
| Cells_Intensity_MADIntensity_Lipid | 2.9E-07 | 7.1E-08 | 4.5E-05 | 4.7E-03 |
| Cells_Intensity_MeanIntensity_Lipid | 2.1E-07 | 7.2E-08 | 2.9E-03 | 2.9E-02 |
| Cells_Intensity_MeanIntensityEdge_Lipid | 2.5E-07 | 7.0E-08 | 3.8E-04 | 1.8E-02 |
| Cells_Intensity_MeanIntensityEdge_Mito | 2.7E-07 | 9.1E-08 | 3.5E-03 | 3.2E-02 |
| Cells_Intensity_MedianIntensity_Lipid | 3.2E-07 | 7.1E-08 | 5.8E-06 | 1.3E-03 |
| Cells_Intensity_MinIntensity_AGP | -1.4E-07 | 4.6E-08 | 1.9E-03 | 2.4E-02 |
| Cells_Intensity_MinIntensity_Lipid | 2.3E-07 | 6.0E-08 | 9.1E-05 | 7.4E-03 |
| Cells_Intensity_MinIntensityEdge_AGP | -1.8E-07 | 5.3E-08 | 7.0E-04 | 1.9E-02 |
| Cells_Intensity_MinIntensityEdge_Lipid | 2.4E-07 | 6.0E-08 | 8.5E-05 | 7.0E-03 |
| Cells_Intensity_StdIntensity_AGP | -1.2E-07 | 4.3E-08 | 6.7E-03 | 4.3E-02 |
| Cells_Intensity_UpperQuartileIntensity_Lipid | 2.5E-07 | 7.5E-08 | 7.1E-04 | 1.9E-02 |
| Cells_LargeLipidObjects_AreaShape_Area | 2.5E-07 | 5.9E-08 | 2.9E-05 | 3.7E-03 |
| Cells_LargeLipidObjects_Intensity_IntegratedIntensity_Lipid | 2.6E-07 | 7.2E-08 | 3.8E-04 | 1.8E-02 |
| Cells_Mean_LargeLipidObjects_Correlation_K_AGP_Mito | 2.1E-07 | 7.3E-08 | 4.3E-03 | 3.5E-02 |
| Cells_Mean_LargeLipidObjects_Correlation_K_Lipid_Mito | 1.7E-07 | 6.0E-08 | 4.8E-03 | 3.7E-02 |
| Cells_Mean_LargeLipidObjects_Correlation_K_DNA_Mito | 2.1E-07 | 7.8E-08 | 7.7E-03 | 4.7E-02 |
| Cells_Mean_LargeLipidObjects_Correlation_Overlap_DNA_AGP | -9.9E-08 | 3.3E-08 | 2.4E-03 | 2.6E-02 |
| Cells_Mean_LargeLipidObjects_Correlation_Overlap_Mito_AGP | -1.3E-07 | 2.6E-08 | 2.1E-07 | 1.1E-04 |
| Cells_Mean_LargeLipidObjects_Granularity_1_Lipid | 1.7E-07 | 4.6E-08 | 2.9E-04 | 1.5E-02 |
| Cells_Mean_LargeLipidObjects_Granularity_10_Lipid | 1.6E-07 | 4.3E-08 | 1.4E-04 | 9.5E-03 |
| Cells_Mean_LargeLipidObjects_Granularity_11_Lipid | 1.2E-07 | 4.4E-08 | 5.1E-03 | 3.8E-02 |
| Cells_Mean_LargeLipidObjects_Granularity_12_Lipid | 1.0E-07 | 3.9E-08 | 8.0E-03 | 4.8E-02 |
| Cells_Mean_LargeLipidObjects_Granularity_2_Lipid | 1.7E-07 | 4.9E-08 | 7.6E-04 | 1.9E-02 |
| Cells_Mean_LargeLipidObjects_Granularity_3_Lipid | 2.0E-07 | 4.7E-08 | 1.8E-05 | 2.7E-03 |
| Cells_Mean_LargeLipidObjects_Granularity_4_Lipid | 2.2E-07 | 4.5E-08 | 1.3E-06 | 4.4E-04 |
| Cells_Mean_LargeLipidObjects_Granularity_5_Lipid | 2.3E-07 | 4.8E-08 | 9.7E-07 | 3.5E-04 |
| Cells_Mean_LargeLipidObjects_Granularity_6_Lipid | 2.4E-07 | 4.9E-08 | 1.2E-06 | 4.1E-04 |
| Cells_Mean_LargeLipidObjects_Granularity_7_Lipid | 2.3E-07 | 4.9E-08 | 2.8E-06 | 7.6E-04 |
| Cells_Mean_LargeLipidObjects_Granularity_8_Lipid | 2.1E-07 | 4.8E-08 | 8.6E-06 | 1.7E-03 |
| Cells_Mean_LargeLipidObjects_Granularity_9_Lipid | 1.9E-07 | 4.7E-08 | 7.1E-05 | 6.2E-03 |
| Cells_RadialDistribution_FracAtD_AGP_1of4 | -4.0E-08 | 1.3E-08 | 2.7E-03 | 2.7E-02 |
| Cells_RadialDistribution_FracAtD_DNA_4of4 | 8.5E-08 | 1.9E-08 | 5.3E-06 | 1.2E-03 |
| Cells_RadialDistribution_FracAtD_Mito_4of4 | 6.3E-08 | 1.6E-08 | 7.7E-05 | 6.5E-03 |
| Cells_RadialDistribution_MeanFrac_DNA_4of4 | 8.8E-08 | 2.1E-08 | 4.1E-05 | 4.4E-03 |
| Cells_RadialDistribution_RadialCV_Lipid_1of4 | 6.1E-08 | 1.6E-08 | 2.0E-04 | 1.2E-02 |
| Cells_Texture_Contrast_Mito_5_01 | 2.2E-07 | 8.5E-08 | 8.1E-03 | 4.9E-02 |
| Cells_Texture_Correlation_AGP_5_02 | -6.1E-08 | 2.3E-08 | 7.9E-03 | 4.8E-02 |
| Cells_Texture_DifferenceEntropy_AGP_20_02 | -1.3E-07 | 4.9E-08 | 6.2E-03 | 4.1E-02 |
| Cells_Texture_DifferenceEntropy_DNA_20_01 | -9.9E-08 | 3.1E-08 | 1.6E-03 | 2.3E-02 |
| Cells_Texture_Entropy_Lipid_20_03 | 1.7E-07 | 6.3E-08 | 5.6E-03 | 4.0E-02 |
| Cells_Texture_InfoMeas1_AGP_20_01 | 7.6E-08 | 2.1E-08 | 3.6E-04 | 1.8E-02 |
| Cells_Texture_InfoMeas1_Lipid_10_01 | 1.3E-07 | 3.2E-08 | 9.8E-05 | 7.7E-03 |
| Cells_Texture_InfoMeas1_DNA_20_02 | 1.1E-07 | 1.7E-08 | 2.1E-10 | 3.9E-07 |
| Cells_Texture_SumAverage_Lipid_20_01 | 2.0E-07 | 7.1E-08 | 4.9E-03 | 3.7E-02 |
| Cells_Texture_SumAverage_DNA_5_02 | -1.1E-07 | 3.3E-08 | 8.5E-04 | 1.9E-02 |
| Cells_Texture_SumEntropy_DNA_20_03 | -9.3E-08 | 3.5E-08 | 8.5E-03 | 5.0E-02 |
| PLIN2 (ENSG00000147872) |  |  |  |  |

| LipocyteProfiler features | beta | se | p-value | q-value |
| --- | --- | --- | --- | --- |
| Cells_AreaShape_MeanRadius | 2.5E-06 | 9.3E-07 | 6.6E-03 | 4.3E-02 |
| Cells_AreaShape_MedianRadius | 2.8E-06 | 1.0E-06 | 6.0E-03 | 4.1E-02 |
| Cells_AreaShape_Zernike_6_4 | 9.2E-07 | 2.6E-07 | 3.3E-04 | 1.7E-02 |
| Cells_LipidObjects_AreaShape_Area | 7.6E-06 | 2.4E-06 | 1.5E-03 | 2.3E-02 |
| Cells_Correlation_K_AGP_Mito | 9.0E-06 | 3.1E-06 | 4.5E-03 | 3.5E-02 |
| Cells_Granularity_10_Mito | -3.2E-06 | 7.8E-07 | 5.2E-05 | 5.0E-03 |
| Cells_Granularity_11_AGP | -1.6E-06 | 3.8E-07 | 2.4E-05 | 3.3E-03 |
| Cells_Granularity_11_Mito | -4.3E-06 | 7.0E-07 | 1.0E-09 | 1.5E-06 |
| Cells_Granularity_12_Mito | -3.7E-06 | 7.1E-07 | 2.4E-07 | 1.2E-04 |
| Cells_Granularity_14_Mito | -2.3E-06 | 8.3E-07 | 5.5E-03 | 3.9E-02 |
| Cells_Granularity_5_Lipid | 3.1E-06 | 1.1E-06 | 7.5E-03 | 4.6E-02 |
| Cells_Granularity_7_AGP | -3.2E-06 | 1.1E-06 | 3.4E-03 | 3.1E-02 |
| Cells_Granularity_8_AGP | -3.0E-06 | 1.1E-06 | 7.0E-03 | 4.4E-02 |
| Cells_Intensity_LowerQuartileIntensity_Lipid | 1.1E-05 | 3.6E-06 | 2.7E-03 | 2.7E-02 |
| Cells_Intensity_MADIntensity_AGP | -6.3E-06 | 1.6E-06 | 4.9E-05 | 4.9E-03 |
| Cells_Intensity_MaxIntensity_AGP | -4.6E-06 | 1.2E-06 | 1.8E-04 | 1.1E-02 |
| Cells_Intensity_MeanIntensity_AGP | -7.8E-06 | 1.8E-06 | 1.1E-05 | 2.0E-03 |
| Cells_Intensity_MedianIntensity_AGP | -5.8E-06 | 2.0E-06 | 4.1E-03 | 3.4E-02 |
| Cells_Intensity_MedianIntensity_Lipid | 9.9E-06 | 3.6E-06 | 6.7E-03 | 4.3E-02 |
| Cells_Intensity_MinIntensity_AGP | -5.8E-06 | 1.9E-06 | 2.5E-03 | 2.7E-02 |
| Cells_Intensity_MinIntensity_Lipid | 7.8E-06 | 2.9E-06 | 7.2E-03 | 4.5E-02 |
| Cells_Intensity_MinIntensityEdge_AGP | -7.7E-06 | 2.2E-06 | 3.6E-04 | 1.8E-02 |
| Cells_Intensity_MinIntensityEdge_Lipid | 7.9E-06 | 2.9E-06 | 7.1E-03 | 4.5E-02 |
| Cells_Intensity_StdIntensity_AGP | -5.8E-06 | 1.5E-06 | 8.6E-05 | 7.1E-03 |
| Cells_Intensity_StdIntensityEdge_AGP | -4.8E-06 | 1.5E-06 | 1.1E-03 | 2.0E-02 |
| Cells_Intensity_UpperQuartileIntensity_AGP | -7.4E-06 | 1.8E-06 | 2.5E-05 | 3.3E-03 |
| Cells_LargeLipidObjects_AreaShape_Area | 8.1E-06 | 2.6E-06 | 2.0E-03 | 2.5E-02 |
| Cells_Mean_LargeLipidObjects_Correlation_K_AGP_Mito | 8.7E-06 | 2.9E-06 | 2.7E-03 | 2.7E-02 |
| Cells_Mean_LargeLipidObjects_Correlation_K_Lipid_Mito | 6.6E-06 | 2.4E-06 | 5.4E-03 | 3.9E-02 |
| Cells_Mean_LargeLipidObjects_Correlation_K_Mito_AGP | -6.3E-06 | 2.3E-06 | 6.0E-03 | 4.1E-02 |
| Cells_Mean_LargeLipidObjects_Correlation_Overlap_Mito_AGP | -4.4E-06 | 1.2E-06 | 2.8E-04 | 1.5E-02 |
| Cells_Mean_LargeLipidObjects_Granularity_10_Lipid | 7.0E-06 | 1.9E-06 | 1.8E-04 | 1.1E-02 |
| Cells_Mean_LargeLipidObjects_Granularity_11_Lipid | 5.7E-06 | 1.8E-06 | 1.7E-03 | 2.3E-02 |
| Cells_Mean_LargeLipidObjects_Granularity_12_Lipid | 4.8E-06 | 1.6E-06 | 3.6E-03 | 3.2E-02 |
| Cells_Mean_LargeLipidObjects_Granularity_3_Lipid | 6.6E-06 | 2.0E-06 | 1.0E-03 | 1.9E-02 |
| Cells_Mean_LargeLipidObjects_Granularity_4_Lipid | 7.9E-06 | 1.8E-06 | 1.2E-05 | 2.1E-03 |
| Cells_Mean_LargeLipidObjects_Granularity_5_Lipid | 8.9E-06 | 1.9E-06 | 2.9E-06 | 7.9E-04 |
| Cells_Mean_LargeLipidObjects_Granularity_6_Lipid | 9.4E-06 | 2.0E-06 | 2.4E-06 | 6.9E-04 |
| Cells_Mean_LargeLipidObjects_Granularity_7_Lipid | 9.6E-06 | 2.1E-06 | 6.9E-06 | 1.4E-03 |
| Cells_Mean_LargeLipidObjects_Granularity_8_Lipid | 9.2E-06 | 2.1E-06 | 1.2E-05 | 2.0E-03 |
| Cells_Mean_LargeLipidObjects_Granularity_9_Lipid | 8.0E-06 | 2.1E-06 | 9.7E-05 | 7.7E-03 |
| Cells_RadialDistribution_FracAtD_AGP_1of4 | -1.9E-06 | 5.3E-07 | 3.7E-04 | 1.8E-02 |
| Cells_RadialDistribution_FracAtD_DNA_4of4 | 3.7E-06 | 7.8E-07 | 2.8E-06 | 7.7E-04 |
| Cells_RadialDistribution_FracAtD_Mito_4of4 | 2.1E-06 | 7.3E-07 | 4.5E-03 | 3.6E-02 |
| Cells_RadialDistribution_MeanFrac_DNA_4of4 | 3.8E-06 | 9.0E-07 | 2.5E-05 | 3.4E-03 |
| Cells_RadialDistribution_RadialCV_Lipid_1of4 | 2.0E-06 | 7.1E-07 | 4.0E-03 | 3.3E-02 |
| Cells_Texture_AngularSecondMoment_AGP_5_00 | 9.5E-06 | 2.5E-06 | 1.7E-04 | 1.1E-02 |
| Cells_Texture_Contrast_AGP_20_01 | -4.7E-06 | 1.4E-06 | 7.9E-04 | 1.9E-02 |
| Cells_Texture_DifferenceEntropy_AGP_20_01 | -6.3E-06 | 1.6E-06 | 8.2E-05 | 6.8E-03 |
| Cells_Texture_DifferenceEntropy_DNA_20_01 | -4.2E-06 | 1.4E-06 | 2.0E-03 | 2.4E-02 |
| Cells_Texture_DifferenceVariance_AGP_5_03 | 8.6E-06 | 1.9E-06 | 6.8E-06 | 1.4E-03 |
| Cells_Texture_Entropy_AGP_20_03 | -6.8E-06 | 1.7E-06 | 6.7E-05 | 5.9E-03 |
| Cells_Texture_InfoMeas2_AGP_10_03 | -2.7E-06 | 1.0E-06 | 7.6E-03 | 4.7E-02 |
| Cells_Texture_InfoMeas1_Lipid_10_01 | 5.2E-06 | 1.2E-06 | 1.1E-05 | 2.0E-03 |
| Cells_Texture_InfoMeas1_DNA_20_02 | 4.0E-06 | 6.7E-07 | 2.1E-09 | 2.8E-06 |
| Cells_Texture_InverseDifferenceMoment_AGP_5_03 | 8.4E-06 | 1.8E-06 | 3.8E-06 | 9.5E-04 |
| Cells_Texture_InverseDifferenceMoment_DNA_20_03 | 3.8E-06 | 1.4E-06 | 6.8E-03 | 4.4E-02 |
| Cells_Texture_SumAverage_AGP_20_01 | -7.4E-06 | 1.8E-06 | 4.1E-05 | 4.4E-03 |
| Cells_Texture_SumAverage_DNA_5_02 | -4.5E-06 | 1.4E-06 | 1.6E-03 | 2.3E-02 |
| Cells_Texture_SumEntropy_AGP_20_01 | -6.1E-06 | 1.7E-06 | 2.8E-04 | 1.5E-02 |
| Cells_Texture_SumEntropy_DNA_20_01 | -4.2E-06 | 1.5E-06 | 6.2E-03 | 4.1E-02 |
| Cells_Texture_SumVariance_AGP_20_02 | -3.9E-06 | 1.5E-06 | 8.0E-03 | 4.8E-02 |
| Cells_Texture_Variance_AGP_20_01 | -4.4E-06 | 1.5E-06 | 2.9E-03 | 2.8E-02 |
| LIPE (ENSG00000079435) |  |  |  |  |
| LipocyteProfiler features | beta | se | p-value | q-value |
| Cells_AreaShape_Zernike_7_5 | 1.9E-06 | 6.3E-07 | 2.2E-03 | 2.5E-02 |
| Cells_LipidObjects_AreaShape_Area | 2.1E-05 | 5.0E-06 | 3.1E-05 | 3.8E-03 |
| Cells_LipidObjects_Intensity_IntegratedIntensity_Lipid | 2.4E-05 | 6.5E-06 | 1.9E-04 | 1.2E-02 |
| Cells_Children_LipidObjects_Count | 1.2E-05 | 4.3E-06 | 4.5E-03 | 3.5E-02 |
| Cells_Children_LargeLipidObjects_Count | 2.2E-05 | 7.1E-06 | 2.3E-03 | 2.5E-02 |
| Cells_Correlation_Correlation_DNA_AGP | -8.9E-06 | 3.3E-06 | 6.0E-03 | 4.1E-02 |
| Cells_Correlation_Correlation_Mito_AGP | -1.5E-05 | 4.7E-06 | 1.1E-03 | 2.0E-02 |
| Cells_Correlation_Correlation_Mito_Lipid | -1.5E-05 | 5.3E-06 | 5.3E-03 | 3.9E-02 |
| Cells_Correlation_K_AGP_Mito | 2.1E-05 | 7.6E-06 | 5.4E-03 | 3.9E-02 |
| Cells_Correlation_K_DNA_Mito | 1.7E-05 | 4.9E-06 | 4.9E-04 | 1.9E-02 |
| Cells_Correlation_Overlap_Mito_AGP | -1.4E-05 | 3.6E-06 | 1.2E-04 | 9.1E-03 |
| Cells_Granularity_10_Mito | -8.4E-06 | 1.7E-06 | 7.2E-07 | 2.8E-04 |
| Cells_Granularity_11_AGP | -3.1E-06 | 9.6E-07 | 1.1E-03 | 2.0E-02 |
| Cells_Granularity_11_Mito | -9.2E-06 | 2.1E-06 | 1.6E-05 | 2.5E-03 |
| Cells_Granularity_12_Mito | -6.3E-06 | 2.2E-06 | 5.3E-03 | 3.9E-02 |
| Cells_Granularity_15_Lipid | 5.4E-06 | 1.6E-06 | 1.1E-03 | 2.0E-02 |
| Cells_Granularity_16_Lipid | 5.0E-06 | 1.7E-06 | 4.1E-03 | 3.4E-02 |
| Cells_Granularity_3_Lipid | 7.4E-06 | 2.6E-06 | 3.6E-03 | 3.2E-02 |
| Cells_Granularity_7_AGP | -7.9E-06 | 2.4E-06 | 8.5E-04 | 1.9E-02 |
| Cells_Granularity_8_AGP | -7.1E-06 | 2.4E-06 | 3.4E-03 | 3.1E-02 |
| Cells_Granularity_9_AGP | -6.5E-06 | 2.4E-06 | 7.7E-03 | 4.7E-02 |
| Cells_Granularity_9_Mito | -6.2E-06 | 2.1E-06 | 2.9E-03 | 2.8E-02 |
| Cells_Intensity_IntegratedIntensity_Lipid | 2.5E-05 | 6.4E-06 | 9.6E-05 | 7.7E-03 |
| Cells_Intensity_IntegratedIntensityEdge_Lipid | 3.0E-05 | 6.1E-06 | 1.1E-06 | 3.8E-04 |
| Cells_Intensity_IntegratedIntensityEdge_Mito | 2.3E-05 | 7.2E-06 | 1.7E-03 | 2.3E-02 |
| Cells_Intensity_LowerQuartileIntensity_Lipid | 3.7E-05 | 5.4E-06 | 5.0E-12 | 1.5E-08 |
| Cells_Intensity_LowerQuartileIntensity_Mito | 2.2E-05 | 7.1E-06 | 2.4E-03 | 2.6E-02 |
| Cells_Intensity_MADIntensity_Lipid | 3.0E-05 | 6.5E-06 | 5.5E-06 | 1.2E-03 |
| Cells_Intensity_MaxIntensityEdge_Mito | 1.9E-05 | 7.2E-06 | 6.9E-03 | 4.4E-02 |
| Cells_Intensity_MeanIntensity_Lipid | 2.3E-05 | 6.6E-06 | 4.1E-04 | 1.8E-02 |
| Cells_Intensity_MeanIntensity_Mito | 1.8E-05 | 6.8E-06 | 7.4E-03 | 4.6E-02 |
| Cells_Intensity_MeanIntensityEdge_Lipid | 2.8E-05 | 6.1E-06 | 6.2E-06 | 1.3E-03 |
| Cells_Intensity_MeanIntensityEdge_Mito | 2.7E-05 | 8.1E-06 | 8.4E-04 | 1.9E-02 |
| Cells_Intensity_MedianIntensity_Lipid | 3.4E-05 | 6.1E-06 | 1.9E-08 | 1.7E-05 |
| Cells_Intensity_MedianIntensity_Mito | 1.9E-05 | 7.0E-06 | 5.7E-03 | 4.0E-02 |
| Cells_Intensity_MinIntensity_AGP | -1.3E-05 | 4.6E-06 | 4.2E-03 | 3.4E-02 |
| Cells_Intensity_MinIntensity_Lipid | 2.5E-05 | 5.3E-06 | 2.9E-06 | 7.9E-04 |
| Cells_Intensity_MinIntensityEdge_AGP | -1.6E-05 | 5.4E-06 | 2.8E-03 | 2.8E-02 |
| Cells_Intensity_MinIntensityEdge_Lipid | 2.5E-05 | 5.3E-06 | 2.4E-06 | 6.8E-04 |
| Cells_Intensity_StdIntensityEdge_Mito | 2.1E-05 | 7.6E-06 | 4.9E-03 | 3.7E-02 |
| Cells_Intensity_UpperQuartileIntensity_Lipid | 2.6E-05 | 7.1E-06 | 2.5E-04 | 1.4E-02 |
| Cells_LargeLipidObjects_AreaShape_Area | 2.3E-05 | 5.9E-06 | 7.2E-05 | 6.2E-03 |
| Cells_LargeLipidObjects_Intensity_IntegratedIntensity_Lipid | 2.5E-05 | 7.0E-06 | 3.2E-04 | 1.6E-02 |
| Cells_Mean_LargeLipidObjects_Correlation_K_AGP_Mito | 2.2E-05 | 6.6E-06 | 9.8E-04 | 1.9E-02 |
| Cells_Mean_LargeLipidObjects_Correlation_K_DNA_Mito | 2.1E-05 | 7.2E-06 | 2.8E-03 | 2.8E-02 |
| Cells_Mean_LargeLipidObjects_Correlation_Overlap_DNA_AGP | -9.3E-06 | 3.2E-06 | 3.3E-03 | 3.1E-02 |
| Cells_Mean_LargeLipidObjects_Correlation_Overlap_Mito_AGP | -1.3E-05 | 2.4E-06 | 7.5E-08 | 5.0E-05 |
| Cells_Mean_LargeLipidObjects_Granularity_1_Lipid | 1.6E-05 | 4.4E-06 | 3.7E-04 | 1.8E-02 |
| Cells_Mean_LargeLipidObjects_Granularity_10_Lipid | 1.3E-05 | 4.7E-06 | 4.7E-03 | 3.6E-02 |
| Cells_Mean_LargeLipidObjects_Granularity_2_Lipid | 1.7E-05 | 4.4E-06 | 8.8E-05 | 7.2E-03 |
| Cells_Mean_LargeLipidObjects_Granularity_3_Lipid | 1.9E-05 | 4.5E-06 | 1.6E-05 | 2.5E-03 |
| Cells_Mean_LargeLipidObjects_Granularity_4_Lipid | 2.0E-05 | 4.6E-06 | 1.7E-05 | 2.6E-03 |
| Cells_Mean_LargeLipidObjects_Granularity_5_Lipid | 2.0E-05 | 5.2E-06 | 1.6E-04 | 1.0E-02 |
| Cells_Mean_LargeLipidObjects_Granularity_6_Lipid | 1.9E-05 | 5.6E-06 | 9.3E-04 | 1.9E-02 |
| Cells_Mean_LargeLipidObjects_Granularity_7_Lipid | 1.8E-05 | 5.7E-06 | 1.9E-03 | 2.4E-02 |
| Cells_Mean_LargeLipidObjects_Granularity_8_Lipid | 1.6E-05 | 5.5E-06 | 3.1E-03 | 3.0E-02 |
| Cells_Mean_LargeLipidObjects_Granularity_9_Lipid | 1.4E-05 | 5.3E-06 | 8.4E-03 | 4.9E-02 |
| Cells_RadialDistribution_FracAtD_DNA_4of4 | 7.5E-06 | 1.9E-06 | 8.8E-05 | 7.2E-03 |
| Cells_RadialDistribution_FracAtD_Mito_4of4 | 5.4E-06 | 1.6E-06 | 6.7E-04 | 1.9E-02 |
| Cells_RadialDistribution_MeanFrac_DNA_4of4 | 7.9E-06 | 2.2E-06 | 2.4E-04 | 1.4E-02 |
| Cells_RadialDistribution_RadialCV_AGP_1of4 | 5.1E-06 | 1.7E-06 | 2.1E-03 | 2.5E-02 |
| Cells_Texture_Contrast_Mito_5_00 | 2.6E-05 | 7.0E-06 | 2.6E-04 | 1.4E-02 |
| Cells_Texture_Correlation_AGP_10_02 | -5.4E-06 | 2.0E-06 | 6.1E-03 | 4.1E-02 |
| Cells_Texture_DifferenceEntropy_DNA_5_02 | -9.5E-06 | 3.3E-06 | 3.9E-03 | 3.3E-02 |
| Cells_Texture_Entropy_Lipid_5_02 | 1.7E-05 | 6.6E-06 | 8.1E-03 | 4.9E-02 |
| Cells_Texture_InfoMeas1_AGP_20_00 | 6.9E-06 | 2.3E-06 | 3.3E-03 | 3.1E-02 |
| Cells_Texture_InfoMeas1_Lipid_5_02 | 1.1E-05 | 3.9E-06 | 7.2E-03 | 4.5E-02 |
| Cells_Texture_InfoMeas1_DNA_10_01 | 8.9E-06 | 2.1E-06 | 1.7E-05 | 2.6E-03 |
| Cells_Texture_InfoMeas1_Mito_5_02 | 9.2E-06 | 3.2E-06 | 3.5E-03 | 3.2E-02 |
| Cells_Texture_SumAverage_Lipid_5_02 | 2.2E-05 | 6.5E-06 | 6.6E-04 | 1.9E-02 |
| Cells_Texture_SumAverage_DNA_5_02 | -1.0E-05 | 3.2E-06 | 1.7E-03 | 2.3E-02 |
| Cells_Texture_SumEntropy_DNA_20_01 | -9.6E-06 | 3.4E-06 | 5.1E-03 | 3.8E-02 |
| TIMM22 (ENSG00000177370) |  |  |  |  |
| LipocyteProfiler features | beta | se | p-value | q-value |
| Cells_Children_LipidObjects_Count | 1.5E-03 | 3.7E-04 | 4.6E-05 | 4.7E-03 |
| Cells_Correlation_K_AGP_Mito | 3.4E-03 | 6.1E-04 | 3.3E-08 | 2.7E-05 |
| Cells_Correlation_K_DNA_Mito | 2.3E-03 | 4.4E-04 | 1.3E-07 | 7.7E-05 |
| Cells_Correlation_Overlap_DNA_AGP | -7.5E-04 | 2.8E-04 | 7.8E-03 | 4.8E-02 |
| Cells_Correlation_Overlap_DNA_Lipid | -1.6E-03 | 4.8E-04 | 1.1E-03 | 2.0E-02 |
| Cells_Granularity_1_Mito | -1.4E-03 | 3.5E-04 | 8.1E-05 | 6.8E-03 |
| Cells_Granularity_3_Lipid | 7.0E-04 | 2.6E-04 | 6.3E-03 | 4.2E-02 |
| Cells_Granularity_4_Lipid | 1.0E-03 | 2.2E-04 | 4.1E-06 | 9.9E-04 |
| Cells_Granularity_5_Lipid | 1.0E-03 | 2.1E-04 | 2.6E-06 | 7.4E-04 |
| Cells_Intensity_IntegratedIntensity_Mito | 2.3E-03 | 5.9E-04 | 9.5E-05 | 7.6E-03 |
| Cells_Intensity_IntegratedIntensityEdge_DNA | 1.1E-03 | 3.6E-04 | 2.2E-03 | 2.5E-02 |
| Cells_Intensity_IntegratedIntensityEdge_Mito | 3.1E-03 | 7.3E-04 | 2.2E-05 | 3.1E-03 |
| Cells_Intensity_LowerQuartileIntensity_DNA | 1.0E-03 | 3.8E-04 | 6.3E-03 | 4.1E-02 |
| Cells_Intensity_LowerQuartileIntensity_Mito | 2.9E-03 | 7.8E-04 | 2.1E-04 | 1.2E-02 |
| Cells_Intensity_MADIntensity_Mito | 3.3E-03 | 5.2E-04 | 4.0E-10 | 6.8E-07 |
| Cells_Intensity_MassDisplacement_Lipid | -4.3E-04 | 1.6E-04 | 8.3E-03 | 4.9E-02 |
| Cells_Intensity_MassDisplacement_DNA | -3.6E-04 | 1.2E-04 | 1.5E-03 | 2.3E-02 |
| Cells_Intensity_MaxIntensity_Mito | 2.4E-03 | 4.3E-04 | 3.5E-08 | 2.8E-05 |
| Cells_Intensity_MaxIntensityEdge_Mito | 3.5E-03 | 5.5E-04 | 3.4E-10 | 5.9E-07 |
| Cells_Intensity_MeanIntensity_Mito | 3.1E-03 | 6.7E-04 | 4.4E-06 | 1.1E-03 |
| Cells_Intensity_MeanIntensityEdge_DNA | 1.1E-03 | 4.1E-04 | 6.4E-03 | 4.2E-02 |
| Cells_Intensity_MeanIntensityEdge_Mito | 3.5E-03 | 8.5E-04 | 4.6E-05 | 4.7E-03 |
| Cells_Intensity_MedianIntensity_Lipid | 2.6E-03 | 9.8E-04 | 7.2E-03 | 4.5E-02 |
| Cells_Intensity_MedianIntensity_Mito | 3.0E-03 | 7.0E-04 | 2.1E-05 | 3.0E-03 |
| Cells_Intensity_StdIntensity_Mito | 3.0E-03 | 5.1E-04 | 2.1E-09 | 2.8E-06 |
| Cells_Intensity_StdIntensityEdge_DNA | 1.0E-03 | 3.8E-04 | 7.6E-03 | 4.7E-02 |
| Cells_Intensity_StdIntensityEdge_Mito | 3.7E-03 | 5.6E-04 | 9.3E-11 | 1.9E-07 |
| Cells_Intensity_UpperQuartileIntensity_Mito | 3.1E-03 | 6.0E-04 | 3.2E-07 | 1.5E-04 |
| Cells_Mean_LargeLipidObjects_Correlation_K_AGP_Mito | 2.8E-03 | 5.7E-04 | 5.8E-07 | 2.4E-04 |
| Cells_Mean_LargeLipidObjects_Correlation_K_Lipid_Mito | 2.3E-03 | 5.6E-04 | 3.8E-05 | 4.2E-03 |
| Cells_Mean_LargeLipidObjects_Correlation_K_DNA_Mito | 2.1E-03 | 7.3E-04 | 4.5E-03 | 3.6E-02 |
| Cells_Mean_LargeLipidObjects_Correlation_K_Mito_AGP | -1.6E-03 | 6.2E-04 | 7.6E-03 | 4.7E-02 |
| Cells_Mean_LargeLipidObjects_Correlation_Overlap_Mito_AGP | -9.9E-04 | 3.5E-04 | 4.5E-03 | 3.5E-02 |
| Cells_Mean_LargeLipidObjects_Granularity_2_Lipid | 1.6E-03 | 5.8E-04 | 7.2E-03 | 4.5E-02 |
| Cells_Mean_LargeLipidObjects_Granularity_3_Lipid | 2.0E-03 | 5.7E-04 | 6.2E-04 | 1.9E-02 |
| Cells_Mean_LargeLipidObjects_Granularity_4_Lipid | 2.1E-03 | 5.5E-04 | 1.2E-04 | 8.9E-03 |
| Cells_Mean_LargeLipidObjects_Granularity_5_Lipid | 2.3E-03 | 5.2E-04 | 6.7E-06 | 1.4E-03 |
| Cells_Mean_LargeLipidObjects_Granularity_6_Lipid | 2.1E-03 | 4.9E-04 | 1.2E-05 | 2.0E-03 |
| Cells_Mean_LargeLipidObjects_Granularity_7_Lipid | 1.7E-03 | 5.5E-04 | 1.7E-03 | 2.3E-02 |
| Cells_RadialDistribution_FracAtD_DNA_4of4 | 5.9E-04 | 2.1E-04 | 5.2E-03 | 3.8E-02 |
| Cells_RadialDistribution_MeanFrac_DNA_4of4 | 6.8E-04 | 2.2E-04 | 2.5E-03 | 2.7E-02 |
| Cells_RadialDistribution_RadialCV_Lipid_1of4 | 8.0E-04 | 1.6E-04 | 3.1E-07 | 1.5E-04 |
| Cells_RadialDistribution_RadialCV_DNA_4of4 | -5.8E-04 | 2.2E-04 | 7.9E-03 | 4.8E-02 |
| Cells_RadialDistribution_RadialCV_Mito_1of4 | 6.0E-04 | 2.0E-04 | 2.9E-03 | 2.8E-02 |
| Cells_Texture_AngularSecondMoment_Mito_20_01 | -1.0E-03 | 3.3E-04 | 2.0E-03 | 2.5E-02 |
| Cells_Texture_Contrast_Mito_5_03 | 3.9E-03 | 5.5E-04 | 3.3E-12 | 1.0E-08 |
| Cells_Texture_Correlation_AGP_5_00 | -7.0E-04 | 2.5E-04 | 4.4E-03 | 3.5E-02 |
| Cells_Texture_DifferenceEntropy_Mito_5_02 | 2.6E-03 | 4.7E-04 | 1.9E-08 | 1.7E-05 |
| Cells_Texture_DifferenceVariance_Mito_10_03 | -1.5E-03 | 5.8E-04 | 8.2E-03 | 4.9E-02 |
| Cells_Texture_Entropy_Lipid_5_00 | 1.8E-03 | 6.5E-04 | 5.3E-03 | 3.9E-02 |
| Cells_Texture_Entropy_Mito_5_00 | 2.5E-03 | 4.6E-04 | 1.2E-07 | 7.1E-05 |
| Cells_Texture_InfoMeas1_Lipid_5_02 | 1.4E-03 | 4.1E-04 | 6.1E-04 | 1.9E-02 |
| Cells_Texture_InfoMeas1_Mito_5_00 | 8.6E-04 | 3.1E-04 | 6.2E-03 | 4.1E-02 |
| Cells_Texture_InverseDifferenceMoment_Lipid_5_02 | -1.8E-03 | 5.8E-04 | 1.8E-03 | 2.3E-02 |
| Cells_Texture_InverseDifferenceMoment_Mito_20_01 | -1.5E-03 | 4.1E-04 | 2.1E-04 | 1.3E-02 |
| Cells_Texture_SumAverage_Mito_5_02 | 3.1E-03 | 6.5E-04 | 2.6E-06 | 7.3E-04 |
| Cells_Texture_SumEntropy_Mito_5_02 | 2.4E-03 | 4.5E-04 | 8.9E-08 | 5.7E-05 |
| Cells_Texture_SumVariance_Mito_5_00 | 3.9E-03 | 6.5E-04 | 2.4E-09 | 3.2E-06 |
| Cells_Texture_Variance_Mito_5_00 | 3.9E-03 | 6.2E-04 | 3.5E-10 | 6.1E-07 |
| INSR (ENSG00000171105) |  |  |  |  |
| LipocyteProfiler features | beta | se | p-value | q-value |
| Cells_AreaShape_MaximumRadius | 1.2E-04 | 4.2E-05 | 3.5E-03 | 3.2E-02 |
| Cells_AreaShape_MeanRadius | 1.5E-04 | 4.1E-05 | 3.8E-04 | 1.8E-02 |
| Cells_AreaShape_MedianRadius | 1.7E-04 | 4.5E-05 | 2.2E-04 | 1.3E-02 |
| Cells_LipidObjects_AreaShape_Area | 4.7E-04 | 1.1E-04 | 8.1E-06 | 1.6E-03 |
| Cells_LipidObjects_Intensity_IntegratedIntensity_Lipid | 5.6E-04 | 1.4E-04 | 3.9E-05 | 4.2E-03 |
| Cells_Children_LipidObjects_Count | 2.8E-04 | 9.4E-05 | 2.7E-03 | 2.7E-02 |
| Cells_Children_LargeLipidObjects_Count | 4.6E-04 | 1.6E-04 | 4.0E-03 | 3.3E-02 |
| Cells_Correlation_Correlation_Mito_AGP | -3.0E-04 | 1.1E-04 | 4.9E-03 | 3.7E-02 |
| Cells_Correlation_Correlation_Mito_Lipid | -3.4E-04 | 1.1E-04 | 2.3E-03 | 2.6E-02 |
| Cells_Correlation_K_DNA_Mito | 3.5E-04 | 1.3E-04 | 5.2E-03 | 3.8E-02 |
| Cells_Correlation_K_Mito_DNA | -2.3E-04 | 8.6E-05 | 7.9E-03 | 4.8E-02 |
| Cells_Correlation_Overlap_Mito_AGP | -2.8E-04 | 8.7E-05 | 1.2E-03 | 2.2E-02 |
| Cells_Granularity_10_Mito | -1.9E-04 | 4.4E-05 | 1.2E-05 | 2.0E-03 |
| Cells_Granularity_11_AGP | -6.9E-05 | 2.1E-05 | 1.0E-03 | 1.9E-02 |
| Cells_Granularity_11_Mito | -2.5E-04 | 4.8E-05 | 1.7E-07 | 9.6E-05 |
| Cells_Granularity_12_Mito | -1.8E-04 | 4.9E-05 | 1.5E-04 | 1.0E-02 |
| Cells_Granularity_14_Lipid | 1.0E-04 | 3.6E-05 | 4.6E-03 | 3.6E-02 |
| Cells_Granularity_15_Lipid | 1.0E-04 | 3.8E-05 | 6.4E-03 | 4.2E-02 |
| Cells_Granularity_7_AGP | -1.7E-04 | 5.2E-05 | 1.5E-03 | 2.3E-02 |
| Cells_Granularity_8_AGP | -1.7E-04 | 5.2E-05 | 1.2E-03 | 2.2E-02 |
| Cells_Granularity_9_AGP | -1.6E-04 | 5.3E-05 | 2.6E-03 | 2.7E-02 |
| Cells_Intensity_IntegratedIntensity_Lipid | 5.7E-04 | 1.3E-04 | 1.6E-05 | 2.4E-03 |
| Cells_Intensity_IntegratedIntensityEdge_Lipid | 5.7E-04 | 1.5E-04 | 1.3E-04 | 9.1E-03 |
| Cells_Intensity_LowerQuartileIntensity_Lipid | 6.3E-04 | 1.7E-04 | 1.5E-04 | 9.9E-03 |
| Cells_Intensity_MADIntensity_Lipid | 6.4E-04 | 1.5E-04 | 1.3E-05 | 2.2E-03 |
| Cells_Intensity_MeanIntensity_Lipid | 4.8E-04 | 1.5E-04 | 1.3E-03 | 2.2E-02 |
| Cells_Intensity_MeanIntensityEdge_Lipid | 5.4E-04 | 1.5E-04 | 2.8E-04 | 1.5E-02 |
| Cells_Intensity_MedianIntensity_Lipid | 6.9E-04 | 1.5E-04 | 4.5E-06 | 1.1E-03 |
| Cells_Intensity_MinIntensity_AGP | -2.9E-04 | 1.0E-04 | 4.6E-03 | 3.6E-02 |
| Cells_Intensity_MinIntensity_Lipid | 4.8E-04 | 1.3E-04 | 2.7E-04 | 1.5E-02 |
| Cells_Intensity_MinIntensityEdge_AGP | -3.5E-04 | 1.2E-04 | 3.2E-03 | 3.0E-02 |
| Cells_Intensity_MinIntensityEdge_Lipid | 4.8E-04 | 1.3E-04 | 2.7E-04 | 1.5E-02 |
| Cells_Intensity_UpperQuartileIntensity_Lipid | 5.7E-04 | 1.5E-04 | 1.9E-04 | 1.2E-02 |
| Cells_LargeLipidObjects_AreaShape_Area | 5.4E-04 | 1.3E-04 | 2.3E-05 | 3.1E-03 |
| Cells_LargeLipidObjects_Intensity_IntegratedIntensity_Lipid | 6.0E-04 | 1.5E-04 | 4.3E-05 | 4.6E-03 |
| Cells_Mean_LargeLipidObjects_Correlation_K_DNA_Mito | 4.4E-04 | 1.7E-04 | 8.3E-03 | 4.9E-02 |
| Cells_Mean_LargeLipidObjects_Correlation_Overlap_DNA_AGP | -2.3E-04 | 6.8E-05 | 7.4E-04 | 1.9E-02 |
| Cells_Mean_LargeLipidObjects_Correlation_Overlap_Mito_AGP | -2.9E-04 | 5.4E-05 | 6.5E-08 | 4.5E-05 |
| Cells_Mean_LargeLipidObjects_Granularity_1_Lipid | 3.2E-04 | 1.1E-04 | 3.7E-03 | 3.3E-02 |
| Cells_Mean_LargeLipidObjects_Granularity_10_Lipid | 3.3E-04 | 9.4E-05 | 3.8E-04 | 1.8E-02 |
| Cells_Mean_LargeLipidObjects_Granularity_2_Lipid | 3.3E-04 | 1.1E-04 | 3.1E-03 | 2.9E-02 |
| Cells_Mean_LargeLipidObjects_Granularity_3_Lipid | 3.9E-04 | 1.1E-04 | 4.9E-04 | 1.9E-02 |
| Cells_Mean_LargeLipidObjects_Granularity_4_Lipid | 4.2E-04 | 1.1E-04 | 2.0E-04 | 1.2E-02 |
| Cells_Mean_LargeLipidObjects_Granularity_5_Lipid | 4.4E-04 | 1.2E-04 | 1.8E-04 | 1.1E-02 |
| Cells_Mean_LargeLipidObjects_Granularity_6_Lipid | 4.6E-04 | 1.1E-04 | 5.7E-05 | 5.4E-03 |
| Cells_Mean_LargeLipidObjects_Granularity_7_Lipid | 4.6E-04 | 1.1E-04 | 3.0E-05 | 3.7E-03 |
| Cells_Mean_LargeLipidObjects_Granularity_8_Lipid | 4.3E-04 | 1.1E-04 | 4.8E-05 | 4.8E-03 |
| Cells_Mean_LargeLipidObjects_Granularity_9_Lipid | 3.8E-04 | 1.0E-04 | 2.4E-04 | 1.4E-02 |
| Cells_RadialDistribution_FracAtD_DNA_4of4 | 1.8E-04 | 4.2E-05 | 2.4E-05 | 3.3E-03 |
| Cells_RadialDistribution_FracAtD_Mito_4of4 | 1.2E-04 | 3.8E-05 | 2.5E-03 | 2.6E-02 |
| Cells_RadialDistribution_MeanFrac_DNA_4of4 | 1.9E-04 | 4.6E-05 | 2.9E-05 | 3.7E-03 |
| Cells_RadialDistribution_RadialCV_Lipid_1of4 | 1.3E-04 | 3.9E-05 | 7.3E-04 | 1.9E-02 |
| Cells_Texture_Contrast_DNA_20_01 | -2.5E-04 | 9.1E-05 | 5.9E-03 | 4.0E-02 |
| Cells_Texture_Correlation_Lipid_20_03 | 1.1E-04 | 3.6E-05 | 1.5E-03 | 2.3E-02 |
| Cells_Texture_DifferenceEntropy_DNA_5_02 | -2.3E-04 | 6.8E-05 | 8.2E-04 | 1.9E-02 |
| Cells_Texture_Entropy_Lipid_20_03 | 3.9E-04 | 1.3E-04 | 3.4E-03 | 3.1E-02 |
| Cells_Texture_Entropy_DNA_20_03 | -2.1E-04 | 7.5E-05 | 5.5E-03 | 3.9E-02 |
| Cells_Texture_InfoMeas1_AGP_20_01 | 1.5E-04 | 5.0E-05 | 2.5E-03 | 2.6E-02 |
| Cells_Texture_InfoMeas1_Lipid_10_01 | 3.1E-04 | 7.4E-05 | 3.7E-05 | 4.1E-03 |
| Cells_Texture_InfoMeas1_DNA_20_02 | 2.6E-04 | 4.3E-05 | 1.0E-09 | 1.6E-06 |
| Cells_Texture_InverseDifferenceMoment_DNA_20_01 | 1.9E-04 | 6.7E-05 | 3.8E-03 | 3.3E-02 |
| Cells_Texture_SumAverage_Lipid_20_01 | 4.4E-04 | 1.5E-04 | 2.6E-03 | 2.7E-02 |
| Cells_Texture_SumAverage_DNA_5_02 | -2.9E-04 | 7.0E-05 | 3.5E-05 | 3.9E-03 |
| Cells_Texture_SumEntropy_DNA_20_01 | -2.3E-04 | 7.1E-05 | 1.4E-03 | 2.2E-02 |
| Cells_Texture_SumVariance_DNA_5_02 | -2.9E-04 | 9.2E-05 | 1.9E-03 | 2.4E-02 |
| Cells_Texture_Variance_DNA_5_01 | -2.8E-04 | 9.2E-05 | 2.2E-03 | 2.5E-02 |
| GLUT4 (ENSG00000181856) |  |  |  |  |
| LipocyteProfiler features | beta | se | p-value | q-value |
| Cells_AreaShape_Zernike_6_4 | 8.1E-06 | 1.7E-06 | 1.8E-06 | 5.5E-04 |
| Cells_LipidObjects_AreaShape_Area | 5.9E-05 | 1.8E-05 | 1.1E-03 | 2.0E-02 |
| Cells_LipidObjects_Intensity_IntegratedIntensity_Lipid | 6.2E-05 | 2.2E-05 | 6.1E-03 | 4.1E-02 |
| Cells_Correlation_Correlation_DNA_AGP | -2.7E-05 | 9.7E-06 | 4.9E-03 | 3.7E-02 |
| Cells_Correlation_Overlap_Mito_AGP | -3.7E-05 | 1.2E-05 | 2.5E-03 | 2.6E-02 |
| Cells_Granularity_10_Mito | -2.3E-05 | 6.3E-06 | 2.9E-04 | 1.5E-02 |
| Cells_Granularity_11_AGP | -1.2E-05 | 3.0E-06 | 9.4E-05 | 7.5E-03 |
| Cells_Granularity_11_Mito | -3.0E-05 | 6.3E-06 | 2.3E-06 | 6.7E-04 |
| Cells_Granularity_12_Mito | -2.6E-05 | 5.8E-06 | 6.1E-06 | 1.3E-03 |
| Cells_Granularity_7_AGP | -2.5E-05 | 8.3E-06 | 2.4E-03 | 2.6E-02 |
| Cells_Granularity_8_AGP | -2.4E-05 | 8.4E-06 | 4.8E-03 | 3.6E-02 |
| Cells_Intensity_IntegratedIntensity_Lipid | 6.9E-05 | 2.5E-05 | 5.4E-03 | 3.9E-02 |
| Cells_Intensity_LowerQuartileIntensity_Lipid | 8.2E-05 | 2.9E-05 | 4.4E-03 | 3.5E-02 |
| Cells_Intensity_MADIntensity_AGP | -4.2E-05 | 1.3E-05 | 1.2E-03 | 2.1E-02 |
| Cells_Intensity_MADIntensity_Lipid | 7.5E-05 | 2.7E-05 | 4.7E-03 | 3.6E-02 |
| Cells_Intensity_MaxIntensity_AGP | -3.1E-05 | 1.0E-05 | 3.2E-03 | 3.0E-02 |
| Cells_Intensity_MeanIntensity_AGP | -4.4E-05 | 1.5E-05 | 4.3E-03 | 3.5E-02 |
| Cells_Intensity_MedianIntensity_AGP | -4.3E-05 | 1.6E-05 | 6.0E-03 | 4.1E-02 |
| Cells_Intensity_MedianIntensity_Lipid | 8.4E-05 | 2.8E-05 | 2.4E-03 | 2.6E-02 |
| Cells_Intensity_MinIntensity_AGP | -4.8E-05 | 1.4E-05 | 5.5E-04 | 1.9E-02 |
| Cells_Intensity_MinIntensity_Lipid | 6.7E-05 | 2.2E-05 | 2.4E-03 | 2.6E-02 |
| Cells_Intensity_MinIntensityEdge_AGP | -6.2E-05 | 1.6E-05 | 8.7E-05 | 7.2E-03 |
| Cells_Intensity_MinIntensityEdge_Lipid | 6.7E-05 | 2.2E-05 | 2.4E-03 | 2.6E-02 |
| Cells_Intensity_StdIntensity_AGP | -4.0E-05 | 1.2E-05 | 8.7E-04 | 1.9E-02 |
| Cells_Intensity_StdIntensityEdge_AGP | -3.2E-05 | 1.2E-05 | 6.4E-03 | 4.2E-02 |
| Cells_Intensity_UpperQuartileIntensity_AGP | -4.6E-05 | 1.5E-05 | 1.6E-03 | 2.3E-02 |
| Cells_LargeLipidObjects_AreaShape_Area | 6.4E-05 | 2.0E-05 | 1.7E-03 | 2.3E-02 |
| Cells_LargeLipidObjects_Intensity_IntegratedIntensity_Lipid | 6.2E-05 | 2.3E-05 | 6.9E-03 | 4.4E-02 |
| Cells_Mean_LargeLipidObjects_AreaShape_Orientation | 1.1E-06 | 4.0E-07 | 7.1E-03 | 4.5E-02 |
| Cells_Mean_LargeLipidObjects_Correlation_K_AGP_Mito | 6.0E-05 | 2.3E-05 | 7.6E-03 | 4.7E-02 |
| Cells_Mean_LargeLipidObjects_Correlation_K_Mito_AGP | -4.9E-05 | 1.7E-05 | 4.3E-03 | 3.5E-02 |
| Cells_Mean_LargeLipidObjects_Correlation_Overlap_DNA_AGP | -3.2E-05 | 9.7E-06 | 1.1E-03 | 2.0E-02 |
| Cells_Mean_LargeLipidObjects_Correlation_Overlap_Mito_AGP | -3.5E-05 | 9.0E-06 | 7.9E-05 | 6.7E-03 |
| Cells_Mean_LargeLipidObjects_Granularity_1_Lipid | 4.1E-05 | 1.5E-05 | 5.3E-03 | 3.9E-02 |
| Cells_Mean_LargeLipidObjects_Granularity_10_Lipid | 4.9E-05 | 1.5E-05 | 1.5E-03 | 2.2E-02 |
| Cells_Mean_LargeLipidObjects_Granularity_3_Lipid | 5.3E-05 | 1.5E-05 | 4.4E-04 | 1.8E-02 |
| Cells_Mean_LargeLipidObjects_Granularity_4_Lipid | 5.9E-05 | 1.4E-05 | 2.6E-05 | 3.5E-03 |
| Cells_Mean_LargeLipidObjects_Granularity_5_Lipid | 6.6E-05 | 1.5E-05 | 6.0E-06 | 1.3E-03 |
| Cells_Mean_LargeLipidObjects_Granularity_6_Lipid | 6.8E-05 | 1.6E-05 | 1.2E-05 | 2.0E-03 |
| Cells_Mean_LargeLipidObjects_Granularity_7_Lipid | 6.7E-05 | 1.7E-05 | 9.9E-05 | 7.8E-03 |
| Cells_Mean_LargeLipidObjects_Granularity_8_Lipid | 6.3E-05 | 1.7E-05 | 3.2E-04 | 1.6E-02 |
| Cells_Mean_LargeLipidObjects_Granularity_9_Lipid | 5.5E-05 | 1.7E-05 | 1.2E-03 | 2.1E-02 |
| Cells_RadialDistribution_FracAtD_AGP_4of4 | 1.5E-05 | 5.0E-06 | 2.0E-03 | 2.5E-02 |
| Cells_RadialDistribution_FracAtD_DNA_4of4 | 2.6E-05 | 6.0E-06 | 1.7E-05 | 2.5E-03 |
| Cells_RadialDistribution_FracAtD_Mito_4of4 | 1.8E-05 | 5.3E-06 | 9.6E-04 | 1.9E-02 |
| Cells_RadialDistribution_MeanFrac_AGP_4of4 | 2.6E-05 | 9.5E-06 | 6.1E-03 | 4.1E-02 |
| Cells_RadialDistribution_MeanFrac_DNA_4of4 | 2.6E-05 | 6.9E-06 | 1.6E-04 | 1.0E-02 |
| Cells_RadialDistribution_RadialCV_AGP_1of4 | 1.6E-05 | 5.3E-06 | 3.2E-03 | 3.0E-02 |
| Cells_RadialDistribution_RadialCV_Lipid_1of4 | 1.6E-05 | 5.2E-06 | 3.1E-03 | 3.0E-02 |
| Cells_Texture_AngularSecondMoment_AGP_5_00 | 7.4E-05 | 1.9E-05 | 1.3E-04 | 9.5E-03 |
| Cells_Texture_Contrast_AGP_20_01 | -3.1E-05 | 1.2E-05 | 7.5E-03 | 4.6E-02 |
| Cells_Texture_Correlation_Lipid_5_00 | 2.7E-05 | 9.1E-06 | 3.4E-03 | 3.1E-02 |
| Cells_Texture_DifferenceEntropy_AGP_20_01 | -4.6E-05 | 1.3E-05 | 3.2E-04 | 1.6E-02 |
| Cells_Texture_DifferenceVariance_AGP_20_00 | 6.6E-05 | 1.5E-05 | 6.1E-06 | 1.3E-03 |
| Cells_Texture_Entropy_AGP_20_03 | -4.9E-05 | 1.4E-05 | 3.8E-04 | 1.8E-02 |
| Cells_Texture_InfoMeas1_AGP_20_01 | 2.1E-05 | 7.2E-06 | 3.5E-03 | 3.2E-02 |
| Cells_Texture_InfoMeas1_Lipid_10_01 | 3.5E-05 | 9.9E-06 | 4.0E-04 | 1.8E-02 |
| Cells_Texture_InfoMeas1_DNA_20_02 | 2.6E-05 | 6.6E-06 | 8.4E-05 | 7.0E-03 |
| Cells_Texture_InverseDifferenceMoment_AGP_10_00 | 6.3E-05 | 1.4E-05 | 6.6E-06 | 1.4E-03 |
| Cells_Texture_InverseDifferenceMoment_DNA_20_01 | 2.9E-05 | 1.1E-05 | 8.1E-03 | 4.9E-02 |
| Cells_Texture_SumAverage_AGP_20_03 | -4.5E-05 | 1.5E-05 | 3.0E-03 | 2.9E-02 |
| Cells_Texture_SumEntropy_AGP_20_01 | -4.4E-05 | 1.3E-05 | 1.1E-03 | 2.0E-02 |
| Cells_Texture_SumVariance_AGP_5_03 | -3.0E-05 | 1.1E-05 | 7.5E-03 | 4.6E-02 |
| Cells_Texture_Variance_AGP_20_02 | -3.1E-05 | 1.2E-05 | 7.9E-03 | 4.8E-02 |

**Supplemental Table 3:** Significant effects of drug perturbation on LP-profiles in AMSCs and PHH (t-test, significance level AMSCs 5%FDR, PHH 0.1% FDR). p-value of t-test, q-value of t-test, t-statistics of t-test.

| mature visceral AMSCs isoproterenol treatment |  |  |  |
| --- | --- | --- | --- |
| LipocyteProfiler features | p-value | q-value | t-statistics |
| Cells_AreaShape_Eccentricity | 1.32E-02 | 3.34E-02 | -5.97E-01 |
| Cells_AreaShape_MaximumRadius | 2.74E-02 | 4.80E-02 | 1.08E+00 |
| Cells_AreaShape_Zernike_2_0 | 1.58E-02 | 3.72E-02 | 1.02E+00 |
| Cells_Correlation_Correlation_DNA_Lipid | 1.72E-02 | 3.90E-02 | 5.55E-01 |
| Cells_Correlation_K_Lipid_AGP | 2.83E-05 | 1.59E-03 | 1.07E+00 |
| Cells_Correlation_K_Lipid_DNA | 1.08E-03 | 1.59E-02 | 7.03E-01 |
| Cells_Granularity_11_Lipid | 1.21E-02 | 3.24E-02 | -8.78E-01 |
| Cells_Granularity_9_AGP | 2.53E-02 | 4.64E-02 | -9.61E-01 |
| Cells_Granularity_6_Lipid | 2.43E-02 | 4.57E-02 | -1.10E+00 |
| Cells_Granularity_8_Mito | 2.29E-02 | 4.48E-02 | -1.10E+00 |
| Cells_Intensity_MassDisplacement_DNA | 1.18E-02 | 3.22E-02 | 1.38E+00 |
| Cells_Intensity_MassDisplacement_Mito | 5.40E-03 | 2.51E-02 | 5.48E-01 |
| Cells_Intensity_MaxIntensity_Lipid | 2.44E-03 | 1.93E-02 | -4.80E-01 |
| Cells_Intensity_MeanIntensityEdge_DNA | 1.88E-02 | 4.04E-02 | -6.04E-01 |
| Cells_Intensity_StdIntensityEdge_DNA | 1.18E-02 | 3.22E-02 | -6.43E-01 |
| Cells_Mean_LargeLipidObjects_AreaShape_Eccentricity | 5.06E-03 | 2.47E-02 | 8.53E-01 |
| Cells_Mean_LargeLipidObjects_AreaShape_FormFactor | 2.01E-02 | 4.17E-02 | -5.46E-01 |
| Cells_Mean_LargeLipidObjects_AreaShape_Solidity | 1.65E-02 | 3.81E-02 | -5.12E-01 |
| Cells_Mean_LargeLipidObjects_Correlation_K_Mito_Lipid | 1.06E-02 | 3.10E-02 | -4.76E-01 |
| Cells_Mean_LargeLipidObjects_Correlation_Overlap_DNA_AGP | 1.21E-02 | 3.24E-02 | 1.10E+00 |
| Cells_Mean_LargeLipidObjects_Correlation_Overlap_Mito_AGP | 9.74E-03 | 3.03E-02 | 1.62E+00 |
| Cells_RadialDistribution_FracAtD_DNA_3of4 | 2.61E-02 | 4.70E-02 | 5.31E-01 |
| Cells_RadialDistribution_MeanFrac_Lipid_1of4 | 1.28E-02 | 3.28E-02 | -1.27E+00 |
| Cells_RadialDistribution_RadialCV_AGP_4of4 | 1.22E-02 | 3.24E-02 | -1.12E+00 |
| Cells_Texture_Correlation_AGP_10_00 | 1.00E-02 | 3.07E-02 | 7.05E-01 |
| Cells_Texture_Correlation_DNA_20_02 | 1.24E-02 | 3.24E-02 | 1.05E+00 |
| Cells_Texture_Correlation_Mito_20_00 | 2.48E-02 | 4.62E-02 | 7.50E-01 |
| Cells_Texture_DifferenceEntropy_AGP_10_01 | 2.69E-02 | 4.75E-02 | -6.33E-01 |
| Cells_Texture_DifferenceEntropy_Lipid_10_03 | 3.20E-03 | 1.97E-02 | -7.73E-01 |
| Cells_Texture_DifferenceVariance_Lipid_5_02 | 8.78E-05 | 2.84E-03 | 1.05E+00 |
| Cells_Texture_Entropy_Lipid_10_00 | 8.36E-03 | 2.85E-02 | -7.72E-01 |
| Cells_Texture_InfoMeas2_Lipid_20_01 | 1.12E-02 | 3.17E-02 | -6.49E-01 |
| Cells_Texture_InfoMeas1_DNA_10_01 | 2.60E-02 | 4.70E-02 | -9.90E-01 |
| Cells_Texture_InfoMeas1_Mito_10_01 | 5.92E-03 | 2.53E-02 | -1.08E+00 |
| Cells_Texture_InverseDifferenceMoment_Lipid_5_00 | 1.76E-02 | 3.95E-02 | 8.65E-01 |
| Cells_Texture_SumAverage_DNA_20_02 | 1.99E-02 | 4.17E-02 | -3.25E-01 |
| Cells_Texture_SumEntropy_Lipid_10_00 | 5.69E-03 | 2.51E-02 | -7.10E-01 |
| Cytoplasm_AreaShape_Compactness | 6.83E-04 | 1.18E-02 | -9.66E-01 |
| Cytoplasm_AreaShape_Eccentricity | 2.55E-02 | 4.64E-02 | -5.27E-01 |
| Cytoplasm_AreaShape_MaximumRadius | 2.69E-02 | 4.75E-02 | 1.09E+00 |
| Cytoplasm_AreaShape_MeanRadius | 1.83E-02 | 4.00E-02 | 1.24E+00 |
| Cytoplasm_AreaShape_Zernike_5_1 | 7.44E-03 | 2.69E-02 | 1.39E+00 |
| Cytoplasm_Correlation_K_Lipid_AGP | 1.06E-02 | 3.10E-02 | 1.17E+00 |
| Cytoplasm_Granularity_11_Lipid | 9.25E-03 | 2.96E-02 | -9.35E-01 |
| Cytoplasm_Granularity_9_AGP | 2.15E-02 | 4.34E-02 | -9.88E-01 |
| Cytoplasm_Granularity_6_Lipid | 2.13E-02 | 4.32E-02 | -1.13E+00 |
| Cytoplasm_Granularity_8_Mito | 2.50E-02 | 4.62E-02 | -1.21E+00 |
| Cytoplasm_Intensity_MassDisplacement_Mito | 2.37E-03 | 1.93E-02 | 5.68E-01 |
| Cytoplasm_Intensity_MaxIntensity_Lipid | 1.74E-03 | 1.78E-02 | -4.96E-01 |
| Cytoplasm_RadialDistribution_FracAtD_AGP_1of4 | 6.85E-03 | 2.68E-02 | 1.38E+00 |
| Cytoplasm_RadialDistribution_FracAtD_DNA_2of4 | 6.82E-04 | 1.18E-02 | 9.33E-01 |
| Cytoplasm_RadialDistribution_FracAtD_Mito_1of4 | 2.62E-02 | 4.70E-02 | 1.29E+00 |
| Cytoplasm_RadialDistribution_MeanFrac_AGP_2of4 | 1.69E-02 | 3.86E-02 | 1.13E+00 |
| Cytoplasm_Texture_Correlation_AGP_5_03 | 2.90E-02 | 4.98E-02 | 1.12E+00 |
| Cytoplasm_Texture_Correlation_DNA_20_03 | 9.27E-03 | 2.96E-02 | 6.50E-01 |
| Cytoplasm_Texture_Correlation_Mito_20_01 | 2.35E-03 | 1.93E-02 | 9.14E-01 |
| Cytoplasm_Texture_DifferenceEntropy_Lipid_20_00 | 3.07E-03 | 1.97E-02 | -7.83E-01 |
| Cytoplasm_Texture_DifferenceVariance_Lipid_10_03 | 1.16E-05 | 1.50E-03 | 1.28E+00 |
| Cytoplasm_Texture_Entropy_Lipid_10_00 | 2.89E-02 | 4.98E-02 | -6.22E-01 |
| Cytoplasm_Texture_InfoMeas2_Lipid_20_01 | 3.63E-04 | 6.95E-03 | -6.58E-01 |
| Cytoplasm_Texture_InfoMeas1_DNA_5_00 | 3.93E-03 | 2.21E-02 | -6.45E-01 |
| Cytoplasm_Texture_InfoMeas1_Mito_10_01 | 1.84E-03 | 1.78E-02 | -1.49E+00 |
| Cytoplasm_Texture_SumEntropy_Lipid_20_00 | 1.00E-02 | 3.07E-02 | -6.53E-01 |
| Nuclei_AreaShape_Eccentricity | 1.83E-03 | 1.78E-02 | -1.09E+00 |
| Nuclei_AreaShape_Zernike_3_3 | 2.90E-02 | 4.98E-02 | -1.30E+00 |
| Nuclei_Correlation_Correlation_Lipid_AGP | 2.60E-02 | 4.70E-02 | 1.06E+00 |
| Nuclei_Correlation_Correlation_DNA_Lipid | 1.73E-03 | 1.78E-02 | 6.58E-01 |
| Nuclei_Correlation_K_Lipid_AGP | 1.52E-02 | 3.65E-02 | 2.22E-01 |
| Nuclei_Correlation_Overlap_Lipid_AGP | 5.33E-03 | 2.51E-02 | 5.25E-01 |
| Nuclei_Correlation_Overlap_DNA_AGP | 2.66E-03 | 1.97E-02 | 1.29E+00 |
| Nuclei_Correlation_Overlap_DNA_Lipid | 7.37E-03 | 2.69E-02 | 5.31E-01 |
| Nuclei_Correlation_Overlap_Mito_Lipid | 7.75E-03 | 2.73E-02 | 6.24E-01 |
| Nuclei_Granularity_11_Lipid | 1.35E-02 | 3.38E-02 | -7.55E-01 |
| Nuclei_Granularity_11_Mito | 1.59E-02 | 3.72E-02 | 3.32E-01 |
| Nuclei_Granularity_9_AGP | 1.63E-02 | 3.80E-02 | -1.26E+00 |
| Nuclei_Granularity_1_Lipid | 1.55E-03 | 1.78E-02 | 1.03E+00 |
| Nuclei_Granularity_4_DNA | 9.51E-03 | 3.02E-02 | -1.10E+00 |
| Nuclei_Intensity_MassDisplacement_AGP | 2.14E-03 | 1.85E-02 | -1.05E+00 |
| Nuclei_Intensity_MinIntensity_DNA | 4.56E-03 | 2.36E-02 | 3.13E-01 |
| Nuclei_Intensity_MinIntensityEdge_DNA | 4.47E-03 | 2.36E-02 | 3.13E-01 |
| Nuclei_RadialDistribution_FracAtD_Lipid_1of4 | 1.57E-02 | 3.71E-02 | 1.03E+00 |
| Nuclei_RadialDistribution_MeanFrac_Lipid_1of4 | 2.78E-02 | 4.84E-02 | 7.83E-01 |
| Nuclei_RadialDistribution_MeanFrac_DNA_3of4 | 1.35E-03 | 1.78E-02 | -1.25E+00 |
| Nuclei_RadialDistribution_RadialCV_AGP_3of4 | 5.66E-03 | 2.51E-02 | -1.33E+00 |
| Nuclei_RadialDistribution_RadialCV_DNA_3of4 | 1.82E-02 | 4.00E-02 | -1.21E+00 |
| Nuclei_Texture_Correlation_Lipid_5_01 | 5.27E-03 | 2.51E-02 | -1.65E+00 |
| Nuclei_Texture_Correlation_DNA_5_02 | 1.18E-02 | 3.22E-02 | -9.49E-01 |
| Nuclei_Texture_Correlation_Mito_5_02 | 6.62E-03 | 2.63E-02 | -8.42E-01 |
| Nuclei_Texture_DifferenceEntropy_Lipid_20_03 | 5.37E-03 | 2.51E-02 | -8.16E-01 |
| Nuclei_Texture_DifferenceVariance_Lipid_20_03 | 2.12E-02 | 4.32E-02 | 4.70E-01 |
| Nuclei_Texture_Entropy_Lipid_20_01 | 5.51E-03 | 2.51E-02 | -7.64E-01 |
| Nuclei_Texture_InfoMeas2_AGP_5_02 | 2.31E-03 | 1.93E-02 | -1.20E+00 |
| Nuclei_Texture_InfoMeas1_Lipid_5_01 | 1.01E-02 | 3.08E-02 | 1.08E+00 |
| Nuclei_Texture_InfoMeas2_DNA_5_02 | 4.02E-03 | 2.21E-02 | -8.35E-01 |
| Nuclei_Texture_InfoMeas1_Mito_5_02 | 1.75E-03 | 1.78E-02 | 1.22E+00 |
| Nuclei_Texture_InverseDifferenceMoment_AGP_20_01 | 1.07E-02 | 3.11E-02 | 1.01E+00 |
| Nuclei_Texture_InverseDifferenceMoment_Lipid_20_00 | 7.56E-04 | 1.26E-02 | 7.43E-01 |
| Nuclei_Texture_SumAverage_DNA_20_03 | 2.78E-03 | 1.97E-02 | 4.54E-01 |
| Nuclei_Texture_SumEntropy_Lipid_20_01 | 6.33E-03 | 2.61E-02 | -7.81E-01 |
| PHH olic acid treatment |  |  |  |
| LipocyteProfiler features | pvalue | q-value | t-statistics |
| Cells_Children_LargeLipidObjects_Count | 2.3E-11 | 3.4E-10 | 1.9E+01 |
| Cells_Correlation_Correlation_Lipid_AGP | 9.5E-13 | 7.7E-11 | -2.4E+01 |
| Cells_Correlation_Correlation_DNA_Lipid | 2.1E-12 | 1.2E-10 | -2.3E+01 |
| Cells_Correlation_Correlation_Mito_Lipid | 5.1E-06 | 1.6E-05 | -7.1E+00 |
| Cells_Correlation_K_AGP_Lipid | 2.6E-07 | 1.0E-06 | 9.2E+00 |
| Cells_Correlation_K_Lipid_AGP | 2.0E-11 | 3.1E-10 | -1.9E+01 |
| Cells_Correlation_K_Lipid_DNA | 2.2E-11 | 3.3E-10 | -1.9E+01 |
| Cells_Correlation_K_Lipid_Mito | 1.2E-08 | 5.5E-08 | -1.2E+01 |
| Cells_Correlation_K_DNA_Lipid | 4.1E-07 | 1.5E-06 | 8.9E+00 |
| Cells_Correlation_K_DNA_Mito | 1.2E-04 | 2.9E-04 | 5.2E+00 |
| Cells_Correlation_K_Mito_Lipid | 8.1E-09 | 3.7E-08 | 1.2E+01 |
| Cells_Correlation_Overlap_Lipid_AGP | 1.3E-11 | 2.3E-10 | -2.0E+01 |
| Cells_Correlation_Overlap_DNA_Lipid | 9.0E-12 | 2.1E-10 | -2.0E+01 |
| Cells_Correlation_Overlap_Mito_Lipid | 2.1E-08 | 9.0E-08 | -1.1E+01 |
| Cells_Granularity_1_AGP | 5.7E-06 | 1.8E-05 | -7.1E+00 |
| Cells_Granularity_2_Lipid | 7.1E-13 | 6.8E-11 | 2.4E+01 |
| Cells_Intensity_IntegratedIntensity_Lipid | 7.1E-10 | 4.0E-09 | 1.5E+01 |
| Cells_Intensity_IntegratedIntensityEdge_Lipid | 1.1E-11 | 2.1E-10 | 2.0E+01 |
| Cells_Intensity_IntegratedIntensityEdge_DNA | 7.9E-06 | 2.4E-05 | -6.9E+00 |
| Cells_Intensity_LowerQuartileIntensity_Lipid | 3.2E-04 | 7.1E-04 | 4.7E+00 |
| Cells_Intensity_LowerQuartileIntensity_DNA | 3.0E-07 | 1.2E-06 | -9.1E+00 |
| Cells_Intensity_MADIntensity_Lipid | 2.8E-09 | 1.4E-08 | 1.3E+01 |
| Cells_Intensity_MADIntensity_DNA | 1.3E-05 | 3.9E-05 | -6.5E+00 |
| Cells_Intensity_MassDisplacement_Lipid | 1.8E-08 | 7.8E-08 | 1.1E+01 |
| Cells_Intensity_MassDisplacement_DNA | 5.3E-05 | 1.3E-04 | 5.7E+00 |
| Cells_Intensity_MaxIntensity_Lipid | 5.1E-10 | 3.0E-09 | 1.5E+01 |
| Cells_Intensity_MaxIntensityEdge_Lipid | 3.3E-11 | 4.6E-10 | 1.8E+01 |
| Cells_Intensity_MeanIntensity_Lipid | 1.2E-09 | 6.6E-09 | 1.4E+01 |
| Cells_Intensity_MeanIntensity_DNA | 1.2E-05 | 3.5E-05 | -6.6E+00 |
| Cells_Intensity_MeanIntensityEdge_Lipid | 1.6E-11 | 2.6E-10 | 1.9E+01 |
| Cells_Intensity_MeanIntensityEdge_DNA | 9.6E-07 | 3.3E-06 | -8.2E+00 |
| Cells_Intensity_MedianIntensity_Lipid | 4.8E-06 | 1.5E-05 | 7.2E+00 |
| Cells_Intensity_MedianIntensity_DNA | 5.9E-07 | 2.1E-06 | -8.6E+00 |
| Cells_Intensity_MinIntensity_DNA | 8.7E-09 | 3.9E-08 | -1.2E+01 |
| Cells_Intensity_MinIntensityEdge_DNA | 2.8E-08 | 1.2E-07 | -1.1E+01 |
| Cells_Intensity_StdIntensity_Lipid | 1.3E-11 | 2.3E-10 | 2.0E+01 |
| Cells_Intensity_StdIntensityEdge_Lipid | 1.7E-12 | 1.1E-10 | 2.3E+01 |
| Cells_Intensity_UpperQuartileIntensity_Lipid | 6.3E-09 | 2.9E-08 | 1.2E+01 |
| Cells_Intensity_UpperQuartileIntensity_DNA | 8.9E-06 | 2.7E-05 | -6.8E+00 |
| Cells_Mean_LargeLipidObjects_AreaShape_Area | 1.1E-11 | 2.1E-10 | 2.0E+01 |
| Cells_Mean_LargeLipidObjects_AreaShape_Center_X | 1.9E-12 | 1.1E-10 | 2.3E+01 |
| Cells_Mean_LargeLipidObjects_AreaShape_Center_Y | 1.1E-11 | 2.1E-10 | 2.0E+01 |
| Cells_Mean_LargeLipidObjects_AreaShape_Center_Z | 6.1E-12 | 1.7E-10 | 2.1E+01 |
| Cells_Mean_LargeLipidObjects_AreaShape_Compactness | 1.0E-11 | 2.1E-10 | 2.0E+01 |
| Cells_Mean_LargeLipidObjects_AreaShape_Eccentricity | 5.1E-12 | 1.7E-10 | 2.1E+01 |
| Cells_Mean_LargeLipidObjects_AreaShape_EulerNumber | 6.1E-12 | 1.7E-10 | 2.1E+01 |
| Cells_Mean_LargeLipidObjects_AreaShape_Extent | 2.5E-12 | 1.3E-10 | 2.2E+01 |
| Cells_Mean_LargeLipidObjects_AreaShape_FormFactor | 7.3E-13 | 6.8E-11 | 2.4E+01 |
| Cells_Mean_LargeLipidObjects_AreaShape_MajorAxisLength | 9.2E-12 | 2.1E-10 | 2.0E+01 |
| Cells_Mean_LargeLipidObjects_AreaShape_MaxFeretDiameter | 9.9E-12 | 2.1E-10 | 2.0E+01 |
| Cells_Mean_LargeLipidObjects_AreaShape_MaximumRadius | 3.0E-12 | 1.4E-10 | 2.2E+01 |
| Cells_Mean_LargeLipidObjects_AreaShape_MeanRadius | 2.8E-12 | 1.3E-10 | 2.2E+01 |
| Cells_Mean_LargeLipidObjects_AreaShape_MedianRadius | 2.8E-12 | 1.3E-10 | 2.2E+01 |
| Cells_Mean_LargeLipidObjects_AreaShape_MinFeretDiameter | 1.4E-11 | 2.4E-10 | 2.0E+01 |
| Cells_Mean_LargeLipidObjects_AreaShape_MinorAxisLength | 1.2E-11 | 2.1E-10 | 2.0E+01 |
| Cells_Mean_LargeLipidObjects_AreaShape_Perimeter | 2.9E-11 | 4.2E-10 | 1.9E+01 |
| Cells_Mean_LargeLipidObjects_AreaShape_Solidity | 3.2E-12 | 1.4E-10 | 2.2E+01 |
| Cells_Mean_LargeLipidObjects_Correlation_Correlation_Lipid_AGP | 5.5E-12 | 1.7E-10 | -2.1E+01 |
| Cells_Mean_LargeLipidObjects_Correlation_Correlation_DNA_AGP | 1.5E-10 | 1.0E-09 | 1.6E+01 |
| Cells_Mean_LargeLipidObjects_Correlation_Correlation_DNA_Lipid | 1.7E-15 | 9.5E-13 | -3.8E+01 |
| Cells_Mean_LargeLipidObjects_Correlation_Correlation_DNA_Mito | 4.2E-12 | 1.6E-10 | 2.1E+01 |
| Cells_Mean_LargeLipidObjects_Correlation_Correlation_Mito_AGP | 2.8E-10 | 1.8E-09 | 1.6E+01 |
| Cells_Mean_LargeLipidObjects_Correlation_Correlation_Mito_Lipid | 1.7E-15 | 9.5E-13 | -3.8E+01 |
| Cells_Mean_LargeLipidObjects_Correlation_K_AGP_Lipid | 3.0E-10 | 1.9E-09 | 1.6E+01 |
| Cells_Mean_LargeLipidObjects_Correlation_K_AGP_DNA | 2.8E-07 | 1.1E-06 | 9.1E+00 |
| Cells_Mean_LargeLipidObjects_Correlation_K_AGP_Mito | 2.2E-08 | 9.5E-08 | 1.1E+01 |
| Cells_Mean_LargeLipidObjects_Correlation_K_DNA_AGP | 3.1E-09 | 1.5E-08 | 1.3E+01 |
| Cells_Mean_LargeLipidObjects_Correlation_K_DNA_Lipid | 7.4E-12 | 1.9E-10 | 2.1E+01 |
| Cells_Mean_LargeLipidObjects_Correlation_K_DNA_Mito | 4.8E-09 | 2.3E-08 | 1.3E+01 |
| Cells_Mean_LargeLipidObjects_Correlation_K_Mito_AGP | 6.6E-08 | 2.7E-07 | 1.0E+01 |
| Cells_Mean_LargeLipidObjects_Correlation_K_Mito_Lipid | 6.0E-13 | 6.8E-11 | 2.5E+01 |
| Cells_Mean_LargeLipidObjects_Correlation_K_Mito_DNA | 1.9E-07 | 7.6E-07 | 9.5E+00 |
| Cells_Mean_LargeLipidObjects_Correlation_Overlap_Lipid_AGP | 5.2E-12 | 1.7E-10 | 2.1E+01 |
| Cells_Mean_LargeLipidObjects_Correlation_Overlap_DNA_AGP | 5.4E-12 | 1.7E-10 | 2.1E+01 |
| Cells_Mean_LargeLipidObjects_Correlation_Overlap_DNA_Lipid | 8.7E-12 | 2.1E-10 | 2.0E+01 |
| Cells_Mean_LargeLipidObjects_Correlation_Overlap_DNA_Mito | 5.9E-12 | 1.7E-10 | 2.1E+01 |
| Cells_Mean_LargeLipidObjects_Correlation_Overlap_Mito_AGP | 5.7E-12 | 1.7E-10 | 2.1E+01 |
| Cells_Mean_LargeLipidObjects_Correlation_Overlap_Mito_Lipid | 8.4E-12 | 2.1E-10 | 2.0E+01 |
| Cells_Mean_LargeLipidObjects_Granularity_13_Lipid | 7.3E-07 | 2.6E-06 | 8.4E+00 |
| Cells_Mean_LargeLipidObjects_Granularity_2_Lipid | 1.7E-14 | 5.9E-12 | 3.2E+01 |
| Cells_Mean_LargeLipidObjects_Intensity_IntegratedIntensity_Lipid | 3.9E-12 | 1.5E-10 | 2.2E+01 |
| Cells_Mean_LargeLipidObjects_Intensity_IntegratedIntensityEdge_Lipid | 1.1E-11 | 2.1E-10 | 2.0E+01 |
| Cells_Mean_LargeLipidObjects_Intensity_LowerQuartileIntensity_Lipid | 7.9E-12 | 2.0E-10 | 2.0E+01 |
| Cells_Mean_LargeLipidObjects_Intensity_MADIntensity_Lipid | 6.7E-13 | 6.8E-11 | 2.5E+01 |
| Cells_Mean_LargeLipidObjects_Intensity_MassDisplacement_Lipid | 1.5E-10 | 1.0E-09 | 1.6E+01 |
| Cells_Mean_LargeLipidObjects_Intensity_MaxIntensity_Lipid | 4.8E-12 | 1.7E-10 | 2.1E+01 |
| Cells_Mean_LargeLipidObjects_Intensity_MaxIntensityEdge_Lipid | 5.6E-12 | 1.7E-10 | 2.1E+01 |
| Cells_Mean_LargeLipidObjects_Intensity_MeanIntensity_Lipid | 3.9E-12 | 1.5E-10 | 2.2E+01 |
| Cells_Mean_LargeLipidObjects_Intensity_MeanIntensityEdge_Lipid | 7.3E-12 | 1.9E-10 | 2.1E+01 |
| Cells_Mean_LargeLipidObjects_Intensity_MedianIntensity_Lipid | 5.0E-12 | 1.7E-10 | 2.1E+01 |
| Cells_Mean_LargeLipidObjects_Intensity_MinIntensity_Lipid | 1.5E-11 | 2.5E-10 | 1.9E+01 |
| Cells_Mean_LargeLipidObjects_Intensity_MinIntensityEdge_Lipid | 1.5E-11 | 2.5E-10 | 1.9E+01 |
| Cells_Mean_LargeLipidObjects_Intensity_StdIntensity_Lipid | 2.7E-12 | 1.3E-10 | 2.2E+01 |
| Cells_Mean_LargeLipidObjects_Intensity_StdIntensityEdge_Lipid | 3.8E-12 | 1.5E-10 | 2.2E+01 |
| Cells_Mean_LargeLipidObjects_Intensity_UpperQuartileIntensity_Lipid | 3.0E-12 | 1.4E-10 | 2.2E+01 |
| Cells_Mean_LargeLipidObjects_Location_Center_X | 1.9E-12 | 1.1E-10 | 2.3E+01 |
| Cells_Mean_LargeLipidObjects_Location_Center_Y | 1.1E-11 | 2.1E-10 | 2.0E+01 |
| Cells_Mean_LargeLipidObjects_Location_CenterMassIntensity_X_Lipid | 1.9E-12 | 1.1E-10 | 2.3E+01 |
| Cells_Mean_LargeLipidObjects_Location_CenterMassIntensity_Y_Lipid | 1.1E-11 | 2.1E-10 | 2.0E+01 |
| Cells_Mean_LargeLipidObjects_Location_MaxIntensity_X_Lipid | 1.9E-12 | 1.1E-10 | 2.3E+01 |
| Cells_Mean_LargeLipidObjects_Location_MaxIntensity_Y_Lipid | 1.1E-11 | 2.1E-10 | 2.0E+01 |
| Cells_Mean_LargeLipidObjects_Number_Object_Number | 5.4E-11 | 6.6E-10 | 1.8E+01 |
| Cells_Neighbors_FirstClosestObjectNumber_10 | 1.7E-04 | 3.8E-04 | -5.1E+00 |
| Cells_Neighbors_FirstClosestObjectNumber_Adjacent | 1.7E-04 | 3.8E-04 | -5.1E+00 |
| Cells_Neighbors_NumberOfNeighbors_10 | 4.4E-04 | 9.5E-04 | -4.6E+00 |
| Cells_Neighbors_SecondClosestDistance_10 | 3.9E-04 | 8.4E-04 | 4.6E+00 |
| Cells_Neighbors_SecondClosestDistance_Adjacent | 3.9E-04 | 8.4E-04 | 4.6E+00 |
| Cells_Neighbors_SecondClosestObjectNumber_10 | 1.5E-04 | 3.4E-04 | -5.1E+00 |
| Cells_Neighbors_SecondClosestObjectNumber_Adjacent | 1.5E-04 | 3.4E-04 | -5.1E+00 |
| Cells_Number_Object_Number | 1.6E-04 | 3.7E-04 | -5.1E+00 |
| Cells_Parent_Nuclei | 1.7E-04 | 3.8E-04 | -5.1E+00 |
| Cells_RadialDistribution_FracAtD_Lipid_2of4 | 1.0E-11 | 2.1E-10 | -2.0E+01 |
| Cells_RadialDistribution_MeanFrac_Lipid_2of4 | 4.1E-11 | 5.5E-10 | -1.8E+01 |
| Cells_RadialDistribution_RadialCV_Lipid_3of4 | 1.6E-10 | 1.1E-09 | 1.6E+01 |
| Cells_RadialDistribution_RadialCV_DNA_4of4 | 3.6E-05 | 9.2E-05 | 5.9E+00 |
| Cells_Texture_AngularSecondMoment_Lipid_20_01 | 4.9E-07 | 1.8E-06 | -8.7E+00 |
| Cells_Texture_AngularSecondMoment_DNA_10_03 | 3.6E-05 | 9.1E-05 | 5.9E+00 |
| Cells_Texture_Contrast_Lipid_20_01 | 6.9E-11 | 7.5E-10 | 1.7E+01 |
| Cells_Texture_Correlation_Lipid_10_02 | 1.9E-08 | 8.1E-08 | -1.1E+01 |
| Cells_Texture_Correlation_DNA_20_03 | 8.5E-06 | 2.6E-05 | 6.8E+00 |
| Cells_Texture_Correlation_Mito_20_00 | 1.1E-04 | 2.6E-04 | 5.3E+00 |
| Cells_Texture_DifferenceEntropy_Lipid_10_00 | 4.9E-11 | 6.2E-10 | 1.8E+01 |
| Cells_Texture_DifferenceEntropy_DNA_10_02 | 3.6E-06 | 1.2E-05 | -7.3E+00 |
| Cells_Texture_DifferenceVariance_Lipid_5_02 | 8.6E-07 | 3.0E-06 | -8.3E+00 |
| Cells_Texture_Entropy_Lipid_10_02 | 2.5E-09 | 1.2E-08 | 1.3E+01 |
| Cells_Texture_Entropy_DNA_10_02 | 1.8E-06 | 6.1E-06 | -7.8E+00 |
| Cells_Texture_InfoMeas1_Lipid_5_00 | 3.9E-12 | 1.5E-10 | -2.2E+01 |
| Cells_Texture_InfoMeas2_DNA_20_00 | 3.0E-05 | 7.8E-05 | -6.1E+00 |
| Cells_Texture_InverseDifferenceMoment_Lipid_20_01 | 6.1E-09 | 2.8E-08 | -1.2E+01 |
| Cells_Texture_InverseDifferenceMoment_DNA_20_02 | 2.5E-07 | 1.0E-06 | 9.2E+00 |
| Cells_Texture_SumAverage_Lipid_20_01 | 8.8E-10 | 4.8E-09 | 1.4E+01 |
| Cells_Texture_SumAverage_DNA_20_02 | 9.7E-06 | 2.9E-05 | -6.7E+00 |
| Cells_Texture_SumEntropy_Lipid_20_01 | 1.5E-10 | 1.1E-09 | 1.6E+01 |
| Cells_Texture_SumEntropy_DNA_10_02 | 5.2E-06 | 1.6E-05 | -7.1E+00 |
| Cells_Texture_SumVariance_Lipid_10_00 | 1.0E-10 | 8.9E-10 | 1.7E+01 |
| Cells_Texture_Variance_Lipid_20_01 | 8.7E-11 | 8.1E-10 | 1.7E+01 |
| Cytoplasm_AreaShape_Zernike_7_1 | 4.2E-05 | 1.1E-04 | -5.8E+00 |
| Cytoplasm_Correlation_Correlation_Lipid_AGP | 2.3E-15 | 9.8E-13 | -3.7E+01 |
| Cytoplasm_Correlation_Correlation_DNA_Lipid | 9.5E-14 | 2.3E-11 | -2.8E+01 |
| Cytoplasm_Correlation_Correlation_Mito_Lipid | 1.4E-12 | 9.9E-11 | -2.3E+01 |
| Cytoplasm_Correlation_K_AGP_Lipid | 2.8E-09 | 1.4E-08 | 1.3E+01 |
| Cytoplasm_Correlation_K_Lipid_AGP | 3.8E-10 | 2.3E-09 | -1.5E+01 |
| Cytoplasm_Correlation_K_Lipid_DNA | 1.5E-10 | 1.0E-09 | -1.6E+01 |
| Cytoplasm_Correlation_K_Lipid_Mito | 2.2E-08 | 9.5E-08 | -1.1E+01 |
| Cytoplasm_Correlation_K_DNA_Lipid | 5.6E-13 | 6.8E-11 | 2.5E+01 |
| Cytoplasm_Correlation_K_DNA_Mito | 1.9E-05 | 5.3E-05 | 6.3E+00 |
| Cytoplasm_Correlation_K_Mito_Lipid | 5.1E-09 | 2.4E-08 | 1.3E+01 |
| Cytoplasm_Correlation_K_Mito_DNA | 1.3E-05 | 3.7E-05 | -6.6E+00 |
| Cytoplasm_Correlation_Overlap_Lipid_AGP | 7.2E-07 | 2.6E-06 | -8.5E+00 |
| Cytoplasm_Correlation_Overlap_DNA_Lipid | 5.7E-06 | 1.8E-05 | -7.1E+00 |
| Cytoplasm_Correlation_Overlap_Mito_Lipid | 2.6E-07 | 1.0E-06 | -9.2E+00 |
| Cytoplasm_Granularity_1_AGP | 1.1E-05 | 3.2E-05 | -6.7E+00 |
| Cytoplasm_Granularity_2_Lipid | 1.4E-15 | 9.5E-13 | 3.8E+01 |
| Cytoplasm_Intensity_IntegratedIntensity_Lipid | 3.3E-10 | 2.0E-09 | 1.6E+01 |
| Cytoplasm_Intensity_IntegratedIntensity_DNA | 2.6E-04 | 5.8E-04 | -4.8E+00 |
| Cytoplasm_Intensity_IntegratedIntensityEdge_Lipid | 3.1E-11 | 4.4E-10 | 1.8E+01 |
| Cytoplasm_Intensity_IntegratedIntensityEdge_DNA | 2.8E-07 | 1.1E-06 | -9.2E+00 |
| Cytoplasm_Intensity_LowerQuartileIntensity_DNA | 2.7E-07 | 1.1E-06 | -9.2E+00 |
| Cytoplasm_Intensity_MADIntensity_Lipid | 4.0E-10 | 2.4E-09 | 1.5E+01 |
| Cytoplasm_Intensity_MADIntensity_DNA | 4.7E-05 | 1.2E-04 | -5.8E+00 |
| Cytoplasm_Intensity_MassDisplacement_Lipid | 2.7E-09 | 1.3E-08 | 1.3E+01 |
| Cytoplasm_Intensity_MaxIntensity_Lipid | 5.0E-10 | 2.9E-09 | 1.5E+01 |
| Cytoplasm_Intensity_MaxIntensity_DNA | 7.5E-05 | 1.8E-04 | -5.5E+00 |
| Cytoplasm_Intensity_MaxIntensityEdge_Lipid | 4.8E-11 | 6.2E-10 | 1.8E+01 |
| Cytoplasm_Intensity_MaxIntensityEdge_DNA | 4.3E-05 | 1.1E-04 | -5.8E+00 |
| Cytoplasm_Intensity_MeanIntensity_Lipid | 3.6E-10 | 2.2E-09 | 1.5E+01 |
| Cytoplasm_Intensity_MeanIntensity_DNA | 2.7E-07 | 1.0E-06 | -9.2E+00 |
| Cytoplasm_Intensity_MeanIntensityEdge_Lipid | 5.4E-11 | 6.6E-10 | 1.8E+01 |
| Cytoplasm_Intensity_MeanIntensityEdge_DNA | 1.4E-07 | 5.6E-07 | -9.7E+00 |
| Cytoplasm_Intensity_MedianIntensity_Lipid | 3.7E-06 | 1.2E-05 | 7.3E+00 |
| Cytoplasm_Intensity_MedianIntensity_DNA | 5.9E-07 | 2.1E-06 | -8.6E+00 |
| Cytoplasm_Intensity_MinIntensity_DNA | 8.7E-09 | 3.9E-08 | -1.2E+01 |
| Cytoplasm_Intensity_MinIntensityEdge_DNA | 2.8E-08 | 1.2E-07 | -1.1E+01 |
| Cytoplasm_Intensity_StdIntensity_Lipid | 9.3E-12 | 2.1E-10 | 2.0E+01 |
| Cytoplasm_Intensity_StdIntensity_DNA | 8.0E-06 | 2.4E-05 | -6.8E+00 |
| Cytoplasm_Intensity_StdIntensityEdge_Lipid | 2.4E-12 | 1.3E-10 | 2.2E+01 |
| Cytoplasm_Intensity_StdIntensityEdge_DNA | 4.8E-05 | 1.2E-04 | -5.8E+00 |
| Cytoplasm_Intensity_UpperQuartileIntensity_Lipid | 5.2E-10 | 3.0E-09 | 1.5E+01 |
| Cytoplasm_Intensity_UpperQuartileIntensity_DNA | 8.3E-07 | 2.9E-06 | -8.3E+00 |
| Cytoplasm_Number_Object_Number | 1.6E-04 | 3.7E-04 | -5.1E+00 |
| Cytoplasm_Parent_Cells | 1.7E-04 | 3.8E-04 | -5.1E+00 |
| Cytoplasm_Parent_Nuclei | 1.7E-04 | 3.8E-04 | -5.1E+00 |
| Cytoplasm_RadialDistribution_FracAtD_AGP_1of4 | 1.7E-05 | 4.8E-05 | 6.4E+00 |
| Cytoplasm_RadialDistribution_MeanFrac_AGP_1of4 | 1.9E-05 | 5.3E-05 | 6.3E+00 |
| Cytoplasm_RadialDistribution_MeanFrac_Lipid_3of4 | 1.9E-07 | 7.7E-07 | -9.4E+00 |
| Cytoplasm_RadialDistribution_RadialCV_Lipid_2of4 | 8.4E-11 | 8.0E-10 | 1.7E+01 |
| Cytoplasm_Texture_AngularSecondMoment_Lipid_20_01 | 2.2E-07 | 9.0E-07 | -9.3E+00 |
| Cytoplasm_Texture_AngularSecondMoment_DNA_20_02 | 1.2E-04 | 2.8E-04 | 5.3E+00 |
| Cytoplasm_Texture_Contrast_Lipid_10_03 | 5.7E-11 | 6.7E-10 | 1.8E+01 |
| Cytoplasm_Texture_Contrast_DNA_5_02 | 1.3E-06 | 4.4E-06 | -8.0E+00 |
| Cytoplasm_Texture_Correlation_Lipid_5_02 | 7.5E-13 | 6.8E-11 | 2.4E+01 |
| Cytoplasm_Texture_DifferenceEntropy_Lipid_10_01 | 9.7E-12 | 2.1E-10 | 2.0E+01 |
| Cytoplasm_Texture_DifferenceEntropy_DNA_10_02 | 1.7E-06 | 5.7E-06 | -7.8E+00 |
| Cytoplasm_Texture_DifferenceVariance_Lipid_20_00 | 3.0E-07 | 1.1E-06 | -9.1E+00 |
| Cytoplasm_Texture_DifferenceVariance_DNA_20_02 | 1.8E-05 | 5.0E-05 | 6.4E+00 |
| Cytoplasm_Texture_Entropy_Lipid_10_02 | 5.5E-10 | 3.2E-09 | 1.5E+01 |
| Cytoplasm_Texture_Entropy_DNA_10_03 | 1.5E-05 | 4.3E-05 | -6.5E+00 |
| Cytoplasm_Texture_InfoMeas1_Lipid_5_00 | 2.8E-13 | 5.4E-11 | -2.6E+01 |
| Cytoplasm_Texture_InfoMeas2_DNA_20_00 | 6.1E-05 | 1.5E-04 | -5.6E+00 |
| Cytoplasm_Texture_InverseDifferenceMoment_Lipid_20_02 | 1.5E-09 | 7.9E-09 | -1.4E+01 |
| Cytoplasm_Texture_InverseDifferenceMoment_DNA_20_02 | 5.7E-06 | 1.8E-05 | 7.1E+00 |
| Cytoplasm_Texture_SumAverage_Lipid_10_02 | 2.7E-10 | 1.8E-09 | 1.6E+01 |
| Cytoplasm_Texture_SumAverage_DNA_10_01 | 2.5E-07 | 9.8E-07 | -9.2E+00 |
| Cytoplasm_Texture_SumEntropy_Lipid_10_01 | 4.1E-11 | 5.5E-10 | 1.8E+01 |
| Cytoplasm_Texture_SumEntropy_DNA_10_03 | 1.8E-05 | 5.1E-05 | -6.3E+00 |
| Cytoplasm_Texture_SumVariance_Lipid_10_00 | 6.8E-11 | 7.5E-10 | 1.7E+01 |
| Cytoplasm_Texture_SumVariance_DNA_10_00 | 2.7E-05 | 7.2E-05 | -6.1E+00 |
| Cytoplasm_Texture_Variance_Lipid_10_03 | 7.4E-11 | 7.7E-10 | 1.7E+01 |
| Cytoplasm_Texture_Variance_DNA_5_00 | 1.4E-05 | 4.1E-05 | -6.5E+00 |
| Nuclei_AreaShape_Zernike_3_3 | 4.4E-05 | 1.1E-04 | 5.8E+00 |
| Nuclei_Correlation_Correlation_Lipid_AGP | 1.0E-11 | 2.1E-10 | -2.0E+01 |
| Nuclei_Correlation_Correlation_DNA_Lipid | 3.4E-11 | 4.7E-10 | -1.8E+01 |
| Nuclei_Correlation_Correlation_Mito_Lipid | 1.3E-06 | 4.3E-06 | 8.0E+00 |
| Nuclei_Correlation_K_Lipid_AGP | 4.6E-05 | 1.1E-04 | -5.8E+00 |
| Nuclei_Correlation_K_Lipid_DNA | 1.7E-09 | 9.2E-09 | -1.4E+01 |
| Nuclei_Correlation_K_DNA_Lipid | 7.0E-08 | 2.9E-07 | 1.0E+01 |
| Nuclei_Correlation_K_DNA_Mito | 1.0E-05 | 3.0E-05 | 6.7E+00 |
| Nuclei_Correlation_K_Mito_DNA | 1.1E-05 | 3.3E-05 | -6.6E+00 |
| Nuclei_Correlation_Overlap_Lipid_AGP | 7.8E-12 | 2.0E-10 | -2.0E+01 |
| Nuclei_Correlation_Overlap_DNA_Lipid | 5.5E-12 | 1.7E-10 | -2.1E+01 |
| Nuclei_Granularity_10_Lipid | 8.3E-05 | 2.0E-04 | 5.5E+00 |
| Nuclei_Granularity_1_Lipid | 2.9E-06 | 9.3E-06 | -7.5E+00 |
| Nuclei_Intensity_IntegratedIntensityEdge_Lipid | 9.3E-07 | 3.2E-06 | 8.3E+00 |
| Nuclei_Intensity_MassDisplacement_Lipid | 5.2E-14 | 1.5E-11 | 3.0E+01 |
| Nuclei_Intensity_MassDisplacement_DNA | 3.6E-05 | 9.3E-05 | 5.9E+00 |
| Nuclei_Intensity_MaxIntensity_Lipid | 7.1E-09 | 3.2E-08 | 1.2E+01 |
| Nuclei_Intensity_MaxIntensityEdge_Lipid | 3.7E-10 | 2.2E-09 | 1.5E+01 |
| Nuclei_Intensity_MeanIntensityEdge_Lipid | 1.3E-06 | 4.4E-06 | 8.0E+00 |
| Nuclei_Intensity_MinIntensity_DNA | 1.7E-05 | 4.9E-05 | -6.4E+00 |
| Nuclei_Intensity_MinIntensityEdge_DNA | 1.7E-05 | 4.9E-05 | -6.4E+00 |
| Nuclei_Intensity_StdIntensity_Lipid | 1.6E-07 | 6.5E-07 | 9.6E+00 |
| Nuclei_Intensity_StdIntensityEdge_Lipid | 2.2E-11 | 3.3E-10 | 1.9E+01 |
| Nuclei_Neighbors_FirstClosestObjectNumber_2 | 1.5E-04 | 3.6E-04 | -5.1E+00 |
| Nuclei_Neighbors_SecondClosestObjectNumber_2 | 1.8E-04 | 4.1E-04 | -5.0E+00 |
| Nuclei_Number_Object_Number | 1.6E-04 | 3.7E-04 | -5.1E+00 |
| Nuclei_RadialDistribution_FracAtD_Lipid_2of4 | 7.2E-10 | 4.0E-09 | -1.5E+01 |
| Nuclei_RadialDistribution_MeanFrac_Lipid_2of4 | 2.0E-09 | 1.0E-08 | -1.3E+01 |
| Nuclei_RadialDistribution_RadialCV_Lipid_4of4 | 1.8E-13 | 3.9E-11 | 2.7E+01 |
| Nuclei_RadialDistribution_RadialCV_DNA_3of4 | 8.3E-07 | 2.9E-06 | 8.3E+00 |
| Nuclei_Texture_Contrast_Lipid_10_01 | 9.1E-10 | 5.0E-09 | 1.4E+01 |
| Nuclei_Texture_Correlation_Lipid_5_03 | 3.7E-12 | 1.5E-10 | 2.2E+01 |
| Nuclei_Texture_InfoMeas1_Lipid_10_03 | 1.2E-04 | 2.7E-04 | -5.3E+00 |
| Nuclei_Texture_SumAverage_Lipid_10_01 | 3.9E-04 | 8.4E-04 | 4.6E+00 |
| Nuclei_Texture_SumEntropy_Lipid_10_01 | 1.7E-05 | 4.9E-05 | 6.4E+00 |
| Nuclei_Texture_SumVariance_Lipid_10_01 | 2.5E-10 | 1.6E-09 | 1.6E+01 |
| Nuclei_Texture_Variance_Lipid_10_01 | 4.2E-10 | 2.5E-09 | 1.5E+01 |
| PHH metformin treatment |  |  |  |
| LipocyteProfiler features | p-value | q-value | t-statistics |
| Cells_AreaShape_Area | 3.9E-11 | 2.0E-11 | -1.8E+01 |
| Cells_AreaShape_Center_Y | 1.1E-03 | 1.1E-04 | 4.1E+00 |
| Cells_AreaShape_Compactness | 3.7E-14 | 5.2E-14 | -3.0E+01 |
| Cells_AreaShape_Eccentricity | 2.8E-09 | 9.0E-10 | -1.3E+01 |
| Cells_AreaShape_EulerNumber | 7.9E-11 | 3.8E-11 | 1.7E+01 |
| Cells_AreaShape_Extent | 7.7E-16 | 2.5E-15 | 4.0E+01 |
| Cells_AreaShape_FormFactor | 1.5E-16 | 7.6E-16 | 4.5E+01 |
| Cells_AreaShape_MajorAxisLength | 3.1E-12 | 2.1E-12 | -2.2E+01 |
| Cells_AreaShape_MaxFeretDiameter | 1.0E-12 | 8.3E-13 | -2.4E+01 |
| Cells_AreaShape_MaximumRadius | 2.8E-06 | 4.6E-07 | -7.5E+00 |
| Cells_AreaShape_MinFeretDiameter | 1.3E-12 | 1.0E-12 | -2.3E+01 |
| Cells_AreaShape_MinorAxisLength | 1.2E-11 | 7.2E-12 | -2.0E+01 |
| Cells_AreaShape_Perimeter | 1.8E-15 | 4.8E-15 | -3.8E+01 |
| Cells_AreaShape_Solidity | 2.0E-17 | 1.9E-16 | 5.2E+01 |
| Cells_AreaShape_Zernike_0_0 | 3.2E-16 | 1.2E-15 | 4.3E+01 |
| Cells_Children_LargeLipidObjects_Count | 2.4E-07 | 4.8E-08 | -9.3E+00 |
| Cells_Correlation_Correlation_Lipid_AGP | 1.4E-07 | 2.9E-08 | 9.7E+00 |
| Cells_Correlation_Correlation_DNA_AGP | 1.0E-18 | 2.8E-17 | -6.4E+01 |
| Cells_Correlation_Correlation_DNA_Lipid | 2.9E-06 | 4.6E-07 | 7.5E+00 |
| Cells_Correlation_Correlation_DNA_Mito | 1.5E-14 | 2.5E-14 | 3.2E+01 |
| Cells_Correlation_Correlation_Mito_AGP | 3.3E-18 | 5.6E-17 | 5.9E+01 |
| Cells_Correlation_Correlation_Mito_Lipid | 1.1E-13 | 1.2E-13 | 2.8E+01 |
| Cells_Correlation_K_AGP_Lipid | 3.0E-03 | 2.7E-04 | 3.6E+00 |
| Cells_Correlation_K_AGP_DNA | 9.2E-05 | 1.1E-05 | 5.4E+00 |
| Cells_Correlation_K_AGP_Mito | 1.3E-06 | 2.2E-07 | 8.0E+00 |
| Cells_Correlation_K_Lipid_AGP | 1.1E-05 | 1.6E-06 | -6.6E+00 |
| Cells_Correlation_K_Lipid_DNA | 2.8E-06 | 4.5E-07 | -7.5E+00 |
| Cells_Correlation_K_Lipid_Mito | 8.5E-08 | 1.9E-08 | 1.0E+01 |
| Cells_Correlation_K_DNA_AGP | 2.0E-06 | 3.2E-07 | -7.8E+00 |
| Cells_Correlation_K_DNA_Lipid | 8.8E-06 | 1.2E-06 | 6.8E+00 |
| Cells_Correlation_K_DNA_Mito | 1.8E-07 | 3.8E-08 | 9.5E+00 |
| Cells_Correlation_K_Mito_AGP | 1.3E-06 | 2.3E-07 | -8.0E+00 |
| Cells_Correlation_K_Mito_Lipid | 2.2E-07 | 4.6E-08 | -9.3E+00 |
| Cells_Correlation_K_Mito_DNA | 3.3E-06 | 5.3E-07 | -7.4E+00 |
| Cells_Correlation_Overlap_DNA_AGP | 2.2E-15 | 5.6E-15 | -3.7E+01 |
| Cells_Correlation_Overlap_DNA_Lipid | 2.3E-09 | 7.6E-10 | -1.3E+01 |
| Cells_Correlation_Overlap_Mito_AGP | 3.2E-17 | 2.6E-16 | 5.0E+01 |
| Cells_Correlation_Overlap_Mito_Lipid | 3.4E-07 | 6.7E-08 | 9.0E+00 |
| Cells_Granularity_10_AGP | 1.2E-12 | 9.4E-13 | 2.3E+01 |
| Cells_Granularity_10_Lipid | 2.5E-10 | 1.1E-10 | 1.6E+01 |
| Cells_Granularity_10_Mito | 2.1E-13 | 2.2E-13 | 2.7E+01 |
| Cells_Granularity_1_AGP | 9.7E-19 | 2.8E-17 | -6.5E+01 |
| Cells_Granularity_4_Lipid | 1.1E-15 | 3.5E-15 | 3.9E+01 |
| Cells_Granularity_3_Mito | 3.8E-18 | 5.8E-17 | 5.9E+01 |
| Cells_Intensity_IntegratedIntensity_AGP | 4.1E-03 | 3.6E-04 | -3.4E+00 |
| Cells_Intensity_IntegratedIntensity_Lipid | 2.1E-09 | 7.1E-10 | -1.3E+01 |
| Cells_Intensity_IntegratedIntensity_DNA | 6.1E-10 | 2.3E-10 | -1.5E+01 |
| Cells_Intensity_IntegratedIntensity_Mito | 6.2E-03 | 5.4E-04 | -3.2E+00 |
| Cells_Intensity_IntegratedIntensityEdge_AGP | 1.1E-05 | 1.5E-06 | -6.7E+00 |
| Cells_Intensity_IntegratedIntensityEdge_Lipid | 1.0E-12 | 8.2E-13 | -2.4E+01 |
| Cells_Intensity_IntegratedIntensityEdge_DNA | 1.3E-12 | 9.8E-13 | -2.3E+01 |
| Cells_Intensity_IntegratedIntensityEdge_Mito | 4.5E-10 | 1.8E-10 | -1.5E+01 |
| Cells_Intensity_LowerQuartileIntensity_AGP | 2.0E-07 | 4.1E-08 | 9.4E+00 |
| Cells_Intensity_LowerQuartileIntensity_Lipid | 4.7E-07 | 8.8E-08 | -8.8E+00 |
| Cells_Intensity_LowerQuartileIntensity_DNA | 1.7E-08 | 4.5E-09 | -1.1E+01 |
| Cells_Intensity_MADIntensity_AGP | 4.0E-07 | 7.8E-08 | 8.9E+00 |
| Cells_Intensity_MADIntensity_Lipid | 3.3E-03 | 2.9E-04 | 3.5E+00 |
| Cells_Intensity_MADIntensity_DNA | 5.6E-08 | 1.3E-08 | 1.0E+01 |
| Cells_Intensity_MADIntensity_Mito | 5.2E-08 | 1.2E-08 | 1.0E+01 |
| Cells_Intensity_MassDisplacement_AGP | 4.0E-12 | 2.6E-12 | -2.2E+01 |
| Cells_Intensity_MassDisplacement_Lipid | 6.5E-04 | 6.6E-05 | 4.4E+00 |
| Cells_Intensity_MassDisplacement_DNA | 1.3E-11 | 8.0E-12 | -2.0E+01 |
| Cells_Intensity_MassDisplacement_Mito | 1.6E-10 | 7.0E-11 | 1.6E+01 |
| Cells_Intensity_MaxIntensity_AGP | 1.4E-07 | 3.0E-08 | -9.7E+00 |
| Cells_Intensity_MaxIntensity_Lipid | 7.0E-07 | 1.3E-07 | -8.5E+00 |
| Cells_Intensity_MaxIntensityEdge_Lipid | 2.5E-09 | 8.3E-10 | -1.3E+01 |
| Cells_Intensity_MeanIntensity_Lipid | 6.1E-06 | 9.0E-07 | -7.0E+00 |
| Cells_Intensity_MeanIntensity_DNA | 1.4E-06 | 2.4E-07 | -8.0E+00 |
| Cells_Intensity_MeanIntensity_Mito | 1.0E-02 | 8.9E-04 | 3.0E+00 |
| Cells_Intensity_MeanIntensityEdge_Lipid | 1.6E-10 | 7.1E-11 | -1.6E+01 |
| Cells_Intensity_MeanIntensityEdge_DNA | 5.0E-09 | 1.5E-09 | -1.3E+01 |
| Cells_Intensity_MeanIntensityEdge_Mito | 1.3E-05 | 1.8E-06 | -6.5E+00 |
| Cells_Intensity_MedianIntensity_AGP | 8.2E-08 | 1.8E-08 | 1.0E+01 |
| Cells_Intensity_MedianIntensity_Lipid | 3.8E-04 | 4.0E-05 | -4.6E+00 |
| Cells_Intensity_MedianIntensity_DNA | 9.5E-04 | 9.3E-05 | -4.2E+00 |
| Cells_Intensity_MedianIntensity_Mito | 1.6E-03 | 1.5E-04 | 3.9E+00 |
| Cells_Intensity_MinIntensity_AGP | 1.8E-09 | 6.3E-10 | -1.4E+01 |
| Cells_Intensity_MinIntensity_Lipid | 1.0E-11 | 6.4E-12 | -2.0E+01 |
| Cells_Intensity_MinIntensity_DNA | 1.4E-14 | 2.3E-14 | -3.3E+01 |
| Cells_Intensity_MinIntensity_Mito | 1.6E-13 | 1.7E-13 | -2.7E+01 |
| Cells_Intensity_MinIntensityEdge_AGP | 1.1E-09 | 4.0E-10 | -1.4E+01 |
| Cells_Intensity_MinIntensityEdge_Lipid | 1.0E-11 | 6.3E-12 | -2.0E+01 |
| Cells_Intensity_MinIntensityEdge_DNA | 3.6E-15 | 8.2E-15 | -3.6E+01 |
| Cells_Intensity_MinIntensityEdge_Mito | 8.4E-14 | 1.0E-13 | -2.9E+01 |
| Cells_Intensity_StdIntensity_AGP | 2.5E-08 | 6.3E-09 | -1.1E+01 |
| Cells_Intensity_StdIntensity_Lipid | 8.6E-05 | 9.9E-06 | -5.5E+00 |
| Cells_Intensity_StdIntensity_DNA | 4.0E-04 | 4.2E-05 | -4.6E+00 |
| Cells_Intensity_StdIntensity_Mito | 1.4E-06 | 2.5E-07 | 8.0E+00 |
| Cells_Intensity_StdIntensityEdge_AGP | 7.0E-05 | 8.2E-06 | 5.6E+00 |
| Cells_Intensity_StdIntensityEdge_Lipid | 3.2E-08 | 8.0E-09 | -1.1E+01 |
| Cells_Intensity_StdIntensityEdge_DNA | 4.4E-07 | 8.4E-08 | 8.8E+00 |
| Cells_Intensity_StdIntensityEdge_Mito | 2.1E-08 | 5.3E-09 | 1.1E+01 |
| Cells_Intensity_UpperQuartileIntensity_AGP | 1.3E-04 | 1.4E-05 | 5.2E+00 |
| Cells_Intensity_UpperQuartileIntensity_Lipid | 7.4E-04 | 7.5E-05 | -4.3E+00 |
| Cells_Intensity_UpperQuartileIntensity_DNA | 1.5E-05 | 2.0E-06 | -6.5E+00 |
| Cells_Intensity_UpperQuartileIntensity_Mito | 1.5E-04 | 1.7E-05 | 5.1E+00 |
| Cells_Location_Center_Y | 9.3E-04 | 9.2E-05 | 4.2E+00 |
| Cells_Location_CenterMassIntensity_Y_AGP | 9.6E-04 | 9.4E-05 | 4.2E+00 |
| Cells_Location_CenterMassIntensity_Y_Lipid | 9.1E-04 | 9.0E-05 | 4.2E+00 |
| Cells_Location_CenterMassIntensity_Y_DNA | 1.0E-03 | 9.8E-05 | 4.1E+00 |
| Cells_Location_CenterMassIntensity_Y_Mito | 8.7E-04 | 8.6E-05 | 4.2E+00 |
| Cells_Location_MaxIntensity_Y_AGP | 1.2E-03 | 1.1E-04 | 4.1E+00 |
| Cells_Location_MaxIntensity_Y_Lipid | 1.0E-03 | 9.8E-05 | 4.1E+00 |
| Cells_Location_MaxIntensity_Y_DNA | 9.5E-04 | 9.3E-05 | 4.2E+00 |
| Cells_Location_MaxIntensity_Y_Mito | 1.2E-03 | 1.2E-04 | 4.0E+00 |
| Cells_Mean_LargeLipidObjects_AreaShape_Area | 3.8E-07 | 7.3E-08 | -8.9E+00 |
| Cells_Mean_LargeLipidObjects_AreaShape_Center_X | 2.8E-07 | 5.7E-08 | -9.1E+00 |
| Cells_Mean_LargeLipidObjects_AreaShape_Center_Y | 4.4E-07 | 8.5E-08 | -8.8E+00 |
| Cells_Mean_LargeLipidObjects_AreaShape_Center_Z | 3.7E-07 | 7.2E-08 | -8.9E+00 |
| Cells_Mean_LargeLipidObjects_AreaShape_Compactness | 4.3E-07 | 8.3E-08 | -8.8E+00 |
| Cells_Mean_LargeLipidObjects_AreaShape_Eccentricity | 3.7E-07 | 7.2E-08 | -8.9E+00 |
| Cells_Mean_LargeLipidObjects_AreaShape_EulerNumber | 3.7E-07 | 7.2E-08 | -8.9E+00 |
| Cells_Mean_LargeLipidObjects_AreaShape_Extent | 4.1E-07 | 7.8E-08 | -8.9E+00 |
| Cells_Mean_LargeLipidObjects_AreaShape_FormFactor | 1.2E-06 | 2.1E-07 | -8.1E+00 |
| Cells_Mean_LargeLipidObjects_AreaShape_MajorAxisLength | 4.1E-07 | 7.9E-08 | -8.9E+00 |
| Cells_Mean_LargeLipidObjects_AreaShape_MaxFerretDiameter | 3.9E-07 | 7.6E-08 | -8.9E+00 |
| Cells_Mean_LargeLipidObjects_AreaShape_MaximumRadius | 5.6E-07 | 1.0E-07 | -8.6E+00 |
| Cells_Mean_LargeLipidObjects_AreaShape_MeanRadius | 5.9E-07 | 1.1E-07 | -8.6E+00 |
| Cells_Mean_LargeLipidObjects_AreaShape_MedianRadius | 5.6E-07 | 1.0E-07 | -8.6E+00 |
| Cells_Mean_LargeLipidObjects_AreaShape_MinFeretDiameter | 3.0E-07 | 6.0E-08 | -9.1E+00 |
| Cells_Mean_LargeLipidObjects_AreaShape_MinorAxisLength | 3.2E-07 | 6.4E-08 | -9.0E+00 |
| Cells_Mean_LargeLipidObjects_AreaShape_Perimeter | 2.1E-07 | 4.2E-08 | -9.4E+00 |
| Cells_Mean_LargeLipidObjects_AreaShape_Solidity | 4.6E-07 | 8.8E-08 | -8.8E+00 |
| Cells_Mean_LargeLipidObjects_Correlation_Correlation_Lipid_AGP | 1.3E-07 | 2.9E-08 | 9.7E+00 |
| Cells_Mean_LargeLipidObjects_Correlation_Correlation_DNA_AGP | 6.0E-07 | 1.1E-07 | -8.6E+00 |
| Cells_Mean_LargeLipidObjects_Correlation_Correlation_DNA_Lipid | 8.4E-08 | 1.9E-08 | 1.0E+01 |
| Cells_Mean_LargeLipidObjects_Correlation_Correlation_DNA_Mito | 2.3E-07 | 4.7E-08 | -9.3E+00 |
| Cells_Mean_LargeLipidObjects_Correlation_Correlation_Mito_AGP | 1.2E-06 | 2.1E-07 | -8.1E+00 |
| Cells_Mean_LargeLipidObjects_Correlation_Correlation_Mito_Lipid | 2.8E-07 | 5.6E-08 | 9.1E+00 |
| Cells_Mean_LargeLipidObjects_Correlation_K_AGP_Lipid | 7.7E-09 | 2.2E-09 | -1.2E+01 |
| Cells_Mean_LargeLipidObjects_Correlation_K_AGP_DNA | 6.2E-09 | 1.8E-09 | -1.2E+01 |
| Cells_Mean_LargeLipidObjects_Correlation_K_AGP_Mito | 3.0E-08 | 7.4E-09 | -1.1E+01 |
| Cells_Mean_LargeLipidObjects_Correlation_K_DNA_Lipid | 8.7E-08 | 1.9E-08 | -1.0E+01 |
| Cells_Mean_LargeLipidObjects_Correlation_K_DNA_Mito | 3.7E-04 | 3.9E-05 | -4.7E+00 |
| Cells_Mean_LargeLipidObjects_Correlation_K_Mito_AGP | 7.4E-05 | 8.7E-06 | -5.5E+00 |
| Cells_Mean_LargeLipidObjects_Correlation_K_Mito_Lipid | 1.1E-07 | 2.3E-08 | -9.9E+00 |
| Cells_Mean_LargeLipidObjects_Correlation_K_Mito_DNA | 1.3E-07 | 2.8E-08 | -9.7E+00 |
| Cells_Mean_LargeLipidObjects_Correlation_Overlap_Lipid_AGP | 5.1E-07 | 9.5E-08 | -8.7E+00 |
| Cells_Mean_LargeLipidObjects_Correlation_Overlap_DNA_AGP | 3.9E-07 | 7.6E-08 | -8.9E+00 |
| Cells_Mean_LargeLipidObjects_Correlation_Overlap_DNA_Lipid | 5.2E-07 | 9.7E-08 | -8.7E+00 |
| Cells_Mean_LargeLipidObjects_Correlation_Overlap_DNA_Mito | 3.8E-07 | 7.3E-08 | -8.9E+00 |
| Cells_Mean_LargeLipidObjects_Correlation_Overlap_Mito_AGP | 4.7E-07 | 8.8E-08 | -8.8E+00 |
| Cells_Mean_LargeLipidObjects_Correlation_Overlap_Mito_Lipid | 6.2E-07 | 1.1E-07 | -8.6E+00 |
| Cells_Mean_LargeLipidObjects_Granularity_16_Lipid | 1.4E-04 | 1.5E-05 | -5.2E+00 |
| Cells_Mean_LargeLipidObjects_Granularity_1_Lipid | 2.0E-08 | 5.2E-09 | -1.1E+01 |
| Cells_Mean_LargeLipidObjects_Intensity_IntegratedIntensity_Lipid | 1.6E-07 | 3.3E-08 | -9.6E+00 |
| Cells_Mean_LargeLipidObjects_Intensity_IntegratedIntensityEdge_Lipid | 2.0E-07 | 4.2E-08 | -9.4E+00 |
| Cells_Mean_LargeLipidObjects_Intensity_LowerQuartileIntensity_Lipid | 2.2E-07 | 4.6E-08 | -9.3E+00 |
| Cells_Mean_LargeLipidObjects_Intensity_MADIntensity_Lipid | 5.7E-08 | 1.4E-08 | -1.0E+01 |
| Cells_Mean_LargeLipidObjects_Intensity_MassDisplacement_Lipid | 1.7E-06 | 2.8E-07 | -7.9E+00 |
| Cells_Mean_LargeLipidObjects_Intensity_MaxIntensity_Lipid | 1.6E-07 | 3.3E-08 | -9.6E+00 |
| Cells_Mean_LargeLipidObjects_Intensity_MaxIntensityEdge_Lipid | 4.9E-07 | 9.3E-08 | -8.7E+00 |
| Cells_Mean_LargeLipidObjects_Intensity_MeanIntensity_Lipid | 1.4E-07 | 2.9E-08 | -9.7E+00 |
| Cells_Mean_LargeLipidObjects_Intensity_MeanIntensityEdge_Lipid | 3.2E-07 | 6.3E-08 | -9.0E+00 |
| Cells_Mean_LargeLipidObjects_Intensity_MedianIntensity_Lipid | 1.2E-07 | 2.7E-08 | -9.8E+00 |
| Cells_Mean_LargeLipidObjects_Intensity_MinIntensity_Lipid | 7.6E-07 | 1.4E-07 | -8.4E+00 |
| Cells_Mean_LargeLipidObjects_Intensity_MinIntensityEdge_Lipid | 6.9E-07 | 1.3E-07 | -8.5E+00 |
| Cells_Mean_LargeLipidObjects_Intensity_StdIntensity_Lipid | 1.3E-07 | 2.8E-08 | -9.7E+00 |
| Cells_Mean_LargeLipidObjects_Intensity_StdIntensityEdge_Lipid | 5.3E-07 | 9.9E-08 | -8.7E+00 |
| Cells_Mean_LargeLipidObjects_Intensity_UpperQuartileIntensity_Lipid | 1.1E-07 | 2.4E-08 | -9.9E+00 |
| Cells_Mean_LargeLipidObjects_Location_Center_X | 2.8E-07 | 5.7E-08 | -9.1E+00 |
| Cells_Mean_LargeLipidObjects_Location_Center_Y | 4.4E-07 | 8.5E-08 | -8.8E+00 |
| Cells_Mean_LargeLipidObjects_Location_CenterMassIntensity_X_Lipid | 2.8E-07 | 5.7E-08 | -9.1E+00 |
| Cells_Mean_LargeLipidObjects_Location_CenterMassIntensity_Y_Lipid | 4.4E-07 | 8.5E-08 | -8.8E+00 |
| Cells_Mean_LargeLipidObjects_Location_MaxIntensity_X_Lipid | 2.8E-07 | 5.7E-08 | -9.1E+00 |
| Cells_Mean_LargeLipidObjects_Location_MaxIntensity_Y_Lipid | 4.5E-07 | 8.5E-08 | -8.8E+00 |
| Cells_Mean_LargeLipidObjects_Number_Object_Number | 8.8E-07 | 1.6E-07 | -8.3E+00 |
| Cells_Neighbors_AngleBetweenNeighbors_10 | 1.0E-12 | 8.3E-13 | -2.4E+01 |
| Cells_Neighbors_AngleBetweenNeighbors_Adjacent | 1.0E-12 | 8.3E-13 | -2.4E+01 |
| Cells_Neighbors_FirstClosestObjectNumber_10 | 6.4E-09 | 1.8E-09 | -1.2E+01 |
| Cells_Neighbors_FirstClosestObjectNumber_Adjacent | 6.4E-09 | 1.8E-09 | -1.2E+01 |
| Cells_Neighbors_NumberOfNeighbors_10 | 3.9E-16 | 1.4E-15 | -4.2E+01 |
| Cells_Neighbors_NumberOfNeighbors_Adjacent | 2.2E-16 | 9.4E-16 | -4.4E+01 |
| Cells_Neighbors_PercentTouching_10 | 1.8E-15 | 4.8E-15 | -3.8E+01 |
| Cells_Neighbors_PercentTouching_Adjacent | 3.8E-11 | 2.0E-11 | -1.8E+01 |
| Cells_Neighbors_SecondClosestDistance_10 | 1.7E-03 | 1.6E-04 | 3.9E+00 |
| Cells_Neighbors_SecondClosestDistance_Adjacent | 1.7E-03 | 1.6E-04 | 3.9E+00 |
| Cells_Neighbors_SecondClosestObjectNumber_10 | 5.6E-09 | 1.7E-09 | -1.2E+01 |
| Cells_Neighbors_SecondClosestObjectNumber_Adjacent | 5.6E-09 | 1.7E-09 | -1.2E+01 |
| Cells_Number_Object_Number | 6.0E-09 | 1.7E-09 | -1.2E+01 |
| Cells_Parent_Nuclei | 6.1E-09 | 1.8E-09 | -1.2E+01 |
| Cells_RadialDistribution_FracAtD_AGP_2of4 | 3.2E-12 | 2.2E-12 | -2.2E+01 |
| Cells_RadialDistribution_FracAtD_Lipid_4of4 | 5.3E-18 | 6.6E-17 | -5.7E+01 |
| Cells_RadialDistribution_FracAtD_DNA_1of4 | 3.4E-14 | 4.9E-14 | 3.0E+01 |
| Cells_RadialDistribution_FracAtD_Mito_1of4 | 7.4E-17 | 4.8E-16 | 4.7E+01 |
| Cells_RadialDistribution_MeanFrac_AGP_2of4 | 4.6E-16 | 1.6E-15 | -4.2E+01 |
| Cells_RadialDistribution_MeanFrac_Lipid_4of4 | 3.9E-16 | 1.4E-15 | -4.2E+01 |
| Cells_RadialDistribution_MeanFrac_DNA_3of4 | 6.3E-16 | 2.1E-15 | -4.1E+01 |
| Cells_RadialDistribution_MeanFrac_Mito_4of4 | 7.5E-16 | 2.5E-15 | -4.0E+01 |
| Cells_RadialDistribution_RadialCV_AGP_3of4 | 2.2E-16 | 9.4E-16 | -4.4E+01 |
| Cells_RadialDistribution_RadialCV_Lipid_4of4 | 2.2E-09 | 7.4E-10 | 1.3E+01 |
| Cells_RadialDistribution_RadialCV_DNA_4of4 | 2.4E-10 | 1.0E-10 | 1.6E+01 |
| Cells_RadialDistribution_RadialCV_Mito_4of4 | 2.3E-16 | 9.4E-16 | 4.4E+01 |
| Cells_Texture_AngularSecondMoment_AGP_5_01 | 8.5E-08 | 1.9E-08 | -1.0E+01 |
| Cells_Texture_AngularSecondMoment_DNA_20_00 | 9.4E-07 | 1.7E-07 | -8.3E+00 |
| Cells_Texture_AngularSecondMoment_Mito_20_02 | 6.7E-05 | 7.9E-06 | -5.6E+00 |
| Cells_Texture_Contrast_AGP_20_00 | 1.6E-09 | 5.4E-10 | -1.4E+01 |
| Cells_Texture_Contrast_Lipid_5_02 | 5.8E-05 | 7.0E-06 | -5.7E+00 |
| Cells_Texture_Contrast_DNA_20_03 | 2.1E-04 | 2.3E-05 | -4.9E+00 |
| Cells_Texture_Contrast_Mito_20_01 | 1.7E-06 | 2.8E-07 | 7.9E+00 |
| Cells_Texture_Correlation_AGP_5_00 | 1.3E-14 | 2.2E-14 | -3.3E+01 |
| Cells_Texture_Correlation_Lipid_20_01 | 6.7E-13 | 5.9E-13 | -2.5E+01 |
| Cells_Texture_Correlation_DNA_20_03 | 1.4E-18 | 3.0E-17 | -6.3E+01 |
| Cells_Texture_Correlation_Mito_20_01 | 4.4E-17 | 3.1E-16 | -4.9E+01 |
| Cells_Texture_DifferenceEntropy_AGP_20_00 | 1.5E-05 | 2.0E-06 | -6.5E+00 |
| Cells_Texture_DifferenceEntropy_Lipid_20_01 | 2.2E-03 | 2.0E-04 | -3.7E+00 |
| Cells_Texture_DifferenceEntropy_DNA_10_03 | 2.9E-07 | 5.8E-08 | 9.1E+00 |
| Cells_Texture_DifferenceEntropy_Mito_10_01 | 1.3E-05 | 1.7E-06 | 6.6E+00 |
| Cells_Texture_DifferenceVariance_Lipid_5_00 | 6.4E-04 | 6.5E-05 | 4.4E+00 |
| Cells_Texture_DifferenceVariance_DNA_10_03 | 4.6E-06 | 7.0E-07 | -7.2E+00 |
| Cells_Texture_DifferenceVariance_Mito_20_01 | 2.7E-04 | 2.9E-05 | -4.8E+00 |
| Cells_Texture_Entropy_AGP_5_00 | 1.7E-05 | 2.3E-06 | 6.4E+00 |
| Cells_Texture_Entropy_Lipid_20_01 | 7.0E-04 | 7.1E-05 | -4.3E+00 |
| Cells_Texture_Entropy_DNA_10_03 | 4.4E-08 | 1.1E-08 | 1.1E+01 |
| Cells_Texture_Entropy_Mito_10_01 | 6.7E-06 | 9.8E-07 | 7.0E+00 |
| Cells_Texture_InfoMeas1_AGP_5_00 | 1.3E-14 | 2.2E-14 | 3.3E+01 |
| Cells_Texture_InfoMeas1_Lipid_5_00 | 1.6E-05 | 2.2E-06 | -6.4E+00 |
| Cells_Texture_InfoMeas1_DNA_20_03 | 7.7E-14 | 9.5E-14 | -2.9E+01 |
| Cells_Texture_InfoMeas1_Mito_20_01 | 3.4E-13 | 3.3E-13 | -2.6E+01 |
| Cells_Texture_InverseDifferenceMoment_AGP_20_03 | 1.0E-08 | 2.8E-09 | -1.2E+01 |
| Cells_Texture_InverseDifferenceMoment_Lipid_20_00 | 1.2E-03 | 1.1E-04 | -4.1E+00 |
| Cells_Texture_InverseDifferenceMoment_DNA_10_03 | 2.2E-10 | 9.4E-11 | -1.6E+01 |
| Cells_Texture_InverseDifferenceMoment_Mito_20_01 | 9.7E-09 | 2.7E-09 | -1.2E+01 |
| Cells_Texture_SumAverage_Lipid_20_01 | 7.9E-07 | 1.4E-07 | -8.4E+00 |
| Cells_Texture_SumAverage_DNA_20_03 | 2.9E-07 | 5.7E-08 | -9.1E+00 |
| Cells_Texture_SumAverage_Mito_10_00 | 2.1E-03 | 2.0E-04 | 3.8E+00 |
| Cells_Texture_SumEntropy_AGP_20_00 | 1.5E-03 | 1.5E-04 | -3.9E+00 |
| Cells_Texture_SumEntropy_Lipid_20_01 | 8.8E-05 | 1.0E-05 | -5.4E+00 |
| Cells_Texture_SumEntropy_DNA_10_01 | 4.7E-06 | 7.2E-07 | 7.2E+00 |
| Cells_Texture_SumEntropy_Mito_5_00 | 2.4E-06 | 4.0E-07 | 7.6E+00 |
| Cells_Texture_SumVariance_AGP_5_01 | 1.3E-09 | 4.7E-10 | -1.4E+01 |
| Cells_Texture_SumVariance_Lipid_20_01 | 9.8E-06 | 1.4E-06 | -6.7E+00 |
| Cells_Texture_SumVariance_DNA_20_03 | 4.1E-05 | 5.2E-06 | -5.9E+00 |
| Cells_Texture_SumVariance_Mito_5_02 | 2.7E-06 | 4.4E-07 | 7.5E+00 |
| Cells_Texture_Variance_AGP_10_03 | 2.3E-09 | 7.6E-10 | -1.3E+01 |
| Cells_Texture_Variance_Lipid_20_01 | 3.2E-05 | 4.1E-06 | -6.0E+00 |
| Cells_Texture_Variance_DNA_20_03 | 1.1E-04 | 1.2E-05 | -5.3E+00 |
| Cells_Texture_Variance_Mito_10_01 | 5.3E-06 | 8.0E-07 | 7.1E+00 |
| Cytoplasm_AreaShape_Area | 2.0E-11 | 1.1E-11 | -1.9E+01 |
| Cytoplasm_AreaShape_Center_Y | 9.2E-04 | 9.1E-05 | 4.2E+00 |
| Cytoplasm_AreaShape_Eccentricity | 3.9E-06 | 6.1E-07 | -7.3E+00 |
| Cytoplasm_AreaShape_EulerNumber | 2.3E-12 | 1.6E-12 | 2.2E+01 |
| Cytoplasm_AreaShape_Extent | 5.4E-10 | 2.1E-10 | 1.5E+01 |
| Cytoplasm_AreaShape_FormFactor | 4.1E-14 | 5.7E-14 | 3.0E+01 |
| Cytoplasm_AreaShape_MajorAxisLength | 6.5E-12 | 4.2E-12 | -2.1E+01 |
| Cytoplasm_AreaShape_MaxFeretDiameter | 1.0E-12 | 8.3E-13 | -2.4E+01 |
| Cytoplasm_AreaShape_MaximumRadius | 7.2E-11 | 3.6E-11 | -1.7E+01 |
| Cytoplasm_AreaShape_MeanRadius | 3.2E-08 | 7.9E-09 | -1.1E+01 |
| Cytoplasm_AreaShape_MedianRadius | 2.0E-05 | 2.6E-06 | -6.3E+00 |
| Cytoplasm_AreaShape_MinFeretDiameter | 1.3E-12 | 9.6E-13 | -2.3E+01 |
| Cytoplasm_AreaShape_MinorAxisLength | 1.4E-11 | 8.4E-12 | -2.0E+01 |
| Cytoplasm_AreaShape_Perimeter | 3.5E-15 | 8.0E-15 | -3.6E+01 |
| Cytoplasm_AreaShape_Solidity | 6.9E-11 | 3.4E-11 | 1.7E+01 |
| Cytoplasm_AreaShape_Zernike_4_0 | 8.3E-17 | 5.2E-16 | 4.7E+01 |
| Cytoplasm_Correlation_Correlation_Lipid_AGP | 2.0E-13 | 2.1E-13 | 2.7E+01 |
| Cytoplasm_Correlation_Correlation_DNA_AGP | 2.7E-08 | 6.7E-09 | -1.1E+01 |
| Cytoplasm_Correlation_Correlation_DNA_Lipid | 4.3E-14 | 5.8E-14 | 3.0E+01 |
| Cytoplasm_Correlation_Correlation_DNA_Mito | 2.0E-07 | 4.0E-08 | 9.4E+00 |
| Cytoplasm_Correlation_Correlation_Mito_AGP | 4.2E-14 | 5.8E-14 | 3.0E+01 |
| Cytoplasm_Correlation_Correlation_Mito_Lipid | 3.3E-13 | 3.2E-13 | 2.6E+01 |
| Cytoplasm_Correlation_K_AGP_Lipid | 3.3E-08 | 8.1E-09 | -1.1E+01 |
| Cytoplasm_Correlation_K_AGP_DNA | 1.8E-07 | 3.7E-08 | -9.5E+00 |
| Cytoplasm_Correlation_K_Lipid_AGP | 4.3E-07 | 8.2E-08 | 8.8E+00 |
| Cytoplasm_Correlation_K_Lipid_DNA | 7.0E-07 | 1.3E-07 | 8.5E+00 |
| Cytoplasm_Correlation_K_Lipid_Mito | 1.5E-07 | 3.1E-08 | 9.6E+00 |
| Cytoplasm_Correlation_K_DNA_AGP | 1.6E-06 | 2.7E-07 | 7.9E+00 |
| Cytoplasm_Correlation_K_DNA_Lipid | 5.8E-07 | 1.1E-07 | -8.6E+00 |
| Cytoplasm_Correlation_K_DNA_Mito | 8.8E-05 | 1.0E-05 | 5.4E+00 |
| Cytoplasm_Correlation_K_Mito_Lipid | 2.6E-07 | 5.3E-08 | -9.2E+00 |
| Cytoplasm_Correlation_K_Mito_DNA | 1.0E-03 | 1.0E-04 | -4.1E+00 |
| Cytoplasm_Correlation_Overlap_Lipid_AGP | 1.6E-07 | 3.4E-08 | 9.6E+00 |
| Cytoplasm_Correlation_Overlap_DNA_AGP | 3.1E-08 | 7.8E-09 | -1.1E+01 |
| Cytoplasm_Correlation_Overlap_DNA_Lipid | 1.3E-08 | 3.4E-09 | 1.2E+01 |
| Cytoplasm_Correlation_Overlap_Mito_AGP | 3.4E-15 | 8.0E-15 | 3.6E+01 |
| Cytoplasm_Correlation_Overlap_Mito_Lipid | 4.5E-07 | 8.6E-08 | 8.8E+00 |
| Cytoplasm_Granularity_10_AGP | 2.2E-12 | 1.5E-12 | 2.2E+01 |
| Cytoplasm_Granularity_10_Lipid | 1.7E-10 | 7.6E-11 | 1.6E+01 |
| Cytoplasm_Granularity_10_Mito | 2.7E-13 | 2.7E-13 | 2.6E+01 |
| Cytoplasm_Granularity_1_AGP | 1.2E-18 | 2.8E-17 | -6.4E+01 |
| Cytoplasm_Granularity_4_Lipid | 1.7E-15 | 4.5E-15 | 3.8E+01 |
| Cytoplasm_Granularity_3_Mito | 4.1E-18 | 5.8E-17 | 5.8E+01 |
| Cytoplasm_Intensity_IntegratedIntensity_Lipid | 1.3E-09 | 4.7E-10 | -1.4E+01 |
| Cytoplasm_Intensity_IntegratedIntensity_DNA | 4.1E-09 | 1.2E-09 | -1.3E+01 |
| Cytoplasm_Intensity_IntegratedIntensity_Mito | 4.3E-04 | 4.5E-05 | -4.6E+00 |
| Cytoplasm_Intensity_IntegratedIntensityEdge_AGP | 3.6E-05 | 4.6E-06 | -5.9E+00 |
| Cytoplasm_Intensity_IntegratedIntensityEdge_Lipid | 2.9E-11 | 1.6E-11 | -1.9E+01 |
| Cytoplasm_Intensity_IntegratedIntensityEdge_DNA | 1.7E-12 | 1.2E-12 | -2.3E+01 |
| Cytoplasm_Intensity_IntegratedIntensityEdge_Mito | 3.3E-07 | 6.6E-08 | -9.0E+00 |
| Cytoplasm_Intensity_LowerQuartileIntensity_AGP | 4.8E-07 | 9.1E-08 | 8.7E+00 |
| Cytoplasm_Intensity_LowerQuartileIntensity_Lipid | 7.9E-08 | 1.8E-08 | -1.0E+01 |
| Cytoplasm_Intensity_LowerQuartileIntensity_DNA | 1.4E-09 | 4.8E-10 | -1.4E+01 |
| Cytoplasm_Intensity_LowerQuartileIntensity_Mito | 6.4E-03 | 5.6E-04 | -3.2E+00 |
| Cytoplasm_Intensity_MADIntensity_AGP | 3.5E-09 | 1.1E-09 | 1.3E+01 |
| Cytoplasm_Intensity_MADIntensity_Lipid | 1.3E-03 | 1.3E-04 | 4.0E+00 |
| Cytoplasm_Intensity_MADIntensity_DNA | 1.8E-07 | 3.7E-08 | 9.5E+00 |
| Cytoplasm_Intensity_MADIntensity_Mito | 2.7E-08 | 6.8E-09 | 1.1E+01 |
| Cytoplasm_Intensity_MassDisplacement_AGP | 5.8E-03 | 5.1E-04 | -3.2E+00 |
| Cytoplasm_Intensity_MassDisplacement_Lipid | 1.7E-04 | 1.9E-05 | 5.1E+00 |
| Cytoplasm_Intensity_MassDisplacement_Mito | 6.9E-11 | 3.4E-11 | 1.7E+01 |
| Cytoplasm_Intensity_MaxIntensity_AGP | 8.4E-03 | 7.2E-04 | 3.1E+00 |
| Cytoplasm_Intensity_MaxIntensity_Lipid | 5.3E-07 | 9.9E-08 | -8.7E+00 |
| Cytoplasm_Intensity_MaxIntensity_DNA | 2.3E-08 | 5.9E-09 | -1.1E+01 |
| Cytoplasm_Intensity_MaxIntensityEdge_Lipid | 1.7E-07 | 3.6E-08 | -9.5E+00 |
| Cytoplasm_Intensity_MaxIntensityEdge_DNA | 5.5E-09 | 1.6E-09 | -1.3E+01 |
| Cytoplasm_Intensity_MeanIntensity_AGP | 5.7E-07 | 1.1E-07 | 8.6E+00 |
| Cytoplasm_Intensity_MeanIntensity_Lipid | 3.0E-06 | 4.8E-07 | -7.5E+00 |
| Cytoplasm_Intensity_MeanIntensity_DNA | 1.5E-05 | 2.1E-06 | -6.4E+00 |
| Cytoplasm_Intensity_MeanIntensityEdge_Lipid | 1.4E-08 | 3.7E-09 | -1.2E+01 |
| Cytoplasm_Intensity_MeanIntensityEdge_DNA | 1.7E-08 | 4.5E-09 | -1.1E+01 |
| Cytoplasm_Intensity_MedianIntensity_AGP | 3.3E-08 | 8.0E-09 | 1.1E+01 |
| Cytoplasm_Intensity_MedianIntensity_Lipid | 5.9E-05 | 7.1E-06 | -5.7E+00 |
| Cytoplasm_Intensity_MedianIntensity_DNA | 2.6E-05 | 3.3E-06 | -6.1E+00 |
| Cytoplasm_Intensity_MinIntensity_AGP | 1.9E-09 | 6.4E-10 | -1.4E+01 |
| Cytoplasm_Intensity_MinIntensity_Lipid | 1.0E-11 | 6.4E-12 | -2.0E+01 |
| Cytoplasm_Intensity_MinIntensity_DNA | 1.4E-14 | 2.3E-14 | -3.3E+01 |
| Cytoplasm_Intensity_MinIntensity_Mito | 1.5E-13 | 1.6E-13 | -2.7E+01 |
| Cytoplasm_Intensity_MinIntensityEdge_AGP | 1.1E-09 | 3.9E-10 | -1.4E+01 |
| Cytoplasm_Intensity_MinIntensityEdge_Lipid | 1.0E-11 | 6.2E-12 | -2.0E+01 |
| Cytoplasm_Intensity_MinIntensityEdge_DNA | 3.6E-15 | 8.2E-15 | -3.6E+01 |
| Cytoplasm_Intensity_MinIntensityEdge_Mito | 8.9E-14 | 1.1E-13 | -2.8E+01 |
| Cytoplasm_Intensity_StdIntensity_AGP | 1.4E-05 | 1.9E-06 | 6.5E+00 |
| Cytoplasm_Intensity_StdIntensity_Lipid | 3.9E-05 | 4.9E-06 | -5.9E+00 |
| Cytoplasm_Intensity_StdIntensity_Mito | 8.7E-07 | 1.6E-07 | 8.3E+00 |
| Cytoplasm_Intensity_StdIntensityEdge_Lipid | 2.4E-05 | 3.1E-06 | -6.2E+00 |
| Cytoplasm_Intensity_StdIntensityEdge_DNA | 6.3E-03 | 5.5E-04 | -3.2E+00 |
| Cytoplasm_Intensity_StdIntensityEdge_Mito | 2.9E-08 | 7.1E-09 | 1.1E+01 |
| Cytoplasm_Intensity_UpperQuartileIntensity_AGP | 1.0E-07 | 2.2E-08 | 9.9E+00 |
| Cytoplasm_Intensity_UpperQuartileIntensity_Lipid | 9.1E-04 | 9.0E-05 | -4.2E+00 |
| Cytoplasm_Intensity_UpperQuartileIntensity_Mito | 4.9E-04 | 5.0E-05 | 4.5E+00 |
| Cytoplasm_Location_Center_Y | 9.0E-04 | 9.0E-05 | 4.2E+00 |
| Cytoplasm_Location_CenterMassIntensity_Y_AGP | 9.0E-04 | 9.0E-05 | 4.2E+00 |
| Cytoplasm_Location_CenterMassIntensity_Y_Lipid | 8.6E-04 | 8.5E-05 | 4.2E+00 |
| Cytoplasm_Location_CenterMassIntensity_Y_DNA | 9.2E-04 | 9.1E-05 | 4.2E+00 |
| Cytoplasm_Location_CenterMassIntensity_Y_Mito | 8.3E-04 | 8.3E-05 | 4.2E+00 |
| Cytoplasm_Location_MaxIntensity_Y_AGP | 1.5E-03 | 1.4E-04 | 3.9E+00 |
| Cytoplasm_Location_MaxIntensity_Y_Lipid | 1.0E-03 | 9.8E-05 | 4.1E+00 |
| Cytoplasm_Location_MaxIntensity_Y_DNA | 1.6E-03 | 1.5E-04 | 3.9E+00 |
| Cytoplasm_Location_MaxIntensity_Y_Mito | 1.2E-03 | 1.2E-04 | 4.0E+00 |
| Cytoplasm_Number_Object_Number | 6.0E-09 | 1.7E-09 | -1.2E+01 |
| Cytoplasm_Parent_Cells | 6.1E-09 | 1.8E-09 | -1.2E+01 |
| Cytoplasm_Parent_Nuclei | 6.1E-09 | 1.8E-09 | -1.2E+01 |
| Cytoplasm_RadialDistribution_FracAtD_AGP_1of4 | 2.3E-14 | 3.4E-14 | 3.1E+01 |
| Cytoplasm_RadialDistribution_FracAtD_Lipid_2of4 | 5.3E-14 | 6.9E-14 | 2.9E+01 |
| Cytoplasm_RadialDistribution_FracAtD_DNA_1of4 | 7.4E-08 | 1.7E-08 | 1.0E+01 |
| Cytoplasm_RadialDistribution_FracAtD_Mito_1of4 | 3.2E-14 | 4.6E-14 | 3.1E+01 |
| Cytoplasm_RadialDistribution_MeanFrac_AGP_1of4 | 6.2E-14 | 7.8E-14 | 2.9E+01 |
| Cytoplasm_RadialDistribution_MeanFrac_Lipid_4of4 | 2.2E-16 | 9.4E-16 | -4.4E+01 |
| Cytoplasm_RadialDistribution_MeanFrac_DNA_2of4 | 3.1E-09 | 1.0E-09 | 1.3E+01 |
| Cytoplasm_RadialDistribution_MeanFrac_Mito_1of4 | 1.5E-15 | 4.3E-15 | 3.8E+01 |
| Cytoplasm_RadialDistribution_RadialCV_AGP_4of4 | 8.0E-08 | 1.8E-08 | 1.0E+01 |
| Cytoplasm_RadialDistribution_RadialCV_Lipid_4of4 | 7.3E-09 | 2.1E-09 | 1.2E+01 |
| Cytoplasm_RadialDistribution_RadialCV_DNA_4of4 | 1.4E-14 | 2.3E-14 | 3.2E+01 |
| Cytoplasm_RadialDistribution_RadialCV_Mito_4of4 | 2.3E-16 | 9.4E-16 | 4.4E+01 |
| Cytoplasm_Texture_AngularSecondMoment_AGP_5_01 | 4.8E-08 | 1.2E-08 | -1.1E+01 |
| Cytoplasm_Texture_AngularSecondMoment_DNA_20_00 | 1.7E-05 | 2.3E-06 | -6.4E+00 |
| Cytoplasm_Texture_AngularSecondMoment_Mito_20_02 | 3.3E-04 | 3.5E-05 | -4.7E+00 |
| Cytoplasm_Texture_Contrast_AGP_5_00 | 1.0E-06 | 1.8E-07 | 8.2E+00 |
| Cytoplasm_Texture_Contrast_Lipid_5_02 | 2.4E-05 | 3.1E-06 | -6.2E+00 |
| Cytoplasm_Texture_Contrast_DNA_20_03 | 5.6E-03 | 4.9E-04 | 3.3E+00 |
| Cytoplasm_Texture_Contrast_Mito_10_03 | 1.6E-06 | 2.6E-07 | 7.9E+00 |
| Cytoplasm_Texture_Correlation_AGP_10_03 | 1.1E-10 | 5.2E-11 | -1.7E+01 |
| Cytoplasm_Texture_Correlation_Lipid_20_01 | 1.1E-13 | 1.2E-13 | -2.8E+01 |
| Cytoplasm_Texture_Correlation_DNA_20_03 | 6.8E-15 | 1.3E-14 | -3.4E+01 |
| Cytoplasm_Texture_Correlation_Mito_20_01 | 3.8E-15 | 8.5E-15 | -3.6E+01 |
| Cytoplasm_Texture_DifferenceEntropy_AGP_5_00 | 6.0E-08 | 1.4E-08 | 1.0E+01 |
| Cytoplasm_Texture_DifferenceEntropy_Lipid_20_01 | 2.8E-03 | 2.5E-04 | -3.6E+00 |
| Cytoplasm_Texture_DifferenceEntropy_DNA_10_01 | 7.7E-04 | 7.7E-05 | 4.3E+00 |
| Cytoplasm_Texture_DifferenceEntropy_Mito_10_01 | 3.6E-06 | 5.7E-07 | 7.3E+00 |
| Cytoplasm_Texture_DifferenceVariance_AGP_10_00 | 3.9E-06 | 6.1E-07 | -7.3E+00 |
| Cytoplasm_Texture_DifferenceVariance_Lipid_5_00 | 1.5E-03 | 1.4E-04 | 3.9E+00 |
| Cytoplasm_Texture_DifferenceVariance_DNA_5_00 | 2.2E-03 | 2.0E-04 | 3.7E+00 |
| Cytoplasm_Texture_DifferenceVariance_Mito_20_02 | 2.0E-04 | 2.1E-05 | -5.0E+00 |
| Cytoplasm_Texture_Entropy_AGP_5_00 | 6.3E-09 | 1.8E-09 | 1.2E+01 |
| Cytoplasm_Texture_Entropy_Lipid_20_01 | 1.5E-03 | 1.4E-04 | -3.9E+00 |
| Cytoplasm_Texture_Entropy_DNA_10_01 | 8.9E-05 | 1.0E-05 | 5.4E+00 |
| Cytoplasm_Texture_Entropy_Mito_10_02 | 1.3E-05 | 1.7E-06 | 6.6E+00 |
| Cytoplasm_Texture_InfoMeas1_AGP_20_01 | 2.8E-10 | 1.2E-10 | -1.6E+01 |
| Cytoplasm_Texture_InfoMeas1_Lipid_5_00 | 7.6E-04 | 7.6E-05 | -4.3E+00 |
| Cytoplasm_Texture_InfoMeas1_DNA_20_03 | 2.3E-13 | 2.4E-13 | -2.7E+01 |
| Cytoplasm_Texture_InfoMeas1_Mito_5_00 | 1.0E-14 | 1.9E-14 | -3.3E+01 |
| Cytoplasm_Texture_InverseDifferenceMoment_AGP_20_03 | 3.6E-10 | 1.5E-10 | -1.5E+01 |
| Cytoplasm_Texture_InverseDifferenceMoment_Lipid_20_00 | 3.6E-04 | 3.8E-05 | -4.7E+00 |
| Cytoplasm_Texture_InverseDifferenceMoment_DNA_20_03 | 9.0E-09 | 2.5E-09 | -1.2E+01 |
| Cytoplasm_Texture_InverseDifferenceMoment_Mito_20_01 | 6.6E-09 | 1.9E-09 | -1.2E+01 |
| Cytoplasm_Texture_SumAverage_AGP_5_00 | 2.4E-07 | 4.9E-08 | 9.2E+00 |
| Cytoplasm_Texture_SumAverage_Lipid_20_01 | 1.1E-06 | 1.9E-07 | -8.2E+00 |
| Cytoplasm_Texture_SumAverage_DNA_20_03 | 5.2E-06 | 7.9E-07 | -7.1E+00 |
| Cytoplasm_Texture_SumEntropy_AGP_5_02 | 2.4E-08 | 6.1E-09 | 1.1E+01 |
| Cytoplasm_Texture_SumEntropy_Lipid_20_01 | 3.7E-04 | 3.9E-05 | -4.7E+00 |
| Cytoplasm_Texture_SumEntropy_DNA_5_02 | 1.3E-04 | 1.5E-05 | 5.2E+00 |
| Cytoplasm_Texture_SumEntropy_Mito_5_00 | 1.8E-06 | 3.0E-07 | 7.8E+00 |
| Cytoplasm_Texture_SumVariance_AGP_5_00 | 1.7E-04 | 1.8E-05 | 5.1E+00 |
| Cytoplasm_Texture_SumVariance_Lipid_20_01 | 1.6E-05 | 2.1E-06 | -6.4E+00 |
| Cytoplasm_Texture_SumVariance_DNA_5_02 | 8.9E-03 | 7.6E-04 | 3.0E+00 |
| Cytoplasm_Texture_SumVariance_Mito_5_00 | 1.9E-06 | 3.1E-07 | 7.8E+00 |
| Cytoplasm_Texture_Variance_AGP_5_03 | 4.9E-05 | 6.0E-06 | 5.8E+00 |
| Cytoplasm_Texture_Variance_Lipid_20_01 | 3.3E-05 | 4.2E-06 | -6.0E+00 |
| Cytoplasm_Texture_Variance_Mito_10_03 | 2.9E-06 | 4.7E-07 | 7.5E+00 |
| Nuclei_AreaShape_Center_Y | 9.7E-04 | 9.6E-05 | 4.2E+00 |
| Nuclei_AreaShape_Compactness | 1.5E-12 | 1.1E-12 | 2.3E+01 |
| Nuclei_AreaShape_Eccentricity | 5.5E-17 | 3.8E-16 | 4.8E+01 |
| Nuclei_AreaShape_EulerNumber | 1.3E-06 | 2.2E-07 | -8.0E+00 |
| Nuclei_AreaShape_Extent | 4.0E-14 | 5.5E-14 | -3.0E+01 |
| Nuclei_AreaShape_FormFactor | 1.8E-13 | 1.9E-13 | -2.7E+01 |
| Nuclei_AreaShape_MajorAxisLength | 1.7E-03 | 1.6E-04 | 3.9E+00 |
| Nuclei_AreaShape_MaxFeretDiameter | 6.8E-04 | 6.9E-05 | 4.3E+00 |
| Nuclei_AreaShape_MaximumRadius | 2.4E-07 | 4.9E-08 | -9.3E+00 |
| Nuclei_AreaShape_MeanRadius | 1.8E-05 | 2.4E-06 | -6.3E+00 |
| Nuclei_AreaShape_MedianRadius | 1.3E-04 | 1.5E-05 | -5.2E+00 |
| Nuclei_AreaShape_MinFeretDiameter | 2.8E-05 | 3.6E-06 | -6.1E+00 |
| Nuclei_AreaShape_MinorAxisLength | 3.8E-06 | 6.0E-07 | -7.3E+00 |
| Nuclei_AreaShape_Solidity | 6.2E-13 | 5.5E-13 | -2.5E+01 |
| Nuclei_AreaShape_Zernike_4_4 | 4.4E-18 | 5.8E-17 | 5.8E+01 |
| Nuclei_Correlation_Correlation_Lipid_AGP | 1.8E-08 | 4.6E-09 | -1.1E+01 |
| Nuclei_Correlation_Correlation_DNA_AGP | 2.2E-12 | 1.5E-12 | -2.3E+01 |
| Nuclei_Correlation_Correlation_DNA_Lipid | 1.9E-07 | 3.9E-08 | -9.4E+00 |
| Nuclei_Correlation_Correlation_DNA_Mito | 3.7E-07 | 7.2E-08 | 8.9E+00 |
| Nuclei_Correlation_Correlation_Mito_AGP | 8.3E-10 | 3.1E-10 | 1.4E+01 |
| Nuclei_Correlation_Correlation_Mito_Lipid | 5.4E-07 | 1.0E-07 | 8.7E+00 |
| Nuclei_Correlation_K_AGP_Lipid | 2.5E-09 | 8.1E-10 | 1.3E+01 |
| Nuclei_Correlation_K_AGP_DNA | 6.0E-06 | 8.9E-07 | 7.0E+00 |
| Nuclei_Correlation_K_AGP_Mito | 1.5E-08 | 3.9E-09 | 1.2E+01 |
| Nuclei_Correlation_K_Lipid_AGP | 5.5E-11 | 2.8E-11 | -1.8E+01 |
| Nuclei_Correlation_K_Lipid_DNA | 3.0E-05 | 3.9E-06 | -6.0E+00 |
| Nuclei_Correlation_K_Lipid_Mito | 9.8E-08 | 2.2E-08 | 1.0E+01 |
| Nuclei_Correlation_K_DNA_AGP | 5.7E-08 | 1.3E-08 | -1.0E+01 |
| Nuclei_Correlation_K_DNA_Lipid | 2.5E-03 | 2.2E-04 | 3.7E+00 |
| Nuclei_Correlation_K_DNA_Mito | 1.0E-07 | 2.2E-08 | 9.9E+00 |
| Nuclei_Correlation_K_Mito_AGP | 3.5E-11 | 1.9E-11 | -1.8E+01 |
| Nuclei_Correlation_K_Mito_Lipid | 6.7E-08 | 1.6E-08 | -1.0E+01 |
| Nuclei_Correlation_K_Mito_DNA | 9.3E-09 | 2.6E-09 | -1.2E+01 |
| Nuclei_Correlation_Overlap_Lipid_AGP | 2.0E-06 | 3.3E-07 | -7.7E+00 |
| Nuclei_Correlation_Overlap_DNA_AGP | 2.8E-13 | 2.8E-13 | -2.6E+01 |
| Nuclei_Correlation_Overlap_DNA_Lipid | 2.7E-10 | 1.1E-10 | -1.6E+01 |
| Nuclei_Correlation_Overlap_DNA_Mito | 2.3E-08 | 5.8E-09 | -1.1E+01 |
| Nuclei_Correlation_Overlap_Mito_AGP | 4.8E-07 | 9.0E-08 | 8.7E+00 |
| Nuclei_Granularity_10_AGP | 9.4E-13 | 7.8E-13 | 2.4E+01 |
| Nuclei_Granularity_10_Lipid | 2.8E-09 | 9.0E-10 | 1.3E+01 |
| Nuclei_Granularity_10_DNA | 1.1E-13 | 1.2E-13 | 2.8E+01 |
| Nuclei_Granularity_10_Mito | 9.5E-14 | 1.1E-13 | 2.8E+01 |
| Nuclei_Granularity_2_AGP | 1.7E-16 | 7.9E-16 | -4.5E+01 |
| Nuclei_Granularity_4_Lipid | 4.2E-15 | 9.3E-15 | 3.5E+01 |
| Nuclei_Granularity_5_DNA | 6.8E-20 | 4.3E-18 | 7.8E+01 |
| Nuclei_Granularity_4_Mito | 8.1E-19 | 2.8E-17 | 6.6E+01 |
| Nuclei_Intensity_IntegratedIntensity_AGP | 1.1E-10 | 5.1E-11 | -1.7E+01 |
| Nuclei_Intensity_IntegratedIntensity_Lipid | 5.9E-05 | 7.1E-06 | -5.7E+00 |
| Nuclei_Intensity_IntegratedIntensity_DNA | 3.3E-09 | 1.0E-09 | -1.3E+01 |
| Nuclei_Intensity_IntegratedIntensity_Mito | 4.2E-06 | 6.4E-07 | 7.3E+00 |
| Nuclei_Intensity_IntegratedIntensityEdge_AGP | 1.0E-06 | 1.8E-07 | -8.2E+00 |
| Nuclei_Intensity_IntegratedIntensityEdge_DNA | 5.1E-11 | 2.6E-11 | -1.8E+01 |
| Nuclei_Intensity_IntegratedIntensityEdge_Mito | 1.2E-04 | 1.3E-05 | 5.3E+00 |
| Nuclei_Intensity_LowerQuartileIntensity_AGP | 1.4E-09 | 4.9E-10 | -1.4E+01 |
| Nuclei_Intensity_LowerQuartileIntensity_Lipid | 9.0E-05 | 1.0E-05 | -5.4E+00 |
| Nuclei_Intensity_LowerQuartileIntensity_DNA | 2.2E-11 | 1.2E-11 | -1.9E+01 |
| Nuclei_Intensity_LowerQuartileIntensity_Mito | 2.3E-06 | 3.7E-07 | 7.7E+00 |
| Nuclei_Intensity_MADIntensity_AGP | 3.2E-12 | 2.2E-12 | -2.2E+01 |
| Nuclei_Intensity_MADIntensity_DNA | 3.6E-04 | 3.8E-05 | 4.7E+00 |
| Nuclei_Intensity_MADIntensity_Mito | 9.2E-09 | 2.6E-09 | 1.2E+01 |
| Nuclei_Intensity_MassDisplacement_AGP | 1.9E-11 | 1.1E-11 | 1.9E+01 |
| Nuclei_Intensity_MassDisplacement_Lipid | 2.6E-13 | 2.6E-13 | 2.6E+01 |
| Nuclei_Intensity_MassDisplacement_DNA | 1.7E-18 | 3.2E-17 | 6.2E+01 |
| Nuclei_Intensity_MassDisplacement_Mito | 1.6E-08 | 4.2E-09 | 1.2E+01 |
| Nuclei_Intensity_MaxIntensity_AGP | 2.4E-10 | 1.0E-10 | -1.6E+01 |
| Nuclei_Intensity_MaxIntensity_Mito | 3.6E-03 | 3.2E-04 | 3.5E+00 |
| Nuclei_Intensity_MaxIntensityEdge_AGP | 1.3E-06 | 2.3E-07 | -8.0E+00 |
| Nuclei_Intensity_MaxIntensityEdge_DNA | 4.5E-09 | 1.4E-09 | -1.3E+01 |
| Nuclei_Intensity_MaxIntensityEdge_Mito | 5.6E-03 | 4.9E-04 | 3.3E+00 |
| Nuclei_Intensity_MeanIntensity_AGP | 1.7E-10 | 7.7E-11 | -1.6E+01 |
| Nuclei_Intensity_MeanIntensity_Lipid | 4.3E-04 | 4.5E-05 | -4.6E+00 |
| Nuclei_Intensity_MeanIntensity_DNA | 1.9E-07 | 3.9E-08 | -9.5E+00 |
| Nuclei_Intensity_MeanIntensity_Mito | 1.7E-06 | 2.9E-07 | 7.8E+00 |
| Nuclei_Intensity_MeanIntensityEdge_AGP | 9.9E-07 | 1.8E-07 | -8.2E+00 |
| Nuclei_Intensity_MeanIntensityEdge_DNA | 2.3E-10 | 9.7E-11 | -1.6E+01 |
| Nuclei_Intensity_MeanIntensityEdge_Mito | 1.4E-04 | 1.5E-05 | 5.2E+00 |
| Nuclei_Intensity_MedianIntensity_AGP | 1.0E-10 | 4.8E-11 | -1.7E+01 |
| Nuclei_Intensity_MedianIntensity_Lipid | 5.3E-04 | 5.4E-05 | -4.5E+00 |
| Nuclei_Intensity_MedianIntensity_DNA | 2.5E-09 | 8.2E-10 | -1.3E+01 |
| Nuclei_Intensity_MedianIntensity_Mito | 3.7E-07 | 7.3E-08 | 8.9E+00 |
| Nuclei_Intensity_MinIntensity_AGP | 1.8E-06 | 3.0E-07 | -7.8E+00 |
| Nuclei_Intensity_MinIntensity_Lipid | 5.6E-07 | 1.0E-07 | -8.6E+00 |
| Nuclei_Intensity_MinIntensity_DNA | 2.8E-10 | 1.2E-10 | -1.6E+01 |
| Nuclei_Intensity_MinIntensity_Mito | 5.3E-04 | 5.4E-05 | 4.5E+00 |
| Nuclei_Intensity_MinIntensityEdge_AGP | 3.1E-06 | 5.0E-07 | -7.5E+00 |
| Nuclei_Intensity_MinIntensityEdge_Lipid | 6.9E-07 | 1.3E-07 | -8.5E+00 |
| Nuclei_Intensity_MinIntensityEdge_DNA | 2.9E-10 | 1.2E-10 | -1.6E+01 |
| Nuclei_Intensity_MinIntensityEdge_Mito | 8.8E-03 | 7.6E-04 | 3.0E+00 |
| Nuclei_Intensity_StdIntensity_AGP | 4.6E-11 | 2.4E-11 | -1.8E+01 |
| Nuclei_Intensity_StdIntensity_DNA | 3.5E-03 | 3.2E-04 | 3.5E+00 |
| Nuclei_Intensity_StdIntensity_Mito | 2.6E-05 | 3.3E-06 | 6.1E+00 |
| Nuclei_Intensity_StdIntensityEdge_AGP | 3.1E-06 | 5.0E-07 | -7.5E+00 |
| Nuclei_Intensity_StdIntensityEdge_Lipid | 1.9E-05 | 2.4E-06 | 6.3E+00 |
| Nuclei_Intensity_StdIntensityEdge_DNA | 7.2E-08 | 1.7E-08 | -1.0E+01 |
| Nuclei_Intensity_StdIntensityEdge_Mito | 1.3E-04 | 1.5E-05 | 5.2E+00 |
| Nuclei_Intensity_UpperQuartileIntensity_AGP | 5.1E-11 | 2.6E-11 | -1.8E+01 |
| Nuclei_Intensity_UpperQuartileIntensity_Lipid | 6.4E-04 | 6.5E-05 | -4.4E+00 |
| Nuclei_Intensity_UpperQuartileIntensity_DNA | 5.8E-05 | 7.0E-06 | -5.7E+00 |
| Nuclei_Intensity_UpperQuartileIntensity_Mito | 7.1E-07 | 1.3E-07 | 8.5E+00 |
| Nuclei_Location_Center_Y | 9.8E-04 | 9.6E-05 | 4.2E+00 |
| Nuclei_Location_CenterMassIntensity_Y_AGP | 1.0E-03 | 9.8E-05 | 4.1E+00 |
| Nuclei_Location_CenterMassIntensity_Y_Lipid | 9.9E-04 | 9.7E-05 | 4.1E+00 |
| Nuclei_Location_CenterMassIntensity_Y_DNA | 9.8E-04 | 9.6E-05 | 4.2E+00 |
| Nuclei_Location_CenterMassIntensity_Y_Mito | 9.3E-04 | 9.1E-05 | 4.2E+00 |
| Nuclei_Location_MaxIntensity_Y_AGP | 1.1E-03 | 1.0E-04 | 4.1E+00 |
| Nuclei_Location_MaxIntensity_Y_Lipid | 1.1E-03 | 1.0E-04 | 4.1E+00 |
| Nuclei_Location_MaxIntensity_Y_DNA | 9.3E-04 | 9.2E-05 | 4.2E+00 |
| Nuclei_Location_MaxIntensity_Y_Mito | 5.1E-04 | 5.3E-05 | 4.5E+00 |
| Nuclei_Neighbors_AngleBetweenNeighbors_2 | 2.3E-12 | 1.6E-12 | -2.2E+01 |
| Nuclei_Neighbors_FirstClosestDistance_2 | 3.6E-03 | 3.2E-04 | -3.5E+00 |
| Nuclei_Neighbors_FirstClosestObjectNumber_2 | 5.7E-09 | 1.7E-09 | -1.2E+01 |
| Nuclei_Neighbors_NumberOfNeighbors_2 | 2.0E-11 | 1.1E-11 | 1.9E+01 |
| Nuclei_Neighbors_PercentTouching_2 | 1.0E-11 | 6.4E-12 | 2.0E+01 |
| Nuclei_Neighbors_SecondClosestDistance_2 | 4.8E-04 | 4.9E-05 | 4.5E+00 |
| Nuclei_Neighbors_SecondClosestObjectNumber_2 | 5.6E-09 | 1.7E-09 | -1.2E+01 |
| Nuclei_Number_Object_Number | 6.0E-09 | 1.7E-09 | -1.2E+01 |
| Nuclei_RadialDistribution_FracAtD_AGP_3of4 | 2.2E-13 | 2.3E-13 | -2.7E+01 |
| Nuclei_RadialDistribution_FracAtD_Lipid_1of4 | 6.7E-09 | 1.9E-09 | -1.2E+01 |
| Nuclei_RadialDistribution_FracAtD_DNA_2of4 | 4.4E-14 | 6.0E-14 | 3.0E+01 |
| Nuclei_RadialDistribution_FracAtD_Mito_3of4 | 6.9E-13 | 6.0E-13 | 2.4E+01 |
| Nuclei_RadialDistribution_MeanFrac_AGP_3of4 | 1.7E-13 | 1.8E-13 | -2.7E+01 |
| Nuclei_RadialDistribution_MeanFrac_Lipid_2of4 | 1.6E-06 | 2.7E-07 | -7.9E+00 |
| Nuclei_RadialDistribution_MeanFrac_DNA_2of4 | 1.6E-15 | 4.4E-15 | 3.8E+01 |
| Nuclei_RadialDistribution_MeanFrac_Mito_3of4 | 3.7E-13 | 3.5E-13 | 2.6E+01 |
| Nuclei_RadialDistribution_RadialCV_AGP_3of4 | 4.6E-13 | 4.2E-13 | 2.5E+01 |
| Nuclei_RadialDistribution_RadialCV_Lipid_3of4 | 2.7E-12 | 1.9E-12 | 2.2E+01 |
| Nuclei_RadialDistribution_RadialCV_DNA_3of4 | 3.4E-21 | 6.3E-19 | 9.7E+01 |
| Nuclei_RadialDistribution_RadialCV_Mito_2of4 | 2.5E-15 | 5.9E-15 | 3.7E+01 |
| Nuclei_Texture_AngularSecondMoment_AGP_10_02 | 6.0E-09 | 1.8E-09 | 1.2E+01 |
| Nuclei_Texture_AngularSecondMoment_Lipid_5_03 | 8.4E-04 | 8.4E-05 | 4.2E+00 |
| Nuclei_Texture_AngularSecondMoment_DNA_20_03 | 1.9E-07 | 4.0E-08 | 9.4E+00 |
| Nuclei_Texture_AngularSecondMoment_Mito_5_02 | 4.4E-11 | 2.3E-11 | -1.8E+01 |
| Nuclei_Texture_Contrast_AGP_10_03 | 5.9E-11 | 3.0E-11 | -1.8E+01 |
| Nuclei_Texture_Contrast_Lipid_20_02 | 6.8E-04 | 6.9E-05 | 4.3E+00 |
| Nuclei_Texture_Contrast_DNA_5_01 | 4.6E-05 | 5.7E-06 | 5.8E+00 |
| Nuclei_Texture_Contrast_Mito_5_03 | 9.2E-06 | 1.3E-06 | 6.8E+00 |
| Nuclei_Texture_Correlation_AGP_10_03 | 3.7E-14 | 5.2E-14 | 3.0E+01 |
| Nuclei_Texture_Correlation_Lipid_5_03 | 1.4E-12 | 1.0E-12 | 2.3E+01 |
| Nuclei_Texture_Correlation_DNA_10_00 | 2.1E-10 | 9.1E-11 | -1.6E+01 |
| Nuclei_Texture_Correlation_Mito_5_02 | 7.5E-10 | 2.8E-10 | 1.5E+01 |
| Nuclei_Texture_DifferenceEntropy_AGP_10_00 | 6.5E-12 | 4.2E-12 | -2.1E+01 |
| Nuclei_Texture_DifferenceEntropy_Lipid_20_01 | 1.1E-06 | 1.9E-07 | 8.2E+00 |
| Nuclei_Texture_DifferenceEntropy_DNA_10_03 | 1.4E-06 | 2.4E-07 | -8.0E+00 |
| Nuclei_Texture_DifferenceEntropy_Mito_20_00 | 1.4E-06 | 2.4E-07 | 8.0E+00 |
| Nuclei_Texture_DifferenceVariance_AGP_10_00 | 1.2E-09 | 4.4E-10 | 1.4E+01 |
| Nuclei_Texture_DifferenceVariance_Lipid_20_00 | 6.5E-05 | 7.7E-06 | 5.6E+00 |
| Nuclei_Texture_DifferenceVariance_DNA_20_03 | 4.2E-06 | 6.4E-07 | 7.3E+00 |
| Nuclei_Texture_DifferenceVariance_Mito_10_00 | 1.2E-06 | 2.1E-07 | -8.1E+00 |
| Nuclei_Texture_Entropy_AGP_5_00 | 1.3E-10 | 5.9E-11 | -1.7E+01 |
| Nuclei_Texture_Entropy_Lipid_20_01 | 3.7E-06 | 5.8E-07 | 7.3E+00 |
| Nuclei_Texture_Entropy_DNA_10_03 | 2.0E-06 | 3.3E-07 | -7.7E+00 |
| Nuclei_Texture_Entropy_Mito_5_02 | 3.7E-09 | 1.2E-09 | 1.3E+01 |
| Nuclei_Texture_InfoMeas1_AGP_10_03 | 2.9E-13 | 2.8E-13 | 2.6E+01 |
| Nuclei_Texture_InfoMeas1_Lipid_20_03 | 6.0E-07 | 1.1E-07 | -8.6E+00 |
| Nuclei_Texture_InfoMeas2_DNA_10_03 | 3.1E-09 | 1.0E-09 | -1.3E+01 |
| Nuclei_Texture_InfoMeas2_Mito_5_01 | 3.4E-09 | 1.1E-09 | 1.3E+01 |
| Nuclei_Texture_InverseDifferenceMoment_AGP_10_00 | 7.1E-12 | 4.5E-12 | 2.1E+01 |
| Nuclei_Texture_InverseDifferenceMoment_Lipid_20_00 | 3.1E-06 | 5.0E-07 | 7.5E+00 |
| Nuclei_Texture_InverseDifferenceMoment_DNA_20_03 | 4.9E-10 | 1.9E-10 | 1.5E+01 |
| Nuclei_Texture_InverseDifferenceMoment_Mito_10_00 | 8.9E-10 | 3.3E-10 | -1.4E+01 |
| Nuclei_Texture_SumAverage_AGP_10_00 | 4.3E-11 | 2.2E-11 | -1.8E+01 |
| Nuclei_Texture_SumAverage_Lipid_10_00 | 1.2E-05 | 1.6E-06 | -6.6E+00 |
| Nuclei_Texture_SumAverage_DNA_10_01 | 3.5E-11 | 1.9E-11 | -1.8E+01 |
| Nuclei_Texture_SumAverage_Mito_20_00 | 6.9E-07 | 1.3E-07 | 8.5E+00 |
| Nuclei_Texture_SumEntropy_AGP_5_00 | 2.3E-11 | 1.3E-11 | -1.9E+01 |
| Nuclei_Texture_SumEntropy_Lipid_20_01 | 1.4E-06 | 2.4E-07 | 8.0E+00 |
| Nuclei_Texture_SumEntropy_DNA_10_03 | 1.8E-06 | 3.0E-07 | -7.8E+00 |
| Nuclei_Texture_SumEntropy_Mito_5_02 | 2.3E-08 | 5.9E-09 | 1.1E+01 |
| Nuclei_Texture_SumVariance_AGP_10_03 | 7.6E-11 | 3.7E-11 | -1.7E+01 |
| Nuclei_Texture_SumVariance_Lipid_20_01 | 3.1E-05 | 3.9E-06 | 6.0E+00 |
| Nuclei_Texture_SumVariance_DNA_5_02 | 2.3E-05 | 3.0E-06 | 6.2E+00 |
| Nuclei_Texture_SumVariance_Mito_5_02 | 3.7E-06 | 5.7E-07 | 7.3E+00 |
| Nuclei_Texture_Variance_AGP_10_03 | 6.0E-11 | 3.0E-11 | -1.8E+01 |
| Nuclei_Texture_Variance_Lipid_20_00 | 3.7E-04 | 3.9E-05 | 4.7E+00 |
| Nuclei_Texture_Variance_DNA_5_00 | 4.9E-05 | 6.0E-06 | 5.8E+00 |
| Nuclei_Texture_Variance_Mito_5_02 | 7.1E-06 | 1.0E-06 | 6.9E+00 |

**Supplemental Table 4:**
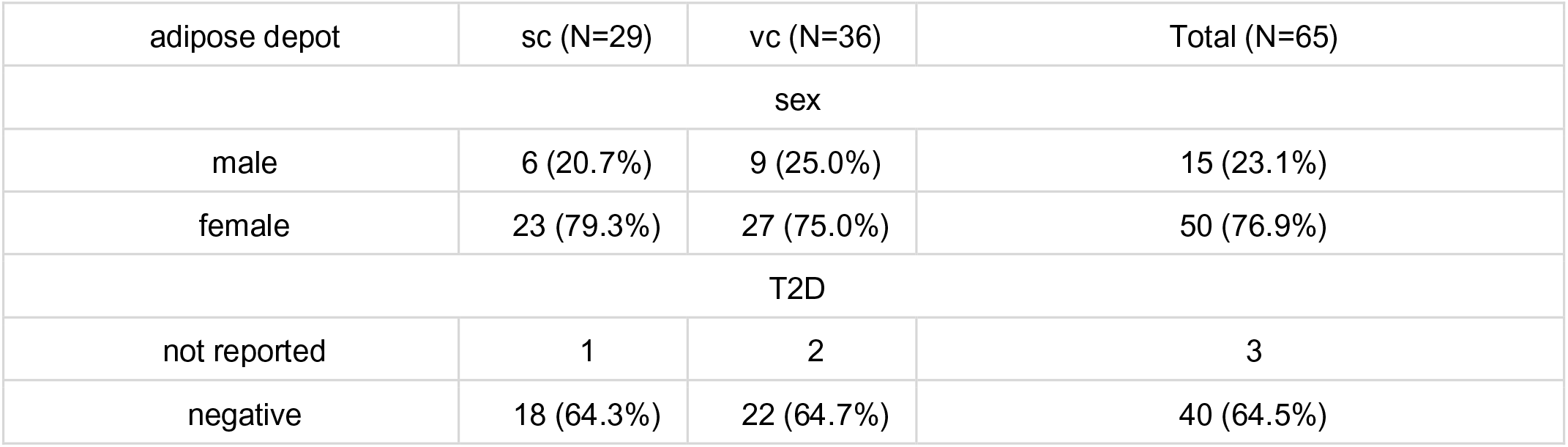
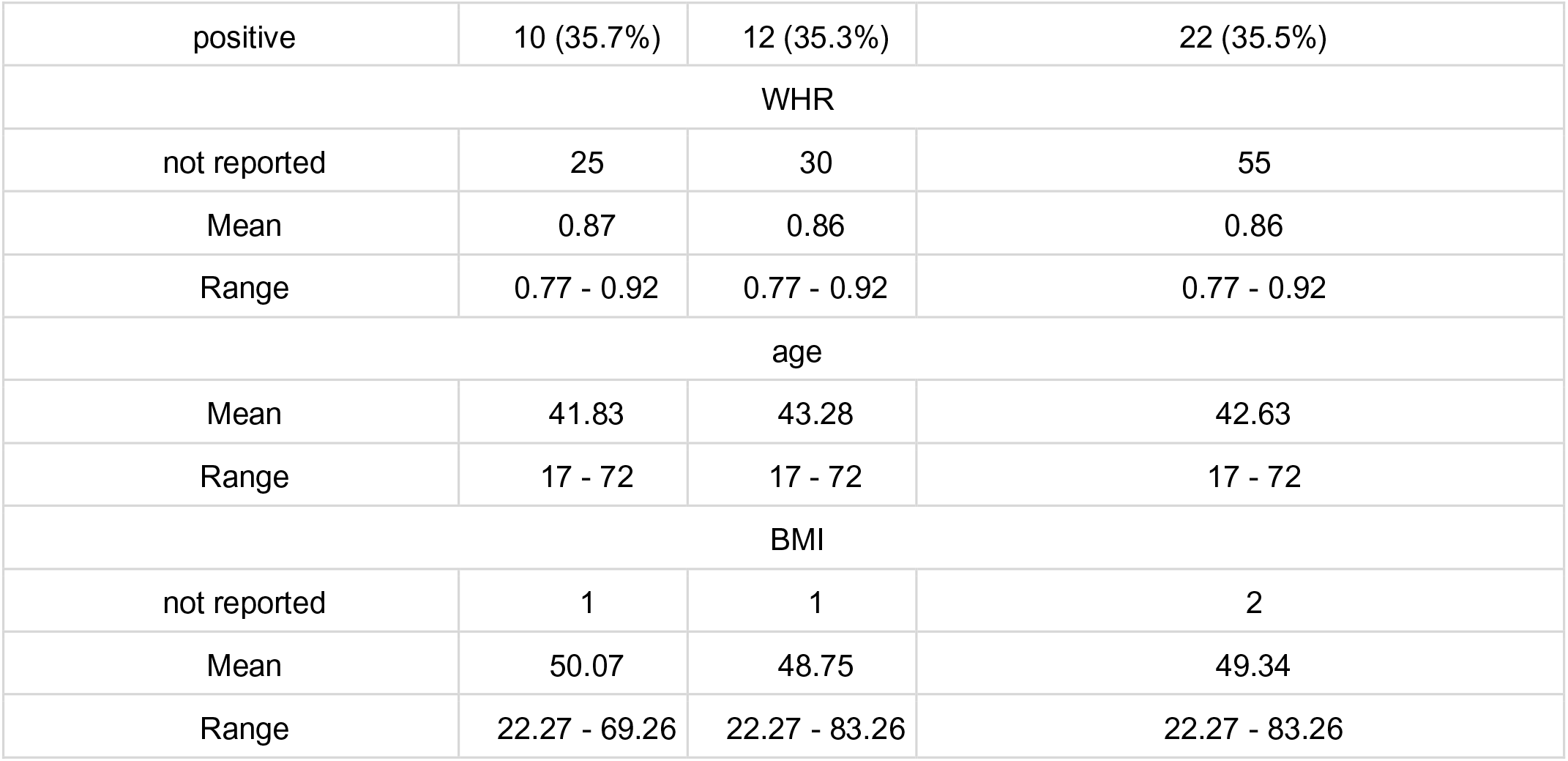
AMSCs summary statistics. Sc, subcutaneous adipose depot; vc, visceral adipose depot; total, total number of samples.

**Supplemental Table 5:**
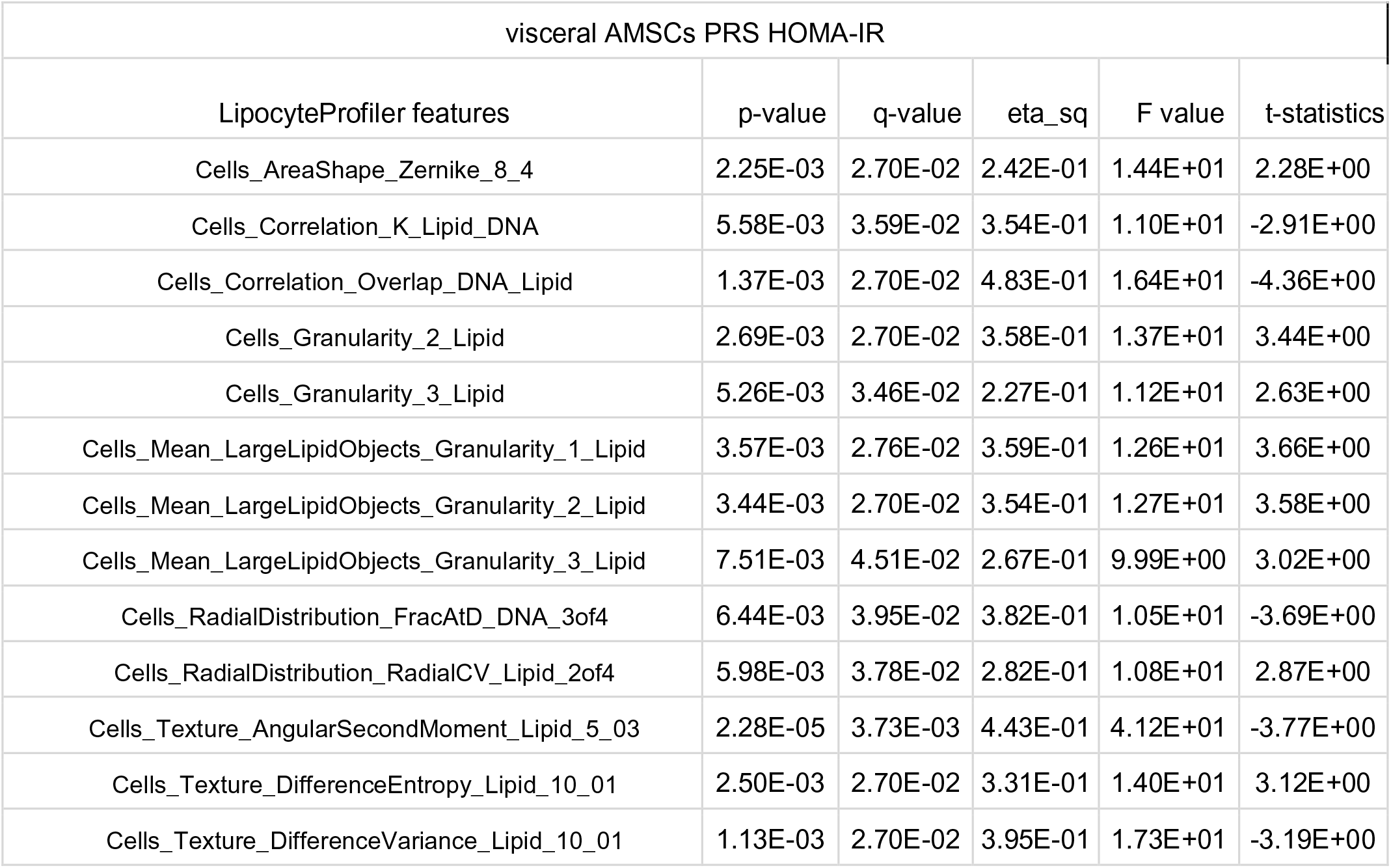
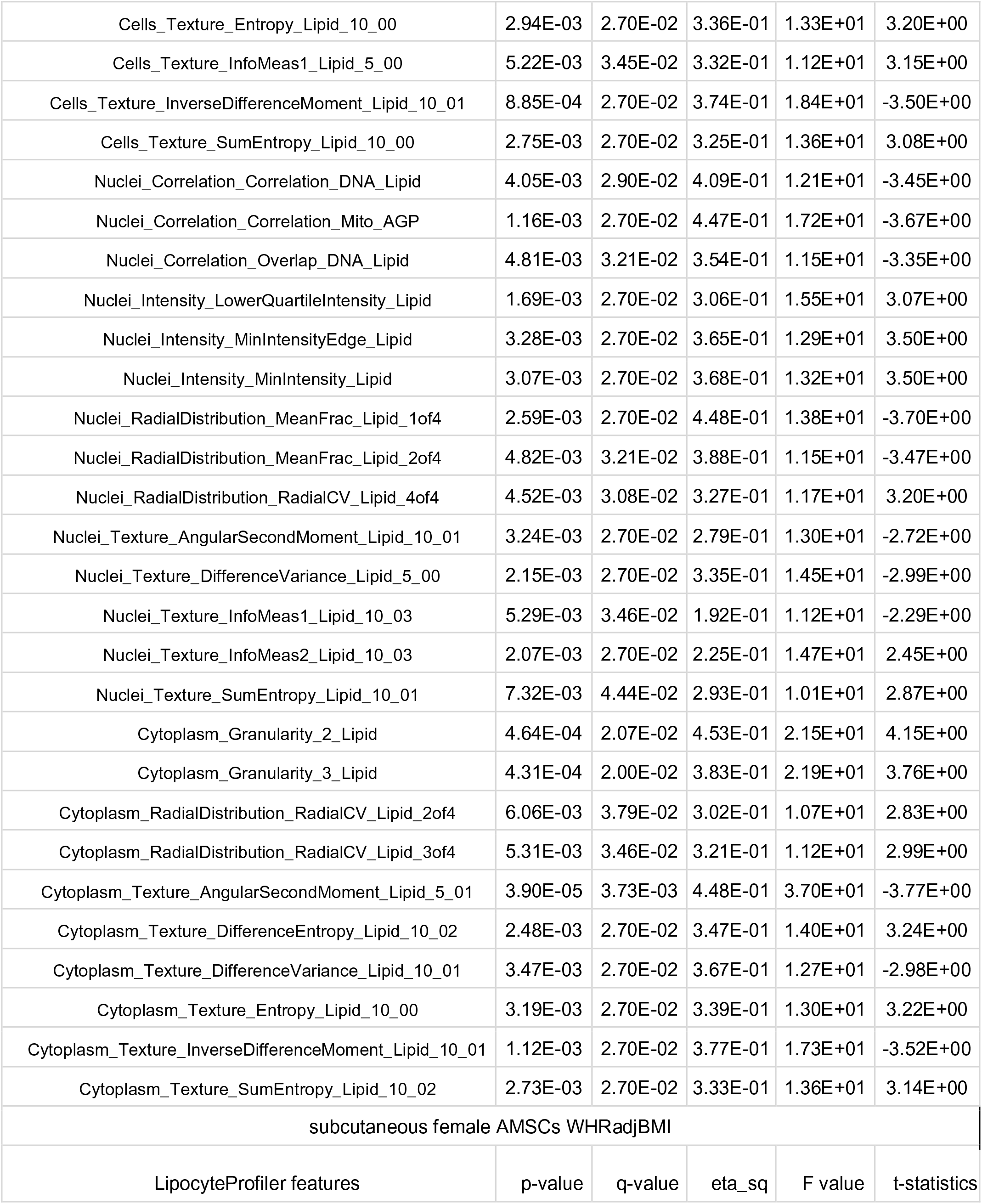
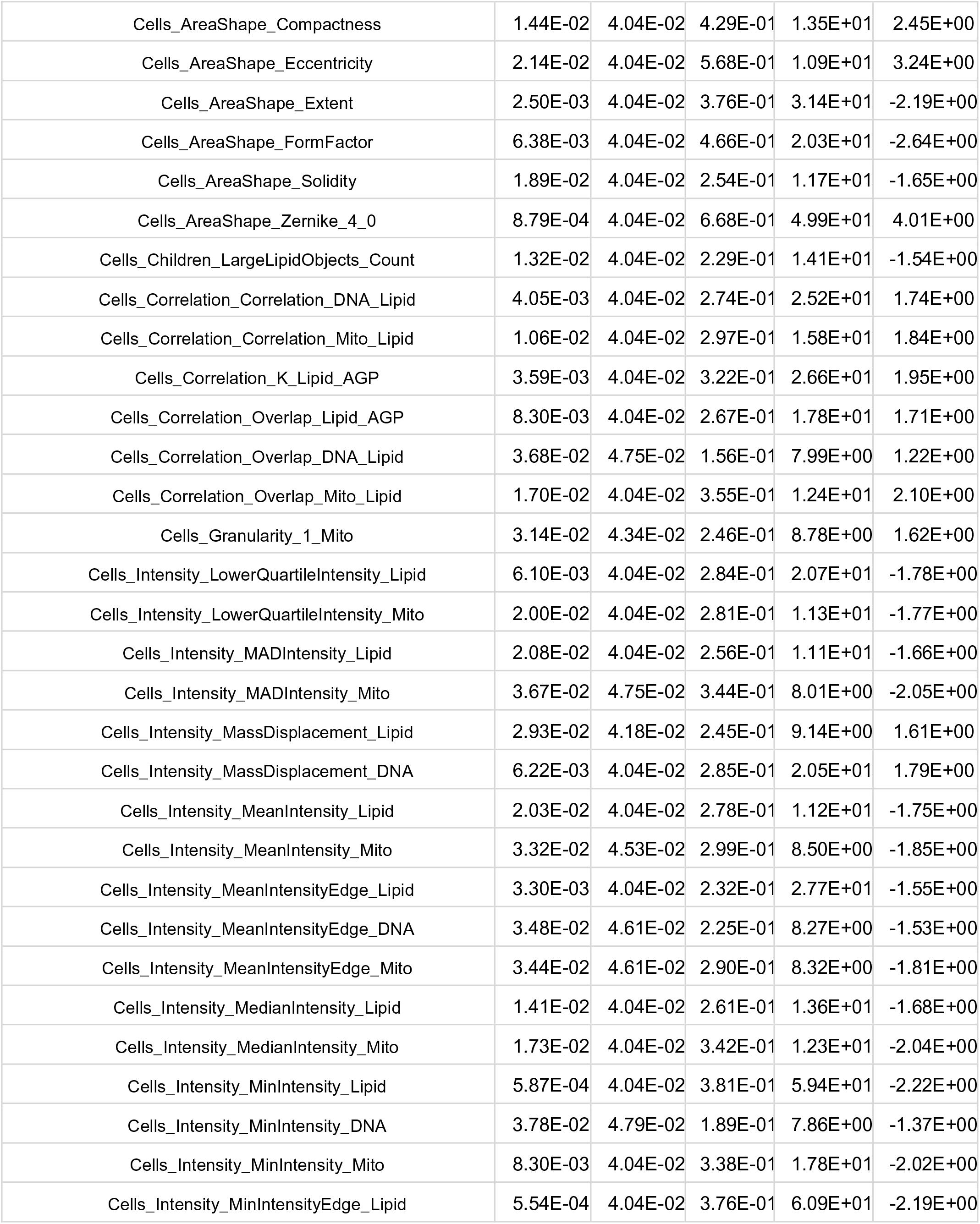
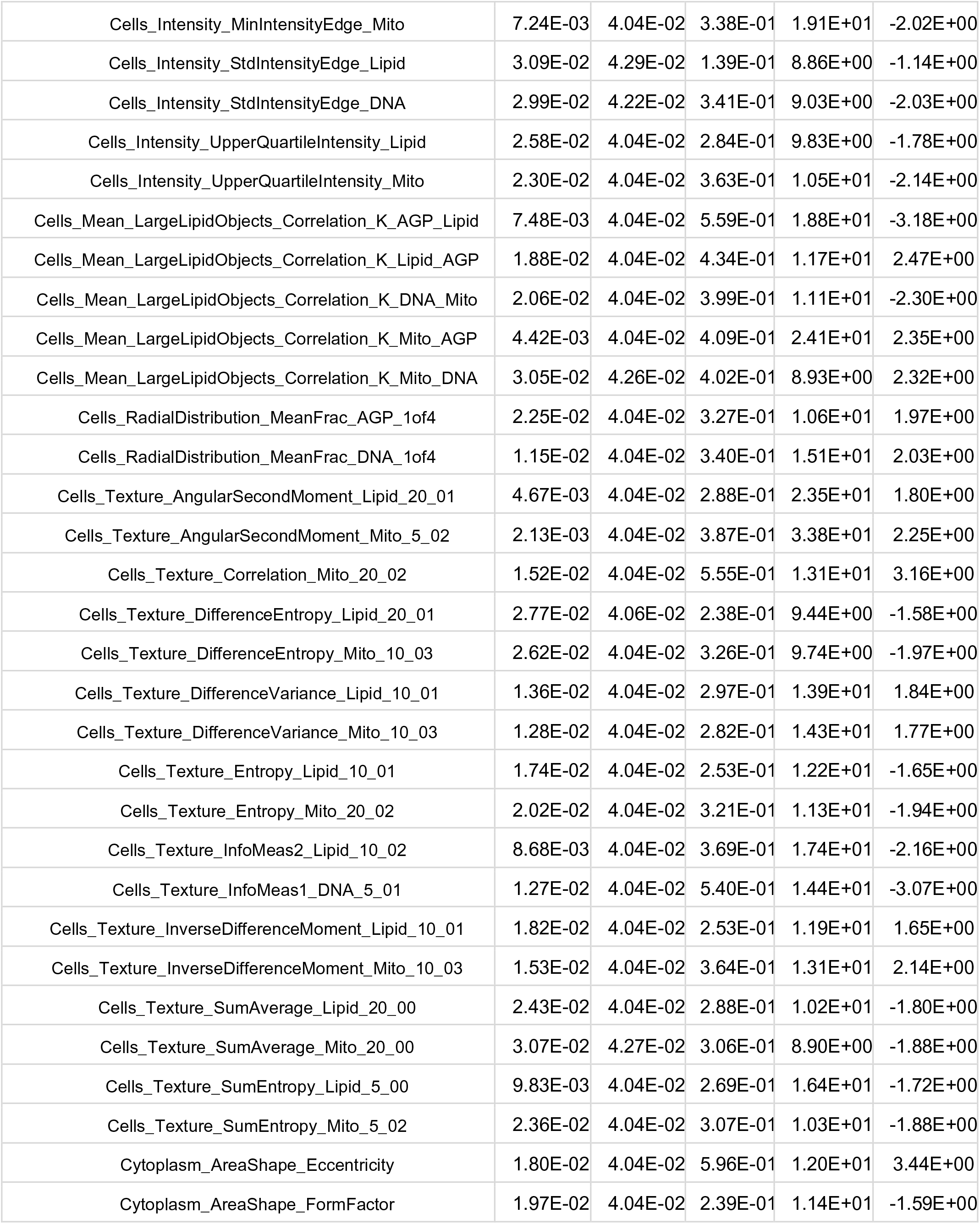
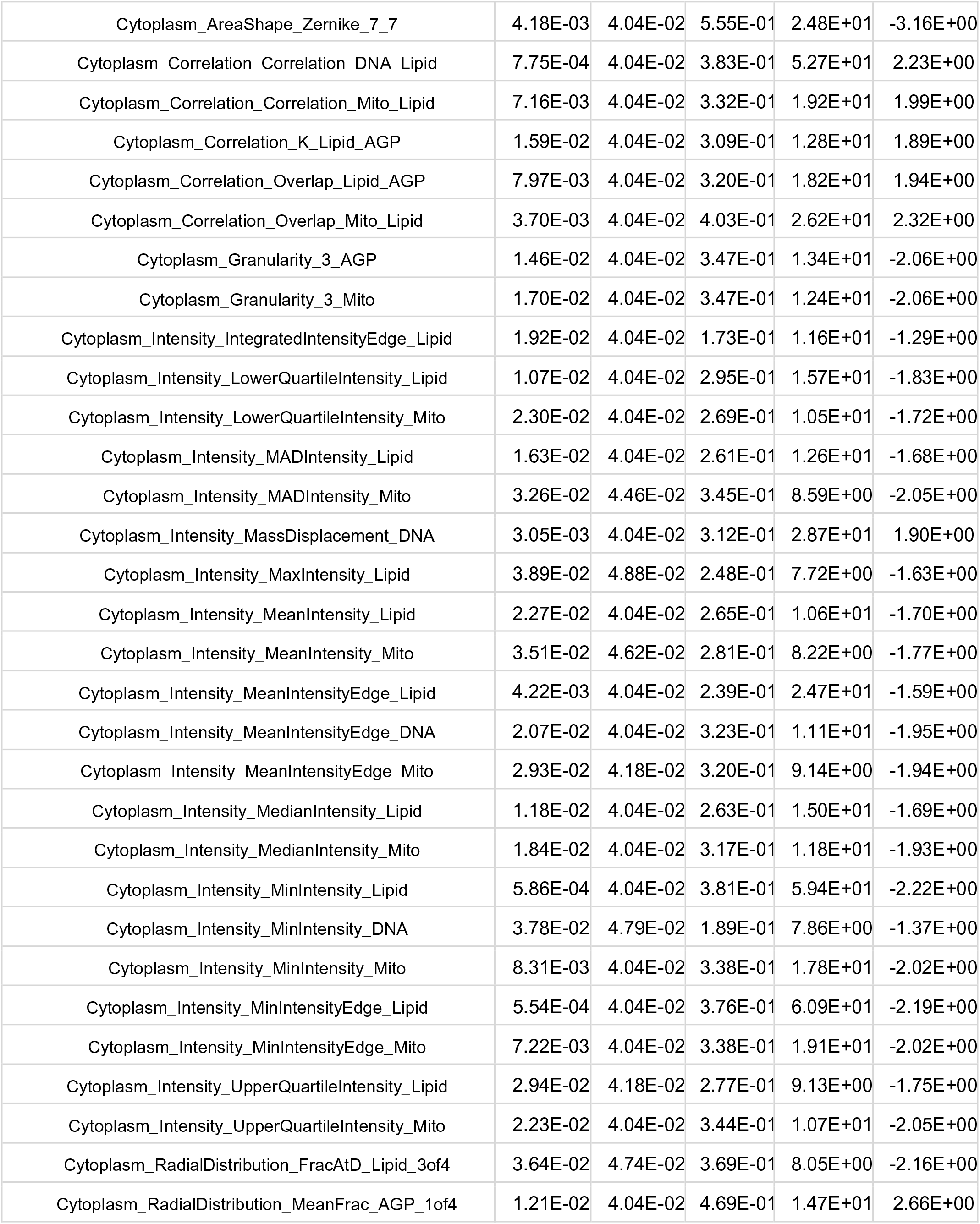
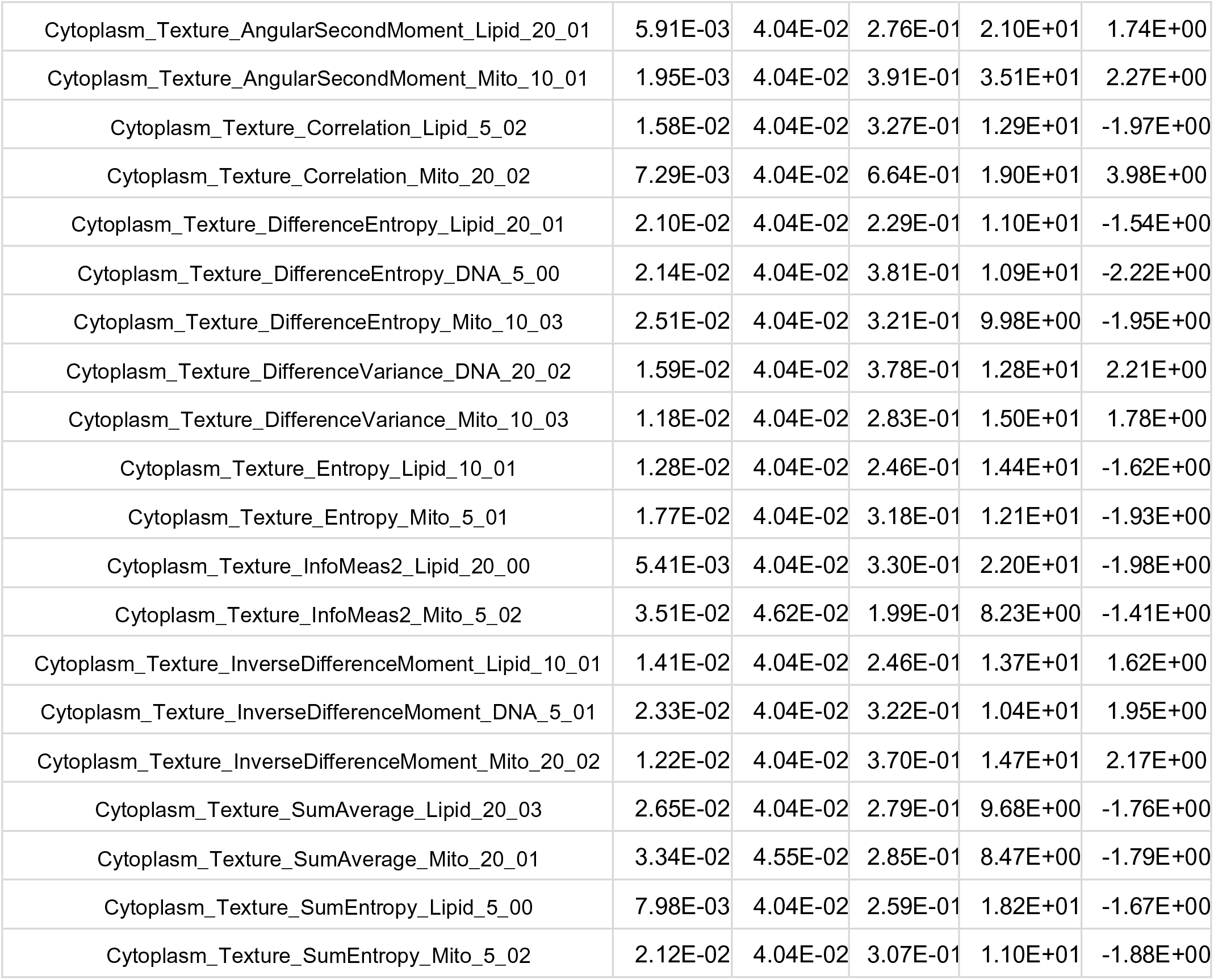
Significant polygenic risk (PRS) effects on LipocyteProfiler for HOMA-IR and WHRadjBMI in terminally differentiated AMSCs. (ANOVA adj. BMI, sex, age, batch, significance level 5%FDR). P-value, p-value of ANOVA, q-value, q-value of ANOVA, FDR; eta_sq, eta square of ANOVA, effect size; F value of ANOVA; t-statistics of t-test.

**Supplemental Table 6:**
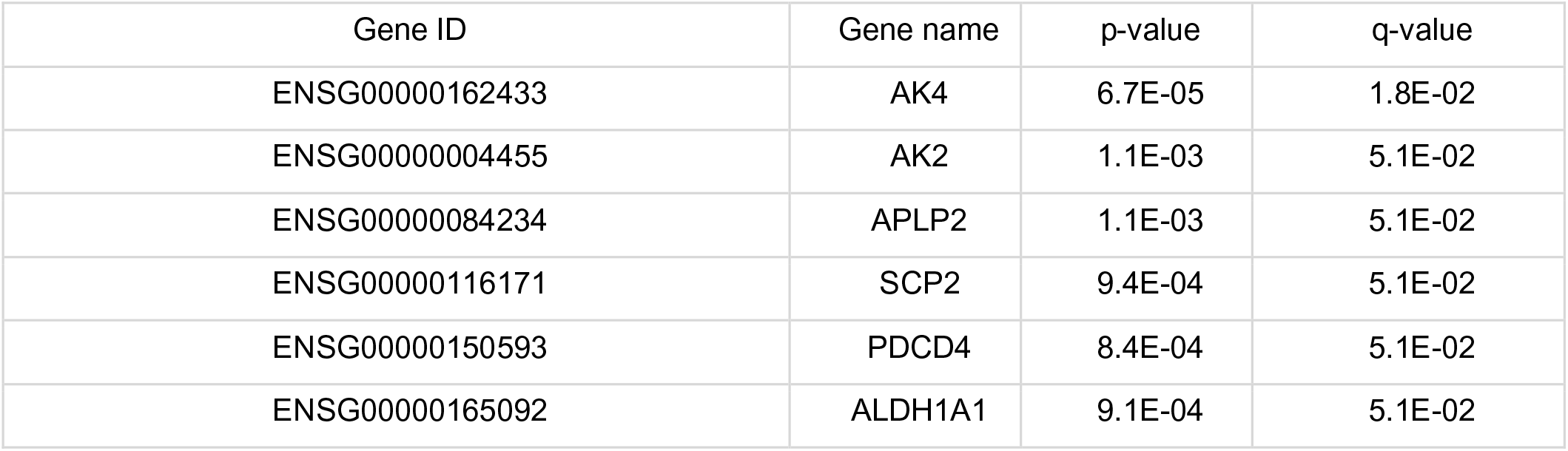
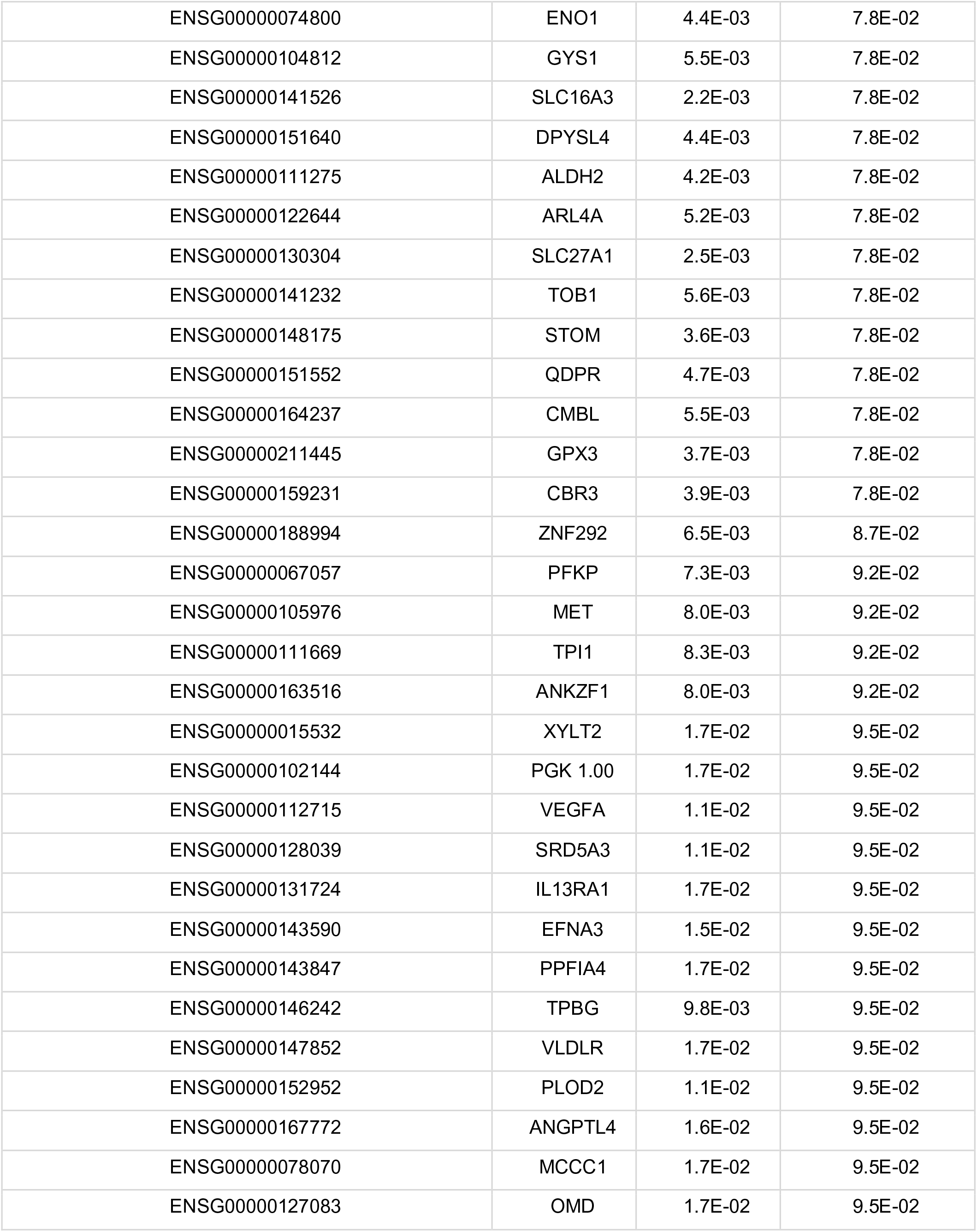
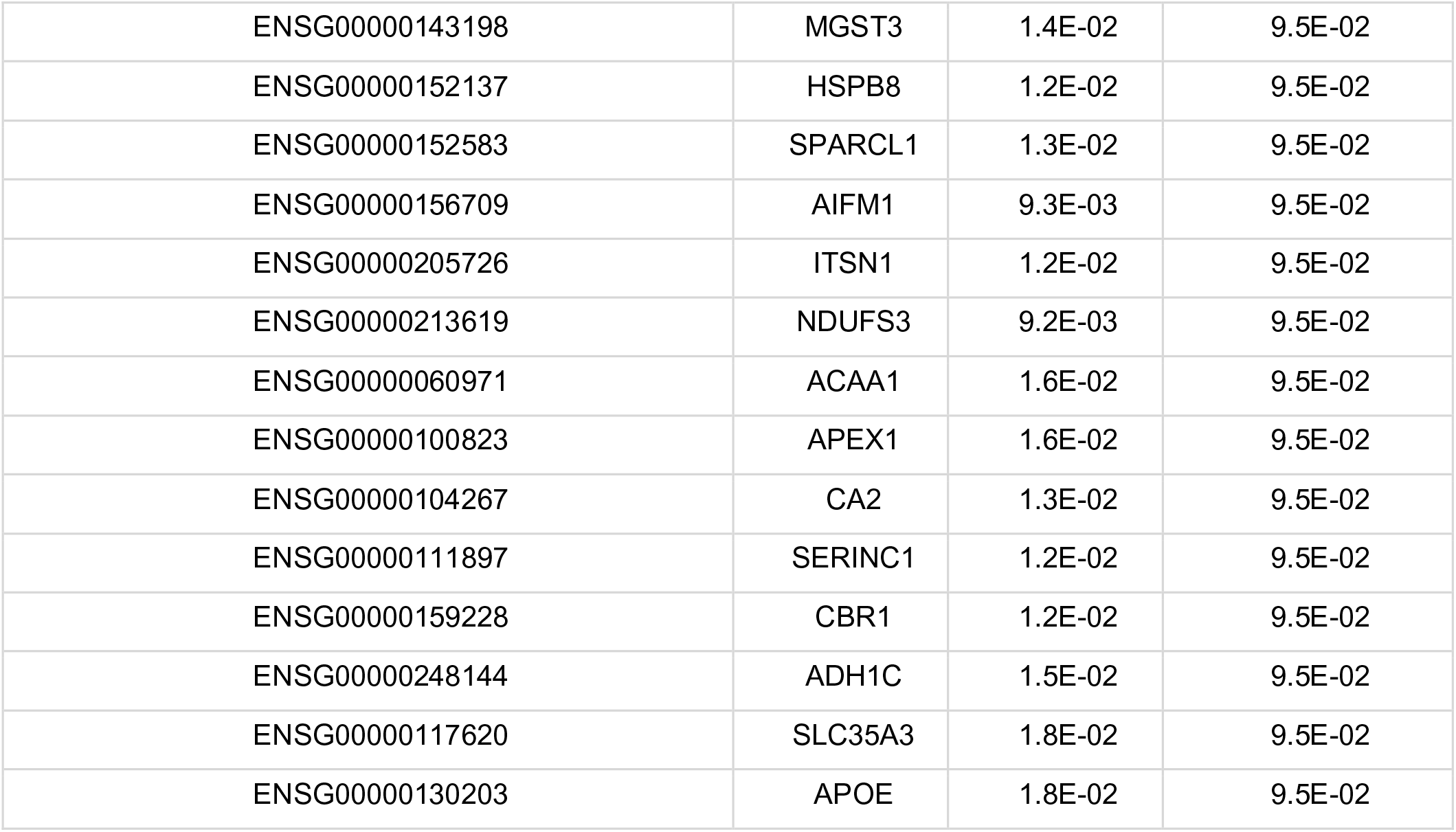
List of significant genes (total 512 genes tested, known to be involved in adipocyte function) for HOMA-IR PRS (linear regression model adj. BMI, sex, age batch, significance level FDR 10%). Gene ID, Ensembl gene identification number; Gene name, gene name; p-value, p-value of linear regression; q-value, q-value of linear regression.

**Supplemental Table 7:**
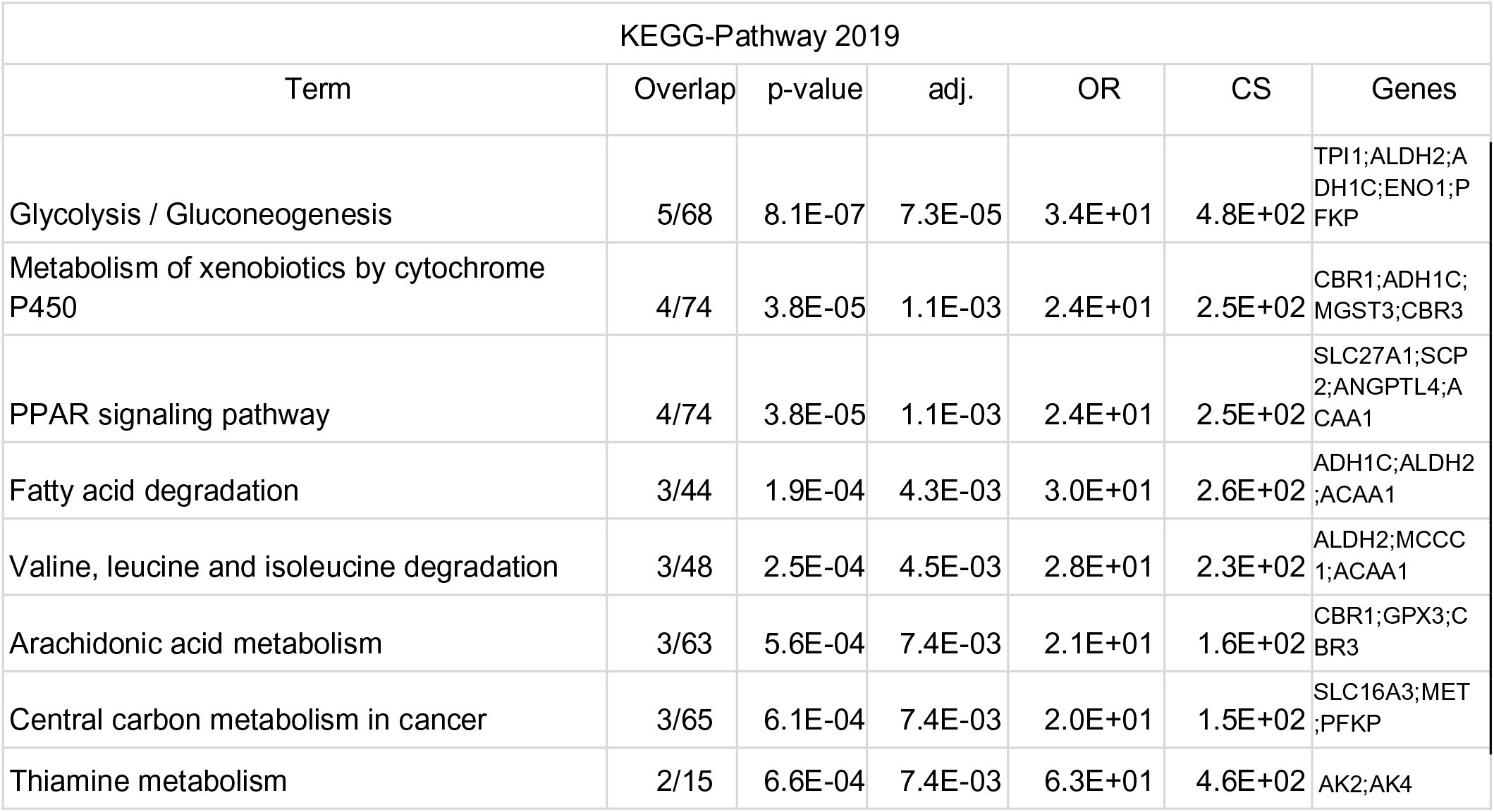
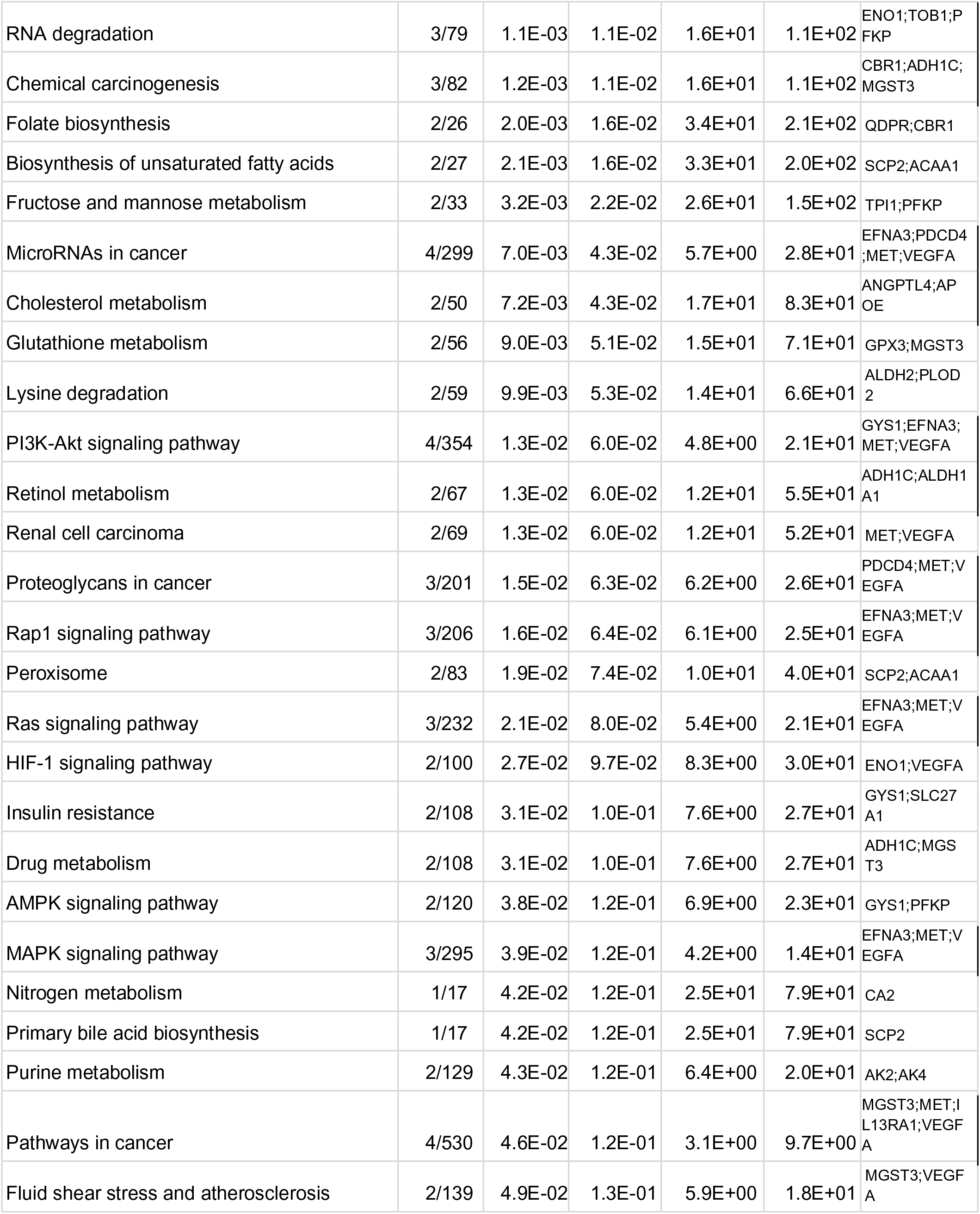
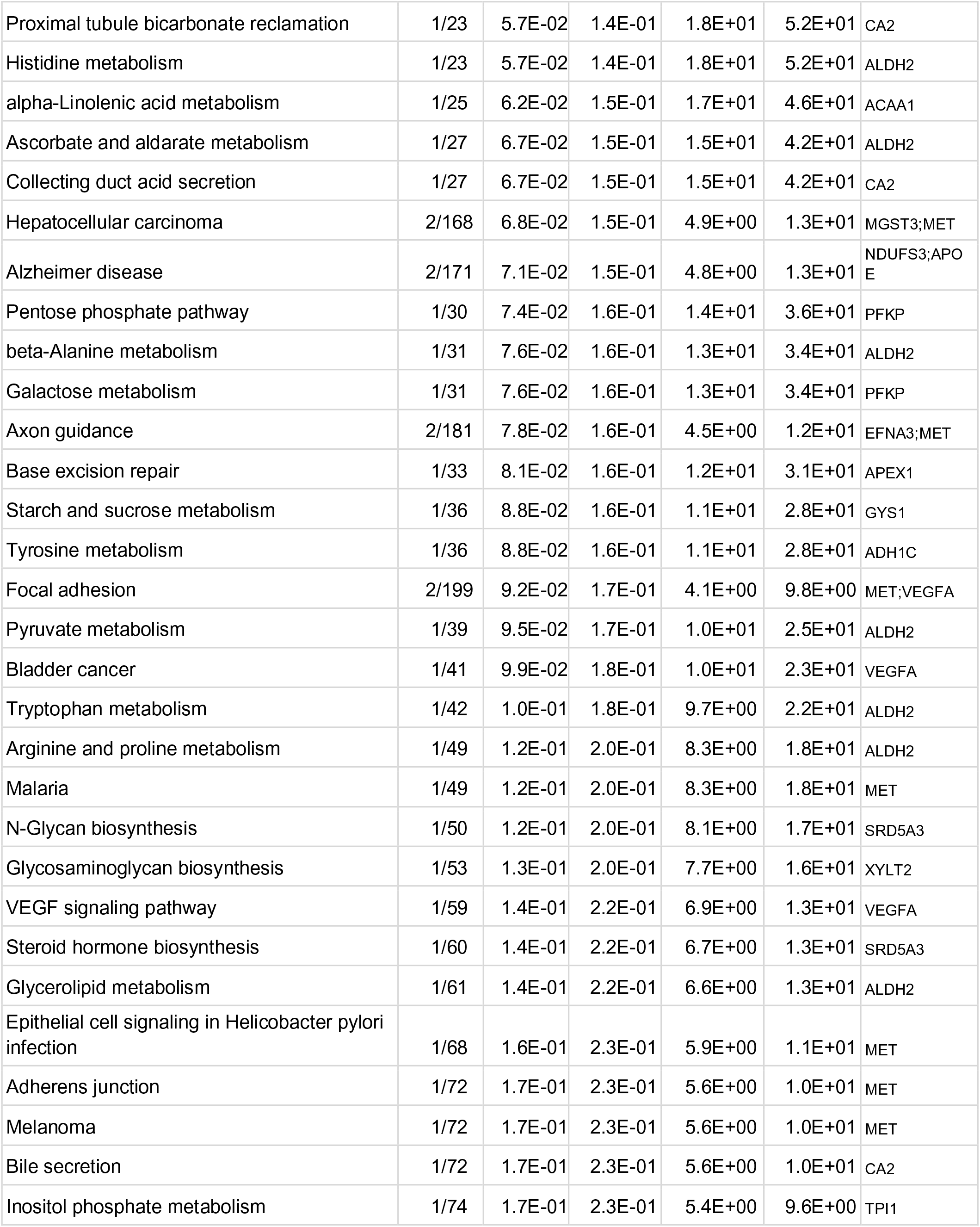
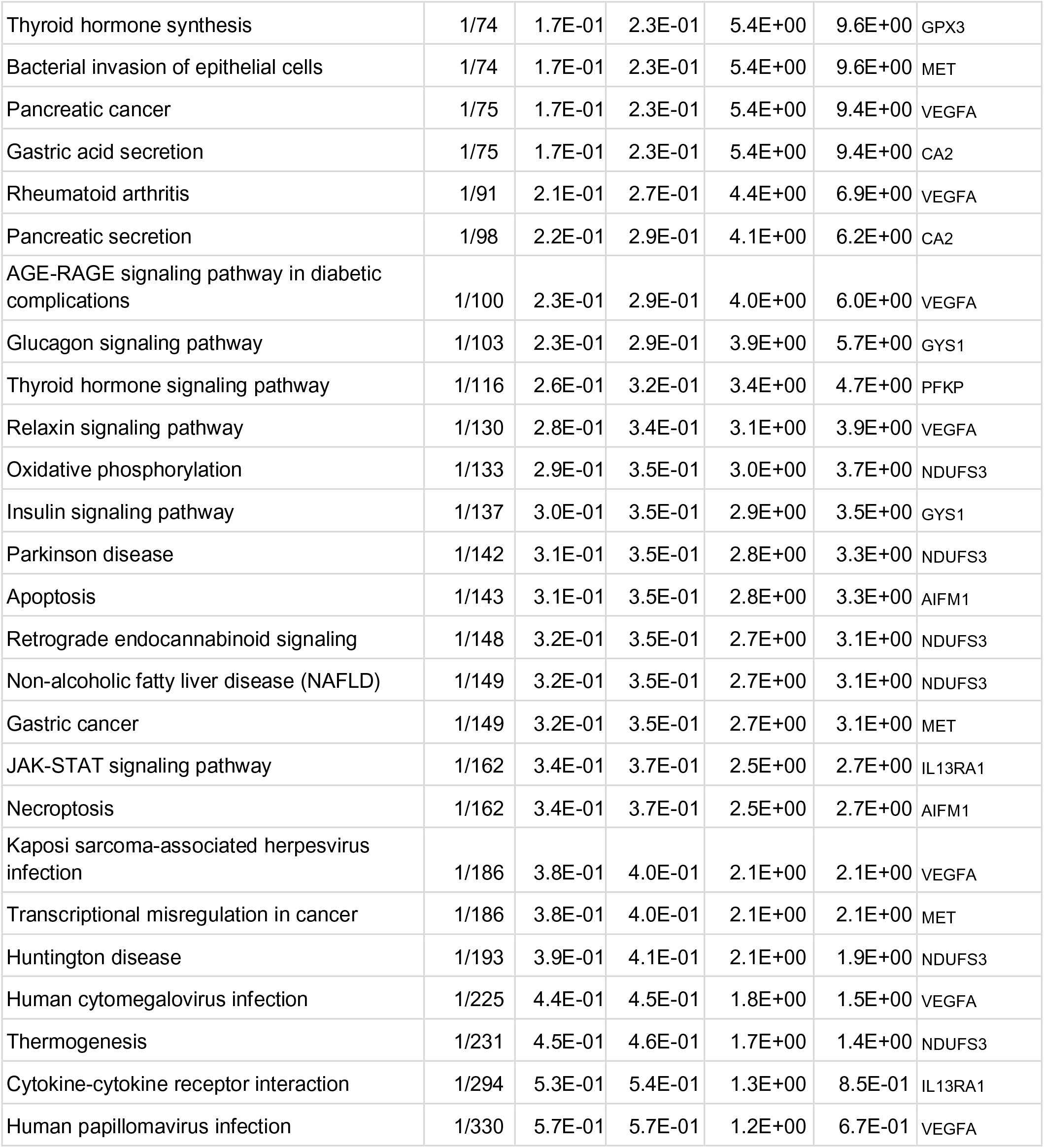
List of KEGG pathways enriched among significantly associated genes with HOMA-IR polygenic risk. Term, which pathway; Overlap, number of genes that overlap and total genes; P-value, enrichment p-value; Adjusted P-value, Q-value; Odds Ratio, enrichment; Combined Score, approximation of overall association (-log10(P) * log(Odds)), Genes, genes in the pathway which are associated with HOMA-IR PRS

**Supplemental Table 8:** Significant polygenic effects on LP profiles for lipodystrophy-like phenotype in terminally differentiated subcutaneous and visceral AMSCs. (linear regression model adj. BMI, sex, age, batch, significance level 5%). P-value, p-value of linear regression; q-value, q-value of linear regression, estimate, estimate of linear regression model.

| subcutaneous AMSCs day14 |  |  |  |
| --- | --- | --- | --- |
| LipocyteProfiler features | pvalue | q-value | estimate |
| Cells_Correlation_Correlation_DNA_AGP | 3.92E-03 | 3.93E-02 | -47.30 |
| Cells_Correlation_Correlation_Mito_AGP | 1.28E-03 | 2.99E-02 | -64.05 |
| Cells_Correlation_Overlap_Mito_AGP | 2.04E-03 | 3.30E-02 | -33.75 |
| Cells_Granularity_14_AGP | 1.88E-03 | 3.30E-02 | 6.32 |
| Cells_Granularity_14_Lipid | 3.60E-03 | 3.93E-02 | 2.14 |
| Cells_Granularity_15_AGP | 1.10E-03 | 2.99E-02 | 12.43 |
| Cells_Granularity_15_Mito | 6.34E-03 | 4.09E-02 | 12.24 |
| Cells_Granularity_16_AGP | 6.36E-03 | 4.09E-02 | 12.59 |
| Cells_Granularity_16_Lipid | 4.22E-03 | 3.93E-02 | -1.55 |
| Cells_Granularity_7_AGP | 5.22E-03 | 3.93E-02 | -11.80 |
| Cells_Intensity_IntegratedIntensity_Mito | 6.31E-03 | 4.09E-02 | 29.51 |
| Cells_Intensity_IntegratedIntensityEdge_Lipid | 1.21E-02 | 4.67E-02 | 2.15 |
| Cells_Intensity_IntegratedIntensityEdge_DNA | 7.80E-03 | 4.33E-02 | 27.05 |
| Cells_Intensity_IntegratedIntensityEdge_Mito | 9.32E-04 | 2.93E-02 | 50.58 |
| Cells_Intensity_LowerQuartileIntensity_Lipid | 4.76E-03 | 3.93E-02 | 5.65 |
| Cells_Intensity_LowerQuartileIntensity_Mito | 8.02E-04 | 2.71E-02 | 38.10 |
| Cells_Intensity_MassDisplacement_Lipid | 1.29E-02 | 4.80E-02 | -10.06 |
| Cells_Intensity_MaxIntensityEdge_Mito | 5.04E-03 | 3.93E-02 | 36.28 |
| Cells_Intensity_MeanIntensity_Mito | 7.54E-03 | 4.33E-02 | 28.08 |
| Cells_Intensity_MeanIntensityEdge_DNA | 7.40E-03 | 4.33E-02 | 15.09 |
| Cells_Intensity_MeanIntensityEdge_Mito | 8.24E-04 | 2.71E-02 | 44.58 |
| Cells_Intensity_MedianIntensity_Mito | 4.08E-03 | 3.93E-02 | 25.75 |
| Cells_Intensity_MinIntensity_Mito | 3.75E-03 | 3.93E-02 | 29.43 |
| Cells_Intensity_MinIntensityEdge_Mito | 2.81E-03 | 3.68E-02 | 30.60 |
| Cells_Intensity_StdIntensityEdge_Mito | 8.63E-03 | 4.33E-02 | 28.68 |
| Cells_Mean_LargeLipidObjects_Correlation_Correlation_DNA_AGP | 1.17E-02 | 4.67E-02 | -23.71 |
| Cells_Mean_LargeLipidObjects_Correlation_Correlation_Mito_AGP | 9.87E-03 | 4.33E-02 | -33.92 |
| Cells_Neighbors_PercentTouching_10 | 1.12E-02 | 4.55E-02 | 28.35 |
| Cells_Neighbors_PercentTouching_Adjacent | 3.38E-03 | 3.93E-02 | 23.72 |
| Cells_RadialDistribution_FracAtD_AGP_2of4 | 1.12E-02 | 4.55E-02 | -16.34 |
| Cells_RadialDistribution_FracAtD_AGP_4of4 | 9.12E-03 | 4.33E-02 | 19.39 |
| Cells_RadialDistribution_FracAtD_Lipid_4of4 | 7.92E-03 | 4.33E-02 | 20.69 |
| Cells_RadialDistribution_MeanFrac_AGP_3of4 | 5.75E-03 | 4.06E-02 | -12.30 |
| Cells_RadialDistribution_MeanFrac_AGP_4of4 | 6.06E-03 | 4.09E-02 | 19.50 |
| Cells_RadialDistribution_MeanFrac_Lipid_3of4 | 1.22E-02 | 4.67E-02 | -19.11 |
| Cells_RadialDistribution_MeanFrac_Lipid_4of4 | 1.03E-02 | 4.41E-02 | 19.84 |
| Cells_RadialDistribution_MeanFrac_Mito_3of4 | 1.34E-02 | 4.91E-02 | -13.26 |
| Cells_RadialDistribution_RadialCV_Lipid_4of4 | 5.30E-03 | 3.93E-02 | -40.29 |
| Cells_Texture_Contrast_Mito_5_03 | 9.54E-03 | 4.33E-02 | 48.68 |
| Cells_Texture_Correlation_AGP_5_03 | 3.75E-03 | 3.93E-02 | -40.89 |
| Cells_Texture_InfoMeas1_Mito_20_01 | 6.83E-05 | 1.15E-02 | -27.70 |
| Cells_Texture_InfoMeas2_AGP_5_02 | 1.18E-02 | 4.67E-02 | -40.71 |
| Cells_Texture_InfoMeas2_Mito_20_00 | 1.21E-03 | 2.99E-02 | 22.90 |
| Cells_Texture_SumAverage_Mito_5_02 | 8.94E-03 | 4.33E-02 | 27.13 |
| Cytoplasm_Correlation_Correlation_DNA_AGP | 2.77E-03 | 3.68E-02 | -56.82 |
| Cytoplasm_Correlation_Correlation_Mito_AGP | 1.17E-03 | 2.99E-02 | -62.74 |
| Cytoplasm_Correlation_Overlap_Mito_AGP | 3.96E-03 | 3.93E-02 | -33.97 |
| Cytoplasm_Granularity_14_AGP | 2.05E-03 | 3.30E-02 | 6.49 |
| Cytoplasm_Granularity_14_Lipid | 3.49E-03 | 3.93E-02 | 2.37 |
| Cytoplasm_Granularity_15_AGP | 1.21E-03 | 2.99E-02 | 12.13 |
| Cytoplasm_Granularity_15_Mito | 7.16E-03 | 4.33E-02 | 11.90 |
| Cytoplasm_Granularity_16_AGP | 6.80E-03 | 4.24E-02 | 12.17 |
| Cytoplasm_Granularity_16_Lipid | 3.76E-03 | 3.93E-02 | -1.35 |
| Cytoplasm_Granularity_7_AGP | 8.87E-03 | 4.33E-02 | -9.04 |
| Cytoplasm_Intensity_IntegratedIntensity_Mito | 4.36E-03 | 3.93E-02 | 31.81 |
| Cytoplasm_Intensity_IntegratedIntensityEdge_DNA | 5.89E-03 | 4.08E-02 | 30.95 |
| Cytoplasm_Intensity_IntegratedIntensityEdge_Mito | 2.28E-03 | 3.36E-02 | 42.61 |
| Cytoplasm_Intensity_LowerQuartileIntensity_Lipid | 4.45E-03 | 3.93E-02 | 10.24 |
| Cytoplasm_Intensity_LowerQuartileIntensity_DNA | 9.86E-03 | 4.33E-02 | 22.46 |
| Cytoplasm_Intensity_LowerQuartileIntensity_Mito | 7.02E-04 | 2.62E-02 | 41.23 |
| Cytoplasm_Intensity_MassDisplacement_Lipid | 1.65E-03 | 3.30E-02 | -14.52 |
| Cytoplasm_Intensity_MeanIntensity_Mito | 4.67E-03 | 3.93E-02 | 32.24 |
| Cytoplasm_Intensity_MeanIntensityEdge_DNA | 8.86E-03 | 4.33E-02 | 12.29 |
| Cytoplasm_Intensity_MeanIntensityEdge_Mito | 2.17E-03 | 3.35E-02 | 37.67 |
| Cytoplasm_Intensity_MedianIntensity_Mito | 2.73E-03 | 3.68E-02 | 29.73 |
| Cytoplasm_Intensity_MinIntensity_Mito | 3.75E-03 | 3.93E-02 | 29.43 |
| Cytoplasm_Intensity_MinIntensityEdge_Mito | 2.82E-03 | 3.68E-02 | 30.61 |
| Cytoplasm_Intensity_UpperQuartileIntensity_Mito | 9.86E-03 | 4.33E-02 | 22.71 |
| Cytoplasm_RadialDistribution_MeanFrac_Mito_2of4 | 1.27E-02 | 4.79E-02 | -8.68 |
| Cytoplasm_RadialDistribution_MeanFrac_Mito_3of4 | 3.87E-03 | 3.93E-02 | -9.92 |
| Cytoplasm_RadialDistribution_MeanFrac_Mito_4of4 | 6.09E-03 | 4.09E-02 | 11.29 |
| Cytoplasm_Texture_Contrast_Mito_5_03 | 5.01E-03 | 3.93E-02 | 57.18 |
| Cytoplasm_Texture_Correlation_AGP_5_03 | 1.30E-02 | 4.80E-02 | -29.05 |
| Cytoplasm_Texture_InfoMeas1_Mito_20_00 | 5.23E-05 | 1.15E-02 | -28.24 |
| Cytoplasm_Texture_InfoMeas2_Mito_20_00 | 3.95E-04 | 2.48E-02 | 24.06 |
| Cytoplasm_Texture_SumAverage_Mito_5_00 | 4.91E-03 | 3.93E-02 | 31.98 |
| Cytoplasm_Texture_SumVariance_Mito_5_02 | 9.83E-03 | 4.33E-02 | 56.60 |
| Cytoplasm_Texture_Variance_Mito_5_02 | 9.51E-03 | 4.33E-02 | 56.87 |
| visceral AMSCs |  |  |  |
| LipocyteProfiler features | pvalue | q-value | estimate |
| Cells_Texture_InfoMeas2_Lipid_20_01 | 3.96E-04 | 3.43E-02 | -25.55 |
| Cytoplasm_Texture_InfoMeas2_Lipid_20_01 | 3.55E-04 | 3.43E-02 | -28.47 |

**Supplemental Table 9:**
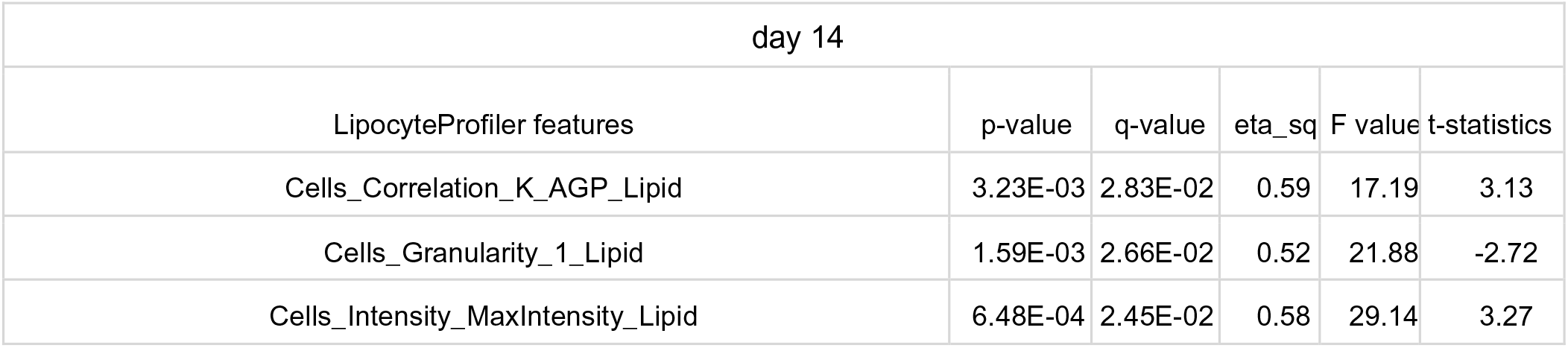
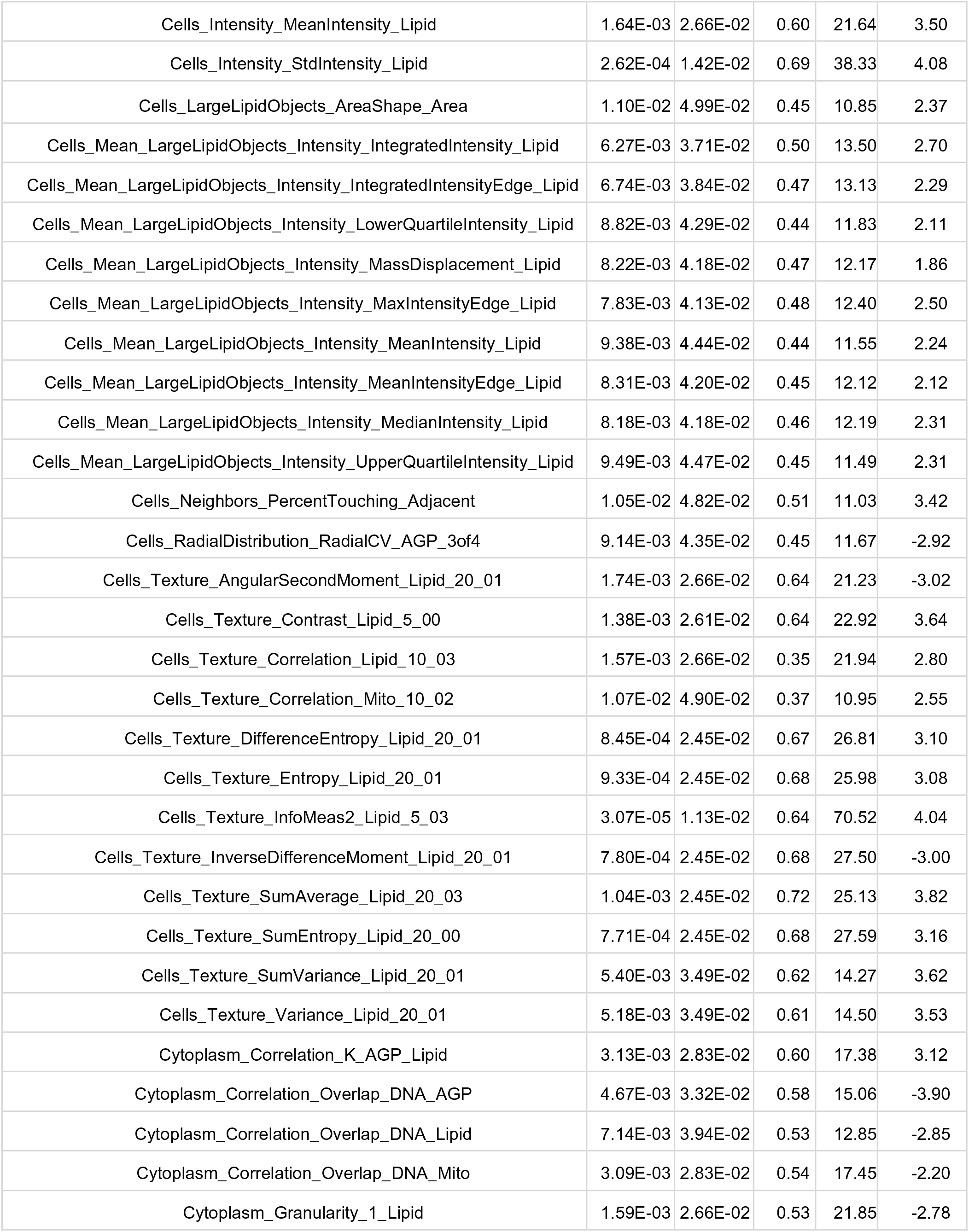
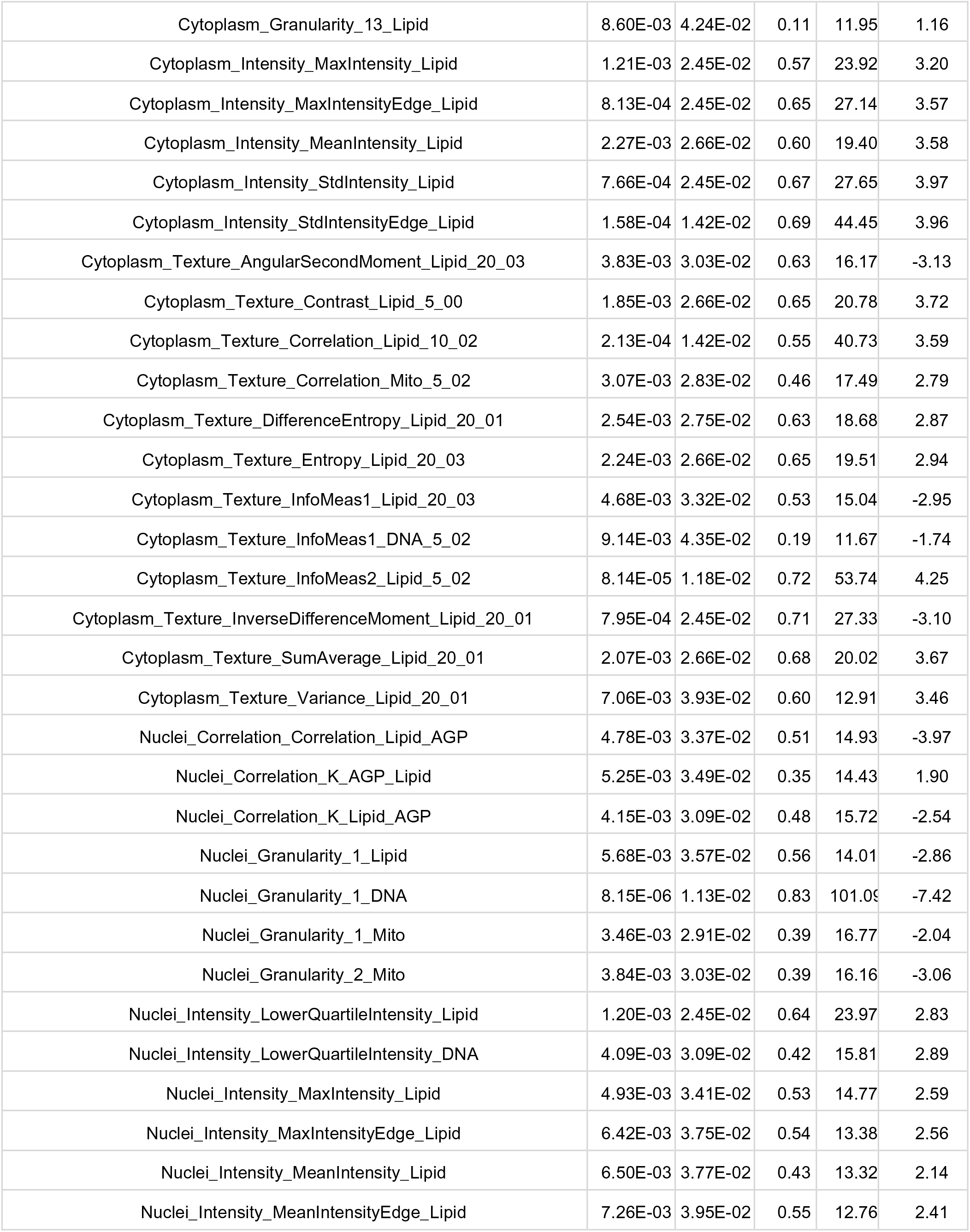
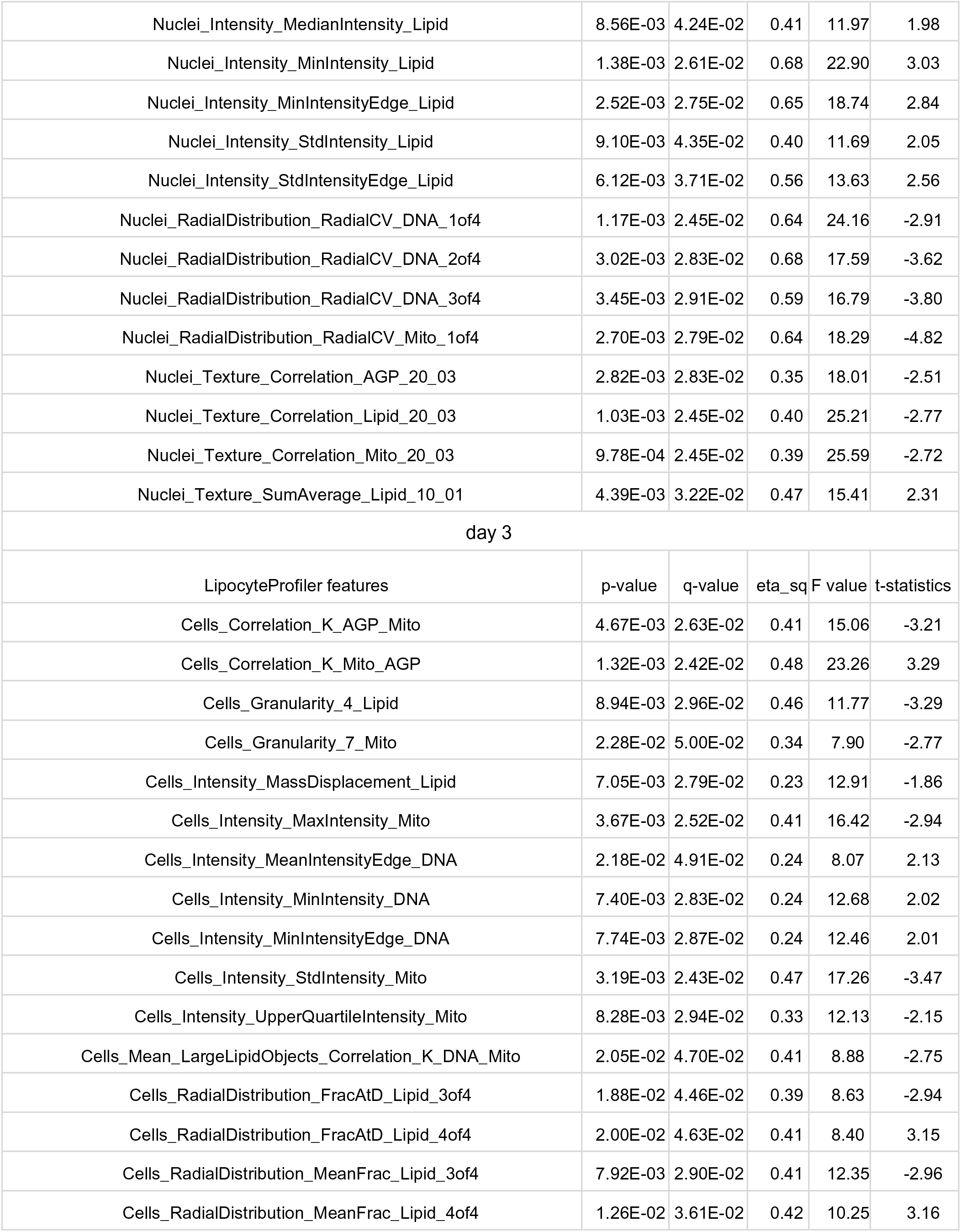
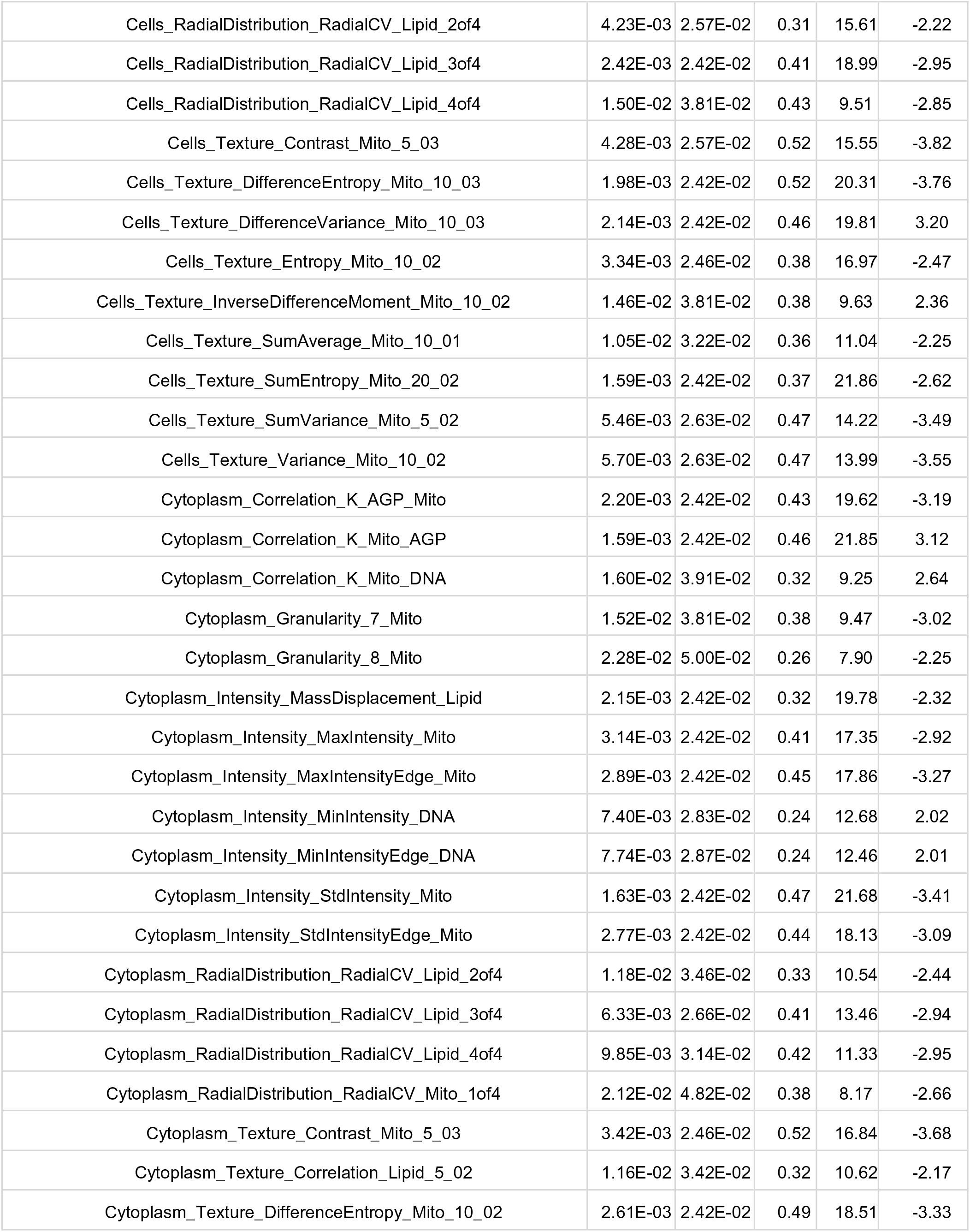
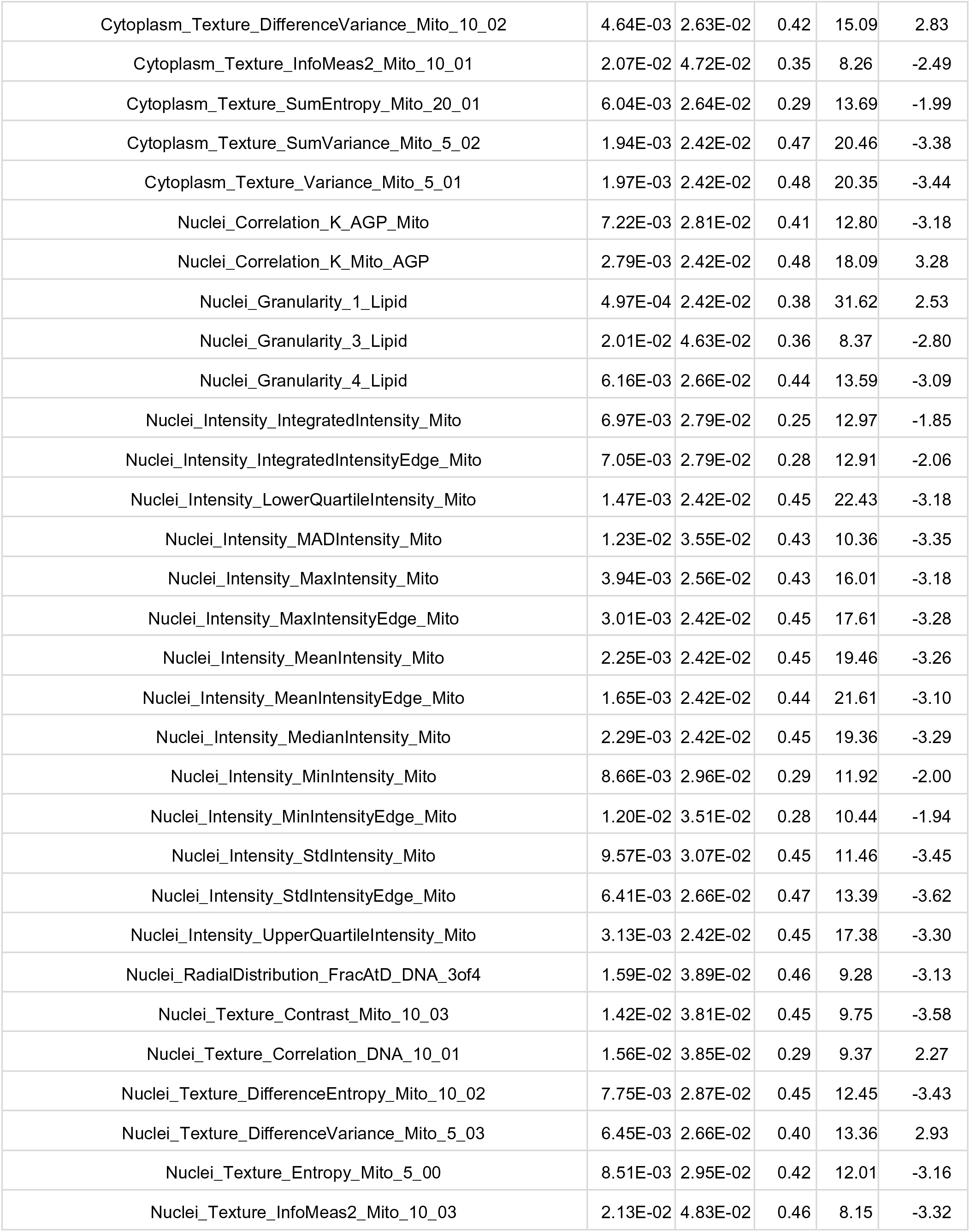
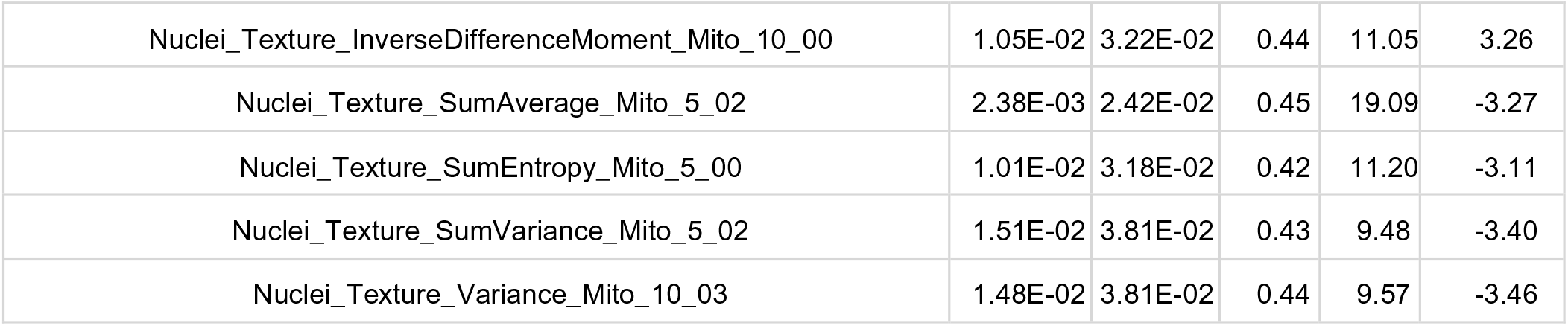
Significant effects on LP profile for 2p23.3 lipodystrophy locus in visceral AMSCs at day3 and day14 of differentiation. (ANOVA adj. BMI, sex, age, batch, significance level 5%FDR). P-value, p-value of ANOVA, q-value, q-value of ANOVA, FDR; eta_sq, eta square of ANOVA, effect size; F value of ANOVA; t-statistics of t-test.

